# Recombination of repeat elements generates somatic complexity in human genomes

**DOI:** 10.1101/2020.07.02.163816

**Authors:** G. Pascarella, K. Hashimoto, A. Busch, J. Luginbühl, C. Parr, C. C. Hon, W. H. Yip, K. Abe, A. Kratz, A. Bonetti, F. Agostini, J. Severin, S. Murayama, Y. Suzuki, S. Gustincich, M. Frith, P. Carninci

## Abstract

Millions of Alu and L1 copies in our genomes contribute to evolution and genetic disorders via non-allelic homologous recombination, but the somatic extent of these rearrangements has not been systematically investigated. Here we combine short and long DNA reads sequencing of repeat elements with a new bioinformatic pipeline to show that somatic recombination of Alu and L1 elements is common in human genomes. We report new tissue-specific recombination hallmarks, and show that retroelements acting as recombination hotspots are enriched in centromeres and cancer genes. We compare recombination profiles in human induced pluripotent stem cells and differentiated neurons and show that neuron-specific recombination of repeat elements accompanies chromatin changes during cell-fate determination. Finally, we find that somatic recombination profiles are altered in Parkinson’s and Alzheimer’s disease, indicating a link between retroelements recombination and genomic instability in neurodegeneration. This work shows that somatic recombination of repeat elements contributes massively to genomic diversity in health and disease.

## Introduction

Alu and Long Interspersed Nuclear Element-1 (LINE-1, abbr. L1) are the two most abundant retrotransposons in humans, with ∼1.2 and ∼1 million annotated copies that together account for almost 30% of the genome (Smith et al.). Key discoveries in recent years have transformed our view of these genomic repeats from inert fossils to evolutionarily co-opted symbionts exerting important functions in chromatin and gene regulation (Chuong et al., 2016; Daniel et al., 2014; Jachowicz et al., 2017; Pontis et al., 2019; Zhang et al., 2019); Alu and L1 can also directly affect genomic integrity via retrotransposition and recombination. Each genome hosts a variable number of young L1 and Alu copies that are competent for the copy-and-paste retrotransposition cycle, but the activity of these elements is usually guarded in adult somatic tissues by several layers of surveillance (Beck et al., 2010; Faulkner and Billon, 2018; Goodier, 2016; Philippe et al., 2016; Tristán-Ramos et al., 2020). In contrast, recombination of repeat elements is not restrained by the same rules governing retrotransposition: any pair of the millions of Alu and L1 elements in the genome can be the substrate of different types of recombination, hence considered a major driving force in human evolution and disease (Gu et al., 2008; Robberecht et al., 2013; Sasaki et al., 2010; Sen et al., 2006). Repair of DNA damage has been proposed as the major trigger of recombination (Currall et al., 2013); the main molecular players of this process have been characterized by landmark studies in yeast and are well-conserved across species (Chapman et al., 2012; Savocco and Piazza, 2021; Scully et al., 2019). Repeat elements have a major role in homology-based repair of double-stranded DNA breaks, during which the regions flanking the break are usually converted to single strands by end resection, assembled with components of the repair complex and used to scan for homologous sequences to be used as a template (Piazza and Heyer, 2019a). If equal loci on sister chromatids or homologous chromosomes are available and are selected as donor, the resolution of the DNA break can be neutral; conversely, if homologous sequences of non-allelic loci are selected as donors, the repair leads to non-allelic homologous recombination (NAHR) (Chen et al., 2007; Piazza and Heyer, 2019b; Savocco and Piazza, 2021). NAHR can disrupt the genetic information causing aberrant phenotypes and repeat elements have often been found at the breakpoints of NAHR events associated with cancer and other genetic disorders (Beck et al., 2011; Kolomietz et al., 2002; Zhang et al., 2009). Considering the substantial number of homologous Alu and L1 elements interspersed throughout the genome of each cell, two long standing questions in the field are how much genomic variation is generated by NAHR, and what is the contribution of different repeat element families to rearrangements caused by NAHR. Although several studies have sought to predict or measure the extent of NAHR and its contribution to diseases (Elliott et al., 2005; Morales et al., 2015; Parks et al., 2015; Startek et al., 2015; Song et al., 2018), a comprehensive investigation of somatic recombination of repeat elements in different cells, tissues or biological contexts is unavailable. Here, we comprehensively investigated NAHR of repeats in the human genome by pairing deep short- and long-read DNA sequencing with a new bioinformatic pipeline. We explored somatic NAHR in a panel of tissues from neurotypical donors and donors affected by neurodegeneration and we profiled recombination in a model of neuronal differentiation. Our work reveals new features of tissue-specific NAHR and show that the recombinogenic activity of Alu and L1 elements is an important contributor of genomic structural variants in normal and pathological conditions.

## Results

### High-efficiency capture and sequencing of Alu and L1 elements from genomic DNA

Considering the low prevalence of somatic structural variants associated with repeat elements (Erwin et al., 2016; Evrony et al., 2012, 2015, 2016) we optimized a riboprobe-based capture protocol (Fisher et al., 2011; Gnirke et al., 2009) to maximize the discovery power and enrich for genomic retroelement sequences prior to sequencing (“capture-seq”, see Supplementary Document 1). Since young repeats with higher similarity may recombine at higher rates compared to older elements we designed tiled DNA capture probes to span the full model sequences of young AluY elements (Hubley et al., 2016) and to cover ∼250bp of the 5’- and 3’-regions of the youngest L1 element consensus sequence (L1HS) (Baillie et al., 2011) (Supplementary Document 2). This design coupled to random shearing of genomic DNA allows for stochastic inclusion of uniquely mapping, non-repeated genomic regions flanking the captured repeats. The hybridization time of target and probes was reduced substantially from several days in previous capture-seq protocols iterations (Carreira et al., 2016; Shukla et al., 2013) down to 5 minutes. This allowed a 1-day library production time while retaining a low number of post-enrichment PCR cycles (n=12) and optimal enrichment efficiency (Figure S1).

We applied our capture-seq workflow to a panel of post-mortem tissues from 10 neurotypical donors (Table S1). For each donor, we selected available tissues derived from the 3 developmental germ layers: kidney (mesoderm), liver (endoderm) and 3 cortical brain regions (frontal cortex, temporal cortex, parietal cortex; ectoderm). Nuclear preparations from the brain samples were labeled with the neuron-specific antibody NeuN and underwent fluorescence-activated nuclei sorting (FANS) to separate the neuronal and non-neuronal fractions (Iwamoto et al., 2011; Matevossian and Akbarian, 2008) (Figure 1A, Figure S2, Supplementary Document 3). Paired-end sequencing of capture-seq libraries at 150bp yielded ∼960 millions raw reads (Table S2); quality control performed on uniquely mapping reads confirmed the consistent and efficient enrichment of a panel of reference L1HS sequences used as benchmark in previous studies (Carreira et al., 2016; Evrony et al., 2012) (Figure S1C). In protocols for enrichment of repeat elements based on capture probes, the repeated nature of the genomic targets overpowers the specificity of the probes. For instance, in our dataset we also detected a comprehensive and highly reproducible enrichment of L1 and Alu elements that were not originally targeted by experimental design. We took advantage of the richness and complexity of our capture-seq libraries by extending downstream analyses to all Alu (n=1,126,901) and primate-specific L1 subfamilies (n=122,626) (Table S3). The three Alu subfamilies, in order of increasing evolutionary age, are AluY, AluS and AluJ (Deininger, 2011). The median capture rate across the whole dataset was 94% for AluY elements, 83% for AluS elements and 45% for AluJ elements with an overall Alu capture rate of 75% ± 3% (Figure 1B). Primate-specific L1 subfamilies are classified as L1PA1-16, from most recent to oldest; L1PA1 (also known as L1HS) includes the only known autonomously active human retrotransposons (Khan et al., 2006). We divided the enriched L1 elements into 3 groups. As expected, L1HS showed the highest capture rate (94%), followed by L1PA2-L1PA7 (85%) and L1PA8-L1PA17 (42%); the aggregate capture rate for L1 elements was 64% ± 2% (Figure 1C). These data show that the capture-seq approach was able to enrich evenly in all libraries a majority of Alu and primate-specific L1 sequences annotated in the human genome.

**Figure 1:**
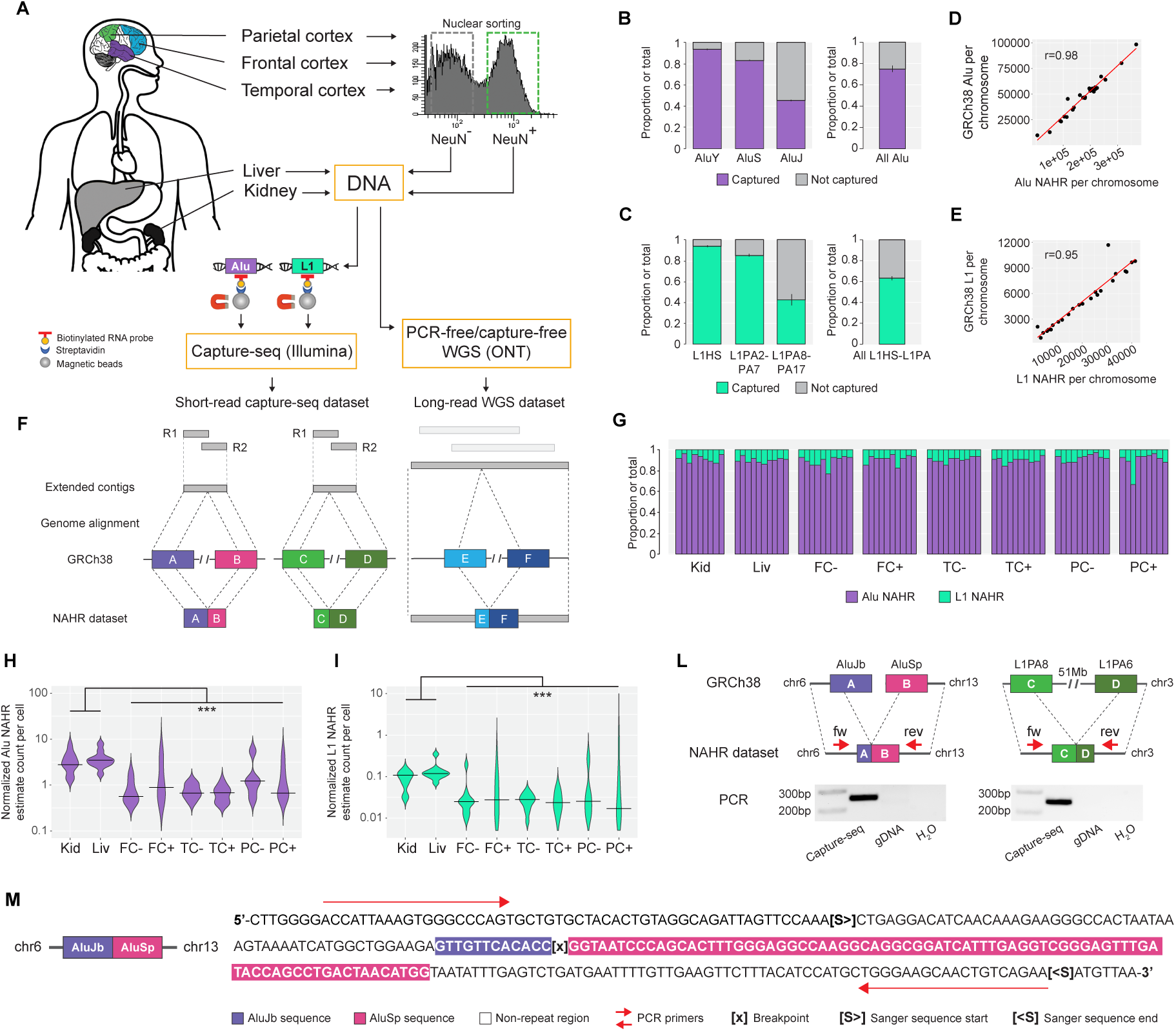
Experimental approach and annotation of somatic recombination of Alu and L1. (A) Schematics of the experimental setup for this study. Genomic DNA samples purified from bulk tissues and sorted nuclei are enriched for Alu and L1 elements by using biotinylated RNA capture probes spanning the entire sequence of young AluY elements and the 5-’ and 3’- regions of L1HS. Alternatively, a selection of samples is subjected to deep whole-genome sequencing on ONT PromethION platform without capture and PCR. (B, C) A majority of Alu and L1 elements annotated in the human genome is efficiently enriched in capture-seq libraries. Capture efficiencies for Alu (B) and L1HS/L1PA2-17 (C) elements are calculated for each library from the coverage of single-mapping reads on repeat elements annotated in the RepeatMasker database. (D, E) Correlation between the number of Alu NAHR events (D) and L1HS/L1PA2-17 (E) per chromosome found by TE-reX and the number of genomic Alu and L1 elements per chromosome annotated in the RepeatMasker database. (F) The TE-reX pipeline identifies NAHR from split-alignments of contigs with breakpoints that are mapped within repeats of the same family, and are located in homologous positions with respect to repeat model sequences. R1, paired read 1; R2, paired read 2. (G) Proportion of Alu and L1 NAHR events in capture-seq libraries. Each bar shows the relative proportion of Alu and L1 NAHR events in one donor. (H, I) Violin plots indicate the estimated counts and quartiles of Alu (H) and L1 (I) NAHR events per cell, annotated in capture-seq libraries of post-mortem samples from neurotypical donors; counts are normalized by amount of input DNA and sequencing depth. (L) Example of PCR validation for one inter-chromosomal Alu NAHR event and one intra-chromosomal L1 NAHR event identified in capture-seq libraries. PCR primers were designed on non-repeat regions flanking the recombined repeats for all targets. fw, forward primer; rev, reverse primer; gDNA, genomic DNA. (M) Example of Sanger validation for the AluJb-AluSp inter-chromosomal recombination in (L). For all panels: FC, frontal cortex; Kid, kidney; Liv, liver; PC, parietal cortex; TC, temporal cortex. PC, parietal cortex; +, neuron-specific antibody (NeuN) positive; –, NeuN, negative. *P < 0.05; **P < 0.01; ***P < 0.001 (Mann–Whitney U test).

### Discovery of genome-wide non-allelic homologous recombination of Alu and L1 elements with TE-reX

De-novo annotation of non-reference structural variants involving repeat elements in short DNA reads libraries is challenging since the repetitiveness and the genomic size of the rearrangements can be at or over the threshold of detection allowed by the short reads. To find NAHR events in Alu and L1 capture-seq libraries we developed TE-reX, a new bioinformatic pipeline based on LAST (Kiełbasa et al., 2011). TE-reX was designed to identify recombination events from split-aligned reads that join repeat elements at homologous positions (Figure 1F); to maximize the stringency of our analyses we relied solely on recombination events with low mismapping probability (p ≤ 1e10-5), reported as “mismap” in LAST alignments (Frith and Kawaguchi, 2015). TE-reX identified 2,131,372 Alu and 251,380 L1 NAHR events in all capture-seq libraries (Table S4). We flagged as “putative somatic” the recombination events identified in single capture-seq libraries (Alu=2,127,834; L1=250,127); conversely, NAHR events joining the same two repeat elements at the exact same base position respect to Dfam repeat models (Hubley et al., 2016) and found in multiple libraries were flagged as “putative polymorphic” (Alu=3,538; L1=1,253). In all libraries the number of NAHR events per chromosome was highly correlated with the number of repeats annotated per chromosome (Figure 1D-1E). The relative abundance of Alu recombination events per library exceeded that of L1 recombination events by several folds as expected from the ratio of Alu:L1 elements included in the analysis (∼9:1, Figure 1G). An estimate count of somatic NAHR events per cell based on the amount of DNA used per library and normalized by sequencing depth showed that the number of Alu and L1 recombination events per sample was higher in kidney and liver compared with brain, whereas we did not detect any difference between the neuronal and non-neuronal fractions (Figure 1H-1I).

We validated NAHR events identified by TE-reX in capture-seq libraries by polymerase chain reaction (PCR) followed by Sanger sequencing. To avoid cross-amplification of homologous repeat sequences, we selected for primers design only those recombination events where we could identify 5’- and 3’-ends non-repeat sequences flanking the recombined repeats in the mapped contigs (Figure 1L). Since across the capture-seq dataset only 1% of all NAHR events satisfied this requirement, at this stage we were not able to find putative polymorphic NAHR events suitable for stringent PCR primers design. For the putative somatic events group we designed primers flanking 112 NAHR events (79 inter-chromosomal, 33 intra-chromosomal); of these, 101 were supported by a single contig, 10 by 2 contigs and 1 by 4 contigs (Table S5). PCR reactions from the respective capture-seq libraries produced amplicons of the expected length for 103/112 targets (92%; Figure 1L and Figure S3) and Sanger sequencing confirmed the target identity of 93/103 targets (90%; Figure 1M, Table S5 and Supplementary Document 4). By definition, somatic structural variants are present only in a fraction of the cells down to a single cell in the tissue of origin, therefore the putative somatic NAHR events validated in capture-seq libraries are not expected to be validated by PCR also in the bulk genomic DNA samples. For 72 putative somatic targets we had sufficient material to attempt direct PCR amplification from the respective bulk DNA; we obtained amplicons of length comparable to the expected values for 17/72 targets (Figure S4A) and Sanger sequencing confirmed the identity of 6/17 amplicons, while the remainder were non-specific PCR products. To test whether these six targets were actually polymorphic events that might have been missed by TE-reX in some capture-seq libraries, we performed additional PCRs using as input the complete panel of eight genomic DNA samples available for each donor. For 5 out of 6 targets we obtained PCR amplicons of the same expected length in all related genomic DNA samples (Figure S4B); we therefore concluded that these NAHR events were true polymorphic, indicating a theoretical 7% (5/72) false positive rate across the somatic NAHR dataset. To confirm this estimate, we compared the genomic location and size of all putative somatic NAHR with that of non-reference polymorphic structural variants (SVs) aggregated in the Database of Genomic Variants (DGV) (MacDonald et al., 2014). This analysis has a few caveats, namely: 1) SVs featured in the DGV derive from studies using different sequencing technologies, hence the breakpoints of the SVs may lack consistency and to compensate for this issue we allowed a ±10% genomic size discrepancy between capture-seq NAHRs and DGV SVs; 2) there is no explicit indication of NAHR events in the DGV metadata, although at least one included study discriminates between insertions and deletions caused by repeat elements (Audano et al., 2019); 3) the comparison can be performed only for intra-chromosomal NAHR events, since the database does not include inter-chromosomal SVs. We found correspondence to SVs annotated in the DGV for 18,417 out of 541,068 intra-chromosomal putative somatic NAHR events (3.4%), a theoretical false positive rate comparable to the estimate by PCR. In comparison, out of 3202 intra-chromosomal putative polymorphic NAHR events we found overlapping and similarly-sized DGV SVs for 372 events (∼12%).

### Genome-wide profiling of Alu and L1 NAHR reveals tissue-specific characteristics of somatic recombination

Somatic NAHR events in brain samples were distinguished by a higher rate of intra-chromosomal events compared with kidney and liver samples; among the 3 tested cortical brain regions the temporal cortex samples had the highest rate of intra-chromosomal recombination (Figure 2A). A comparison of the intra-chromosomal recombination rate of each sample with random Alu and L1 pairs generated *in silico* confirmed that intra-chromosomal NAHR events were enriched in all chromosomes, and the enrichment was higher in brain samples versus kidney and liver samples (Figure 2B). We additionally observed that the relative abundance of somatic NAHR events involving young AluY and L1HS elements was overall higher in brain samples (Figure 2C). However, separating the recombination events involving young repeat elements in their intra-chromosomal and inter-chromosomal components revealed that only the inter-chromosomal recombination events exhibited tissue-specific differences, while the intra-chromosomal recombination rates of young elements were similar in all samples (Figure 2D). Next we explored the landscape of intra-chromosomal recombination by calculating the original genomic distance (“d”) between members of each pair of Alu and L1 elements that we found recombined in the dataset, based on their RepeatMasker annotation. We observed that the majority of intra-chromosomal recombination events were established between retroelements that are either proximal to (d<100 kb) or far away (d>100 kb) from each other, suggesting different NAHR mechanisms for close and distant repeats (Figure 2E). The distance profiles for kidney and liver samples were overall similar, with ∼95% of the intra-chromosomal recombination involving repeats distanced more than 100 kb. In contrast, somatic NAHR in brain samples showed a significantly higher abundance of proximal intra-chromosomal recombination; this was additionally more pronounced in the temporal cortex than the frontal or parietal cortex.

**Figure 2:**
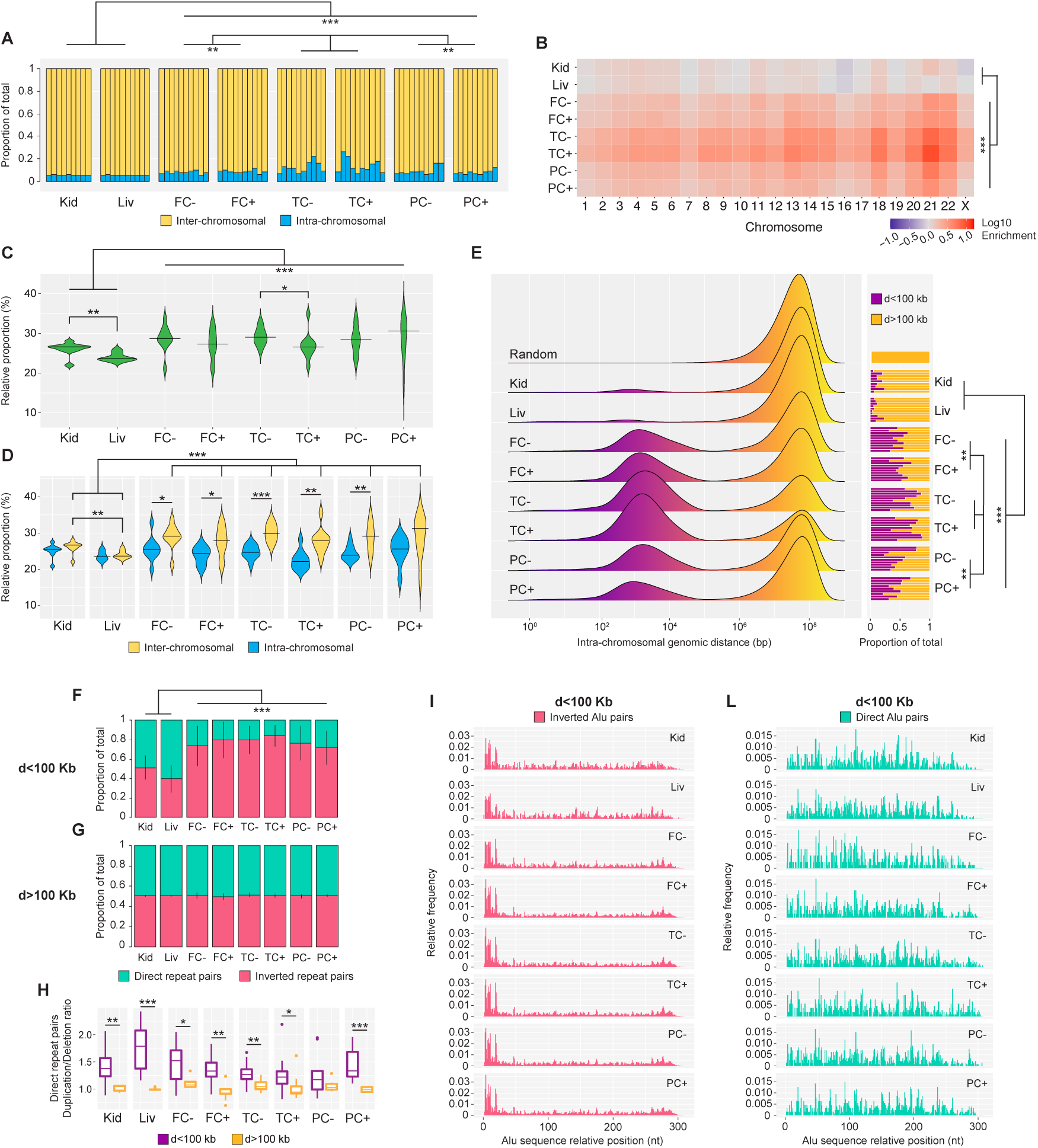
Genome-wide profiling of Alu and L1 NAHR reveals tissue-specific characteristics of somatic recombination. (A) Intra- and inter-chromosomal rates for somatic Alu and L1 NAHR events. Each bar shows the relative proportion of intra- and inter-chromosomal events in one donor. (B) Intra-chromosomal recombination rates for all chromosomes and all samples. Observed values for each chromosome were normalized by the expected values obtained from 100 random recombination dataset of sizes comparable to capture-seq dataset size. Plotted normalized values are in the Log10 scale. Chromosome Y was excluded from this analysis due to its low mappability score. (C) Violin plots of the relative proportion and quartiles of somatic recombination events involving young repeat elements (AluY and L1HS) indicate a tissue-specific contribution of young repeats to NAHR annotated in capture-seq libraries. (D) Violin plots of the relative proportion and quartiles for the contribution of young repeat elements from (C), separated in its intra- and inter-chromosomal components, indicating that the tissue-specificity of young repeats recombination is restricted to the inter-chromosomal recombination. (E) Profiling of genomic distances of repeats involved in somatic intra-chromosomal NAHR events shows a distinct enrichment of recombination of proximal repeat elements in all brain samples. “Random” is the distance profile for 1 million random intra-chromosomal repeat pairs. Side bar plots show the relative proportions of intra-chromosomal NAHR events involving repeat pairs distanced less or more than 100 kilobases. Each bar shows the relative proportion of intra- and inter-chromosomal events in one library. (F, G) Analysis of orientation for recombined repeat elements distanced less than 100 kilobases showed a strong bias in the brain samples for recombination of elements in inverted configuration (F). No orientation bias was observed for recombined pairs distanced more than 100 kilobases (G). (H) Box-and-whisker plots of the ratio between duplications and deletions and quartiles for NAHR of repeats in direct configuration indicates a bias for duplications for recombined repeats distanced less than 100 kilobases. (I, L) Breakpoints frequency displayed along Alu model sequence show different profiles for somatic NAHR events involving Alu elements distanced less than 100 kilobases in inverted (I) and direct (L) configurations. For all panels: d, genomic distance. For tissue and sample abbreviations, see Figure 1. For all panels: *P < 0.05; **P < 0.01; ***P < 0.001 (Mann–Whitney U test).

The human genome sequence is depleted of inverted proximal Alu pairs, an evolutionary consequence of the genomic instability of close Alu elements in this configuration (Lobachev et al., 2000; Stenger et al., 2001). In our dataset the recombination of inverted proximal repeat elements was more frequent than that of repeats in direct configuration, and this bias was stronger in brain samples compared with non-brain samples (Figure 2F). Moreover, the bias was dependent on the distance of recombined elements and it was undetected in recombined repeat pairs distanced more than 100 kb (Figure 2G). These observations confirm at the somatic level that close inverted repeats are indeed a source of genomic instability and that their activity is characterized by tissue specificity.

While recombination of intra-chromosomal repeats in inverted configuration can only generate genomic inversions, recombination of repeats in direct configuration can result in deletions and/or duplications. By comparing the order of each pair of recombined direct repeats in our dataset with their original genomic location we were able to distinguish NAHR events generating duplications or deletions; we found that recombination of proximal direct repeats was biased towards duplications consistently in all samples, while recombination of distal direct repeat pairs showed no bias between deletions and duplications (Fig 2H).

Previous analyses of Alu-mediated deletions in the human genome have reported an enrichment of breakpoints in the 5’-region of Alu sequences (Morales et al., 2015; Sen et al., 2006; Song et al., 2018). We detected a similar enrichment of breakpoints frequency in the 5’- region of all samples for recombination events involving intra-chromosomal Alu pairs separated by less than 100 kb, but the enrichment was exclusive to inverted Alu pairs (Figure 2I). Recombined direct Alu pairs in the same distance interval exhibited markedly different breakpoint profiles without 5’-enrichment (Figure 2L). The enrichment was also absent in the breakpoint profiles of distal Alu pairs, irrespective of their configuration (Figure S9A-S9B). Taken together, these data show that somatic NAHR of Alu and L1 is characterized by complex tissue-specific profiles suggestive of differential regulatory mechanisms and/or dynamics of NAHR in the assayed tissues.

### Genome-wide annotation of NAHR events identifies hot Alu and L1 elements acting as recombination hotspots

To delve deeper into the genomic distribution of somatic NAHR, we next investigated the enrichment and depletion of Alu and L1 NAHR events in gene and regulatory regions by comparing the real dataset (“O”, observed) to datasets comprising random permutations of the genomic coordinates of NAHR breakpoints (“E”, expected) (FigureS5-S6). We did not observe any substantial enrichment or depletion within the gene body, except for a mild depletion of L1 NAHR events in 3’ untranslated regions (UTR). L1 events, but not Alu events, were significantly enriched in transcription start sites (TSS) of genes. A similar trend of enrichment, but to a larger extent, was observed in promoters (Kundaje et al., 2015), in particular across the brain tissues (log2 O/E ratio 1.5 to 2.9). In contrast, we observed an overall mild but significant depletion of both Alu and L1 NAHR events within enhancers (Kundaje et al., 2015) across all tissues and samples (log2 O/E ratio –0.4 to –1.2, P < 0.05, two-tailed Student’s *t*-test). Intriguingly, separating the regulatory regions according to their cell-type and tissue activity (Kundaje et al., 2015) (Figure S7-S8) showed that the enrichment of L1 events in promoters was prominently attributed to the promoter active in stem cells, independent of the tissues in which the NAHR events were observed (Figure S8B). We did not observe consistent enrichment or depletion of NAHR events in regulatory regions active in the matched tissues (e.g. liver NAHR events in liver active promoters).

Alu and L1 elements have frequently been found at the boundaries of structural variants associated with genetic disorders, however the genomic prevalence of somatic NAHR hotspots in normal genomes has never been addressed systematically. To find and annotate recombination hotspots genome-wide, we merged the whole NAHR dataset and screened for Alu and L1 elements involved in recombination events recurring more than expected from random genomic distribution. After disjoining the Alu-Alu and L1-L1 pairing information we calculated the number of individual recombination events per each repeat (“recombination index”, RI) and we flagged as recombination hotspots (“hot” repeat elements) individual repeats with an RI exceeding the threshold of random genomic distribution. About 81% of all recombination hotspots were Alu (82,292/101,539) and the most represented subfamily was AluY (57,408/82,292, 70%). Overall, among all Alu subfamilies ∼40% of annotated AluY and AluS and ∼15% of AluJ copies were flagged as hot elements; the L1 subfamilies with the highest rate of hot elements were L1PA5, L1PA6 and L1PA7 (∼50% each) (Table S6). We observed that several hot Alu and L1 elements were proximal to centromeres as in the case of chromosome 21, where a high density of hot elements localized in a genomic region involved in translocations underlying Down syndrome (Shaw et al., 2008) (Figure 3A, green shaded area). To systematically rank genomic regions based on their local Alu and L1 recombination activity, we binned the genome in non-overlapping windows of 0.1Mb and we normalized the total RI of each bin by the number of annotated Alu and L1 per bin. The top 100 genomic bins with the highest normalized recombination activity (100/28885, 0.34% of all bins) were highly enriched in centromeric regions (69/100, 69%); the rate of centromeric regions in the remaining genomic bins was significantly lower (386/28785, 1.3%, p< 2.2e-16, Mann–Whitney U test) (Figure 3B-3C). Genomic bins with low recombination activity generally corresponded to regions with low mappability scores where structural variants may be difficult to detect relying solely on short DNA reads. The observed frequency of NAHR events with both recombined repeats mapped within or in proximity of centromeres (centromeres ± 1Mb, “C-C” in Figure 3D) was higher than expected in all brain samples while kidney and liver samples did not show any enrichment. In contrast, observed and expected rates for recombination events with only one of two recombined repeats within or in proximity of centromeres (“C-NC” in Figure 3D) were similar for all samples. Additionally, in both brain and non-brain samples recombined “C-C” repeat pairs were distinguished by a higher intra-chromosomal recombination rate in comparison with repeat pairs not in centromeric regions (“NC-NC”), with the brain samples showing the highest enrichment as expected also from previous observations (Figure 3E and Figure 2A). In summary, these data show that intra-chromosomal NAHR of Alu and L1 is enriched at human centromeres.

**Figure 3:**
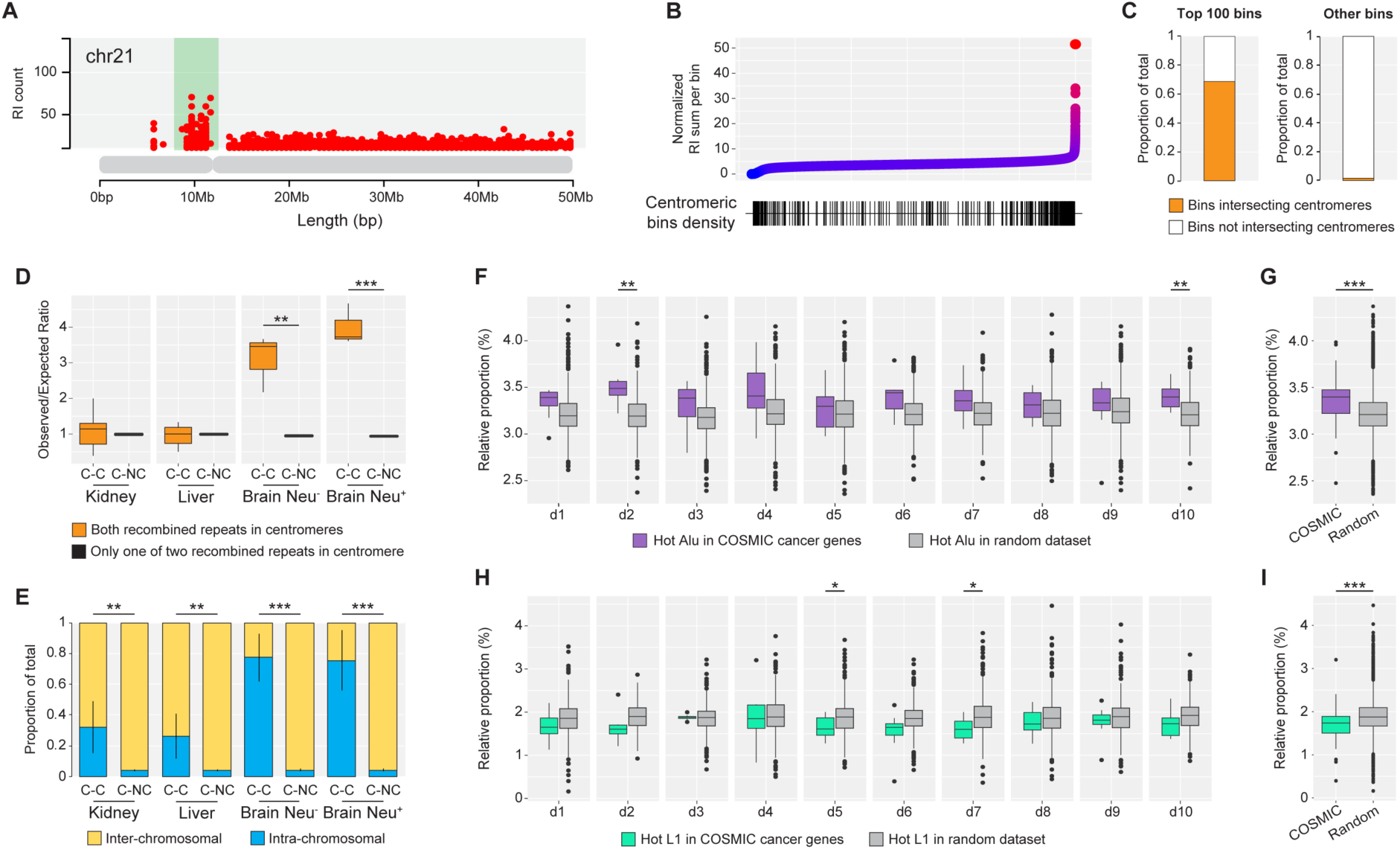
Genome-wide annotation of NAHR events identifies hot Alu and L1 elements acting as recombination hotspots. (A) Representative example of a cluster of hot repeat elements with RI≥10 (green shaded area) proximal to the centromeric region of chromosome 21. Each dot represents the genomic location of a single repeat element associated with its raw RI (y axis). (B) Centromeric regions are enriched in genomic bins with high relative recombination activity. GRCh38 bins (top panel, one dot = one 100 kilobases bin) are ranked by their relative recombination activity, calculated by dividing the sum of the RI of all repeat elements in a given bin by the total bin count of repeat elements annotated in the RepeatMasker. Genomic bins that overlap with centromeres ± 1Mb (lower panel) are marked with a vertical line in the bottom density plot. (C) Relative proportion of genomic bins intersecting centromeres ± 1Mb in the top 100 bins ranked by relative recombination activity from b) and in all remaining genomic bins. (D) Box-and-whisker plots of observed/expected ratio and quartiles for recombined repeat pairs with both repeats (“C-C”) or only one out of two repeats (“C-NC”) located within centromeres ± 1Mb. To compensate for overall low counts of NAHR events in centromeric regions, data of all brain samples were merged in NeuN^-^ and NeuN^+^ fractions. C-C, centromere-centromere; C-NC, centromere-non centromere. **P < 0.01; ***P < 0.001 (Two-way ANOVA). (E) Comparison of intra- and inter-chromosomal recombination rates for recombined repeat pairs with both repeats located within centromeres ± 1Mb (“C-C”) or in non-centromeric regions (“NC-NC”). For tissue and sample abbreviations, see c). **P < 0.01; ***P < 0.001 (Mann–Whitney U test). (F, G) Box-and-whisker plots of relative proportion and quartiles of hot Alu elements with RI ≥10 in cancer genes from the COSMIC database compared to relative proportion in random control datasets generated in silico (100 random datasets per donor. Panel (F): individual donors; panel (G): merged count for all donors. d1-10, donors 1 to 10. *P < 0.05; **P < 0.01; ***P < 0.001 (Mann–Whitney U test). (H, I) Box-and-whisker plots of relative proportion and quartiles of hot L1 elements with RI ≥10 in cancer genes from the COSMIC database compared to relative proportion in random control datasets generated in silico (100 random datasets per donor. Panel (H): individual donors; panel (I): merged count for all donors. d1-10, donors 1 to 10. *P < 0.05; **P < 0.01; ***P < 0.001 (Mann–Whitney U test).

Alu and L1 are responsible for recurrent mutations in several cancer types (Hastings et al., 2009; Kitada et al., 2013; Robberecht et al., 2013; Smida et al., 2017; Wang et al., 2019); in 2/10 donors and in the merged donors count, hot Alu elements (RI≥10) were enriched in oncogenes and tumor suppressor genes included in the COSMIC database (Sondka et al., 2018) compared to a random dataset generated from protein-coding genes not included in the COSMIC database (Figure 3F-3G). In contrast, hot L1 elements were not enriched in COSMIC genes and were depleted from COSMIC genes compared to the random background in the merged donors count (Figure 3H-3I).

### Differentiation of induced pluripotent stem cells to neurons triggers emergence of cell-specific recombination profiles

To gain insights into the origins of tissue-specific recombination profiles observed in post-mortem tissues, we exploited an *in vitro* model of neuronal differentiation (Liu et al., 2013). This protocol allows for the differentiation of induced pluripotent stem cells (iPSCs) into medial ganglionic eminence (MGE)-progenitor cells within 26 days, after which the specific induction towards GABAergic interneurons is started and prolonged for an additional 24 days (Figure S10). We applied our capture-seq workflow to 3 biological replicas of iPSCs and differentiated neurons (“iNEU”; induced from iPSCs) and paired-end sequenced the libraries on Illumina Miseq platform at 300 bp reads yielding a total of ∼32 millions of raw reads (Table S2). TE-reX analysis and annotation of NAHR events in iPSCs and iNEUs revealed that the differentiation triggered significant changes in the recombination profiles of the induced neurons. Although several features of Alu and L1 NAHR did not differ between iPSCs and iNEUs (Figure S11), the analysis of intra-chromosomal recombination distance intervals showed significantly higher recombination rates of proximal Alu and L1 pairs in iNEUs compared with iPSCs (Figure 4A), reminiscent of the differences observed in postmortem samples and suggesting that the recombination of Alu and L1 elements may accompany chromatin restructuring during cell-fate commitment. To test this hypothesis, we analyzed the recombination promoted by Alu and L1 elements in active (“A”) and inactive (“B”) chromatin compartments during the iPSC-to-neurons transition. Mammalian embryonic development has been associated with global rearrangements in chromatin structure (Dixon et al., 2015; Kishi and Gotoh, 2018); for example, neuronal differentiation from embryonic stem cells in mice results in reduced A-A contacts and an increase in B-B contacts (Bonev et al., 2017). Recent data has also shown that L1 and Alu sequences are involved in the organization of A and B compartments in human and mouse genomes, and that the segregation of L1 and Alu elements in defined nuclear compartments instructs genome folding during embryogenesis (Lu et al., 2021). Using the genomic coordinates of A and B compartments computed from high-resolution chromatin maps of iPSCs and induced neurons (Lu et al., 2020), in our iNEU dataset we observed a decreased recombination of Alu and L1 in A compartments mirrored by an increase of recombination in B compartments, compared to iPSC (Figure 4B-4C). Recombination of Alu-Alu pairs respectively located in A-A and A-B were decreased in iNEU, while the recombination rate of pairs in B-B was increased compared to iPSC (Figure 4D); the results for L1-L1 pairs recombination were similar although they did not reach statistical significance for the A-A and A-B combinations (Figure 4E). These results show that tissue-specific recombination profiles are dynamically established during early developmental stages and that 3D chromatin restructuring during cell-fate commitment is accompanied by significant changes in recombination profiles of Alu and L1 elements.

**Figure 4:**
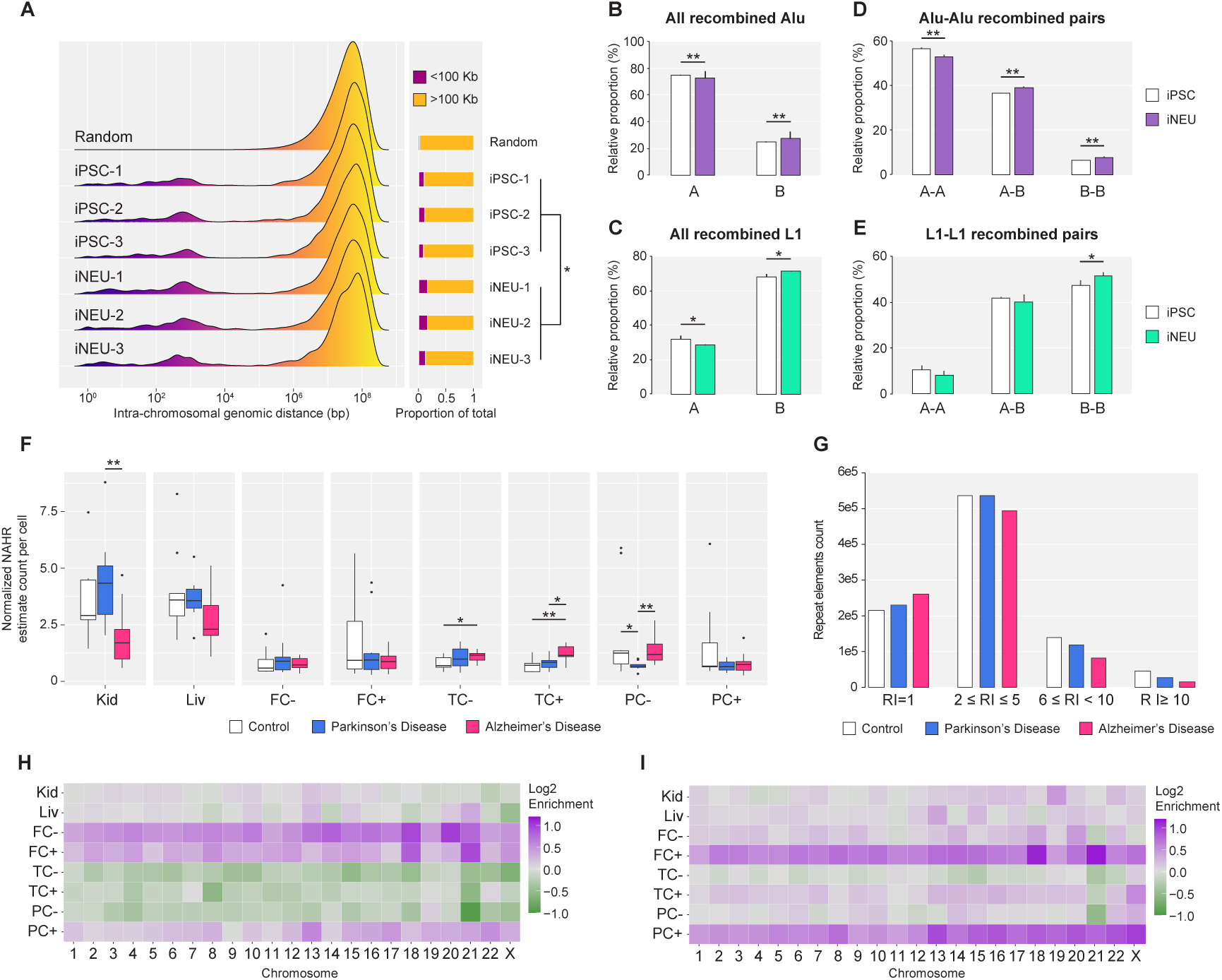
Alu and L1 recombination profiles are shaped by in vitro neuronal differentiation and are altered in neurodegeneration. (A) Genomic distances of repeats involved in intra-chromosomal NAHR events annotated in capture-seq libraries of human induced pluripotent stem cells and differentiated GABAergic cortical interneurons (iPSC and iNEU, three biological replicas for each). Side bar plots show the relative proportions of intra-chromosomal NAHR events involving repeat pairs distanced less or more than 100 kilobases. Each bar shows the relative proportion of intra- and inter- chromosomal events in one biological replica. d, genomic distance. *P < 0.05 (two-tailed Student’s *t*-test). (B, C) Relative proportion of Alu (B) and L1 (C) repeat elements involved in NAHR events annotated in iSPC and iNEU, located in A or B compartments from Lu et al., 2020. Bars show pooled values for 3 biological replicas ± SD. *P < 0.05; **P < 0.01 (One-way ANOVA). (D, E) Relative proportion of all combinations of recombined pairs of Alu (D) and L1 (E) elements in iPSC and iNEU, located in A or B compartments from Lu et al.(Lu et al., 2020). A-A, both repeats of a recombined pair in A compartments; A-B, one repeat in A and one repeat in B; B-B, both repeats of a recombined pair in B compartments. Bars show pooled values for 3 biological replicas ± SD. *P < 0.05; **P < 0.01 (One-way ANOVA). (F) Box-and-whisker plots of estimated counts of NAHR event per cell and quartiles, annotated in capture-seq libraries from post-mortem samples of control, Parkinson’s and Alzheimer’s disease donors; counts are normalized by the amount of input DNA and sequencing depth. For tissue and samples abbreviations, see Figure 1. *P < 0.05; **P < 0.01; ***P < 0.001 (Mann– Whitney U test). (G) Comparison of counts of cold and hot repeat elements involved in NAHR events annotated in capture-seq libraries of post-mortem samples from control, Parkinson’s and Alzheimer’s disease donors. Each bar indicates the count of elements with a given RI value or interval in the respective merged dataset. (H, I) Comparison of intra-chromosomal recombination rates for Parkinson’s (H) and Alzheimer’s disease (I) capture-seq datasets, and control donors capture-seq dataset. Colors show Log2 fold-enrichment of intra-chromosomal recombination rates for each individual chromosome compared with recombination rate values of respective samples in the control dataset. Data for chromosome Y not shown. For tissue and samples abbreviations, see Figure 1.

### Somatic recombination is altered in neurodegeneration

The observation that somatic NAHR is pervasive in normal physiological conditions provokes questions about how this tissue-specific, complex network of recombinations is affected in disease. We extended our investigation of NAHR of Alu and L1 to the two most common forms of neurodegeneration, sporadic Parkinson’s disease (PD) and sporadic Alzheimer’s disease (AD). We obtained post-mortem tissue samples with equal composition to the main dataset from an equal number of PD and AD donors (Table S1). Sequencing of 159 PD and AD capture-seq libraries yielded ∼960 and ∼990 millions of raw reads, respectively; quality control of PD and AD libraries showed capture efficiency and enrichment of Alu and L1 elements comparable to that of control donor libraries (Figure S1D-S1I, S12-S13). Genome-wide annotation and profiling of somatic NAHR events in PD and AD datasets recapitulated the main findings of the control dataset (Figure S9C-S9F, S14-S17). The estimate of normalized number of NAHR events per cell highlighted some relevant differences among the three groups, the most interesting being a significantly higher count of recombination events in the temporal cortex of AD donors (Figure 4F). A comparison of the recombination activity of individual repeats across the three datasets showed a consistent depletion of Alu and L1 hotspots in PD and AD respect to control donors, with AD samples having the lowest count of hot elements (Figure 4G). This striking depletion of recombination hotspots in the genome of AD and PD donors is of particular interest in light of the yet unexplained inverse relationship between these neurodegenerative disorders and cancer (Ording et al., 2019; Lanni et al., 2021). Moreover, the analysis of intra-chromosomal recombination rates in PD and AD revealed a significant enrichment of intra-chromosomal recombination specific for the NeuN+ fraction of parietal cortex samples, for both Alu and L1 NAHR, compared with the respective control samples. A similar result was observed also for the frontal cortex samples of AD, while in PD both the NeuN- and NeuN+ fractions showed an increase of intra-chromosomal NAHR compared with the control dataset (Fig 4H-4I). Collectively, these findings indicate that genome-wide NAHR profiles are affected in a cell- and tissue-specific fashion in neurodegeneration and underline key differences in the genomic distribution of somatic recombination events in PD and AD genomes.

### Recombination of repeat elements is confirmed in capture-free and PCR-free long-read whole-genome sequencing libraries

For an independent confirmation that somatic NAHR detected in capture-seq data is not a technical artefact generated by PCR or by any other step of the capture-seq workflow, we performed long-read whole-genome sequencing (WGS) of 26 DNA samples from 20 donors, including 20 temporal cortex samples (NeuN^+^ fraction, 10 control donors and 10 AD donors), 3 kidney and 3 liver samples (all from control donors) on the PromethION platform from Oxford Nanopore Technologies. This approach allowed us to investigate NAHR events in PCR-free and capture-free conditions, in the same genomic DNA samples used to generate the respective capture-seq libraries. The PromethION platform generated ∼270 millions high-quality reads with an average length of 7.7 kilobases and average genome coverage of 20-folds (Table S2). Since the preparation of these WGS libraries did not include enrichment of any repeat sequence, we expanded the search for NAHR to all repeat elements annotated in GRCh38. Overall, TE-reX identified 151,342 NAHR events across all 26 WGS libraries; the count of events per library was positively correlated with the number of reads per library (Figure S18A). A normalized count of NAHR events per million reads confirmed that the liver samples, but not the kidney samples, had a higher normalized count of recombination events per sample than the temporal cortex samples (Figure 5A). The vast majority of the NAHR events was from Alu and L1 (respective averages of 81% and 16%, Figure 5B), showing that other repeats have a minor NAHR contribution in the human genome (∼ 3%). Libraries with higher NAHR count had a higher complexity, underlined by a higher proportion of events found in single libraries; conversely, libraries with low NAHR count had higher rates of recombination events found in the same configuration in two or more libraries (Figure 5C). A comparison of all NAHR events found in PromethION WGS and in all capture-seq libraries returned 787 events found in the same configuration in both datasets, all of which flagged as putative polymorphic; this number represented the 0.5% and the 0.01% of the entire PromethION and capture-seq NAHR datasets, respectively (Figure 5D). The mean supporting reads count for these shared events was 31.4 across all PromethION libraries; in comparison, the mean supporting reads count for recombination events detected only by PromethION was 1.1. We did not find any putative somatic NAHR event detected in both respective PromethION WGS and capture-seq libraries: these observations indicate that the bulk of NAHR of repeat elements in the human genome is represented by rare somatic events with an extremely low copy number.

**Figure 5:**
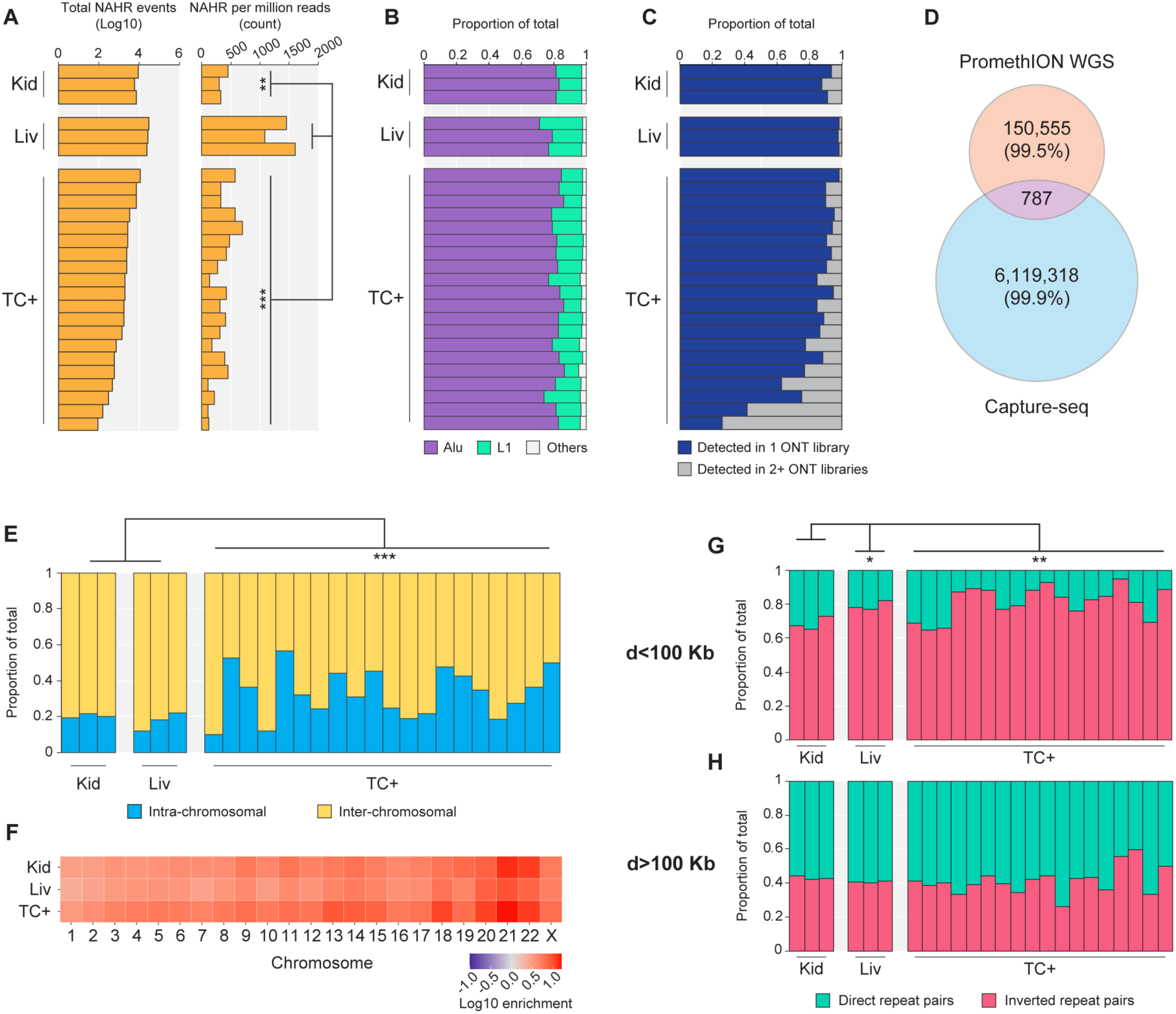
Recombination of repeat elements is confirmed in capture-free and PCR-free long-read whole-genome sequencing libraries. (A) Raw (left) and per-million-reads normalized counts (right) of NAHR events annotated by TE-reX in PromethION WGS libraries sequenced on Oxford Nanopore Technologies PromethION platform. Each bar represents the value relative to a single library. Kid, kidney; Liv, liver; TC+, temporal cortex, NeuN^+^ fraction. (B) Relative proportion of the main repeat elements families involved in NAHR events detected in PromethION WGS libraries. Each bar represents the values relative to a single library. Kid, kidney; Liv, liver; TC+, temporal cortex, NeuN^+^ fraction. (C) Relative proportion of NAHR events found in single or in 2+ PromethION WGS libraries. Each bar represents the values relative to a single library. Kid, kidney; Liv, liver; TC+, temporal cortex, NeuN^+^ fraction. (D) Overlap between PromethION WGS and capture-seq NAHR datasets. The comparison was performed by including all NAHR events detected in capture-seq libraries from control, Parkinson’s and Alzheimer’s donors in order to maximize the identification of putative polymorphic NAHR events in the PromethION WGS NAHR dataset. (E) Intra- and inter-chromosomal rates for NAHR events detected in PromethION WGS libraries. Each bar shows the relative proportion of intra- and inter-chromosomal events in one library. (F) Intra-chromosomal NAHR rates for all chromosomes in PromethION WGS data. Observed values for each chromosome were normalized by the expected values obtained from 10 in silico-generated random recombination dataset of sizes comparable to capture-seq dataset size. Plotted normalized values are in Log10 scale. Chromosome Y was excluded from this analysis due to its low mappability score. Kid, kidney; Liv, liver; TC+, temporal cortex, NeuN+ fraction. (G, H) Relative proportion of repeat pairs found recombined in PromethION WGS data in direct or inverted configuration and distanced less (G) or more (H) than 100 kilobases. Each bar shows the relative proportion of repeat pairs in direct or inverted configuration in one library. d, genomic distance. Kid, kidney; Liv, liver; TC+, temporal cortex, NeuN^+^ fraction. For all panels: *P < 0.05; **P < 0.01; ***P < 0.001 (Mann–Whitney U test).

Out of the 787 NAHR events found in PromethION and capture-seq data, 751 were intra-chromosomal, and for 225/751 (30%) we found comparably-sized SVs annotated in the DGV. In comparison, we found matching DGV SVs for 1222 out of 30,570 intra-chromosomal NAHR events detected in single PromethION WGS libraries (4%), a theoretical false positive rate close to that calculated for putative somatic NAHR events in the capture-seq dataset (3.4%). While capture-seq data did not provide sufficient sequence information to design PCR primers for putative polymorphic NAHR, the long DNA reads in PromethION WGS data expanded the genomic windows surrounding each NAHR event. We designed primers for 20 putative polymorphic NAHR events identified by both approaches; PCR from representative bulk DNA samples produced clean amplicons for 17/20 targets (85%, Figure S19A), and capillary sequencing confirmed the sequence identity of 16/17 amplicons (94%, Table S5 and Supplementary Document 4). For 13 of these verified targets we performed additional PCR from the panel of eight samples available for a single representative donor per target; the amplification of equal PCR products in all 8 samples confirmed the polymorphic status of these targets (Figure S19B).

PromethION WGS and capture-seq are distinguished by significant technical differences including the capture of repeat elements, PCR, reads length, the sequencing depth and sequencing platform. Nevertheless, a genome-wide profiling of NAHR events annotated in the PromethION WGS dataset remarkably confirmed some of the properties of somatic NAHR observed in capture-seq data. The comparison of intra- and inter-chromosomal NAHR rates in WGS data revealed a significant enrichment of intra-chromosomal recombination in the brain samples compared to kidney and liver samples (Figure 5E); observed intra-chromosomal recombination rates were higher than expected in all samples, compared to a background of random pairs (Figure 5F). The genomic distance of intra-chromosomally recombined repeats was characterized by a strong enrichment for recombination of proximal repeat elements, although this was similar for brain and non-brain samples (Figure S18B). In agreement with capture-seq NAHR, the recombination of intra-chromosomal proximal repeat was biased for pairs in inverted configuration (Fig 5G). Distal intra-chromosomal recombined repeats in WGS data showed a mild bias for pairs in direct configuration for all samples (Figure 5H). Moreover, although NAHR in WGS data was sparser compared to capture-seq data due to lack of enrichment and different sequencing depths, the analysis of normalized NAHR activity in individual genomic bins confirmed that the top 100 bins with the highest normalized RI count were enriched in centromeric regions compared to all other bins (31/100 vs 414/28778, p=4.262e-05, Mann–Whitney U test) (Figure S18C-S18D). In conclusion, the annotation of NAHR of Alu and L1 in somatic tissues by long DNA reads corroborated the discovery of NAHR events by TE-reX in capture-seq data. These data demonstrate that somatic NAHR of repeat elements can be effectively investigated in long-read WGS data and set a reference for future studies of somatic NAHR in health and disease.

## Discussion

One of the most surprising discoveries in the first draft of the human genome sequence was that roughly half of the sequenced bases belonged to repeat elements, for long considered as inert remnants of our evolution. Evidence accumulated during the intervening 20 years has changed this narrative, as we’re now aware that repeats are responsible for genomic diversity and biological mechanisms fundamental for life. Here, we show that somatic mosaicism caused by recombination of Alu and L1 in the human genome is extensive and complex, adding a new page to the developing story of how a myriad of genomic variants coexist in the same individual. A conservative estimate from our capture-seq libraries of neurotypical donors adjusted by the PCR validation results is that there are ∼4 and ∼1.4 NAHR somatic events per cell in non-brain and brain tissues respectively. These figures are coherent with estimates of somatic retrotransposition in the brain (Erwin et al., 2016; Evrony et al., 2016) and suggest an important contribution of somatic NAHR of Alu and L1 to genome diversity; however, dedicated and more sensitive technical approaches will be required to confirm our data at the single-cell level. Besides being pervasive, somatic NAHR profiled in capture-seq data exhibits tissue-specific characteristics that distinguish the brain regions from other tissues assayed; brain-specific NAHR is characterized by a higher rate of intra-chromosomal recombination and by higher recombination rates between close repeat elements. In addition, close-range recombination in the brain exhibits a strong orientation bias in favor of repeats in inverted configuration. These differences could stem from several factors, including the preferential usage of different DNA repair pathways, chromatin architecture, epigenetic modifications and developmental differences. Our analyses of recombination in iPSC and differentiated neurons suggest that the specific recombination profiles observed in post-mortem samples may be generated in progenitor cells during early developmental stages; this notion is supported by the homogeneity of intra-sample profiles and in the lack of fundamental differences in the NeuN- and NeuN+ fractions in the control donor samples. It is nevertheless an open question whether tissue-specific recombination profiles are a cause or consequence of cell-fate determination. Moreover, our data shows that neuronal differentiation is accompanied by distinctive dynamics of repeats recombination in A and B compartments. Since the differentiation of stem cells into a given progenitor cell type triggers specific changes in chromatin conformation (Bonev et al., 2017; Dixon et al., 2015; Lu et al., 2020), it is possible to imagine that the DNA damage occurring throughout this process engages the recombination of repeat elements and results in discrete patterns of recombination. Several lines of evidence have illustrated the existence of a complex relationship between DNA damage and neuronal development (Alt and Schwer, 2018). Repair of DNA double-strand breaks is instrumental for normal neurogenesis (Gao et al., 1998) and post-mitotic neurons retain the ability to express components of both the Non-homologous Ends Joining (NHEJ) and the homologous recombination pathway upon induction of DNA damage (Merlo et al., 2005), although at present it is unknown whether NAHR of repeat elements may participate in the repair of DNA DSBs generated during the activation of developmental programs and even by normal brain physiology (Stott et al., 2021). Similarly, under the assumption that DNA DSBs are the main trigger for NAHR of repeat elements, the specific profiles of intra-chromosomal recombination in Parkinson’s and Alzheimer’s brains may be the consequence of cell-type specific neurodegenerative processes causing differential DNA damage and cell-type specific alterations of recombination profiles. We additionally found that the genome of PD and AD donors is significantly depleted from NAHR hotspots compared to neurotypical donors and this may be consequential to complex alterations of both DNA damage and DNA repair in neurodegeneration (Madabhushi et al., 2014). More work will be required to understand the significance and the potential implications of neurodegeneration-specific alterations of NAHR profiles.

The annotation and analyses of NAHR in capture-free and PCR-free long-read WGS confirmed important genomic features of NAHR observed also in capture-seq data, regardless of the substantial technical differences between the two approaches. The cost of long-read sequencing has been steadily decreasing over time, however at present it is still prohibitive to sequence a large collection of samples at depths adequate to fully describe the complexity of somatic structural variants. Future studies will certainly be able to apply long-read technologies in a more comprehensive way and improve our understanding of tissue-specific NAHR. The extent of the repeats-driven recombination detected in short and long reads data poses the problem of compatibility of the observed rearrangements with genome fitness. Pending a precise count of NAHR frequency in single cells, somatic Alu and L1 NAHR events are responsible for a vast array of deletions, duplications, inversions and translocations. Considering the genomic extent of the observed rearrangements, inter- and intra-genic regulatory elements are likely to be affected by somatic NAHR events, with potential cell-specific effects on close and distant chromatin architecture and gene expression (Rigau et al., 2019). Spatial genome organization at single cell level is characterized by high variability (Finn et al., 2019); on the basis of its magnitude we speculate that somatic NAHR, as well as other structural variants coexisting in the same genomic environment, may contribute to this heterogeneity.

Seminal works on NAHR mediated by Ty elements in the yeast genome postulated that this class of rearrangements may be associated with evolution and disease in the human genome (Argueso et al., 2008; Hoang et al., 2010). Building on this hypothesis, we report here for the first time that somatic NAHR of repeat elements is abundant in normal human genomes and that NAHR hotspots are enriched in centromeres and cancer genes. We propose that stochastic somatic recombination of Alu and L1 may occasionally prime the genome of individual cells at vulnerable sites and drive the transition from healthy to pathological states. This scenario becomes even more plausible when considering possible complex interplay between different types of somatic mutation events; for example, a somatic retrotransposition event may be accompanied by a local genomic destabilization and consequent recombination if the newly inserted retroelement finds itself flanked by inverted homologous repeats (Gilbert et al., 2002). This is of particular interest also for estimates of the rate of retrotransposition in the human genome, because insertion of a young retroelement followed by recombination may create complex rearrangements masking the structural hallmarks of canonical retrotransposition events, namely target site duplications and a poly(A) tail, possibly resulting in underestimation of germ-line and somatic retrotransposition rates.

The evolutionary reason why genomes carry so many repeat element sequences is still a matter of debate (Brunet and Doolittle, 2015), but in all likelihood the answer is a complex, multifaceted one. Purely from a DNA repair perspective, the vast pool of interspersed homologous repeat elements may represent a universal “emergency kit” readily available to help stitching the DNA. Furthermore Alu, L1 and LTR (Long terminal repeat) sequences can form G-quadruplex structures that may help to stabilize complexes formed with components of the DNA double-strand break repair machinery (Hall et al., 2019; Lexa et al., 2014; Pessina et al., 2019). Our results further support the possibility that the repertoire of repeat elements may increase organismal fitness (Brunet and Doolittle, 2015) by increasing somatic diversity. It is conceivable that diversified metabolism of e.g. liver or neural cells is beneficial, much as genetic diversity of a microbial population can enhance its resilience (Giraud et al., 2001). The characterization of somatic recombination of Alu and L1 elements in this study paves the way to future experiments that will explore the dynamics of somatic NAHR events and their impact on the structure and function of our genomes.

## Acknowledgements

The authors are indebted to Eric Arner (RIKEN IMS) and Charles Plessy (OIST) for help revising the manuscript and useful comments.

## Authors contributions

P.C., M.F. and G.P. conceived the original idea and managed the project; G.P. and P.C. developed the capture-seq protocol and designed the experiments; M.F. developed the TE-reX pipeline and supervised the analyses of TE-reX data; G.P. and A.Bu. processed all samples and prepared the capture-seq libraries; G.P., K.H. and M.F. conceived and performed the computational analyses; J.L. performed the iPSC differentiation experiments; Y.S. and K.A. prepared the libraries and performed the sequencing on ONT PromethION platform; C.P. and Y.H.W. provided support for ONT and Illumina libraries preparation; C.C.H. performed computational analyses and provided insightful comments on data interpretation; A.K., F.A. and J.S. provided support for computational analyses; A.Bo. provided valuable support for interpretation of results; S.M. provided human post-mortem samples; S.G. provided critical feedback; G.P. produced the figures and wrote the manuscript with valuable contributions from M.F., A.Bo., F.A., K.H., P.C. and S.G.

## Supplemental figures and legends

**Figure S1:**
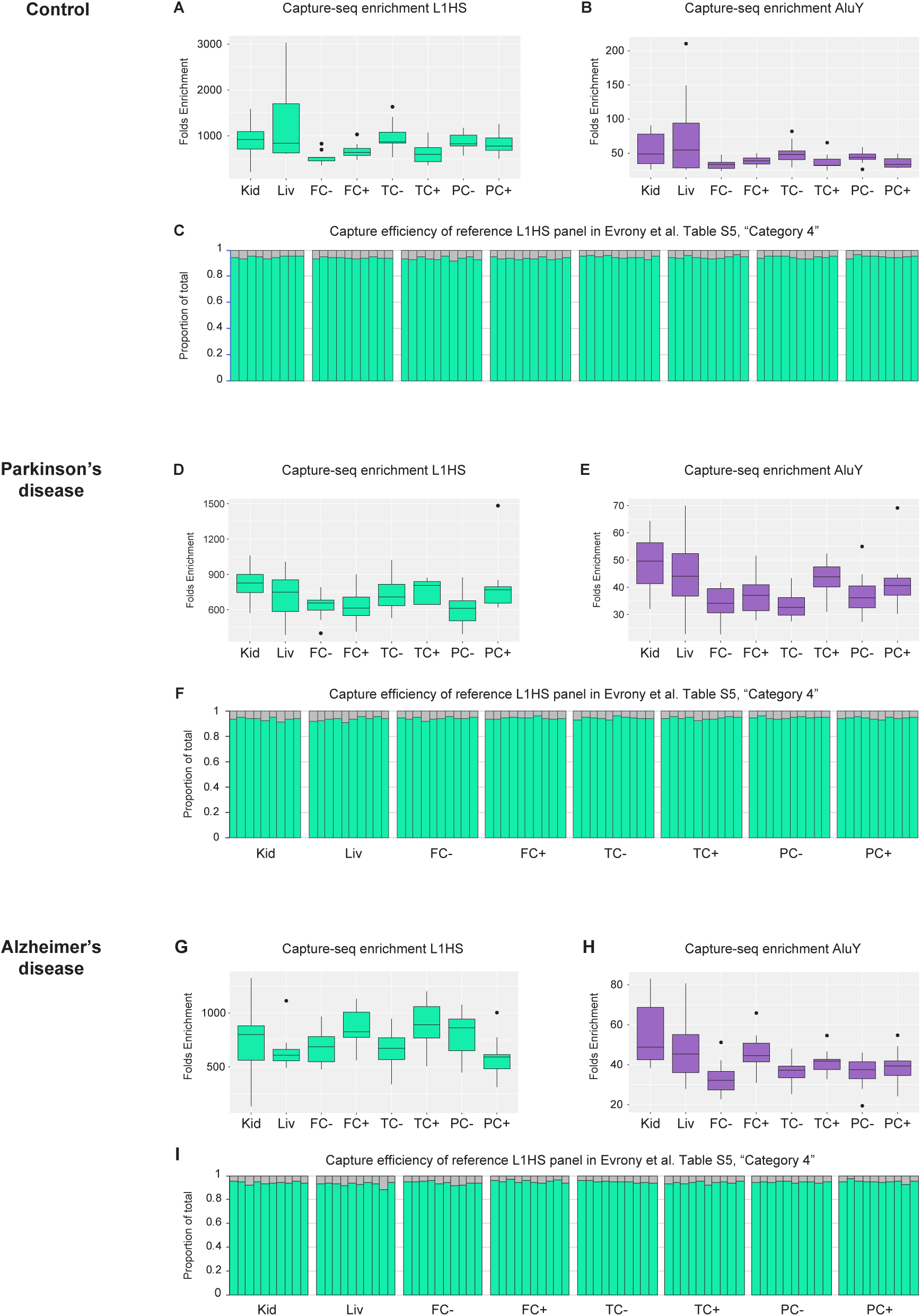
Enrichment efficiency in capture-seq libraries. (A, B) Box-and-whisker plots indicate the enrichment efficiency and quartiles measured by qPCR for L1HS (A) and AluY (B) elements in capture-seq libraries of control donor samples (C) Efficiency of enrichment in capture-seq libraries of control donor samples for a panel of 959 reference L1HS previously used to benchmark L1HS enrichment in previous targeted sequencing protocols(Carreira et al., 2016; Evrony et al., 2012). (D, E) Box-and-whisker plots indicate the enrichment efficiency and quartiles measured by qPCR for L1HS (D) and AluY (E) elements in capture-seq libraries of Parkinson’s disease donor samples (F) Efficiency of enrichment in capture-seq libraries of Parkinson’s disease samples for a panel of 959 reference L1HS previously used to benchmark L1HS enrichment in previous targeted sequencing protocols(Carreira et al., 2016; Evrony et al., 2012). (G, H) Box-and-whisker plots indicate the enrichment efficiency and quartiles measured by qPCR for L1HS (G) and AluY (H) elements in capture-seq libraries of Alzheimer’s disease donor samples (I) Efficiency of enrichment in capture-seq libraries of Alzheimer’s disease samples for a panel of 959 reference L1HS previously used to benchmark L1HS enrichment in previous targeted sequencing protocols(Carreira et al., 2016; Evrony et al., 2012). For all panels: Kid, kidney; Liv, liver; FC, frontal cortex; TC, parietal cortex; PC, temporal cortex. + and –, NeuN+ and NeuN- fractions of the brain tissues.

**Figure S2:**
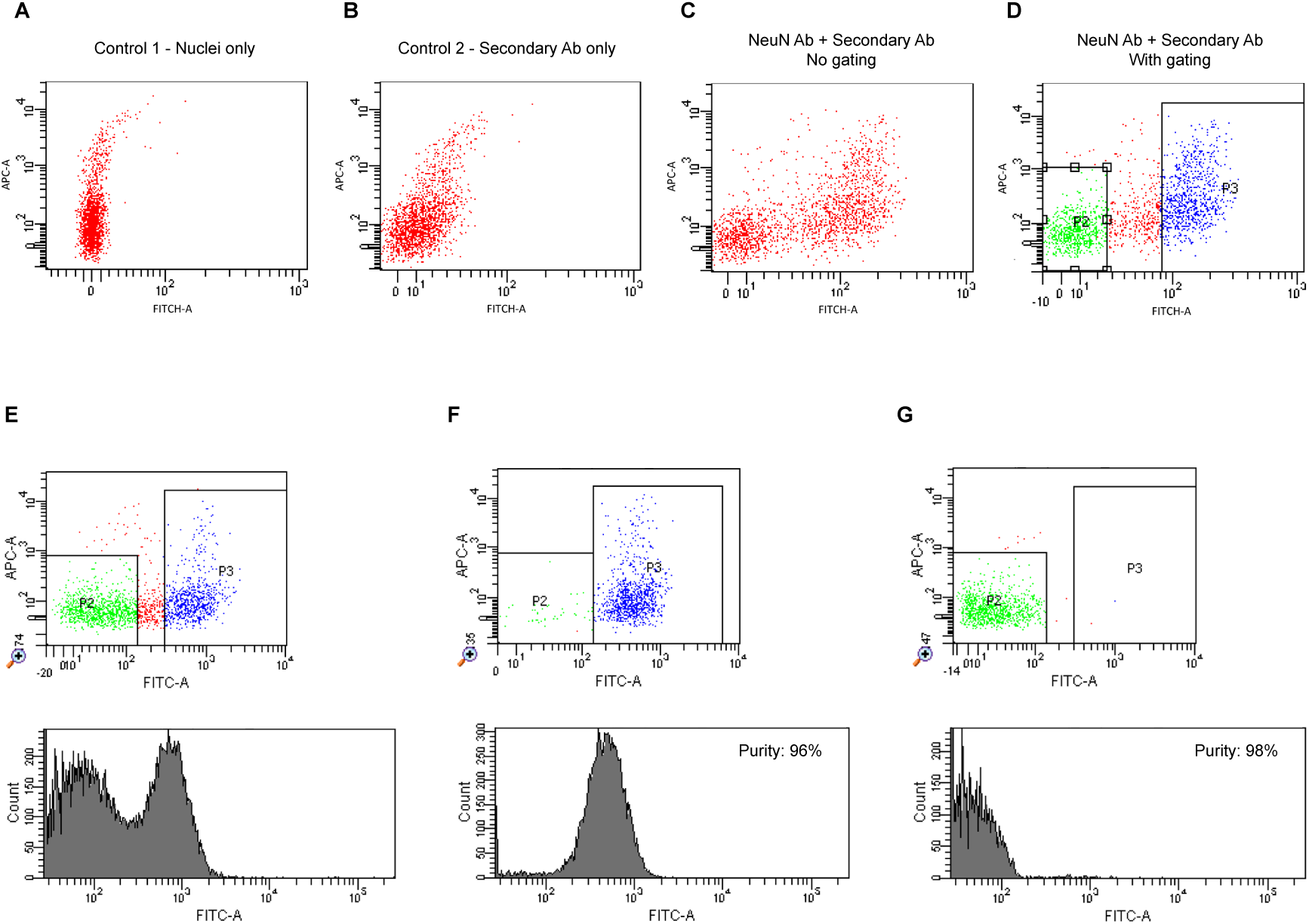
Sorting of nuclei stained with NeuN. (A) Illustrative baseline signal for a preparation of nuclei purified from post-mortem frozen bulk tissue, no staining. (B) Illustrative background signal given by the secondary antibody. (C) Illustrative signal for nuclei stained with NeuN and secondary antibody, before applying gating. (D, E) Illustrative examples of gating on NeuN signal for two different nuclei preparations. The positive and negative gates are set wide apart to avoid collection of low-signal NeuN+ tail and maximize the purity of specific fractions. (F, G) After-sorting purity check for nuclei preparation in (E). NeuN- and NeuN+ fractions show high purity and minimal cross-contamination. For all panels: APC, Allophycocyanin; FITC, Fluorescein isothiocyanate.

**Figure S3:**
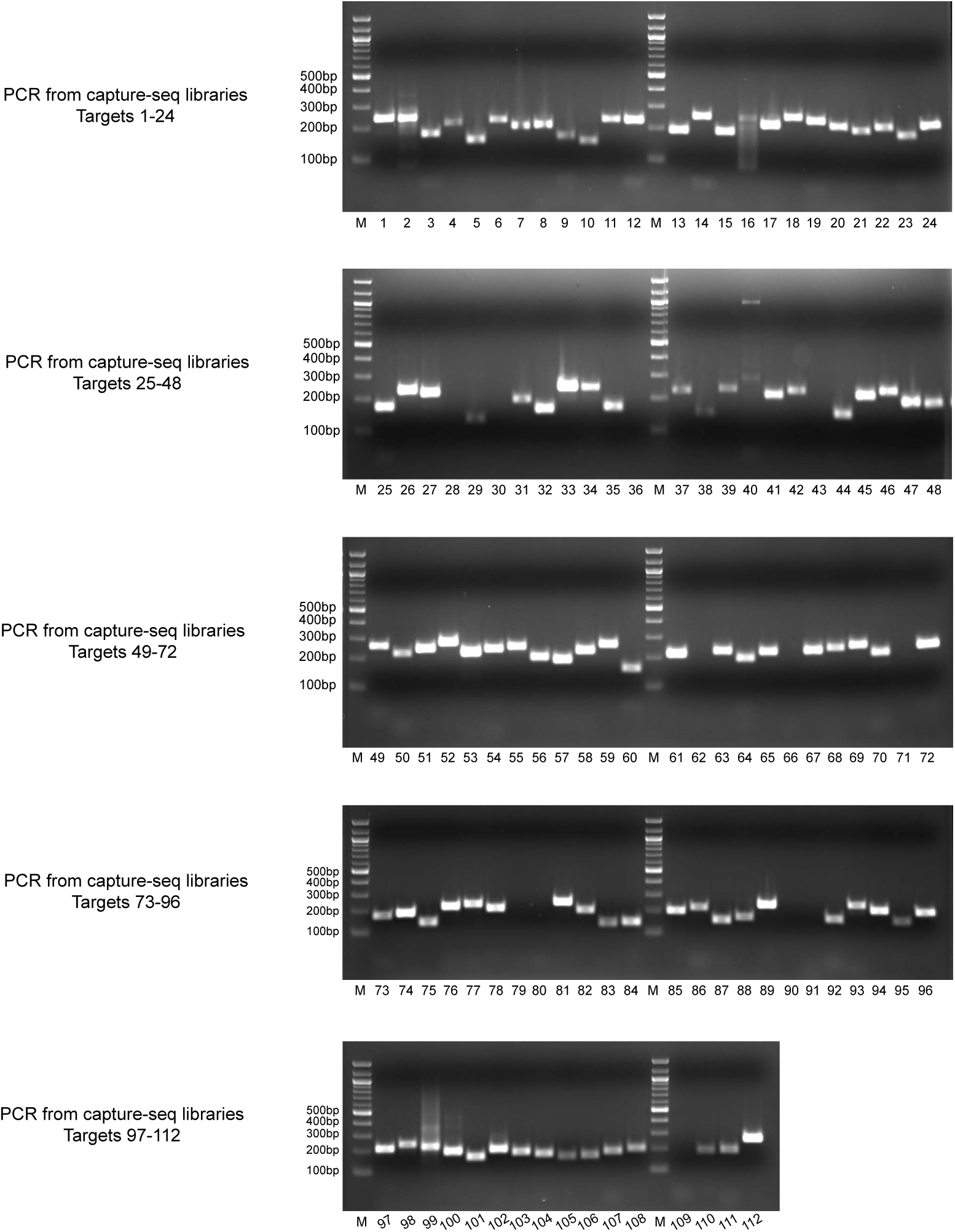
PCR validation of putative somatic NAHR events from capture-seq libraries. PCR amplicons for all targets of the PCR validation from capture-seq libraries on standard 1.5% agarose gel stained with ethidium bromide. Some of the targets presenting multiple bands or low signal underwent PCR optimization and were separately re-amplified prior Sanger sequencing. Extended details for all targets are in Table S5.

**Figure S4:**
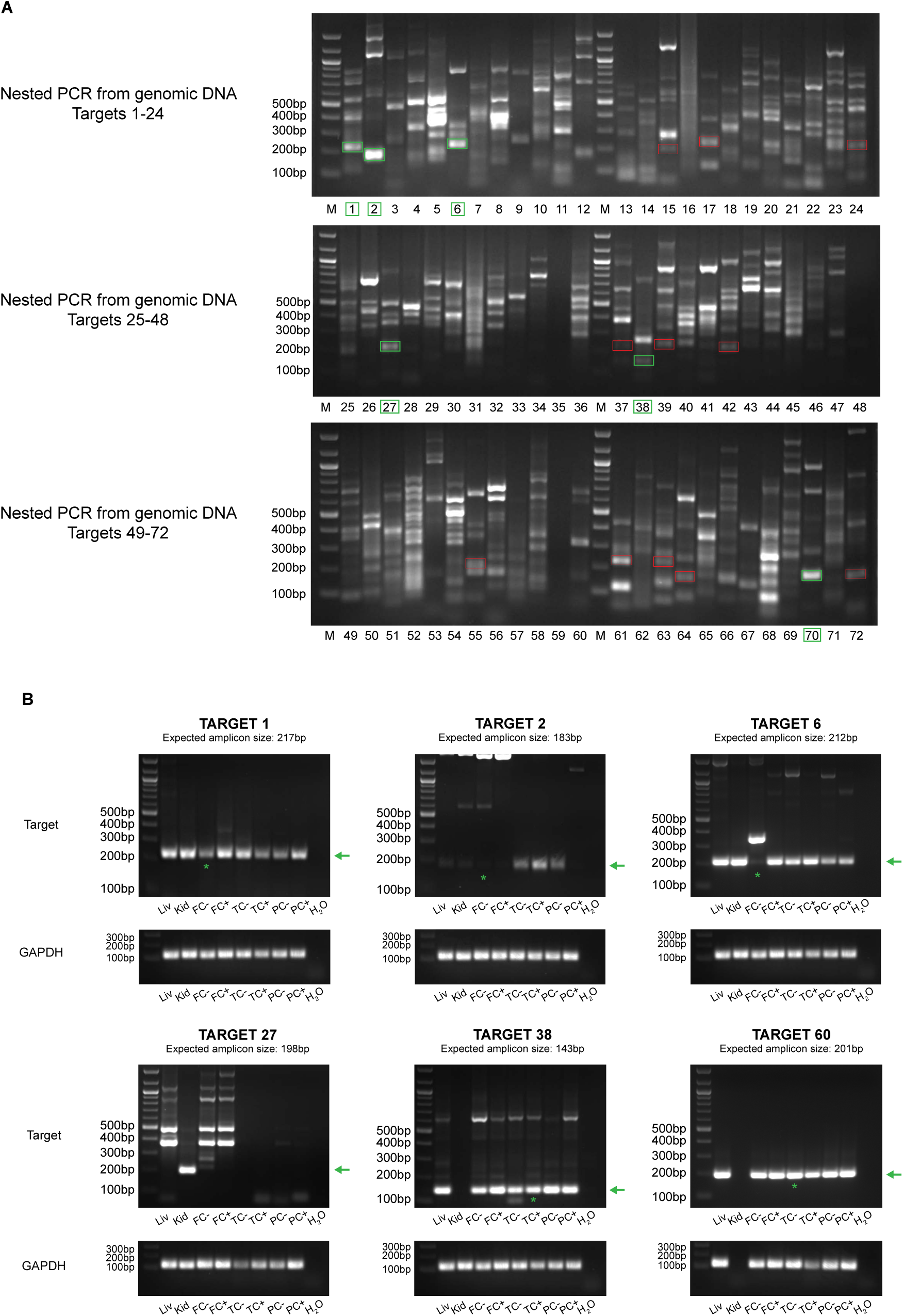
PCR from genomic DNA samples. Selected targets from Figure S3 for which genomic DNA material was available were used to verify the somatic state of the NAHR events. (A) Nested PCR amplicons for 72 selected targets from genomic DNA samples. Amplicons in the green and red boxes had size compatible to expected amplicons according to primers design and were tested by Sanger sequencing. Sanger sequencing confirmed the identity of the 6 targets in the green boxes. (B) The somatic state of the 6 targets validated by Sanger sequencing in (A) was tested by amplifying again the same targets with the same PCR primers and exact same experimental conditions as above from the panel of 8 genomic DNA samples available for each donor. GAPDH was amplified as positive control from the same input DNA. For targets 1, 2, 6, 38 and 60 we observed comparable amplicons of the expected size (green arrows) for all genomic DNA samples tested, hence supporting the polymorphic state of the assayed targets.

**Figure S5:**
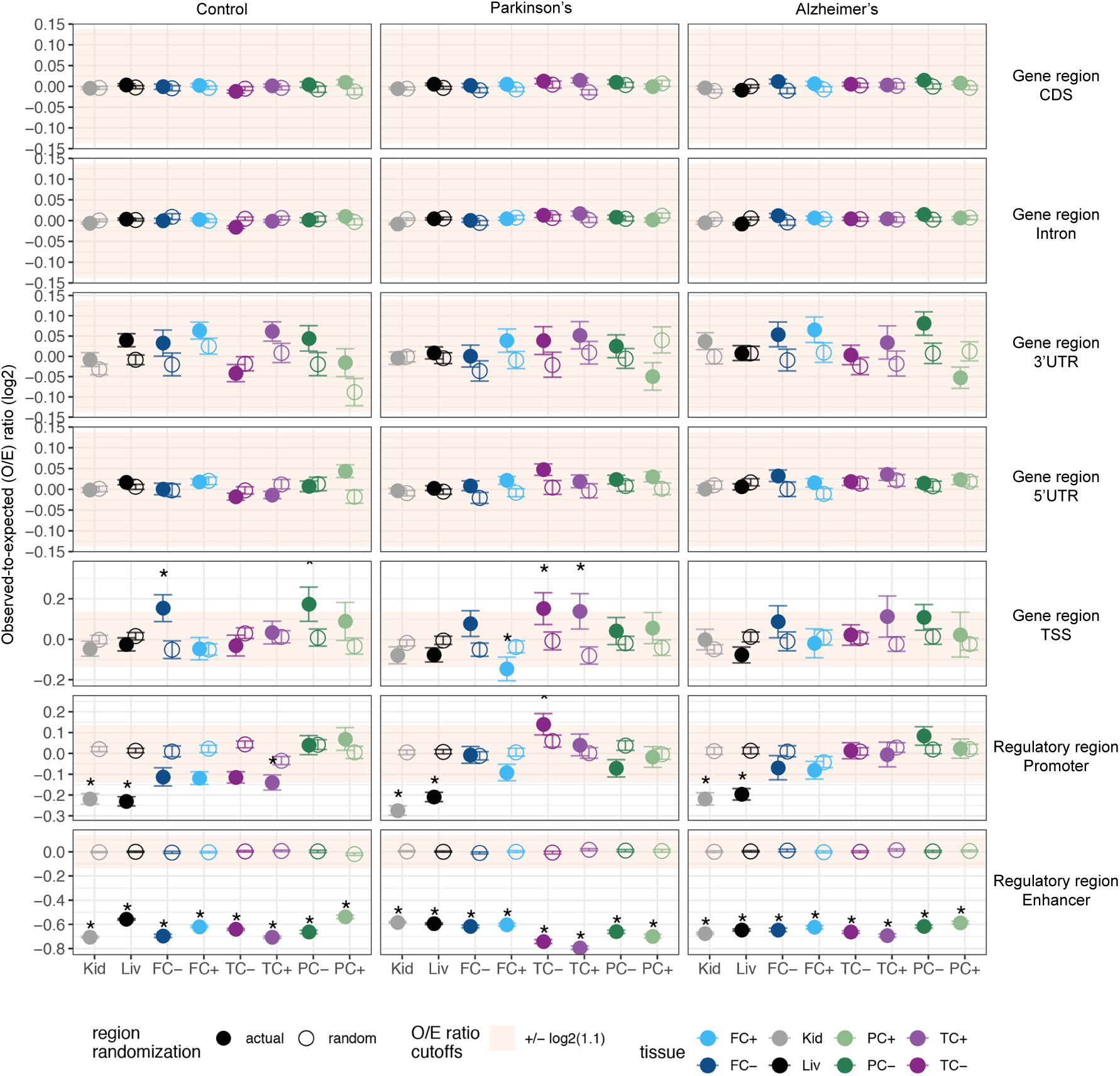
Enrichment of Alu NAHR events in genomic features. Enrichment or depletion of breakpoints was measured as the ratio of observed-to-expected breakpoints (O/E ratio) within certain genomic features. Y-facet, genomic features. X-facet, donor groups. Y-axis, log2 O/E ratio, from 100 permutations of breakpoints. Solid and hollow circles, mean log2 O/E ratio of actual and random regions of the corresponding genomic features. Error bars, standard deviation. Shaded area, predefined O/E ratio cutoffs between 1.1 and –1/(1.1). Asterisks, O/E ratio beyond predefined cutoffs and P < 0.05, in Student’s t-test between actual and randomized regions. X-axis, tissues. Kid, kidney; Liv, liver; FC, frontal cortex; TC, parietal cortex; PC, temporal cortex. + and –, NeuN+ and NeuN- fractions of the brain tissues. CDS, coding sequence; UTR, untranslated region; TSS, transcription start sites.

**Figure S6:**
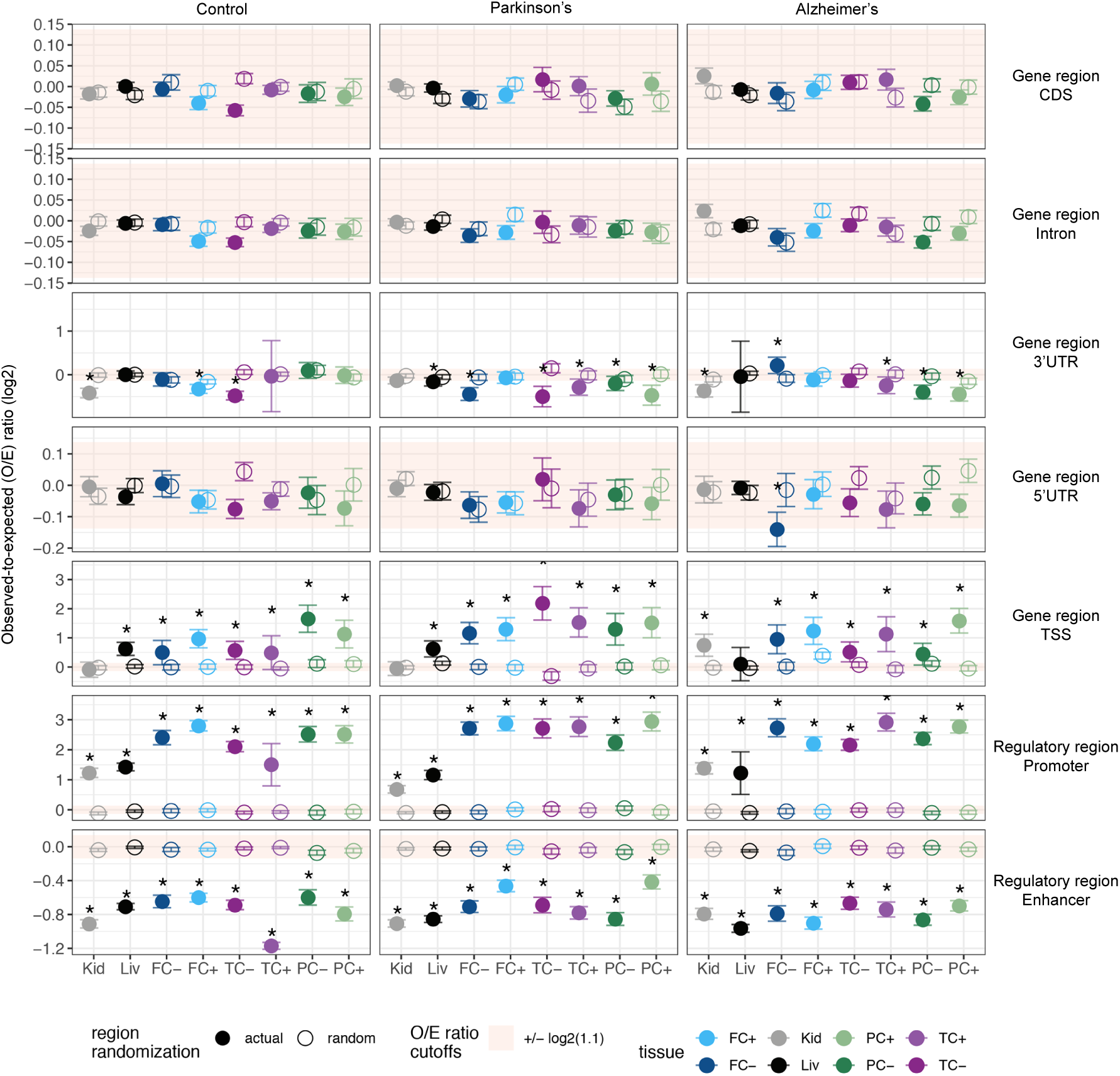
Enrichment of L1 NAHR events in genomic features. Enrichment or depletion of breakpoints was measured as the ratio of observed-to-expected breakpoints (O/E ratio) within certain genomic features. Y-facet, genomic features. X-facet, donor groups. Y-axis, log2 O/E ratio, from 100 permutations of breakpoints. Solid and hollow circles, mean log2 O/E ratio of actual and random regions of the corresponding genomic features. Error bars, standard deviation. Shaded area, predefined O/E ratio cutoffs between 1.1 and –1/(1.1). Asterisks, O/E ratio beyond predefined cutoffs and P < 0.05, in Student’s t-test between actual and randomized regions. X-axis, tissues. Kid, kidney; Liv, liver; FC, frontal cortex; TC, parietal cortex; PC, temporal cortex. + and –, NeuN+ and NeuN- fractions of the brain tissues. CDS, coding sequence; UTR, untranslated region; TSS, transcription start sites.

**Figure S7:**
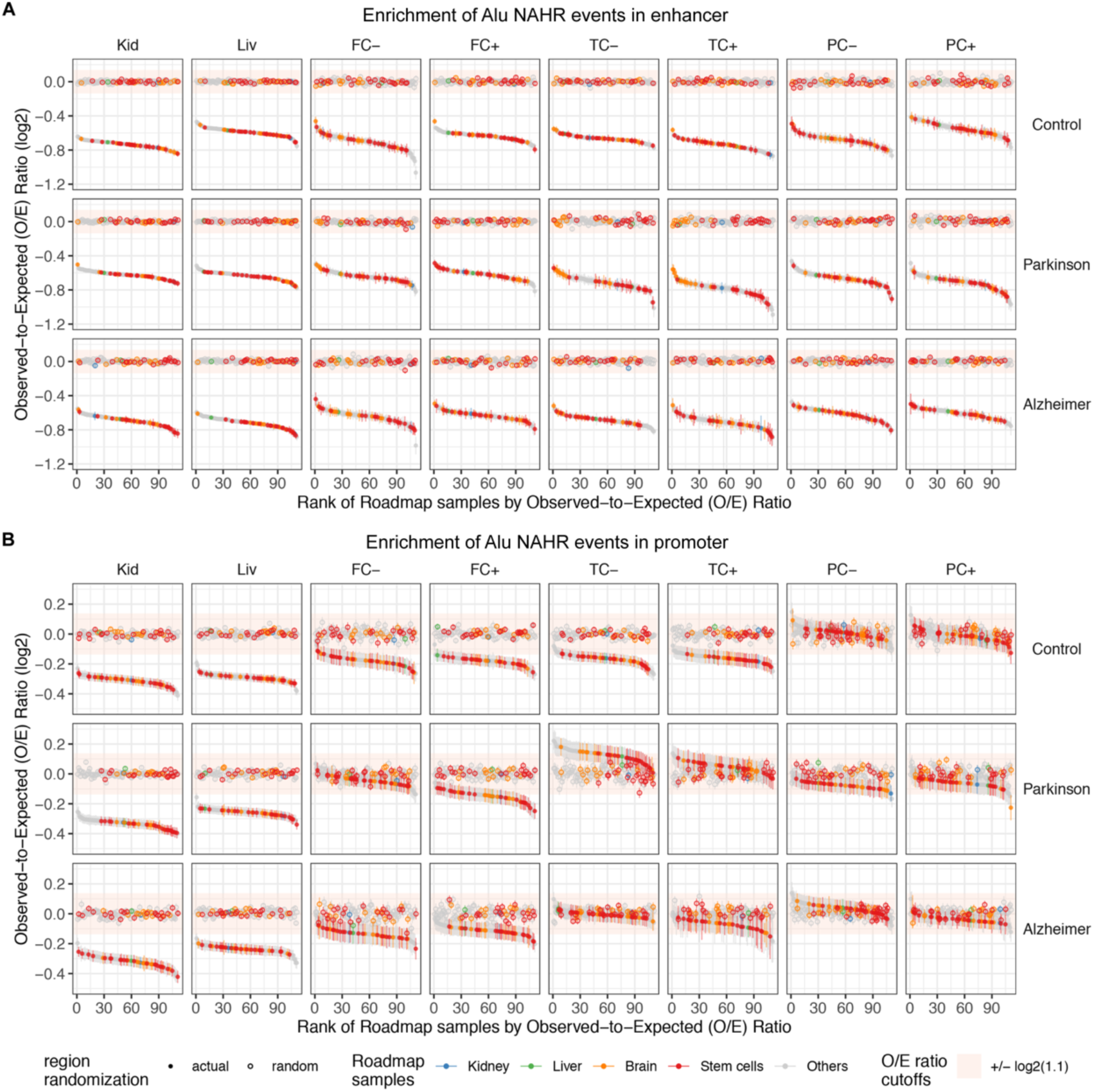
Enrichment of Alu NAHR events in regulatory regions active in various cell-types and tissues. Enrichment or depletion of breakpoints was measured as the ratio of observed-to-expected breakpoints (O/E ratio) within the regulatory regions that are active in certain Roadmap samples (n=111). a) Enrichment in enhancers; b) Enrichment in promoters. Y-facet, donor groups. X-facet, tissues. Kid, kidney; Liv, liver; FC, frontal cortex; TC, parietal cortex; PC, temporal cortex. + and –, NeuN+ and NeuN- fractions of the brain tissues. Y-axis, log2 O/E ratio, from 100 permutations of breakpoints. Solid and hollow circles, mean log2 O/E ratio of actual and random regions of the corresponding genomic features. Error bars, standard deviation. Shaded area, predefined O/E ratio cutoffs between 1.1 and –1/(1.1). X-axis, Roadmap samples, ranked by O/E ratio. Roadmap samples were grouped and colored as kidney, liver, brain, stem cell and others.

**Figure S8:**
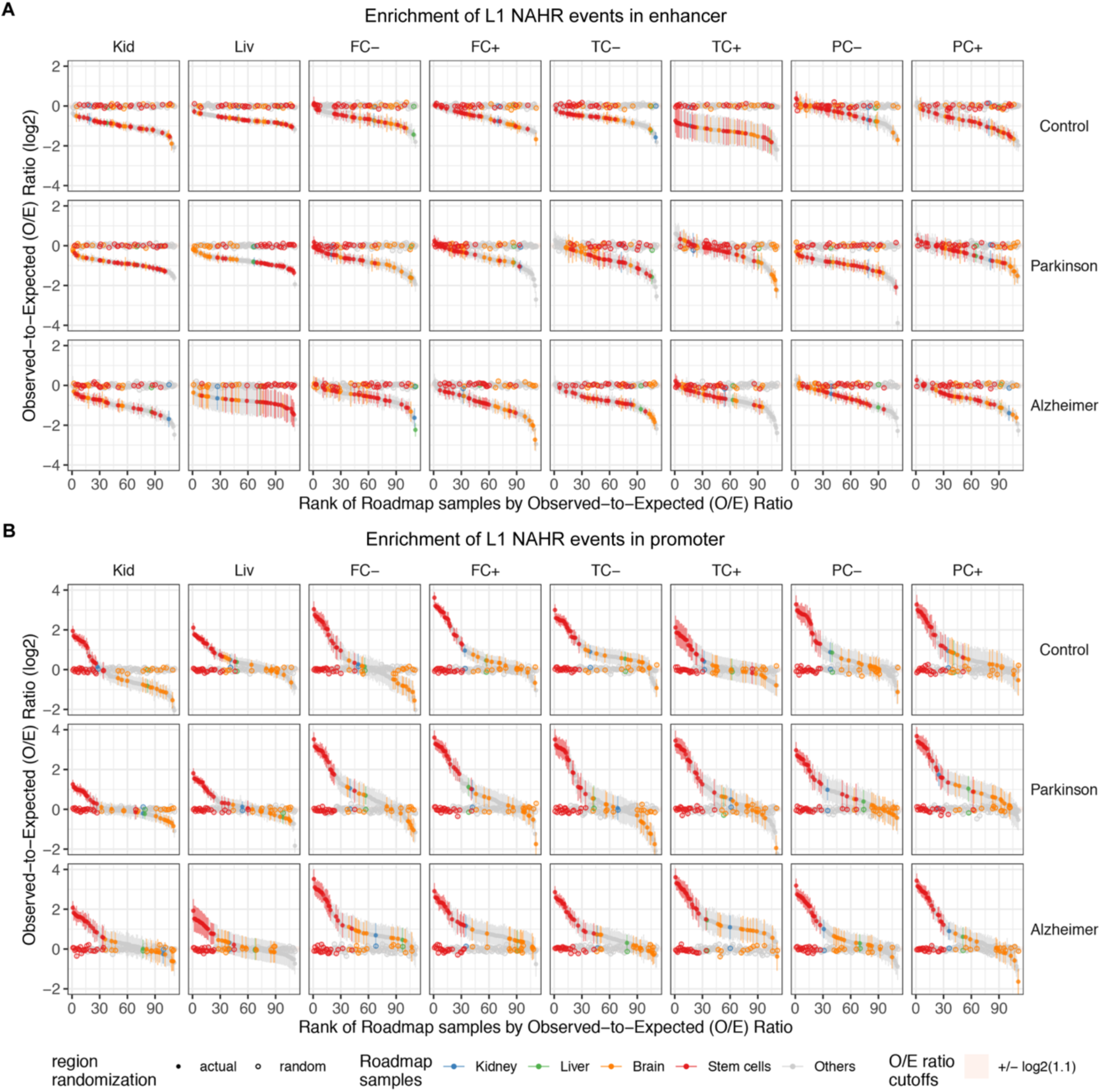
Enrichment of L1 NAHR events in regulatory regions active in various cell-types and tissues. Enrichment or depletion of breakpoints was measured as the ratio of observed-to-expected breakpoints (O/E ratio) within the regulatory regions that are active in certain Roadmap samples (n=111). A) Enrichment in enhancer; B) Enrichment in promoters. Y-facet, donor groups. X-facet, tissues. Kid, kidney; Liv, liver; FC, frontal cortex; TC, parietal cortex; PC, temporal cortex. + and –, NeuN+ and NeuN- fractions of the brain tissues. Y-axis, log2 O/E ratio, from 100 permutations of breakpoints. Solid and hollow circles, mean log2 O/E ratio of actual and random regions of the corresponding genomic features. Error bars, standard deviation. Shaded area, predefined O/E ratio cutoffs between 1.1 and –1/(1.1). X-axis, Roadmap samples, ranked by O/E ratio. Roadmap samples were grouped and colored as kidney, liver, brain, stem cell and others.

**Figure S9:**
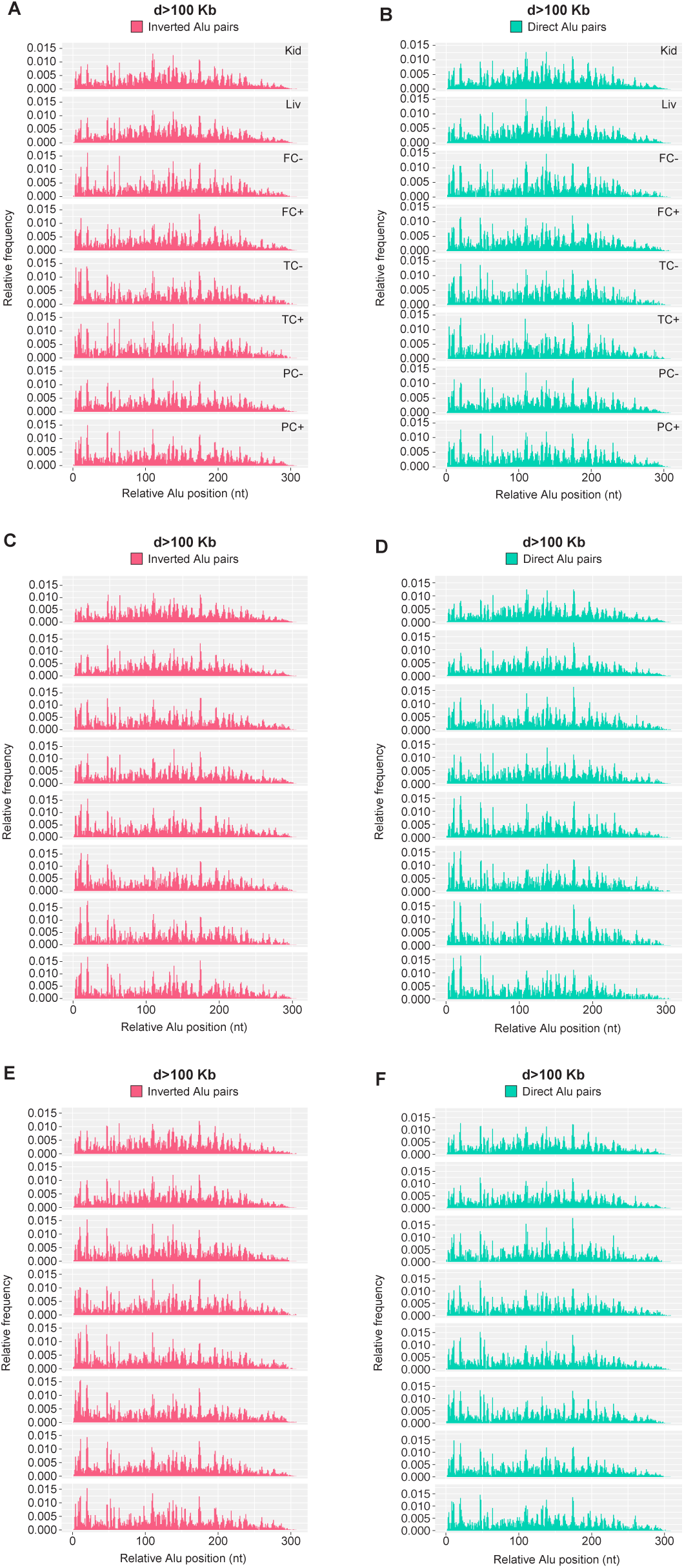
Breakpoint usage frequency profiles along Alu model sequences for NAHR events involving Alu pairs distanced more than 100kb. (A, B) Breakpoint frequency profiles for control dataset NAHR events of Alu-Alu pairs in inverted (A) and direct (B) configuration and distanced more than 100kb. (C, D) Breakpoint frequency profiles for Parkinson’s disease dataset NAHR events of Alu-Alu pairs in inverted (C) and direct (D) configuration and distanced more than 100kb. (E, F) Breakpoint frequency profiles for Alzheimer’s disease dataset NAHR events of Alu-Alu pairs in inverted (E) and direct (F) configuration and distanced more than 100kb. For all panels: Kid, kidney; Liv, liver; FC, frontal cortex; TC, parietal cortex; PC, temporal cortex. + and –, NeuN+ and NeuN- fractions of the brain tissues.

**Figure S10:**
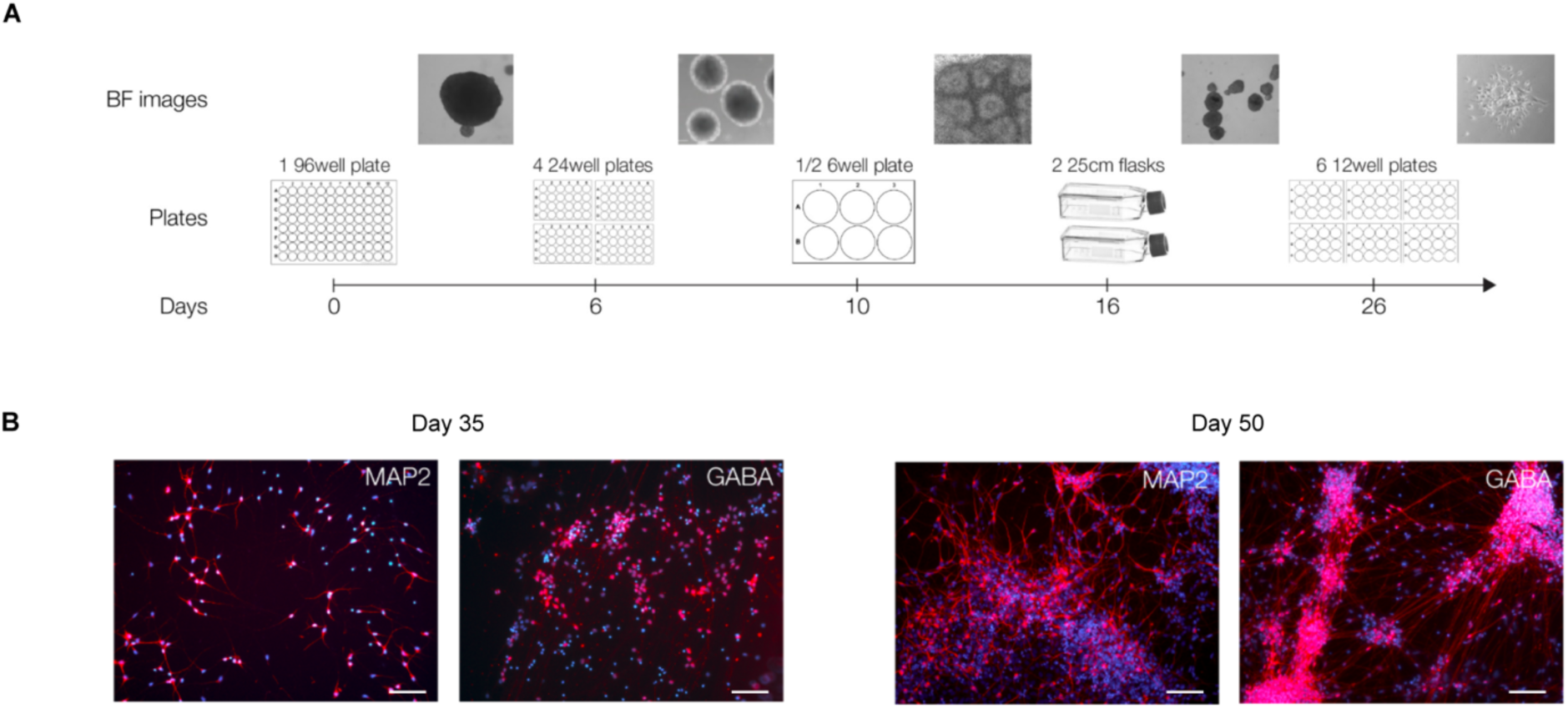
Schematic of the protocol for differentiation of GABAergic cortical interneurons from human IPSCs. (A) Bright field (BF) images, plating strategy, and timescale of the pre-induction iPSC priming phase (days 1-26, see Methods section). (B) Representative immunostaining detection of markers specific for mature neurons, MAP2 and GABA (red), and nuclei (DAPI, 4’,6-diamidino-2-phenylindole, blue) in differentiated neurons at progressing timepoints (day 35 and day 50). Day 50 is the cell collection time point. BF, bright field. Scale bar: 200 μm.

**Figure S11:**
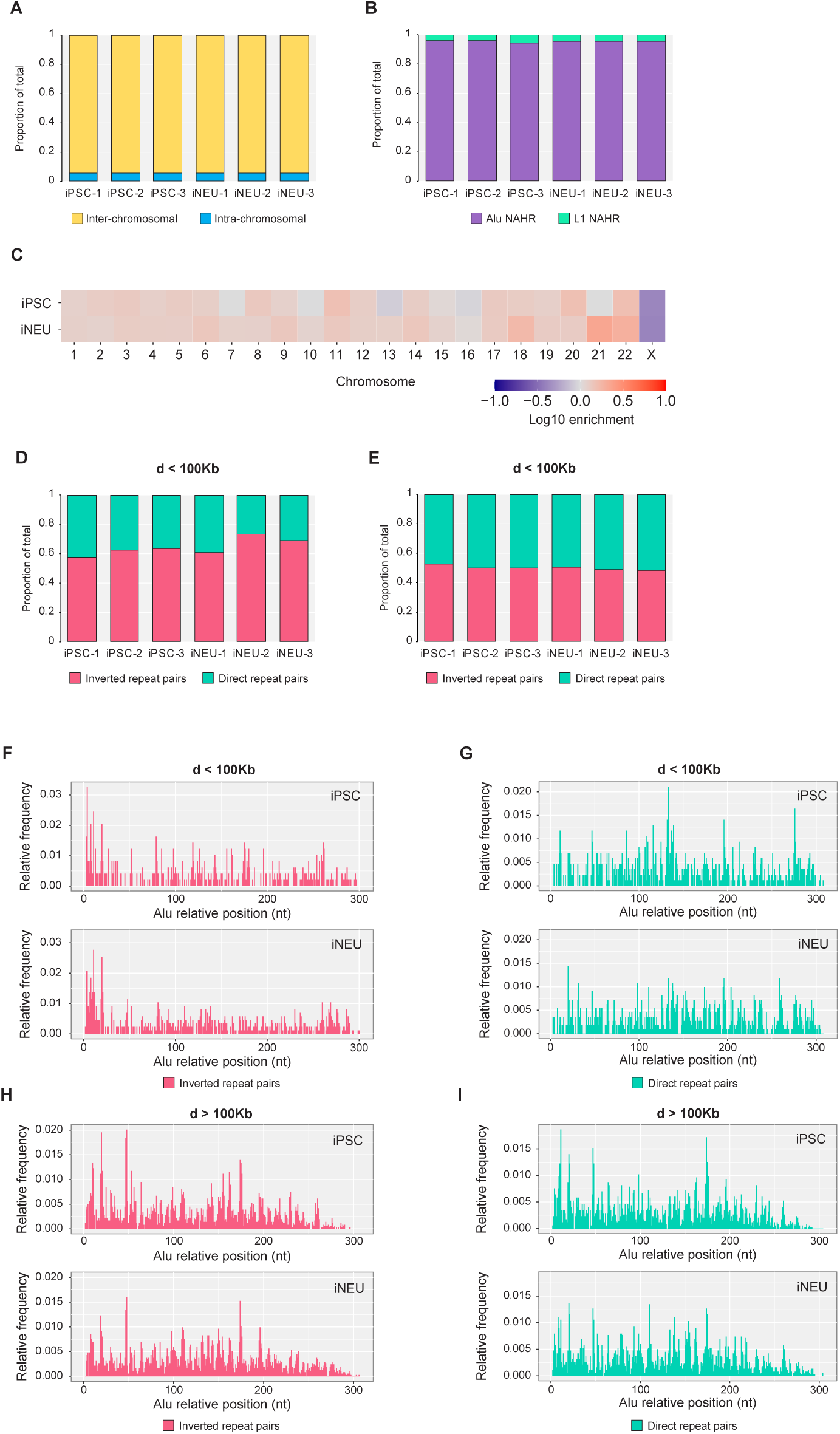
Profiling of NAHR of Alu and L1 elements in human induced pluripotent stem cells (iPSC) and differentiated neurons (iNEU). (A) Relative proportion of intra-chromosomal and inter-chromosomal NAHR events in iPSC and iNEU. Each bar shows the relative proportion of intra-and inter-chromosomal events in one biological replica. (B) Relative proportion of NAHR of Alu and L1 in capture-seq libraries of iPSC and iNEU. Each barplot show the relative proportion of Alu and L1 NAHR events in one biological replica. (C) Intra-chromosomal recombination rate for all chromosomes in iPSC and iNEU capture-seq libraries. Observed values for each chromosome were normalized by the expected values obtained from 100 random recombination dataset of sizes comparable to capture-seq dataset size. Plotted normalized values are in the Log10 scale. Chromosome Y was excluded from this analysis due to its low mappability score. The depletion of recombination in chrX compared to expected values reflects the male derivation of the human iPSCs used in this experiment. (D, E) Analysis of orientation for recombined repeat elements distanced less (D) and more (E) than 100 kilobases in capture-seq libraries of iPSC and iNEU. d, genomic distance. (F, G) Breakpoints frequency displayed along Alu model sequence for NAHR events in iPSC and iNEU capgture-seq libraries, involving Alu elements distanced less than 100 kilobases in inverted (F) and direct (G) configurations. d, genomic distance. (H, I) Breakpoints frequency displayed along Alu model sequence for NAHR events in iPSC and iNEU capgture-seq libraries, involving Alu elements distanced more than 100 kilobases in inverted (H) and direct (I) configurations. d, genomic distance.

**Figure S12:**
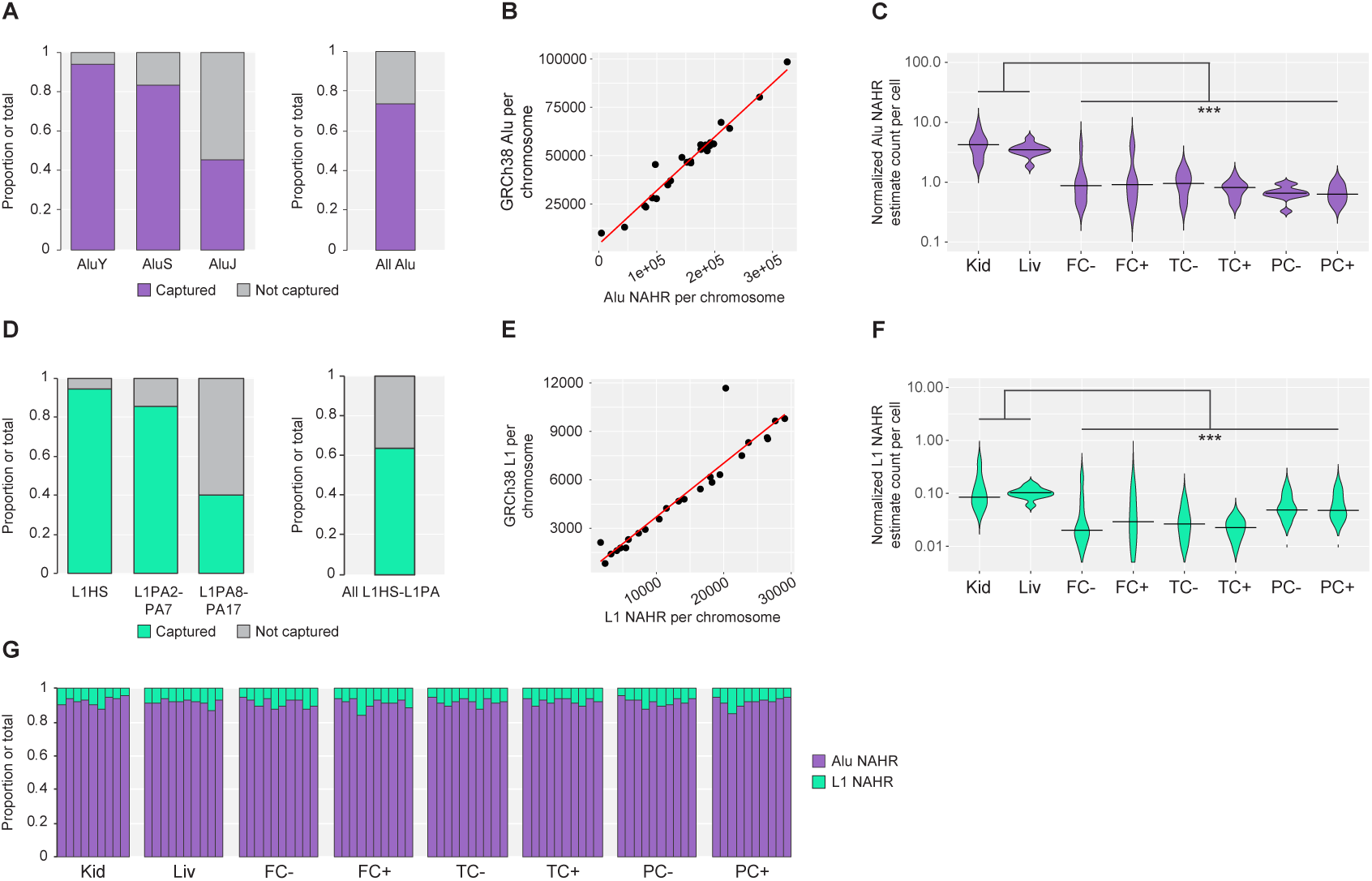
Quality control of capture-seq libraries and estimate normalized NAHR events count for Parkinson’s disease capture-seq libraries. (A, D) Capture efficiency for Alu (A) and L1HS/L1PA2-17 (D) elements annotated in GRCh38 (B, E) Correlation between the number of Alu NAHR events (B) and L1HS/L1PA2-17 (E) per chromosome annotated by TE-reX and the number of genomic Alu and L1 elements per chromosome annotated in the RepeatMasker database. (C, F) Violin plots of the estimated counts of Alu (C) and L1 (F) NAHR events per cell and quartiles annotated in capture-seq libraries of post-mortem samples from Parkinson’s disease donors; counts are normalized by amount of input DNA and sequencing depth. FC, frontal cortex; Kid, kidney; Liv, liver; PC, parietal cortex;TC, temporal cortex. PC, parietal cortex; +, neuron-specific antibody (NeuN) positive; –, NeuN, negative. *P < 0.05; **P < 0.01; ***P < 0.001 (Mann–Whitney U test). (G) Relative proportion of Alu and L1 NAHR events. Each barplot show the relative proportion of Alu and L1 NAHR events in a single capture-seq library.

**Figure S13:**
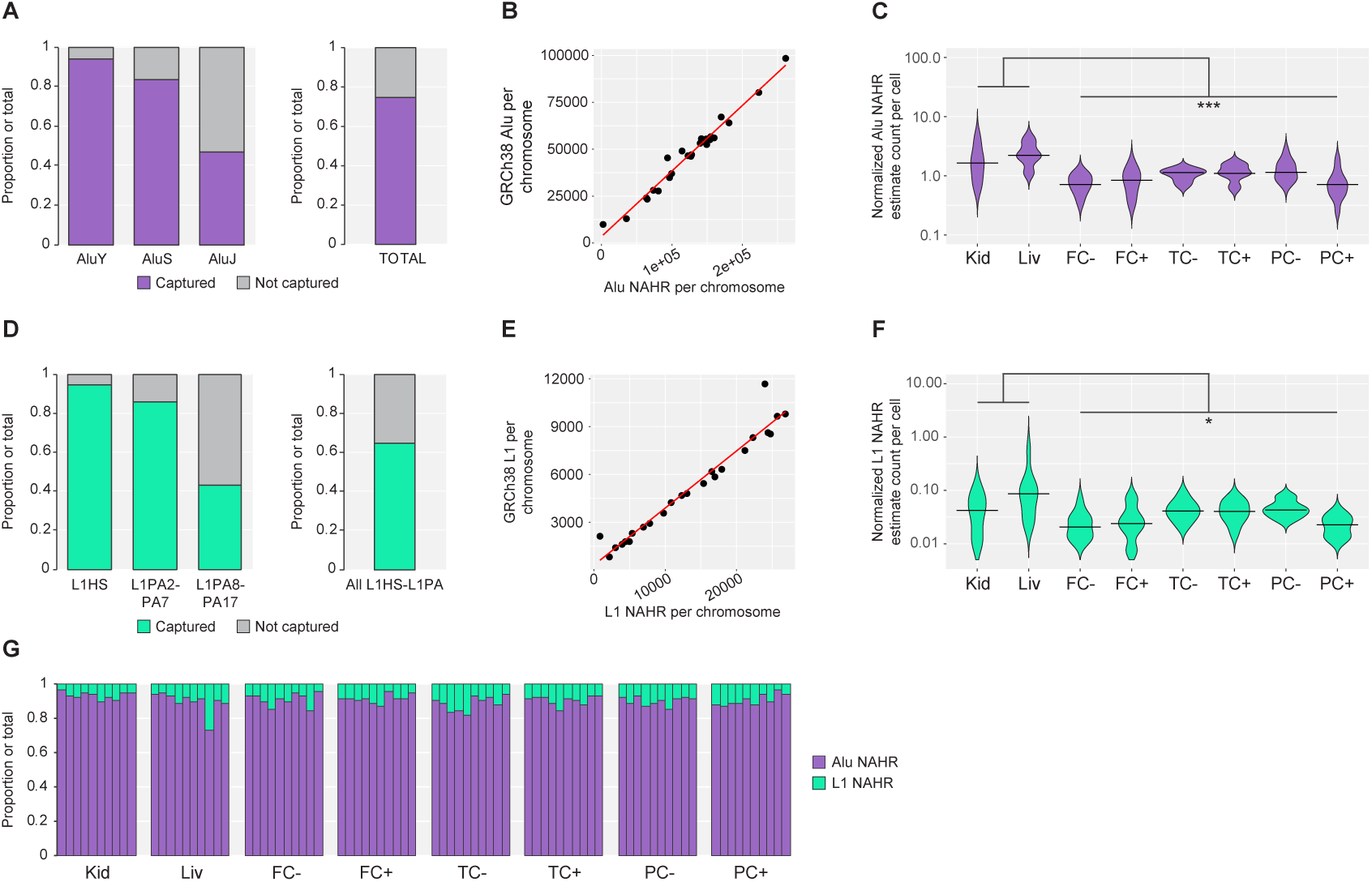
Quality control of capture-seq libraries and estimate normalized NAHR events count for Alzheimer’s disease capture-seq libraries. (A, D) Capture efficiency for Alu (A) and L1HS/L1PA2-17 (D) elements annotated in GRCh38 (B, E) Correlation between the number of Alu NAHR events (B) and L1HS/L1PA2-17 (E) per chromosome annotated by TE-reX and the number of genomic Alu and L1 elements per chromosome annotated in the RepeatMasker database. (C, F) Violin plots of the estimated counts of Alu (C) and L1 (F) NAHR events per cell and quartiles annotated in capture-seq libraries of post-mortem samples from Alzheimer’s disease donors; counts are normalized by amount of input DNA and sequencing depth. FC, frontal cortex; Kid, kidney; Liv, liver; PC, parietal cortex;TC, temporal cortex. PC, parietal cortex; +, neuron-specific antibody (NeuN) positive; –, NeuN, negative. *P < 0.05; **P < 0.01; ***P < 0.001 (Mann–Whitney U test). (G) Relative proportion of Alu and L1 NAHR events. Each barplot show the relative proportion of Alu and L1 NAHR events in a single capture-seq library.

**Figure S14:**
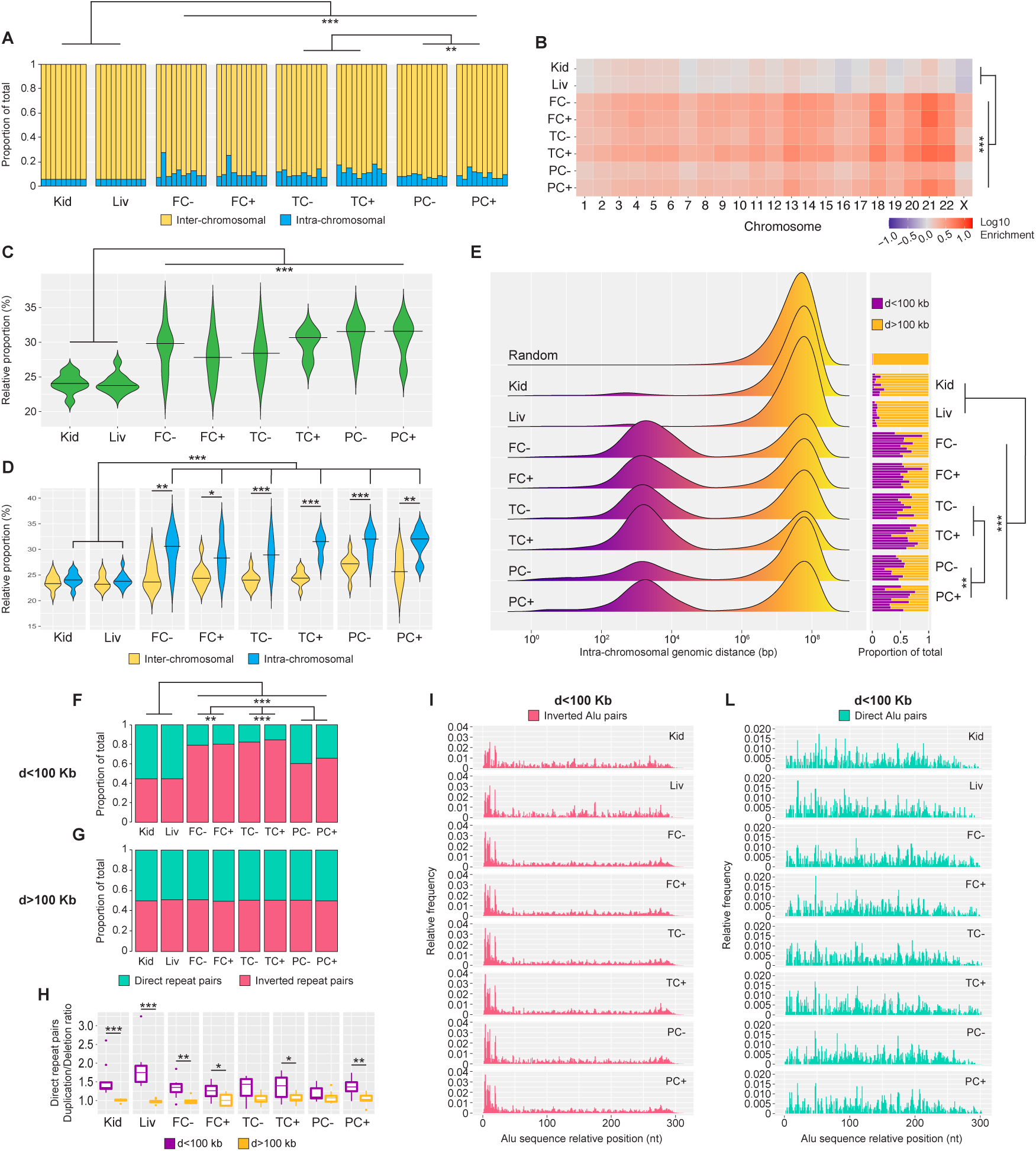
Genome-wide profiling of somatic NAHR events in Parkinson’s disease capture-seq dataset. (A) Relative proportion of intra- and inter-chromosomal for somatic NAHR events of Alu and L1. Each bar shows the relative proportion of intra-and inter-chromosomal events in one donor. (B) Intra-chromosomal recombination rate for all chromosomes and all samples. Observed values for each chromosome were normalized by the expected values obtained from 100 random recombination dataset of sizes comparable to capture-seq dataset size. Plotted normalized values are in the Log10 scale. Chromosome Y was excluded from this analysis due to its low mappability score. (C) Violin plots of the contribution of young repeat elements (AluY and L1HS) and quartiles to NAHR events annotated in capture-seq libraries of Parkinson’s disease donor samples. (D) Violin plots of the contribution of young repeat elements (AluY and L1HS) and quartiles to NAHR events annotated in capture-seq libraries of Parkinson’s disease donor samples, separated in intra- and inter-chromosomal components. (E) Genomic distances of repeats involved in intra-chromosomal NAHR events annotated in capture-seq dataset for all samples. Random is the distance profile for 1 million random intra-chromosomal repeat. Side bar plots show the relative proportions of intra-chromosomal NAHR events involving repeat pairs distanced less or more than 100 kilobases. Each bar shows the relative proportion of intra-and inter-chromosomal events in one library. (F, G) Analysis of orientation for recombined repeat elements distanced less than 100 kilobases showed a strong bias in the brain samples for recombination of elements in inverted configuration (F). No orientation bias was observed for recombined pairs distanced more than 100 kilobases (G). (H) Box-and-whisker plots of the ratio between duplications and deletions for NAHR of repeats in direct configuration and quartiles indicates a bias for duplications for recombined repeats distanced less than 100 kilobases. (I, L) Breakpoints frequency displayed along Alu model sequence show different profiles for somatic NAHR events involving Alu elements distanced less than 100 kilobases in direct (I) and inverted (L) configurations. For tissue and sample abbreviations, see Figure 1. For all panels: d, genomic distance. *P < 0.05; **P < 0.01; ***P < 0.001 (Mann–Whitney U test).

**Figure S15:**
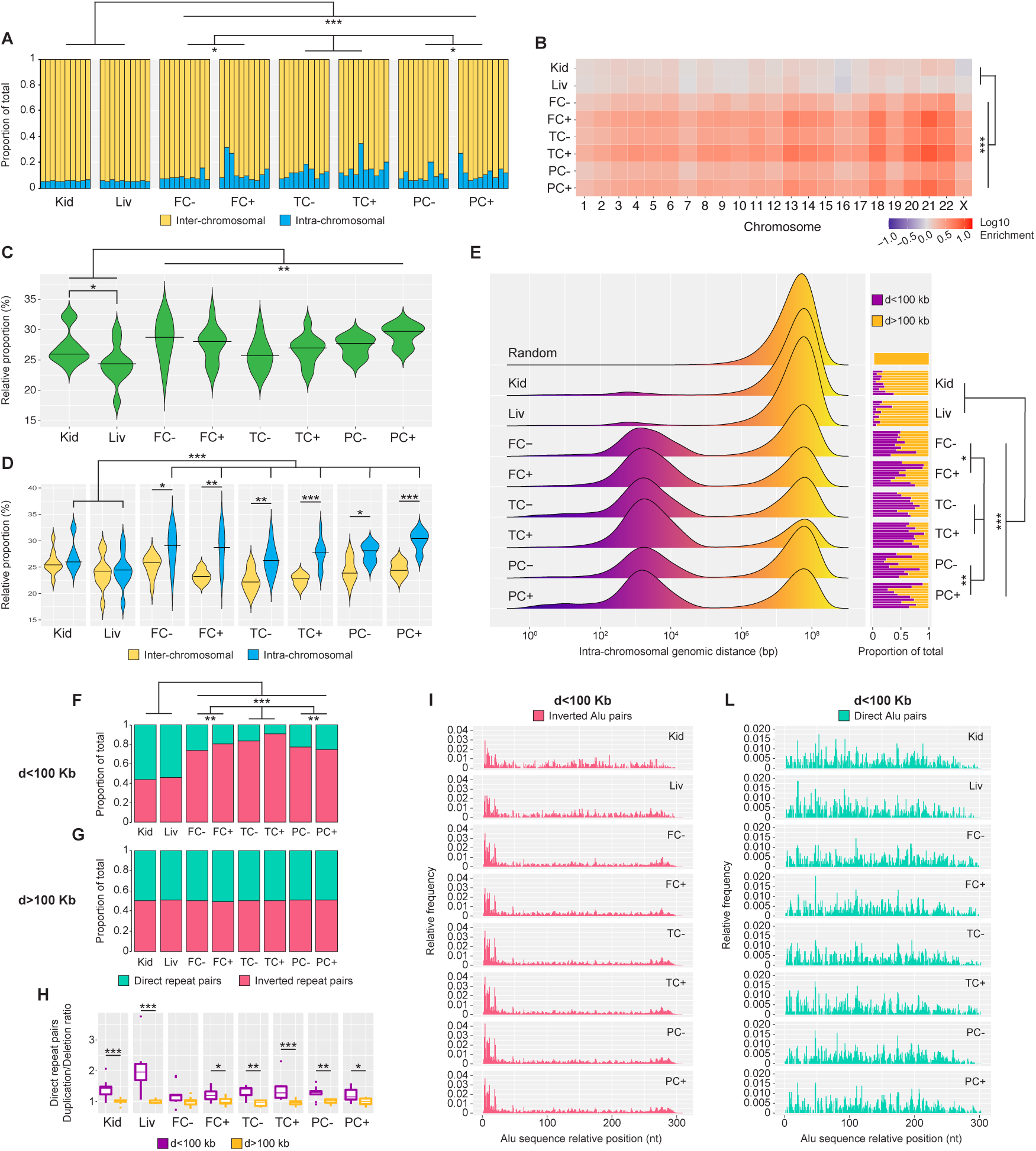
Genome-wide annotation of somatic intra-chromosomal recombination in Alzheimer’s disease dataset. (A) Relative proportion of intra- and inter-chromosomal for somatic NAHR events of Alu and L1. Each bar shows the relative proportion of intra-and inter-chromosomal events in one donor. (B) Intra-chromosomal recombination rate for all chromosomes and all samples. Observed values for each chromosome were normalized by the expected values obtained from 100 random recombination dataset of sizes comparable to capture-seq dataset size. Plotted normalized values are in the Log10 scale. Chromosome Y was excluded from this analysis due to its low mappability score. (C) Violin plots of the contribution of young repeat elements (AluY and L1HS) and quartiles to NAHR events annotated in capture-seq libraries of Alzheimer’s disease donor samples. (D) Violin plots of the contribution of young repeat elements (AluY and L1HS) and quartiles to NAHR events annotated in capture-seq libraries of Alzheimer’s disease donor samples, separated in intra- and inter-chromosomal components. (E) Genomic distances of repeats involved in intra-chromosomal NAHR events annotated in capture-seq dataset for all samples. Random is the distance profile for 1 million random intra-chromosomal repeat pairs. Side bar plots show the relative proportions of intra-chromosomal NAHR events involving repeat pairs distanced less or more than 100 kilobases. Each bar shows the relative proportion of intra-and inter-chromosomal events in one library. (F, G) Analysis of orientation for recombined repeat elements distanced less than 100 kilobases showed a strong bias in the brain samples for recombination of elements in inverted configuration (F). No orientation bias was observed for recombined pairs distanced more than 100 kilobases (G). (H) Box-and-whisker plots of the ratio between duplications and deletions for NAHR of repeats in direct configuration and quartiles indicates a bias for duplications for recombined repeats distanced less than 100 kilobases. (I, L) Breakpoints frequency displayed along Alu model sequence show different profiles for somatic NAHR events involving Alu elements distanced less than 100 kilobases in direct (I) and inverted (L) configurations. For tissue and sample abbreviations, see Figure 1. For all panels: d, genomic distance. *P < 0.05; **P < 0.01; ***P < 0.001 (Mann–Whitney U test).

**Figure S16:**
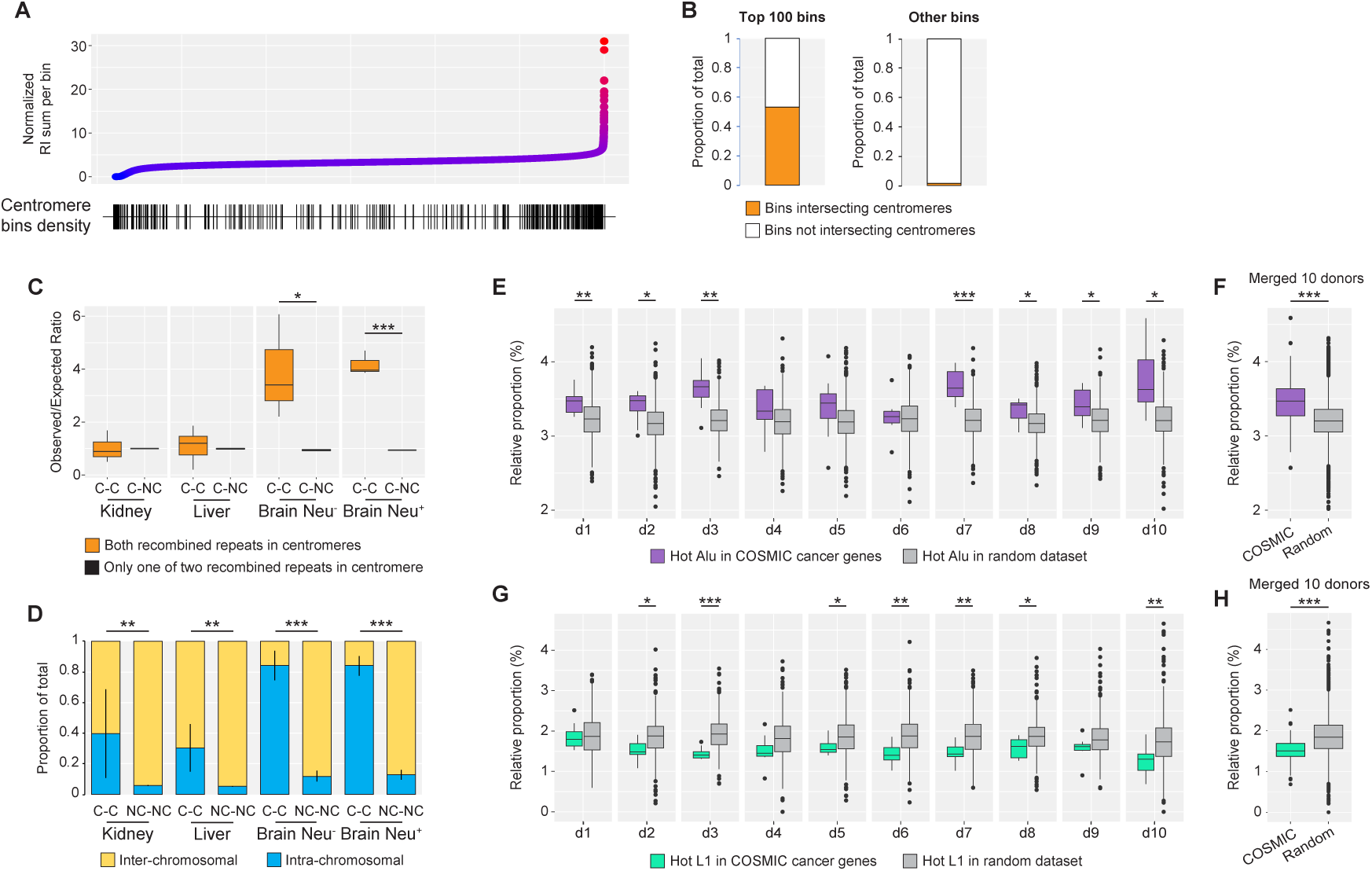
NAHR hotspots analysis in capture-seq libraries of Parkinson’s disease donors. (A) Binning of GRCh38 in 100 kilobases bins (one dot = one bin), ranked by their relative recombination activity, calculated by dividing the sum of the RI of all repeat elements in a given bin by the total bin count of repeat elements annotated in the RepeatMasker. Genomic bins that overlap with centromeres (± 1Mb) are marked with a vertical line in the bottom density plot. (B) Relative proportion of genomic bins intersecting centromeres ± 1Mb in the top 100 bins ranked by relative recombination activity from (A) and in all remaining genomic bins. (C) Box-and-whisker plots of observed/expected ratio and quartiles for recombined repeat pairs with both repeats (“C-C”) or only one out of two repeats (“C-NC”) located within centromeres ± 1Mb. To compensate for overall low counts of NAHR events in centromeric regions, data of all brain samples were merged in NeuN- and NeuN+ fractions. C-C, centromere-centromere; C-NC, centromere-non centromere. **P < 0.01; ***P < 0.001 (Two-way ANOVA). (D) Comparison of intra- and inter-chromosomal recombination rates for recombined repeat pairs with both repeats located within centromeres ± 1Mb (“C-C”) or in non-centromeric regions (“NC-NC”). For tissue and sample abbreviations, see c). **P < 0.01; ***P < 0.001 (Mann–Whitney U test). (E, F) Box-and-whisker plots of relative proportion and quartiles of hot Alu elements with RI ≥10 in cancer genes from the COSMIC database compared to relative proportion in random control datasets generated in silico (100 random datasets per donor. Panel (F): individual donors; panel (G): merged count for all donors. d1-10, donors 1 to 10. *P < 0.05; **P < 0.01; ***P < 0.001 (Mann–Whitney U test). (G, H) Box-and-whisker plots of relative proportion and quartiles of hot L1 elements with RI ≥10 in cancer genes from the COSMIC database compared to relative proportion in random control datasets generated in silico (100 random datasets per donor. Panel (H): individual donors; panel (I): merged count for all donors. d1-10, donors 1 to 10. *P < 0.05; **P < 0.01; ***P < 0.001 (Mann–Whitney U test).

**Figure S17:**
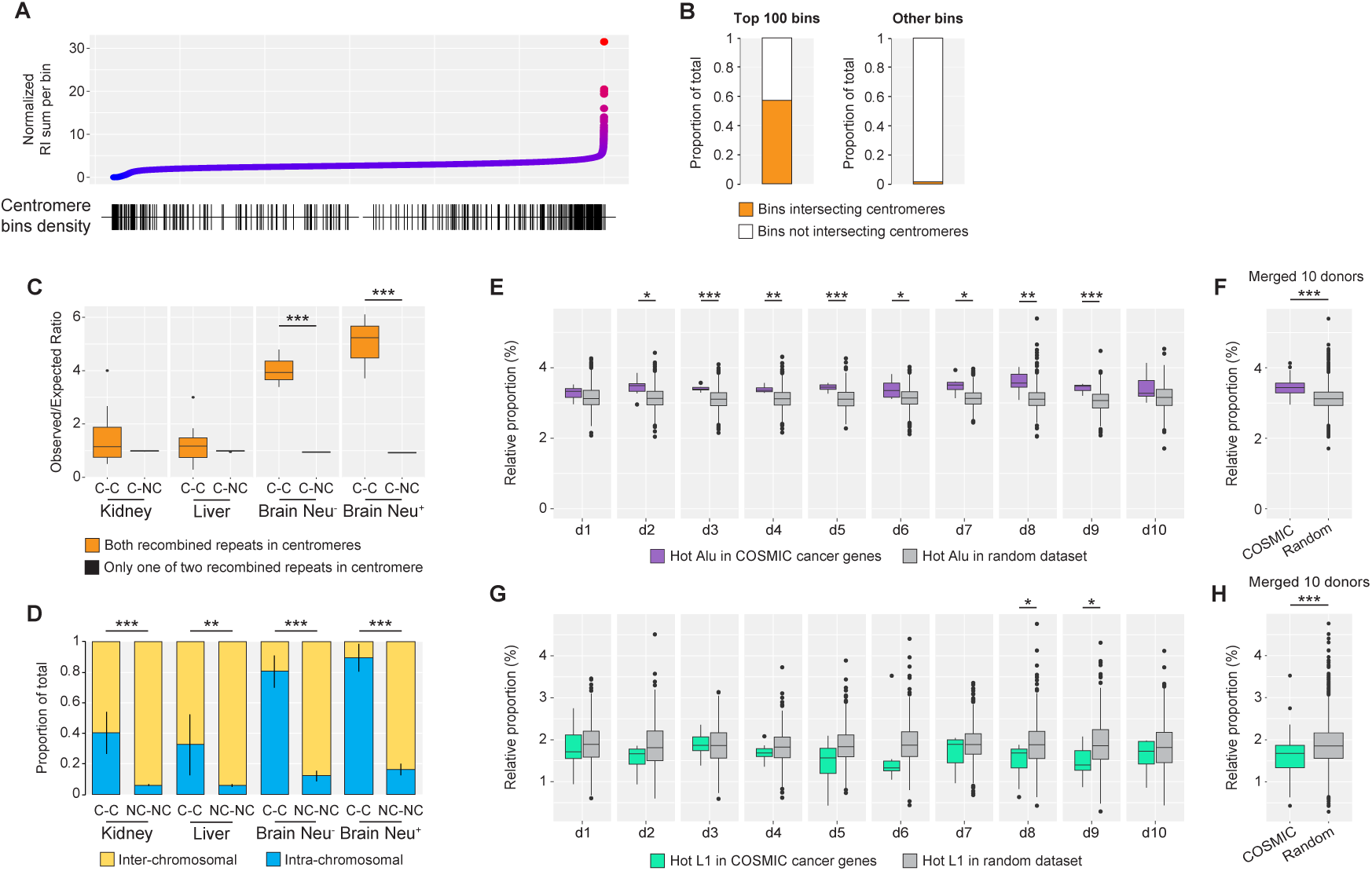
NAHR hotspots analysis in capture-seq libraries of Alzheimer’s disease donors. (A) Binning of GRCh38 in 100 kilobases bins (one dot = one bin), ranked by their relative recombination activity, calculated by dividing the sum of the RI of all repeat elements in a given bin by the total bin count of repeat elements annotated in the RepeatMasker. Genomic bins that overlap with centromeres (± 1Mb) are marked with a vertical line in the bottom density plot. (B) Relative proportion of genomic bins intersecting centromeres ± 1Mb in the top 100 bins ranked by relative recombination activity from (A) and in all remaining genomic bins. (C) Box-and-whisker plots of observed/expected ratio and quartiles for recombined repeat pairs with both repeats (“C-C”) or only one out of two repeats (“C-NC”) located within centromeres ± 1Mb. To compensate for overall low counts of NAHR events in centromeric regions, data of all brain samples were merged in NeuN- and NeuN+ fractions. C-C, centromere-centromere; C-NC, centromere-non centromere. **P < 0.01; ***P < 0.001 (Two-way ANOVA). (D) Comparison of intra- and inter-chromosomal recombination rates for recombined repeat pairs with both repeats located within centromeres ± 1Mb (“C-C”) or in non-centromeric regions (“NC-NC”). For tissue and sample abbreviations, see c). **P < 0.01; ***P < 0.001 (Mann–Whitney U test). (E, F) Box-and-whisker plots of relative proportion and quartiles of hot Alu elements with RI ≥10 in cancer genes from the COSMIC database compared to relative proportion in random control datasets generated in silico (100 random datasets per donor. Panel (F): individual donors; panel (G): merged count for all donors. d1-10, donors 1 to 10. *P < 0.05; **P < 0.01; ***P < 0.001 (Mann–Whitney U test). (G, H) Box-and-whisker plots of relative proportion and quartiles of hot L1 elements with RI ≥10 in cancer genes from the COSMIC database compared to relative proportion in random control datasets generated in silico (100 random datasets per donor. Panel (H): individual donors; panel (I): merged count for all donors. d1-10, donors 1 to 10. *P < 0.05; **P < 0.01; ***P < 0.001 (Mann–Whitney U test).

**Figure S18:**
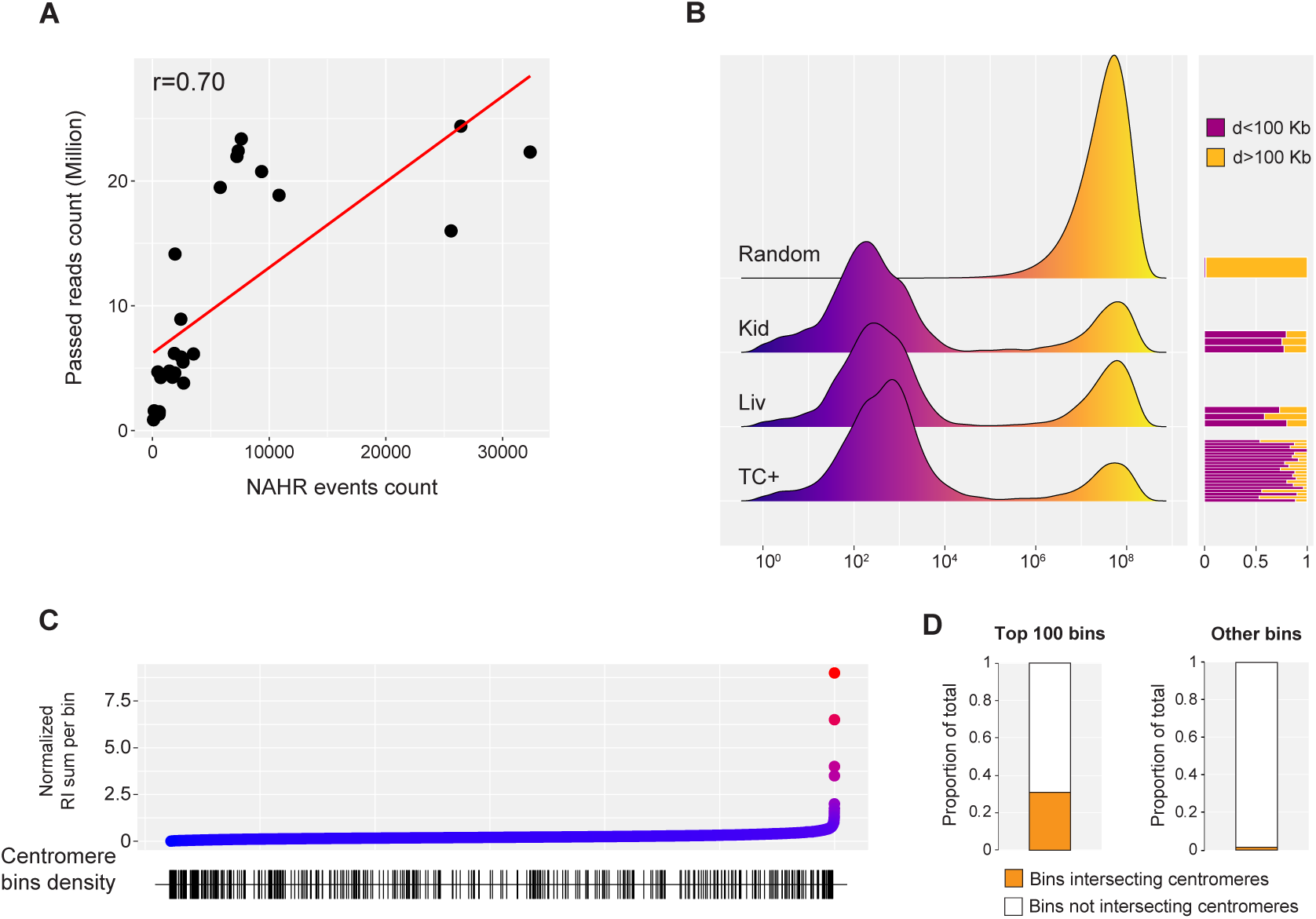
Additional analyses of NAHR in PromethION WGS libraries. (A) Correlation between number of NAHR events annotated by TE-reX in PromehtION WGS library and the number of reads per library that passed QC. (B) Genomic distances of repeats involved in intra-chromosomal NAHR events annotated in PromethION WGS for all samples. Random is the distance profile for 1000 datasets of random intra-chromosomal repeat pairs of size comparable to real datasets. Side bar plots show the relative proportions of intra-chromosomal NAHR events involving repeat pairs distanced less or more than 100 kilobases. Each bar shows the relative proportion of intra-and inter-chromosomal events in one library. d, genomic distance. (C) Binning of GRCh38 in 100 kilobases bins (one dot = one bin), ranked by their relative recombination activity, calculated by dividing the sum of the RI of all repeat elements in a given bin by the total bin count of repeat elements annotated in the RepeatMasker. Genomic bins that overlap with centromeres (± 1Mb) are marked with a vertical line in the bottom density plot. (D) Relative proportion of genomic bins intersecting centromeres ± 1Mb in the top 100 bins ranked by relative recombination activity from (C) and in all remaining genomic bins.

**Figure S19:**
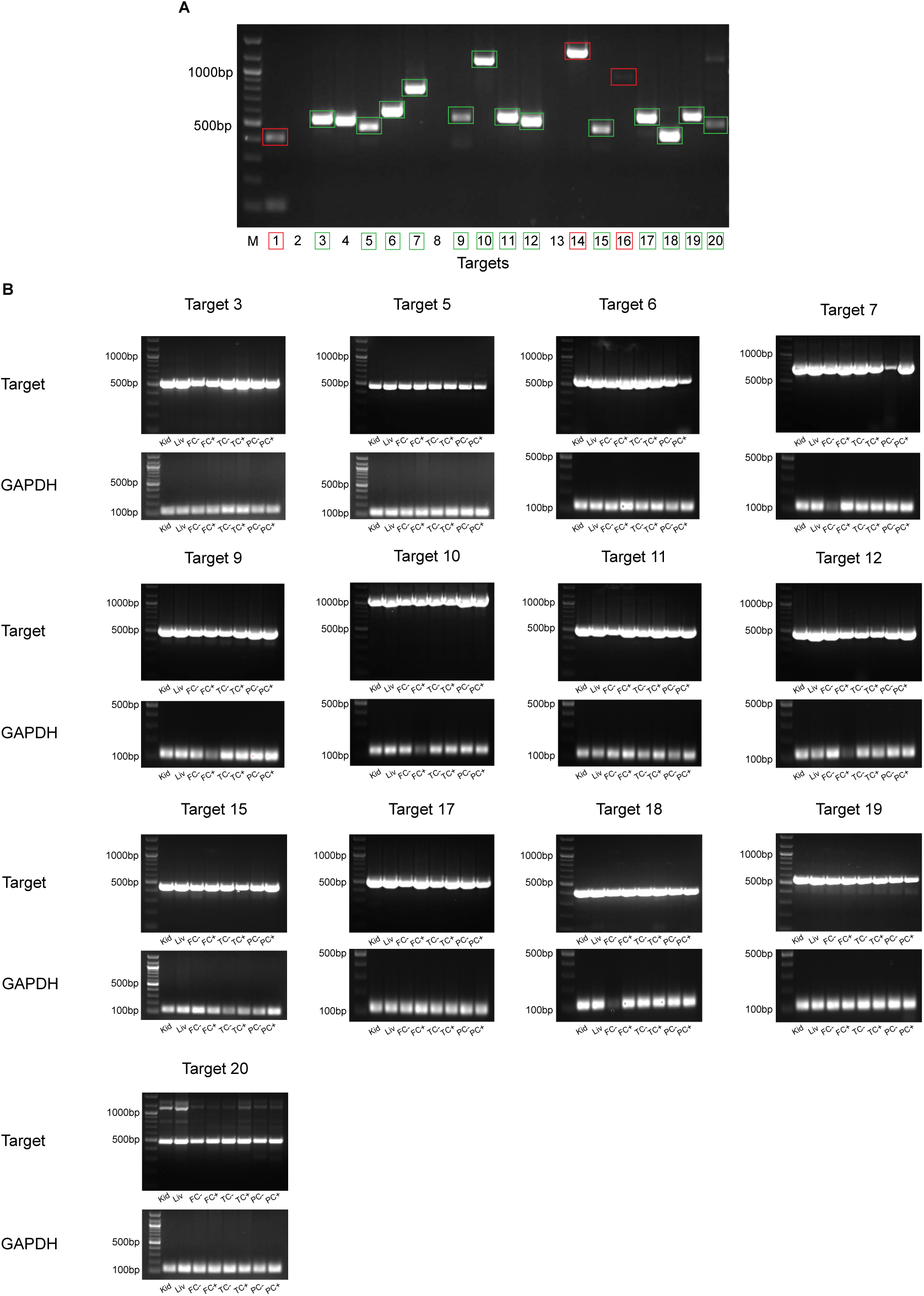
PCR validation of putative polymorphic NAHR events. (A) PCR validation for 20 putative polymorphic NAHR events found in capture-seq and PromethION WGS datasets. The input DNA was one genomic DNA sample from one representative donor. Details about all targets are in Table S5. Amplicons in colored boxes were confirmed by Sanger sequencing; amplicons in green boxes were selected for subsequent PCR amplification from the full panel of 8 samples available for one representative donor. (B) PCR for the full panel of 8 samples available for one representative donor, for targets in green boxes in a). All panels are photos of PCR products run on 1.5% agarose gel stained with ethidium bromide.

**Figure S20:**
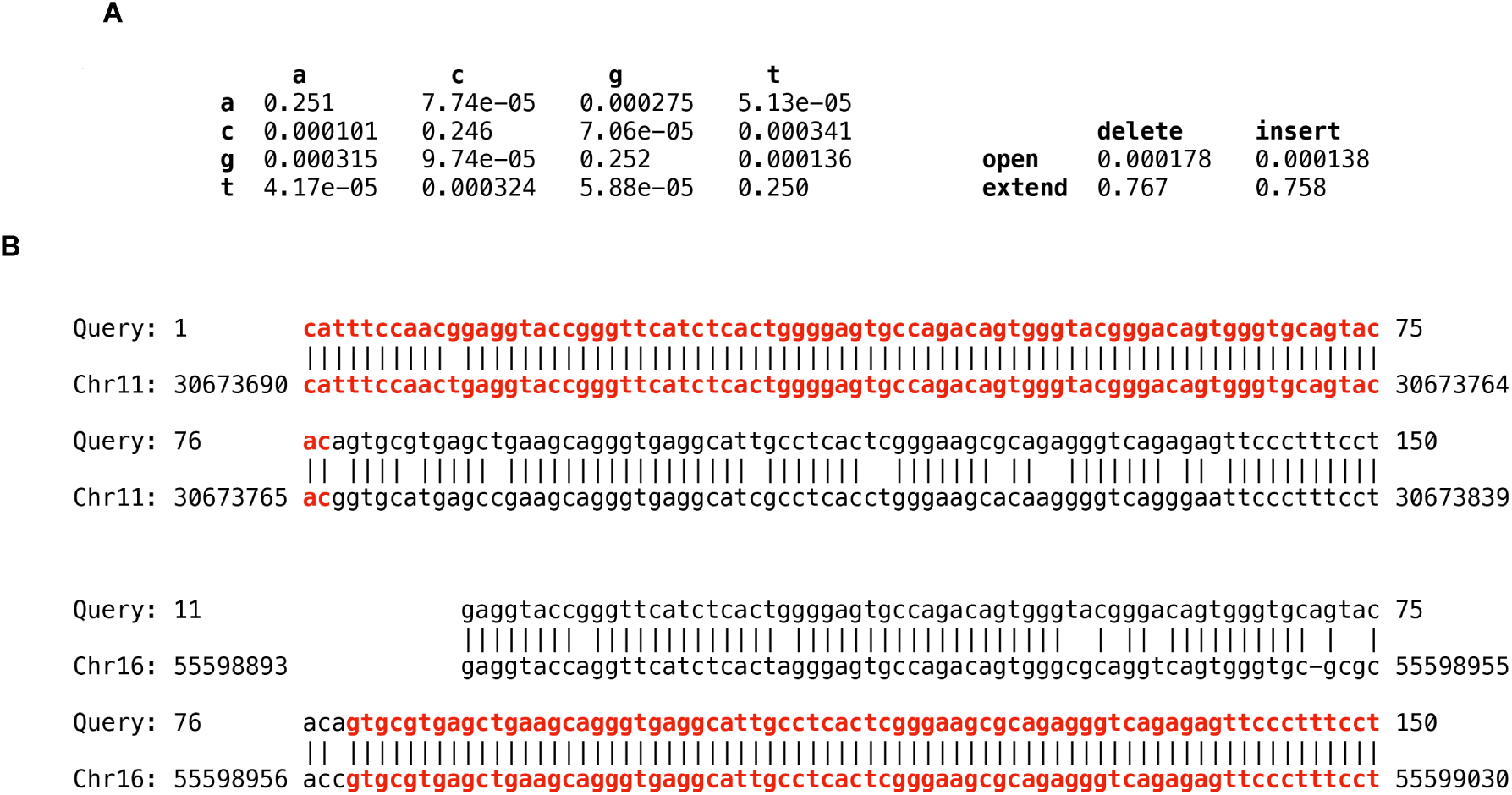
Alignment of DNA sequences to a genome, allowing for homologous recombination. (A) Rates (probabilities) of substitutions, deletions and insertions between one set of DNA sequences and the genome. The 4×4 matrix shows substitution probabilities: rows correspond to genome bases, columns correspond to query-sequence bases. (B) Example of a query-to-genome alignment that indicates homologous recombination. The whole query sequence (150 bp) can be aligned to a LINE-1 element in chromosome 11, and most of it can be aligned to an L1 element in chromosome 16. The query can also be aligned to many other L1 elements (not shown). The alignment produced by last-split is shown in red: the left half of the query comes from this L1 in chromosome 11, rather than anywhere else in the genome, with error probability 10^-6, and the right half comes from this chromosome 16 L1 with error probability p=10e-8.

## Data and code availability

**Data:** all sequencing data for this project have been deposited in the NCBI Sequence Read Archive (SRA) database under accession number PRJNA636606, and are accessible at the following link: https://www.ncbi.nlm.nih.gov/sra/PRJNA636606.

**Code:** TE-reX and related documentation can be accessed at: https://gitlab.com/mcfrith/te-rex.

## Ethics statement and post-mortem human samples

The use of human samples in this study was approved by Riken Research Ethics Committee with permission number H23-16. All information related to the donors are listed in Table S1

## Methods details

### Purification of genomic DNA from post-mortem frozen tissues

All post-mortem human samples were obtained from the Brain Bank for Ageing Research, part of Tokyo Metropolitan Hospital. Upon reception samples were stored at -80C until usage. For genomic DNA extraction from liver and kidney samples ∼200mg of frozen tissue were reduced to fine powder in a liquid nitrogen-cooled mortar and transferred to 10ml lysis buffer(Laird et al., 1991). Genomic DNA extraction was performed as described previously with minor modifications(Wood, 1983). DNA pellets were resuspended in DNA rehydration solution (Promega) and after quantitation with Nanodrop (Thermo Scientific) quality control of each sample was performed by loading 300ng of resuspended DNA on 1% agarose gel. Due to scarcity of available starting material 3 DNA samples were exhausted during processing and could not be replaced (kidney sample for control donor 4, kidney sample for Parkinson’s disease donor 6).

### Purification of genomic DNA from sorted nuclei

After sorting each nuclei pellet was added with 900ul of lysis buffer (1% SDS, 10mM EDTA 0.5M pH 8, 50mM Tris HCl pH 8, 150mM NaCl) and thoroughly resuspended by pipetting until no clumps were visible. Samples were incubated at 65C for 20 minutes, after which RNAse A (Sigma, 10ug/ml) was added and the digestion was carried on for 1 hour. After adjusting the temperature to 55C Proteinase K (NEB, 100ug/ml) was added to each sample and digestion was continued for 3 hours. Samples were then let cool down at RT. An equal volume of Phenol:Chloroform:Isoamyl alcohol (25:24:1 v/v, Sigma) was added to the tubes and after 20 seconds vortexing the samples were centrifuged at 13000g for 7 minutes. The supernatant was carefully transferred to a fresh tube and the DNA was precipitated by adding NaAc 0.3M (Sigma) and 2.5 volumes of EtOH (Wako). After overnight incubation the samples were centrifuged at 13000g, 4C for 30 minutes. The supernatant was discarded and the pellets were washed 2x with EtOH 75%. After removal of residual EtOH droplets the pellets were air dried for 10 minutes and resuspended in DNA rehydration solution (Promega). After quantitation with Nanodrop (Thermo Scientific) 300ng of DNA from each sample were loaded on 1% agarose gel for quality control. Due to scarcity of starting material one sample was exhausted during testing and processing and could not be replaced (Control donor 2, Parietal cortex NeuN+).

### Differentiation of human cortical interneurons from human iPSCs

Differentiation of iPSC into human cortical interneurons was adapted from Liu et al (Liu et al., 2013). On day 0, iPSCs (HUBi001-A, Sigma-Aldrich) were dissociated into a single-cell suspension using Accutase (Life Technologies) and seeded into ultra-low attachment 96-well plates at 9 x 103 cells per well in StemFit containing 10 µM Y-27632 (STEMCELL Technologies). Neuroepithelial induction of self-assembled embryoid bodies (EB) was initiated on day 7 by transferring single EBs into low-attachment 24-well plates containing 500 μl of neuronal induction medium (NIM) consisting of DMEM/F12 (Wako), 1X N2-supplement (GIBCO), 1X non-essential amino acids (NEAA, Thermo Fisher Scientific) and 20 μg/mL heparin (Sigma). On day 11, EBs were attached to mouse laminin (15 μg/mL, Sigma)-coated 6-well plates at approximately 30 EBs per well and maintained in NIM until day 14 when neuronal rosettes appeared. Patterning of neuroepithelial cells into medial ganglionic eminence (MGE) progenitors was initiated on day 14 by replacing old medium with NIM containing 1.5 μM Purmorphamine (Stemgent). On day 16, neuronal rosettes, formed by multiple layers of columnar epithelia, were dissociated by gently blowing off the colonies using a 1 ml pipette tip, transferred into T25 flasks and maintained in NIM containing 1X B-27 supplement w/o vitamin A (Thermo Fisher Scientific) until day 26. Differentiation into GABAergic cortical interneurons was initiated on day 26 by dissociation of MGE progenitors using 1X TrypLE (Thermo Fisher Scientific) and plating MGE progenitors into Poly-L-ornithine (PLO, Sigma) and laminin (Sigma)-coated 12-well plates at 4.5 x 10^4^ cells per well containing neuronal differentiation medium consisting of DMEM/F12 (Wako), 1X N2-supplement (GIBCO), 1X non-essential amino acids (NEAA, Thermo Fisher Scientific), 1X B-27 supplement with vitamin A (Thermo Fisher Scientific), 10 ng/ml BDNF (GIBCO), 10 ng/ml GDNF (Thermo Fisher Scientific), 10 ng/ml IGF (PeproTech) and 1 µM cAMP (Sigma). Proliferation of cells was inhibited by supplementing the medium for the first 3 days with 0.2 μM Compound E (EMD Biosciences). Half of the medium was replaced with fresh medium every third day until day 50. Undifferentiated human iPSC used for comparison in capture-seq were maintained in StemFit (Ajinomoto Co., Inc) on iMatrix-511 (Funakoshi)-coated 6-well plates. At the time of collection iPSC and differentiated neurons pellets were washed with PBS and stored at -80C.

### Construction and mapping of capture-seq and long-read WGS libraries

Construction and quality control of capture-seq libraries are described in Supplementary Document 1. All capture-seq libraries from post-mortem tissues were sequenced on Illumina Hiseq4000 platform by BGI Genomics and demultiplexed according to respective indexes.

For capture-seq libraries from iPSC and differentiated neurons, cell pellets were dissolved in lysis buffer (100mM Tris Hcl pH 8.5, 50mM EDTA, 0.1% SDS, 100mM NaCl) and genomic DNA was purified with Wizard Genomic DNA Purification Kit (Promega). Samples were quantified with Qubit dsDNA BR Assay Kit (Thermo Fisher) and capture-seq libraries were constructed as described in Supplementary Document 1 with no modifications. Libraries were sequenced at 600 cycles on Illumina Miseq using Reagent Kit v3 (Illumina).

PromethION libraries were prepared from a total of 1 microgram of intact genomic using the Ligation Sequencing Kit (SQK-LSK109, Oxford Nanopore Technologies) following the manufacturer’s protocol. Sequencing was done on the PromethION sequencer for 72 hours with the MinKNOW software, and the base calling of the FAST5 data from PromethION was performed with Guppy v4.0.11 and subsequently converted into FASTQ files.

For capture-seq libraries, overlapping paired reads (median across libraries ∼98%) were merged in longer contigs using FLASH (Magoč and Salzberg, 2011); non-overlapping reads were merged with respective FLASH-extended contigs to generate single FASTQ files. FASTQ files from PromethION were aligned directly. Alignment on GRCh38 was performed with LAST(Kiełbasa et al., 2011) using standard parameters.

### Capture efficiency for capture-seq libraries

To calculate capture efficiency across the capture-seq dataset we extracted from all mapped capture-seq libraries reads with a low mismap score (P < 10e-4), converted the reads to BED format, intersected the genomic coordinates of the reads with the genomic coordinates of subfamilies Alu and L1 elements annotated in RepeatMasker (Table S3) and finally calculated the relative proportion of captured elements for each capture-seq library.

### Identification of non-allelic homologous recombination with TE-reX

Finding DNA sequence rearrangements involving repeat elements, especially homologous recombination, by comparing DNA reads to a reference genome requires accurate probability-based reads-to-genome alignment. We first determined the rates (probabilities) of substitutions, deletions, and insertions between reads and genome using last-train (Hamada et al., 2017) (Figure S20A). In order to find genome sequence rearrangements involving repeats, TE-reX needs three input files: 1) Alignments of DNA reads to the genome; 2) Alignments of the genome to repeat consensus sequences; 3) Alignments between pairs of repeat consensus sequences. The read-to-genome alignments are made with last-split (Frith and Kawaguchi, 2015), so each read base is aligned to at most one genome base. Each read has one or more alignments, e.g. the read in Figure S20B has two alignments (shown in red). The genome-to-repeat alignments are made with last-split as well, so each genome base-pair is aligned to at most one base-pair in the repeat database. The repeat-to-repeat alignments retain at most one highest-scoring alignment between any pair of repeats. To identify the rearrangements, for each read we checked whether any pair of alignments that occur consecutively in the read indicate homologous recombination following these steps:

- Require at least one alignment of the pair to have error probability (mismap score) P < 10e-4.
- Require <= *slop* unaligned read bases between the two alignments. *slop* is a parameter with default value 4, however for maximum stringency all TE-reX datasets in this work have been produced with slop set to 0.
- Call the end of the 1st alignment endX, and the beginning of the 2nd alignment begY.
- Require endX to lie within a genomic segment that is aligned to a repeat (call it repeatX); likewise require begY to lie within some repeatY.
- Require that repeatX and repeatY are of types that have an alignment in the repeat-to-repeat alignments.
- Based on the repeat-to-repeat alignments, find endXh = the homologous coordinate in repeatY of endX, and begYh = the homologous coordinate in repeatX of begY. Require that at least one of (endXh, begY) and (endX, begYh) differ by <= *slop*.
- If repeatX and repeatY are of the same type (same repeat consensus sequence), check whether the alignments could be explained by reference-specific deletion. For example: if the read comes from repeatX only, and part of repeatX is deleted in the reference genome (but not in the read), that part of the read may get wrongly aligned to a paralog, in a way that looks just like homologous recombination. Reference-specific deletion is considered possible if the remaining genomic length of repeatX after endX is less than the genomic length of repeatY that is aligned to the read (or likewise with X and Y swapped).

### Post-processing of TE-reX output files

TE-reX output files are in .maf format and in order to annotate and analyze recombination events genome-wide we first processed the .maf files using custom scripts to perform the following operations: a) convert 4-lined .maf files in 1-lined BED-like format; b) collapse PCR duplicates into single entries; c) generate a unique identifier for each recombination event, by joining the modified RepeatMasker annotations of each pair of recombined repeats and the breakpoint junction information from TE-reX; d) select for recombination events having both split reads with mismap score = P < 10e-4; e) calculate the genomic distance of the recombined repeat pairs (<100 kb, >100kb, inter-chromosomal).

### Subsampling of capture-seq libraries and estimate of NAHR events per cell

In order to perform inter-library and inter-dataset comparisons of NAHR events counts we subsampled the FASTQ files of all capture-seq libraries according to the library with the lowest FASTQ reads count (∼8.3 million FASTQ reads, Table S2), re-aligned the subsampled files and run again TE-reX on the subsampled mapped libraries.

To estimate the number of NAHR events per cell in each library, we first calculated the theoretical number of cells used to construct each library based on the amount of input processed DNA. The pre-capture input DNA for each library consisted of fragmented genomic DNA (∼220bp), adapters and sequencing linkers (119bp). Hence, each pre-capture library construct contained ∼65% of genomic DNA sequence and ∼35% of accessory DNA sequences. For Alu and L1 capture libraries we used respectively 100ng and 300ng of input processed DNA, we calculated a total amount of genomic DNA used as a pool for capture of 65ng and 195ng for Alu and L1 libraries, respectively. Considering that a single diploid nucleus contains ∼6pg of genomic DNA, the amount of genomic DNA used as input for Alu and L1 capture in each library corresponded to a theoretical individual cells number of respectively ∼10833 and ∼32500. The values plotted in Figure 1H, 1I, 4f, S12C, S12F, S13C and S13F were obtained by normalizing the count of putative somatic NAHR in the subsampled capture-seq libraries by the theoretical estimate of individual cells used per library.

### Isolation of putative somatic and polymorphic/germline NAHR events

Putative somatic and putative polymorphic/germline NAHR datasets were isolated by merging NAHR events annotated in all capture-seq libraries, and by collapsing the unique identifiers. Recombination events with unique identifiers found only in a single library were flagged as putative somatic. For the definition of putative polymorphic/germline NAHR events, we considered unlikely for a somatic recombination event to involve the same 2 repeat elements using the same breakpoint junction. However, we cannot completely exclude that this may happen in some instances, given that the junctions of putative somatic NAHR events present some positional bias (Figure 2I). By definition, polymorphic/germline NAHR events should be detected in all 8 libraries available for each donor (kidney, liver, frontal cortex NeuN-, frontal cortex NeuN+, temporal cortex NeuN-, temporal cortex NeuN+, parietal cortex NeuN-, parietal cortex NeuN+). However, accounting for potential detection/sensitivity bias due to the short reads we decided to relax this criterion and we preventively flagged as putative polymorphic all the NAHR events with the same unique identifier found in 2+ capture-seq libraries.

### PCR validation

For the PCR validation we selected recombination events for which we could identify non-repeat flanking genomic regions at both 5’- and 3-ends of each contig encompassing one single recombination event. The primers were manually designed using Primer3 (Untergasser et al., 2012); all sequences annealed on unique genomic regions, thus suppressing cross-amplification of non-specific sequences. PCR reactions were performed with Q5 High-Fidelity Master Mix (NEB) in 25ul using 25ng of input (capture-seq library or genomic DNA) and 10uM of forward/reverse primers. Amplification was carried out with initial 10 cycles of touchdown PCR (1x initial denaturation 98C for 30s, 10x 98C for 10s, primers-specific Tm +10C for 30s decreasing by 1C each cycle, 72C for 7s) followed by 30 cycles of standard PCR (98C for 10s, primers-specific Tm for 30s, 72C for 7s, final elongation 72C for 120s). PCR products were cleaned with AMPure XP beads (Beckman Coulter) and quantified with Qubit High Sensitivity dsDNA assay (Thermo Fisher). 20ng of each PCR products were transferred in 96-wells plates, mixed with 5uM forward primers and submitted for Sanger sequencing (Genewiz). Where needed, nested PCR was performed with LongAmp Hot Start Taq 2X Master Mix (New England Biolabs) following manufacturer’s protocol. The first PCR was carried out in 25ul volume using 20ng input DNA; amplification included initial 10 cycles of touchdown PCR (1x initial denaturation 94C for 30s, 10x 94C for 30s, primers-specific Tm +10C for 30s decreasing by 1C each cycle, 65C for 20s) followed by 10 cycles of standard PCR (94C for 10s, primers-specific Tm for 30s, 65C for 20s, final extension 65C for 10 minutes). The nested PCR used 2ul of input from the first PCR and was performed for additional 35 cycles (94C for 10s, primers-specific Tm for 30s, 65C for 20s, final extension 65C for 10 minutes).

### Enrichment of NAHR events in gene and regulatory regions

Enrichment, or depletion, of Alu and L1 NAHR events in gene and regulatory regions (i.e. genomic features) were assessed by random permutation of their breakpoints. These genomic features include 1) the exonic coding sequence (CDS), 3’ untranslated region (UTR), 5’UTR, intron and transcription start sites of coding transcripts from GENCODE v34, and 2) regulatory regions (i.e. enhancer and promoter) from Roadmap Epigenomics consortium (38), either from individual samples (n=111, in FigureS13-14) or all samples pooled (in FigureS11-12). A set of control regions, generated by randomly sampling of 1 kb windows (n=1,000,000) from the masked genome, was used to normalize potential bias in the permutation procedure (described below). The masked genome is referred to as chromosome 1 to 22 and X in GRCh38, subtracted with gap regions defined in UCSC genome browser. Breakpoints (n=2 per NAHR event) of the same tissue fraction from multiple donors of the same disease/control group were pooled and permuted (i.e. shuffled, using BEDTools (Quinlan and Hall, 2010) *shuffle*) into their corresponding TE elements (i.e. AluY/S/J and L1HS/PAx) in the masked genome for 100 times. Only the TE elements with at least 1 observed breakpoint in all datasets were included in the shuffling scope. The number of observed and expected (i.e. permuted) breakpoints within each set of genomic features, and the control regions, was counted. The observed-to-expected (O/E) ratio of each set of genomic features, is defined as the ratio of the observed to expected breakpoints within the genomic feature, normalized by that of the control region. The permutation procedure was repeated with each set of the genomic feature regions (i.e. actual regions, solid dots in FigureS11-14) randomly shuffled within the whole genome (i.e. random regions, hollow circles in FigureS11-14). A Student’s t-test was used to compare the O/E ratio of the actual region to the random region. We consider the NAHR events are significantly enriched, or depleted, in a set of genomic features when P < 0.05 and O/E ratio is >1.1 or <1/(1.1).

### Genome-wide profiling of NAHR events

All analyses were performed using a combination of BEDtools and custom bash scripts. For the analysis of intra-chromosomal recombination rates the real data was compared against the genomic distribution of 100 random dataset generated *in silico* by shuffling RepeatMasker annotations of Alu and L1 elements included in the analysis and sized to match the real data. For the comparison of intra-chromosomal recombination rates in Parkinson’s disease (PD) and Alzheimer’s disease (AD) datasets vs the control dataset in Figure 4h and 4i, we first divided the rate of intra-chromosomal recombination of each chromosome for each tissue in PD and AD datasets (average of 10 donors) by the respective values in the control dataset, then we calculated and plotted the Log2 of the ratios.

To identify recombination hotspots, we maximized the discovery power by merging recombination events detected in the control, Parkinson’s disease and Alzheimer’s disease capture-seq datasets. Calculations for the Recombination Index (RI) and hotness thresholds for each repeat subfamily and each dataset is in Table S6. For instance, in the NAHR dataset for control donors we detected 1191417 individual recombination events involving AluY elements; since there are 139234 AluY elements annotated in the RepeatMasker, a random distribution of the 1191417 NAHR events would theoretically result in ∼8.6 recombination events per AluY (rounded to 9). Hence, we considered as “hot” all the AluY elements participating in a number of NAHR events exceeding this threshold. In the NAHR dataset for control donors, ∼41% of all annotated AluY elements (57408/139234) exceeded this threshold. The chromosome 21 plot with overlay of NAHR events in Figure 3a was made with the R package chromoMap(Anand and Lopez, 2020). To calculate the enrichment of NAHR events in centromeres, we first binned the GRCh38 genome in 100kb bins and we normalized the recombination index of each bin by the repeat elements content of each bin. This was done by first calculating the recombination index sum for all the repeat elements annotated in a bin and participating in at least one NAHR event, and then we divided the RI sum of each bin by the number of repeat elements annotated per bin. This normalization allowed us to rank all genomic bins based on their relative recombination activity. Coordinates for human centromeres were obtained from the UCSC Table Browser, they were extended by ±1Mb with BEDTools *slop* to include peri-centromeric regions and then the extended coordinates were intersected using BEDTools *intersect* with GRCh38 100kb genomic bins in order to annotate the bins proximal to centromeres. Next, for each capture-seq library we counted using BEDTools *intersect* the number of NAHR events in which both repeats, only one repeat or no repeats were contained within a centromeric genomic bin (“observed”). The expected values were calculated by generating 1000 datasets of randomly paired repeat elements to simulate random NAHR events, and similarly counting simulated events having both repeats, only one repeat or no repeats contained within a centromeric genomic bin (“expected”). Finally, for each sample we calculated and plotted the ratio between “observed” and “expected”.

For the analysis of enrichment of recombination hotspots in COSMIC cancer genes we first obtained the COSMIC Cancer Genes Census from the COSMIC website (https://cancer.sanger.ac.uk/cosmic/download) and modified it in BED-like format. We then intersected with BEDTools *intersect* the RepeatMasker database with the COSMIC genes coordinates in order to obtain a list of all Alu and L1HS/PA annotated within COSMIC cancer genes (for Alu: n=36354; for L1HS/PA: n=2134). Next we aggregated the list of 8 capture-seq libraries related to each single donor and obtained the list of Alu and L1 elements with RI≥10 in each aggregated donor dataset. For each donor we then calculated the percentage of hot repeat elements mapped within COSMIC genes using BEDTools *intersect.* To understand how these values compared to non-cancer genes, the percentages of hot elements in COSMIC genes for each donor were compared with the 100 random datasets obtained in the following way. We first compiled a list of Alu and L1HS/PA in non-cancer genes by subtracting the COSMIC cancer genes set from all Refseq genes using BEDTools *subtract* and subsequently intersecting this non-cancer genes list with the RepeatMasker annotations, again with BEDTools *intersect*. We then generated 100 random datasets of Alu and L1HS/PA contained in non-cancer genes, sized as the number of Alu and L1 in COSMIC genes listed above (n=36354 and n=2134). For each tissue of each donor we next calculated the percentage of hot Alu and L1 elements int each random dataset with BEDTools *intersect* and these values were used for the plots in Figure 3e, 3f, 16e, 16f, 17e and 17f).

For the analysis of recombination in A and B compartments in human iPSC and differentiated neurons, we first obtained the genomic coordinates of A and B compartments from Lu et al.(Lu et al., 2020) and converted them from GRCh37 to GRCh38 using UCSC LiftOver program (https://genome-store.ucsc.edu/). We next intersected with BEDTools intersect the compartments coordinates in human iPSC and neurons from Lu et al., in order to obtain a set of compartments that stay constant during the differentiation process as to avoid potential confounding effect from different proportions of A and B compartments in the two cell types. Next, using BEDTools intersect we intersected the genomic coordinates of NAHR events in capture-seq datasets of iPSC and differentiated neurons with the modified set of A/B compartments described above and calculated all combinations the relative abundance of NAHR events in each compartment.

### Quantification and statistical analysis

All details about the statistical analyses used in this study are indicated in the respective Figure legends. Mann–Whitney U test was performed in R. Student’s T-test (in all analyses: unpaired, two-tailed) and One-Way ANOVA test were performed in Excel. Randomized datasets were generated from the real datasets with varying number of iterations tailored for each specific testing and always comparably sized according to real dataset to avoid any potential bias.

## Supplemental items titles and description

**Supplementary Document 1: Capture-seq protocol**

This document contains the extended technical description of the capture-seq protocol performed in this paper, including all steps and required reagents.

**Supplementary Document 2:** Sequence of oligonucleotides for in vitro transcription of biotinylated riboprobes used for capture-seq

**Supplementary Document 3: FANS protocol**

This document contains a detailed technical description of steps and required reagents related to sorting of nuclear fractions performed in this paper.

**Supplementary Document 4: Sanger sequencing results**

This document contains all sequences related to capillary sequencing of purified PCR amplicons for the validation of recombination events.

## Supplementary Tables

Table S1: Information related to all human donors for the post-mortem tissues used in this study

Table S2: raw counts for all FASTQ files produced by sequencing experiments performed in this study

Table S3: list of all repeat elements included in the analyses of capture-seq datasets

Table S4: Raw and normalized count of recombination events per library

Table S5: Extended details of PCR validations including oligonucleotide sequences for all targets

Table S6: Extended details for the calculation of Recombination Index and thresholds in Control, Parkinson and Alzheimer capture-seq datasets

