## Supplementary Tables S1-S6 for "Recombination of repeat elements generates somatic complexity in human genomes"

#### Table S1 - Donor information

| Group | Donor | Age | Sex | Brain weight (grams) | ApoE | Grain | Lewy bodies | Time after death | Braak NFT | Braak stage |
|---|---|---|---|---|---|---|---|---|---|---|
| Control | 1 | 79 | M | 1180 | e3/e3 | 0 | 0 | 1:20:00 | 2 | 0 |
| Control | 2 | 79 | F | 1178 | e3/e3 | 0 | 0 | 1:35:00 | 1 | 0 |
| Control | 3 | 81 | M | 1170 | e2/e3 | 0 | 0 | 2:14:00 | 1 | 0 |
| Control | 4 | 84 | F | 1079 | e2/e3 | 0.5/ 0 | 0 | 4:36:00 | 1/2.5* | 1 |
| Control | 5 | 76 | M | 1314 | e3/e3 | 1 | 0 | 3:45:00 | 2 | 1 |
| Control | 6 | 82 | F | 1470 | e3/e3 | 0? | 0 | 17:53:00 | 1 | 1 |
| Control | 7 | 89 | M | 1180 | e3/e3 | 0.5 | 0 | 4:10:00 | 2 | 1 |
| Control | 8 | 90 | M | 1186 | e3/e4 | 0 | 0 | 5:05:00 | 1 | 1 |
| Control | 9 | 83 | M | 1440 | e3/e3 | 1 | 0 | 17:43:00 | 1 | 0 |
| Control | 10 | 82 | F | 1300 | e3/e3 | 1 | 0 | 12:04:00 | 2/2.5 | 1 |
| Parkinson | 1 | 85 | M | 1440 | e3/e3 | 0 | 4 | 14:42:00 | 2 | 0 |
| Parkinson | 2 | 79 | F | 1094 | e3/e3 | 0.5 | 4 | 4:52:00 | 1 | 1 |
| Parkinson | 3 | 79 | M | 1438 | e3/e3 | 0 | 4 | 19:21:00 | 2 | 0 |
| Parkinson | 4 | 80 | F | 1194 | e2/e3 | 0.5/1 | 4 | 35:36:00 | 2/1 | 1 |
| Parkinson | 5 | 81 | M | 1208 | e3/e3 | 0.5 | 3 | 12:32:00 | 2 | 1 |
| Parkinson | 6 | 92 | M | 1005 | e3/e4 | 3 | 5 | 10:02:00 | 2 | 1 |
| Parkinson | 7 | 83 | F | 1158 | e3/e3 | 0 | 4 | 4:18:00 | 3 | 0 |
| Parkinson | 8 | 83 | M | 1170 | e3/e3 | 0.5 | 3 | 3:02:00 | 3? | 1 |
| Parkinson | 9 | 88 | M | 1308 | e2/e3 | 1 | 3 | 18:45:00 | 2/2 | 1 |
| Parkinson | 10 | 89 | M | 1202 | e3/e3 | 0.5 | 4?(R4?) | 2:02:00 | 2 | 1 |
| Alzheimer | 1 | 84 | M | 1320 | e3/e4 | 0.5 | 0 | 9:50:00 | 5 | 3 |
| Alzheimer | 2 | 81 | M | 1450 | e3/e3 | 0.5 | 0 | 6:00:00 | 6 | 3 |
| Alzheimer | 3 | 79 | F | 1100 | e3/e3 | 0 | 0 | 3:23:00 | 5 | 3 |
| Alzheimer | 4 | 82 | F | 1091 | e4/e4 | 1 | 0 | 19:45:00 | 5 | 3 |
| Alzheimer | 5 | 81 | F | 1127 | e3/e3 | 1/0.5 | 0 | 7:22:00 | 4/>=5 | 3 |
| Alzheimer | 6 | 83 | F | 1280 | e3/e3 | 2 | 0 | 59:25:00 | 5 | 3 |
| Alzheimer | 7 | 85 | F | 991 | e3/e4 | 2 | 0 | 14:20:00 | 4.5 | 3 |
| Alzheimer | 8 | 87 | M | 1040 | e3/e4 | 2.5 | 0 | 3:17:00 | 6 | 3 |
| Alzheimer | 9 | 88 | M | 1206 | e4/e4 | 0 | 0 | 2:08:00 | 6 | 3 |
| Alzheimer | 10 | 86 | M | 1248 | e3/e3 | 1 | 0 | 12:55:00 | 6 | 3 |

Legend:

M: male; F: female
ApoE: apolipoprotein E genetic alleles assessment
Grain: evaluation score of Argyrophilic grain disease
Lewy Bodies: evaluation score for the presence of Lewy Bodies
Braak NFT: evaluation of presence of Neurofibrillary Tangles (NFT)
Braak stage: evaluation of disease progression stage based on Braak scale.

### Table S2 - Fastq raw reads count for all datasets in this study

#### Capture-seq of post-mortem samples (Illumina Hiseq4000, 150 bp reads, paired-end)

| | Control | | | | | | | |
|---|---|---|---|---|---|---|---|---|
| | Kidney | Liver | FC NeuN- | FC NeuN+ | TC NeuN- | TC NeuN+ | PC NeuN- | PC NeuN+ |
| Donor 1 | 13886876 | 16751401 | 8319762 | 9971769 | 12168696 | 10577084 | 14388452 | 15562239 |
| Donor 2 | 12174825 | 8667954 | 9506536 | 12624906 | 11923342 | 13325304 | 10846104 | n/a |
| Donor 3 | 16181126 | 9182801 | 11300903 | 10514003 | 11563502 | 13082196 | 13896835 | 12582243 |
| Donor 4 | n/a | 12550797 | 10051818 | 16264770 | 12737695 | 11085470 | 14085403 | 11986430 |
| Donor 5 | 12864001 | 11953807 | 16166805 | 10012704 | 11409267 | 10489071 | 10941895 | 15203963 |
| Donor 6 | 11180354 | 11795299 | 9147665 | 12329208 | 12559417 | 10289026 | 16759026 | 14294414 |
| Donor 7 | 11350117 | 12328431 | 10329965 | 10391009 | 10346476 | 13894117 | 17117030 | 12820690 |
| Donor 8 | 11715626 | 11700812 | 13122410 | 10887185 | 9614989 | 12628239 | 11880716 | 12709182 |
| Donor 9 | 20248909 | 13597636 | 10358636 | 12293499 | 11550556 | 9785749 | 15388466 | 11938734 |
| Donor 10 | 11445517 | 11727255 | 14124099 | 13304916 | 13032848 | 9994702 | 13188491 | 11973950 |

| | Parkinson's disease | | | | | | | |
|---|---|---|---|---|---|---|---|---|
| | Kidney | Liver | FC NeuN- | FC NeuN+ | TC NeuN- | TC NeuN+ | PC NeuN- | PC NeuN+ |
| Donor 1 | 12344160 | 12954968 | 10355780 | 11693291 | 12047292 | 13116712 | 12432237 | 13269917 |
| Donor 2 | 14101171 | 12430550 | 14150420 | 11344129 | 12233687 | 11152273 | 10145519 | 13007388 |
| Donor 3 | 12656431 | 11935434 | 10414741 | 12299653 | 11596810 | 12561777 | 9940556 | 13668649 |
| Donor 4 | 10704656 | 12360217 | 11587749 | 11385792 | 11522856 | 10825427 | 12977562 | 16353631 |
| Donor 5 | 13536778 | 13796981 | 12207324 | 10534437 | 15608646 | 10297276 | 11258821 | 12354136 |
| Donor 6 | n/a | 12854592 | 11594692 | 15570649 | 13148044 | 12579676 | 12673691 | 14465204 |
| Donor 7 | 9444830 | 12072063 | 12992834 | 11163934 | 10807007 | 11058964 | 11505456 | 10235752 |
| Donor 8 | 11526061 | 11386772 | 10992237 | 11902682 | 11323240 | 13946355 | 11531732 | 12442652 |
| Donor 9 | 14716508 | 10358168 | 12970858 | 12061672 | 9434818 | 11450533 | 11364536 | 12970614 |
| Donor 10 | 13208829 | 11402997 | 12889050 | 12257597 | 10800522 | 9768341 | 11353316 | 13626482 |

| | Alzheimer's disease | | | | | | | |
|---|---|---|---|---|---|---|---|---|
| | Kidney | Liver | FC NeuN- | FC NeuN+ | TC NeuN- | TC NeuN+ | PC NeuN- | PC NeuN+ |
| Donor 1 | 13386248 | 13823449 | 10642376 | 10632884 | 12878543 | 12887943 | 11595802 | 15213277 |
| Donor 2 | 10519575 | 9826953 | 11207746 | 15279100 | 12274921 | 12576258 | 13015796 | 11012717 |
| Donor 3 | 12942521 | 11222391 | 12963439 | 11353513 | 12960713 | 12296578 | 12165658 | 13800619 |
| Donor 4 | 13812976 | 12432782 | 9030947 | 14121160 | 12080019 | 9827534 | 12679010 | 12912298 |
| Donor 5 | 10411139 | 10212822 | 12558552 | 12683504 | 13714124 | 10388631 | 11520780 | 13856359 |
| Donor 6 | 11461019 | 14323377 | 9519033 | 12345642 | 12761110 | 12210782 | 12189359 | 17544840 |
| Donor 7 | 11696846 | 12595115 | 10298690 | 11253199 | 11505489 | 13659153 | 12556228 | 14619612 |
| Donor 8 | 12974066 | 15131936 | 9703391 | 11880676 | 10820722 | 11252484 | 11525328 | 12580843 |
| Donor 9 | 11944623 | 9922838 | 10838347 | 12722938 | 10928914 | 18478067 | 12572465 | 12938716 |
| Donor 10 | 12779835 | 13798837 | 11949143 | 15995381 | 13130482 | 10767656 | 12285620 | 12880689 |

Legend:
FC = Frontal Cortex
TC = Temporal Cortex
PC = Parietal Cortex
n/a = sample not available for sequencing due to exhaustion of genomic DNA during library preparation.
Library with lowest reads count used as reference for normalization by subsampling

#### Capture-seq of induced pluripotent stem cells (iPSC) differentiated to neurons (iNEU) (Illumina Miseq, 300 bp, paired-end)

| | iPSC | Neurons |
|---|---|---|
| rep1 | 6075266 | 6174685 |
| rep2 | 5466470 | 5060940 |
| rep3 | 5152590 | 4254782 |

#### Oxford Nanopore Technologies PromethION WGS libraries (count of fastq reads that passed QC)

| | | | TC NeuN+ | |
|---|---|---|---|---|
| | Kidney | Liver | Control | AD |
| Donor 1 | | | 4697267 | 4268929 |
| Donor 2 | | | 6137932 | 3807476 |
| Donor 3 | | | 5887275 | 6180965 |
| Donor 4 | | | 1507391 | 4766859 |
| Donor 5 | | | 4250178 | 1491867 |
| Donor 6 | 19481750 | 22315607 | 22408736 | 1584423 |
| Donor 7 | | | 4613006 | 14144382 |
| Donor 8 | 23369605 | 24391131 | 21959340 | 5491734 |
| Donor 9 | | | 1306817 | 859778 |
| Donor 10 | 20752367 | 15998265 | 18852702 | 8924069 |

TC = Temporal Cortex

**Table S3 - List of repeat elements included in the analyses.**

NOTE: Color-coding for L1 elements indicates the 3 L1 groups in Figure 1c
* subfamilies annotated in RepeatMasker not included in the analyses because they were not present in Dfam Database.

| ALU | |
|---|---|
| AluJb | 131759 |
| AluSx1 | 123492 |
| AluSx | 123022 |
| AluY | 110881 |
| AluSz | 107707 |
| AluJr | 88503 |
| AluJo | 81375 |
| AluSq2 | 63875 |
| AluSp | 53809 |
| AluSz6 | 49944 |
| AluSg | 38681 |
| AluSc | 36338 |
| AluSx3 | 34198 |
| AluSc8 | 23028 |
| AluJr4 | 20966 |
| AluSq | 19866 |
| AluSx4 | 11540 |
| AluSg7 | 9159 |
| AluSg4 | 7603 |
| AluSc5 | 7018 |
| AluYm1 | 5041 |
| AluYc | 4810 |
| Alu * | 4658 |
| AluYa5 | 3986 |
| AluYj4 | 3860 |
| AluYb8 | 2962 |
| AluYh3 | 2798 |
| AluSq10 | 2165 |
| AluYf1 | 2025 |
| AluSq4 | 1906 |
| AluYe5 | 1378 |
| AluYk11 | 1341 |
| AluYk3 | 1207 |
| AluYk4 | 1051 |
| AluYg6 | 899 |
| AluYk2 | 824 |
| AluYc3 | 566 |
| AluYi6 | 470 |
| AluYa8 | 368 |
| AluYh3a3 * | 342 |
| AluYb9 | 339 |
| AluYd8 | 241 |
| AluYk12 | 219 |
| AluYe6 | 199 |
| AluYh7 | 183 |
| AluYh9 | 165 |
| AluYi6_4d | 153 |

| L1 | |
|---|---|
| L1HS | 1686 |
| L1PA2 | 5113 |
| L1PA3 | 11089 |
| L1PA4 | 12272 |
| L1PA5 | 11616 |
| L1PA6 | 6143 |
| L1PA7 | 13381 |
| L1PA8 | 8376 |
| L1PA8A | 2514 |
| L1PA10 | 7367 |
| L1PA11 | 4207 |
| L1PA12 | 1811 |
| L1PA13 | 9208 |
| L1PA14 | 3116 |
| L1PA15 | 8569 |
| L1PA16 | 14421 |
| L1PA17 | 4863 |
| L1PA15-16 * | 1454 |

#### Table S4 - Raw and normalized count of recombination events per library.

NOTE: Normalization by sequencing depth for capture-seq libraries performed by subsampling as indicated in Table S3.
Normalization for ONT libraries perfomred by recombination-per-million reads

#### Raw Counts

##### Capture-seq libraries, post-mortem tissues, putative somatic

**ALU**

| Donor | Control Kidney | Control Liver | Control FC NeuN- | Control FC NeuN+ | Control TC NeuN- | Control TC NeuN+ | Control PC NeuN- | Control PC NeuN+ |
|---|---|---|---|---|---|---|---|---|
| D1 | 41969 | 71083 | 6605 | 4087 | 8824 | 4948 | 87518 | 50511 |
| D2 | 39526 | 30271 | 5310 | 15000 | 14665 | 9936 | 14715 | n/a |
| D3 | 78601 | 65516 | 5485 | 8053 | 9383 | 6086 | 73260 | 76668 |
| D4 | n/a | 117327 | 5289 | 51766 | 9534 | 10005 | 6864 | 6476 |
| D5 | 21229 | 24330 | 32459 | 68684 | 6230 | 4977 | 6748 | 9234 |
| D6 | 44374 | 51593 | 17138 | 65600 | 8391 | 8130 | 25050 | 10530 |
| D7 | 35935 | 44306 | 9135 | 9592 | 10341 | 18877 | 22762 | 9934 |
| D8 | 19284 | 45638 | 6386 | 6364 | 14050 | 13174 | 9501 | 9151 |
| D9 | 154018 | 57390 | 6443 | 7296 | 5604 | 4737 | 20315 | 22480 |
| D10 | 59333 | 37803 | 16056 | 21205 | 16400 | 9555 | 18967 | 16203 |

| Donor | Parkinson Kidney | Parkinson Liver | Parkinson FC NeuN- | Parkinson FC NeuN+ | Parkinson TC NeuN- | Parkinson TC NeuN+ | Parkinson PC NeuN- | Parkinson PC NeuN+ |
|---|---|---|---|---|---|---|---|---|
| D1 | 28524 | 27795 | 10032 | 12528 | 8066 | 6881 | 8818 | 10597 |
| D2 | 67900 | 78147 | 6200 | 6129 | 20970 | 11632 | 7368 | 10153 |
| D3 | 67532 | 45408 | 11361 | 16626 | 13114 | 13989 | 11519 | 15032 |
| D4 | 27095 | 45695 | 21919 | 49954 | 6017 | 7740 | 4732 | 7690 |
| D5 | 131209 | 63507 | 6191 | 5820 | 6513 | 11162 | 9667 | 6589 |
| D6 | n/a | 56102 | 6550 | 10158 | 25246 | 11643 | 13568 | 5997 |
| D7 | 58673 | 49315 | 14337 | 13997 | 14935 | 10527 | 9328 | 13106 |
| D8 | 39272 | 40173 | 53189 | 59175 | 12545 | 10379 | 8467 | 8303 |
| D9 | 60958 | 45008 | 15688 | 12450 | 17421 | 17379 | 9382 | 13030 |
| D10 | 83735 | 56553 | 8149 | 4082 | 11932 | 4764 | 7516 | 8524 |

| Donor | Alzheimer Kidney | Alzheimer Liver | Alzheimer FC NeuN- | Alzheimer FC NeuN+ | Alzheimer TC NeuN- | Alzheimer TC NeuN+ | Alzheimer PC NeuN- | Alzheimer PC NeuN+ |
|---|---|---|---|---|---|---|---|---|
| D1 | 56993 | 31398 | 12683 | 8163 | 10958 | 15717 | 12053 | 4055 |
| D2 | 21890 | 13215 | 6251 | 6498 | 18661 | 15862 | 25813 | 9503 |
| D3 | 35036 | 28404 | 7985 | 12860 | 9398 | 22416 | 22617 | 13352 |
| D4 | 70276 | 29346 | 10752 | 26588 | 11782 | 12099 | 10381 | 6842 |
| D5 | 19867 | 44002 | 4824 | 18777 | 17380 | 19225 | 17373 | 7658 |
| D6 | 12318 | 61539 | 8413 | 7230 | 19550 | 14740 | 11480 | 37807 |
| D7 | 26368 | 67257 | 14493 | 16368 | 16149 | 20841 | 22573 | 11236 |
| D8 | 10563 | 41013 | 12399 | 4176 | 15165 | 7797 | 35973 | 16629 |
| D9 | 8163 | 28616 | 9449 | 13450 | 12744 | 32210 | 14960 | 10822 |
| D10 | 16316 | 18235 | 9481 | 12869 | 17315 | 8963 | 13304 | 6208 |

**L1**

| Donor | Control Kidney | Control Liver | Control FC NeuN- | Control FC NeuN+ | Control TC NeuN- | Control TC NeuN+ | Control PC NeuN- | Control PC NeuN+ |
|---|---|---|---|---|---|---|---|---|
| D1 | 1409 | 3753 | 786 | 256 | 1111 | 430 | 13243 | 6087 |
| D2 | 5894 | 4140 | 919 | 1386 | 1793 | 1812 | 2045 | n/a |
| D3 | 4034 | 5505 | 936 | 723 | 1635 | 794 | 9559 | 38173 |
| D4 | n/a | 15089 | 535 | 4714 | 820 | 1051 | 528 | 452 |
| D5 | 1442 | 3943 | 9440 | 2848 | 581 | 480 | 465 | 658 |
| D6 | 4309 | 5670 | 1351 | 13431 | 929 | 768 | 1130 | 587 |
| D7 | 4420 | 4931 | 798 | 896 | 985 | 2545 | 608 | 401 |
| D8 | 2712 | 4091 | 451 | 336 | 891 | 1672 | 769 | 770 |
| D9 | 6611 | 5768 | 491 | 488 | 356 | 290 | 1706 | 3134 |
| D10 | 5331 | 4779 | 1237 | 3599 | 1513 | 946 | 1342 | 1318 |

| Donor | Parkinson Kidney | Parkinson Liver | Parkinson FC NeuN- | Parkinson FC NeuN+ | Parkinson TC NeuN- | Parkinson TC NeuN+ | Parkinson PC NeuN- | Parkinson PC NeuN+ |
|---|---|---|---|---|---|---|---|---|
| D1 | 1803 | 2650 | 723 | 992 | 752 | 781 | 608 | 964 |
| D2 | 5843 | 4560 | 752 | 358 | 2488 | 900 | 571 | 1782 |
| D3 | 5025 | 3803 | 666 | 3085 | 1123 | 1329 | 1603 | 1689 |
| D4 | 2905 | 3864 | 3080 | 5877 | 391 | 517 | 407 | 648 |
| D5 | 18013 | 4930 | 730 | 428 | 562 | 711 | 1148 | 575 |
| D6 | n/a | 4454 | 459 | 970 | 3611 | 1098 | 1364 | 454 |
| D7 | 3012 | 4827 | 978 | 1279 | 993 | 1235 | 614 | 1094 |
| D8 | 2462 | 6226 | 7611 | 4362 | 1230 | 673 | 831 | 551 |
| D9 | 2793 | 3411 | 1823 | 1644 | 1377 | 1486 | 608 | 747 |
| D10 | 8802 | 5437 | 396 | 250 | 586 | 300 | 348 | 440 |

| Donor | Alzheimer Kidney | Alzheimer Liver | Alzheimer FC NeuN- | Alzheimer FC NeuN+ | Alzheimer TC NeuN- | Alzheimer TC NeuN+ | Alzheimer PC NeuN- | Alzheimer PC NeuN+ |
|---|---|---|---|---|---|---|---|---|
| D1 | 4108 | 1519 | 961 | 750 | 1339 | 1342 | 1454 | 595 |
| D2 | 1717 | 896 | 690 | 693 | 3720 | 1323 | 1844 | 1206 |
| D3 | 1743 | 3491 | 1397 | 1222 | 1682 | 2709 | 3402 | 1616 |
| D4 | 4504 | 2278 | 1046 | 3382 | 2629 | 2268 | 1308 | 596 |
| D5 | 2250 | 5141 | 531 | 2854 | 1251 | 1677 | 1821 | 1009 |
| D6 | 1024 | 5565 | 462 | 296 | 2039 | 1522 | 1967 | 2370 |
| D7 | 2707 | 24522 | 1108 | 1558 | 1390 | 2844 | 2103 | 1264 |
| D8 | 512 | 4139 | 2190 | 364 | 2095 | 574 | 2949 | 579 |
| D9 | 389 | 3700 | 439 | 736 | 725 | 2430 | 1454 | 618 |
| D10 | 597 | 1077 | 636 | 1247 | 1811 | 854 | 1113 | 816 |

##### Capture-seq libraries, iPSC and induced Neurons

| | Alu iPSC | Alu iNEU | L1 iPSC | L1 iNEU |
|---|---|---|---|---|
| Rep1 | 23337 | 33945 | 1131 | 1460 |
| Rep2 | 23725 | 23507 | 1119 | 970 |
| Rep3 | 22783 | 18881 | 1047 | 1123 |

##### Oxford Nanopore Technologies PromethION WGS libraries

| Donor | Kidney | Liver | TC NeuN+ Control | TC NeuN+ AD |
|---|---|---|---|---|
| D1 | | | 469 | 1717 |
| D2 | | | 3520 | 2672 |
| D3 | | | 2471 | 1881 |
| D4 | | | 594 | 1454 |
| D5 | | | 715 | 317 |
| D6 | 5813 | 32356 | 7353 | 157 |
| D7 | | | 1921 | 1938 |
| D8 | 7615 | 26420 | 7239 | 2626 |
| D9 | | | 579 | 99 |
| D10 | 9356 | 25582 | 10847 | 2437 |

#### Normalized Counts

##### Capture-seq, post-mortem tissues, subsampled libraries, putative somatic . Estimated NAHR count per cell based on genomic DNA input per library (See Methods section for details).

**ALU**

| Donor | Control Kidney | Control Liver | Control FC NeuN- | Control FC NeuN+ | Control TC NeuN- | Control TC NeuN+ | Control PC NeuN- | Control PC NeuN+ |
|---|---|---|---|---|---|---|---|---|
| D1 | 2.69353846 | 3.82144615 | 0.61984615 | 0.34070769 | 0.62852308 | 0.41492308 | 5.38698462 | 2.93926154 |
| D2 | 2.76590769 | 2.68015385 | 0.45203077 | 1.03338462 | 1.05443077 | 0.70661538 | 0.92270769 | n/a |
| D3 | 4.47166154 | 5.52424615 | 0.41926154 | 0.636 | 0.68224615 | 0.41326154 | 5.64036923 | 5.20107692 |
| D4 | n/a | 7.9308 | 0.43375385 | 2.95107692 | 0.65316923 | 0.76596923 | 0.43504615 | 0.45553846 |
| D5 | 1.46289231 | 1.74443077 | 1.90504615 | 5.57030769 | 0.46255385 | 0.40412308 | 0.5268 | 0.54507692 |
| D6 | 3.29132308 | 3.71298462 | 1.50221538 | 4.57726154 | 0.59141538 | 0.65510769 | 1.35378462 | 0.65778462 |
| D7 | 2.61812308 | 3.02095385 | 0.73310769 | 0.76476923 | 0.85947692 | 1.22963077 | 1.20313846 | 0.66627692 |
| D8 | 1.37981538 | 3.2508 | 0.42830769 | 0.48618462 | 1.20701538 | 0.93276923 | 0.69083077 | 0.62169231 |
| D9 | 7.35590769 | 3.70873846 | 0.51230769 | 0.50390769 | 0.40966154 | 0.39978462 | 1.24624615 | 1.62147692 |
| D10 | 4.34455385 | 2.70378462 | 1.01529231 | 1.38369231 | 1.08849231 | 0.77556923 | 1.254 | 1.16723077 |

| Donor | Parkinson Kidney | Parkinson Liver | Parkinson FC NeuN- | Parkinson FC NeuN+ | Parkinson TC NeuN- | Parkinson TC NeuN+ | Parkinson PC NeuN- | Parkinson PC NeuN+ |
|---|---|---|---|---|---|---|---|---|
| D1 | 1.99836923 | 1.85206154 | 0.83658462 | 0.918 | 0.58569231 | 0.46615385 | 0.62676923 | 0.69535385 |
| D2 | 4.21190769 | 5.39963077 | 0.39267692 | 0.48470769 | 1.47 | 0.89003077 | 0.60101538 | 0.70181538 |
| D3 | 4.58723077 | 3.17169231 | 0.90821538 | 1.19206154 | 0.95390769 | 0.96812308 | 0.95326154 | 0.98667692 |
| D4 | 2.08227692 | 3.13901538 | 1.61981538 | 3.78867692 | 0.44132308 | 0.58366154 | 0.32621538 | 0.43670769 |
| D5 | 8.40516923 | 4.05203077 | 0.43883077 | 0.46532308 | 0.37809231 | 0.90027692 | 0.72230769 | 0.46264615 |
| D6 | n/a | 3.72156923 | 0.48018462 | 0.58827692 | 1.68138462 | 0.80243077 | 0.92658462 | 0.37366154 |
| D7 | 5.01664615 | 3.4428 | 0.97938462 | 1.05747692 | 1.14387692 | 0.82449231 | 0.68390769 | 1.03929231 |
| D8 | 2.89689231 | 2.91230769 | 4.04215385 | 4.25150769 | 0.94790769 | 0.66507692 | 0.60526154 | 0.57406154 |
| D9 | 3.75332308 | 3.48609231 | 1.05166154 | 0.90406154 | 1.47683077 | 1.29415385 | 0.70910769 | 0.87341538 |
| D10 | 5.51566154 | 4.10898462 | 0.55024615 | 0.28892308 | 0.92187692 | 0.40809231 | 0.55929231 | 0.55255385 |

| Donor | Alzheimer Kidney | Alzheimer Liver | Alzheimer FC NeuN- | Alzheimer FC NeuN+ | Alzheimer TC NeuN- | Alzheimer TC NeuN+ | Alzheimer PC NeuN- | Alzheimer PC NeuN+ |
|---|---|---|---|---|---|---|---|---|
| D1 | 3.76938462 | 1.97750769 | 0.98593846 | 0.68861538 | 0.75756923 | 1.07695385 | 0.89723077 | 0.25366154 |
| D2 | 1.71544615 | 1.07353846 | 0.46329231 | 0.39784615 | 1.338 | 1.11830769 | 1.70704615 | 0.72895385 |
| D3 | 2.36907692 | 2.10064615 | 0.54396923 | 0.95233846 | 0.64643077 | 1.58861538 | 1.57836923 | 0.87553846 |
| D4 | 4.59036923 | 2.0268 | 0.94846154 | 1.68055385 | 0.85310769 | 1.07196923 | 0.70024615 | 0.46873846 |
| D5 | 1.57790769 | 3.48018462 | 0.33498462 | 1.27790769 | 1.09809231 | 1.52713846 | 1.27015385 | 0.51369231 |
| D6 | 0.91910769 | 3.87110769 | 0.7128 | 0.50713846 | 1.36624615 | 1.05147692 | 0.85550769 | 1.88012308 |
| D7 | 1.89886154 | 4.55307692 | 1.14775385 | 1.10575385 | 1.18107692 | 1.32978462 | 1.55953846 | 0.69609231 |
| D8 | 0.70246154 | 2.47273846 | 1.05701538 | 0.30701538 | 1.17987692 | 0.58633846 | 2.61203077 | 1.19224615 |
| D9 | 0.58569231 | 2.32153846 | 0.71455385 | 1.03015385 | 0.99064615 | 1.67436923 | 1.0284 | 0.74363077 |
| D10 | 1.11895385 | 1.18181538 | 0.68538462 | 0.73643077 | 1.1628 | 0.70070769 | 0.92889231 | 0.45867692 |

**L1**

| Donor | Control Kidney | Control Liver | Control FC NeuN- | Control FC NeuN+ | Control TC NeuN- | Control TC NeuN+ | Control PC NeuN- | Control PC NeuN+ |
|---|---|---|---|---|---|---|---|---|
| D1 | 0.03095385 | 0.07012308 | 0.02541538 | 0.00766154 | 0.02827692 | 0.0136 | 0.28347692 | 0.12553846 |
| D2 | 0.14083077 | 0.12193846 | 0.02563077 | 0.03138462 | 0.04310769 | 0.04393846 | 0.04387692 | n/a |
| D3 | 0.07753846 | 0.15495385 | 0.02436923 | 0.01873846 | 0.04061538 | 0.01913846 | 0.25369231 | 0.87187692 |
| D4 | n/a | 0.34809231 | 0.01467692 | 0.09249231 | 0.01821538 | 0.02676923 | 0.01083077 | 0.01123077 |
| D5 | 0.03332308 | 0.09486154 | 0.19366154 | 0.07929231 | 0.01406154 | 0.01301538 | 0.01258462 | 0.01175385 |
| D6 | 0.10867692 | 0.13661538 | 0.04012308 | 0.32649231 | 0.02353846 | 0.02129231 | 0.02101538 | 0.01196923 |
| D7 | 0.10803077 | 0.11224615 | 0.02193846 | 0.02433846 | 0.02861538 | 0.05701538 | 0.01046154 | 0.00873846 |
| D8 | 0.06633846 | 0.09892308 | 0.00923077 | 0.00830769 | 0.02787692 | 0.04255385 | 0.02036923 | 0.01695385 |
| D9 | 0.10806154 | 0.12612308 | 0.01267692 | 0.012 | 0.00830769 | 0.0076 | 0.03710769 | 0.07673846 |
| D10 | 0.13175385 | 0.11596923 | 0.02655385 | 0.07978462 | 0.03301538 | 0.02609231 | 0.03070769 | 0.03153846 |

| Donor | Parkinson Kidney | Parkinson Liver | Parkinson FC NeuN- | Parkinson FC NeuN+ | Parkinson TC NeuN- | Parkinson TC NeuN+ | Parkinson PC NeuN- | Parkinson PC NeuN+ |
|---|---|---|---|---|---|---|---|---|
| D1 | 0.04206154 | 0.05947692 | 0.02363077 | 0.02532308 | 0.01843077 | 0.01818462 | 0.01378462 | 0.02107692 |
| D2 | 0.12366154 | 0.10516923 | 0.01569231 | 0.01113846 | 0.0588 | 0.02572308 | 0.01587692 | 0.04323077 |
| D3 | 0.11569231 | 0.08910769 | 0.01827692 | 0.07621538 | 0.02713846 | 0.03218462 | 0.04449231 | 0.03827692 |
| D4 | 0.07510769 | 0.08806154 | 0.08061538 | 0.15310769 | 0.00898462 | 0.01295385 | 0.00895385 | 0.01233846 |
| D5 | 0.39076923 | 0.10772308 | 0.01713846 | 0.01067692 | 0.01104615 | 0.02089231 | 0.02843077 | 0.01350769 |
| D6 | n/a | 0.10018462 | 0.01116923 | 0.01953846 | 0.08101538 | 0.02430769 | 0.03101538 | 0.0096 |
| D7 | 0.08486154 | 0.11378462 | 0.02206154 | 0.03350769 | 0.02572308 | 0.03249231 | 0.01510769 | 0.02827692 |
| D8 | 0.06126154 | 0.15206154 | 0.20249231 | 0.11073846 | 0.03369231 | 0.01541538 | 0.01901538 | 0.01236923 |
| D9 | 0.05732308 | 0.08803077 | 0.04172308 | 0.04070769 | 0.03975385 | 0.03744615 | 0.01510769 | 0.01704615 |
| D10 | 0.19513846 | 0.13116923 | 0.00852308 | 0.00544615 | 0.016 | 0.00886154 | 0.00855385 | 0.00907692 |

| Donor | Alzheimer Kidney | Alzheimer Liver | Alzheimer FC NeuN- | Alzheimer FC NeuN+ | Alzheimer TC NeuN- | Alzheimer TC NeuN+ | Alzheimer PC NeuN- | Alzheimer PC NeuN+ |
|---|---|---|---|---|---|---|---|---|
| D1 | 0.09243077 | 0.03163077 | 0.02535385 | 0.02516923 | 0.03033846 | 0.03058462 | 0.03809231 | 0.01116923 |
| D2 | 0.04464615 | 0.02446154 | 0.01673846 | 0.01643077 | 0.09141538 | 0.03212308 | 0.04138462 | 0.03098462 |
| D3 | 0.03984615 | 0.08547692 | 0.03166154 | 0.03113846 | 0.04110769 | 0.06418462 | 0.07996923 | 0.03492308 |
| D4 | 0.10083077 | 0.05298462 | 0.03190769 | 0.07433846 | 0.06498462 | 0.07356923 | 0.03083077 | 0.01316923 |
| D5 | 0.05969231 | 0.13646154 | 0.012 | 0.06716923 | 0.02584615 | 0.0472 | 0.04421538 | 0.02452308 |
| D6 | 0.02575385 | 0.12113846 | 0.01329231 | 0.00661538 | 0.05024615 | 0.03652308 | 0.05298462 | 0.03950769 |
| D7 | 0.06566154 | 0.56141538 | 0.02916923 | 0.03572308 | 0.03409231 | 0.06286154 | 0.04978462 | 0.02643077 |
| D8 | 0.01135385 | 0.08707692 | 0.06403077 | 0.00843077 | 0.05593846 | 0.01516923 | 0.07483077 | 0.01341538 |
| D9 | 0.00944615 | 0.09987692 | 0.01086154 | 0.02064615 | 0.01975385 | 0.04501538 | 0.03415385 | 0.01523077 |
| D10 | 0.01378462 | 0.02415385 | 0.01523077 | 0.02289231 | 0.04083077 | 0.02258462 | 0.02621538 | 0.02089231 |

##### Capture-seq libraries, iPSC and induced Neurons, NAHR per million reads

| | Alu iPSC | Alu iNEU | L1 iPSC | L1 iNEU |
|---|---|---|---|---|
| Rep1 | 3841 | 5497 | 186 | 236 |
| Rep2 | 4340 | 4645 | 205 | 192 |
| Rep3 | 4422 | 4438 | 203 | 264 |

##### Oxford Nanopore Technologies PromethION WGS libraries, NAHR per million reads

| Donor | Kidney | Liver | TC NeuN+ Control | TC NeuN+ AD |
|---|---|---|---|---|
| D1 | | | 100 | 403 |
| D2 | | | 574 | 703 |
| D3 | | | 420 | 305 |
| D4 | | | 394 | 305 |
| D5 | | | 168 | 214 |
| D6 | 299 | 1602 | 329 | 99 |
| D7 | | | 417 | 137 |
| D8 | 327 | 1453 | 330 | 479 |
| D9 | | | 445 | 116 |
| D10 | 452 | 1085 | 576 | 273 |

### Table S5 - PCR primers and targets details

NOTE: Detailed Sanger sequencing results available in Supplementary Document 1

**PCR validation of putative somatic NAHR events in capture-seq libraries - LIBRARY PCR**

| Target Name | Donor | Library | Repeat | Exp. Amplicon size | PCR amplicon | Validated by Sanger | FW primer sequence | FW primer name | REV primer sequence | REV primer name | Contig ID | Supporting Contigs |
|---|---|---|---|---|---|---|---|---|---|---|---|---|
| Inter-1 | AD1 | FA1plus | ALU | 262 | Yes | Yes | CATTAAAGTGGGCCCAGTG | HR-INTER-1-FW | TTCTGACAGTTGCTTCCCA | HR-INTER-1-REV | K00114:354:HC225BBXX:5:2106:26646:247 | 1 |
| Inter-2 | AD1 | PA1minus | ALU | 228 | Yes | Yes | CCAAATGACTGCTCTCCCTG | HR-INTER-2-FW | CGCAGTAGGTGGACAATAAGC | HR-INTER-2-REV | K00114:354:HC225BBXX:5:1106:8065:4581 | 1 |
| Inter-3 | Control1 | FC1minus | ALU | 245 | Yes | Yes | TGAAACCAGATATTTGAAACTTACC | HR-INTER-3-FW | CGACCCTCCCATCTCAGC | HR-INTER-3-REV | K00114:354:HC225BBXX:3:2214:16985:300 | 1 |
| Inter-4 | Control1 | FC1plus | ALU | 221 | Yes | Yes | AAACTATGAGGAGCAGGCTT | HR-INTER-4-FW | GGTGGTCAGCAGCTCTAAAC | HR-INTER-4-REV | K00114:354:HC225BBXX:7:1222:15371:225 | 1 |
| Inter-5 | Control10 | LC10 | ALU | 149 | Yes | Yes | GCAATACTAAGAGAAAGGATGGA | HR-INTER-5-FW | GGCAAGAATCCTGACCCA | HR-INTER-5-REV | K00114:354:HC225BBXX:2:2224:17472:439 | 1 |
| Inter-6 | Control10 | TC10plus | ALU | 235 | Yes | Yes | AGATAAGCATAGCATAGCATGAT | HR-INTER-6-FW | ACAAATGAACAAAAGACGCACT | HR-INTER-6-REV | K00114:354:HC225BBXX:4:1110:31456:303 | 1 |
| Inter-7 | Control2 | LC2 | ALU | 229 | Yes | Yes | TGATTTACTGATGAAACCTACCA | HR-INTER-7-FW | AGGAGATGTGGTGTAGAGTTGA | HR-INTER-7-REV | K00114:354:HC225BBXX:1:2219:21805:745 | 1 |
| Inter-8 | Control2 | TC2plus | ALU | 194 | Yes | Yes | ACCCCTGATTTAGTGATCTCA | HR-INTER-8-FW | AATATGCCCTATTAAGATTCGCT | HR-INTER-8-REV | K00114:354:HC225BBXX:3:1214:21227:326 | 1 |
| Inter-9 | Control3 | PC3minus | ALU | 263 | No | / | GCCTGTAAAGCAACCAGTG | HR-INTER-9-FW | GGAAAGTTACACAGGGACATT | HR-INTER-9-REV | K00114:354:HC225BBXX:7:1209:15270:551 | 1 |
| Inter-10 | Control3 | KC3 | ALU | 224 | Yes | Yes | CACTACGAATTGGTATGTGGGT | HR-INTER-10-FW | TCCCTTAACAGCACTGCAGA | HR-INTER-10-REV | K00114:354:HC225BBXX:2:2204:28392:321 | 1 |
| Inter-11 | Control3 | TC3plus | ALU | 236 | Yes | Yes | ACCCACACATCCATCTATTGC | HR-INTER-11-FW | AGAGAAACCAGTGGAATAGAAGT | HR-INTER-11-REV | K00114:354:HC225BBXX:3:2105:10774:442 | 1 |
| Inter-12 | Control4 | LC4 | ALU | 253 | Yes | Yes | AAGCAGCCTAAATGTCTAACTT | HR-INTER-12-FW | CGAAGGCAGTATTATGGACAAG | HR-INTER-12-REV | K00114:354:HC225BBXX:1:2222:2280:3734 | 1 |
| Inter-13 | Control4 | TC4minus | ALU | 197 | Yes | Yes | ACATCTGGGCTGTGACTGTC | HR-INTER-13-FW | TCAGAGAATGGGCCCTACAG | HR-INTER-13-REV | K00114:354:HC225BBXX:5:2212:8349:2587 | 1 |
| Inter-14 | Control5 | FC5minus | ALU | 231 | Yes | Yes | TCCCATGGTAGAGCTTAGCA | HR-INTER-14-FW | TGAATAACAAGGCAATGGTGGG | HR-INTER-14-REV | K00114:354:HC225BBXX:3:1225:24830:488 | 1 |
| Inter-15 | Control5 | PC5plus | ALU | 251 | Yes | Yes | AAAACGCAGGGGAAGGCA | HR-INTER-15-FW | AGCAGGATTTCACTCGATTTATT | HR-INTER-15-REV | K00114:354:HC225BBXX:5:2209:15757:32 | 1 |
| Inter-16 | Control6 | FC6minus | ALU | 262 | Yes | Yes | CCTTCTCCTGTTAGTTGCCC | HR-INTER-16-FW | ACCCTTTCCCGTGTCCTT | HR-INTER-16-REV | K00114:354:HC225BBXX:4:2202:10389:192 | 1 |
| Inter-17 | Control6 | KC6 | ALU | 248 | No | / | CAGAAAGAGGAGAAAAGTTGC | HR-INTER-17-FW | CTCAAGTAATCCACCTGCCT | HR-INTER-17-REV | K00114:354:HC225BBXX:1:1121:9475:2206 | 1 |
| Inter-18 | Control10 | TC10plus | L1 | 216 | Yes | Yes | CATCTTGACGAGCTGTTTGGA | HR-INTER-18-FW | AGATTTCAGGCTGCATTAATTTG | HR-INTER-18-REV | K00114:354:HC225BBXX:4:1118:20324:120 | 1 |
| Inter-19 | Control2 | TC2plus | L1 | 253 | Yes | Yes | AGTGACCTTAGTTTTCAGTGTT | HR-INTER-19-FW | TGGGAATGCTGAAGGATAATAAA | HR-INTER-19-REV | K00114:354:HC225BBXX:3:2110:21471:205 | 1 |
| Inter-20 | Control4 | TC4minus | L1 | 191 | Yes | No | GGTTATTTACTACCCCTCCACT | HR-INTER-20-FW | TTGTGAAAGCCAACCAGACA | HR-INTER-20-REV | K00114:354:HC225BBXX:5:1204:22171:338 | 1 |
| Inter-21 | Control6 | TC6minus | L1 | 242 | Yes | Yes | ACTGGCCAGAGTGAGATTC | HR-INTER-21-FW | AGGACCCAGAAACAGAAGCT | HR-INTER-21-REV | K00114:354:HC225BBXX:6:1110:23997:221 | 1 |
| Inter-22 | Control7 | KC7 | L1 | 237 | Yes | Yes | TCAGGTCAACAAATTTACTCTGT | HR-INTER-22-FW | ACCCATAACCTGCTCCAG | HR-INTER-22-REV | K00114:354:HC225BBXX:1:1114:26900:479 | 2 |
| Inter-23 | PD10 | FP10minus | ALU | 154 | Yes | Yes | GGGGTAGGCAGATGGGTG | HR-INTER-23-FW | CAGTTTTGAATGTGGCATTCTCA | HR-INTER-23-REV | K00114:354:HC225BBXX:6:1210:13494:651 | 1 |
| Inter-24 | PD10 | LP10 | ALU | 130 | Yes | Yes | TTAGGAGTCCATTGCAAGATG | HR-INTER-24-FW | CTTAGGGAACAAGCACAGTCA | HR-INTER-24-REV | K00114:354:HC225BBXX:1:2101:7750:1369 | 1 |
| Inter-25 | PD10 | TP10plus | ALU | 218 | Yes | Yes | CAAAGCATGAAAGTGATGTCA | HR-INTER-25-FW | ACAGATGCCTCTTGAGAAGTG | HR-INTER-25-REV | K00114:354:HC225BBXX:6:2206:23033:658 | 1 |
| Inter-26 | PD2 | LP2 | ALU | 220 | Yes | Yes | AGCATCACTCTCATTCCTGGT | HR-INTER-26-FW | AGAGATGGGAAGGGAAAACA | HR-INTER-26-REV | K00114:354:HC225BBXX:2:2217:12378:264 | 1 |
| Inter-27 | PD3 | FP3plus | ALU | 237 | Yes | Yes | TCTCTAATGCCTCCTTCCCT | HR-INTER-27-FW | TGTGCATCTTAACTCTCTTCCA | HR-INTER-27-REV | K00114:354:HC225BBXX:3:1205:6197:4842 | 1 |
| Inter-28 | PD1 | FP1plus | L1 | 192 | Yes | Yes | AGCTGGAGTCCGTCATTAGA | HR-INTER-28-FW | ACTCATGTTCAAATATGGCAGTC | HR-INTER-28-REV | K00114:354:HC225BBXX:3:2223:30198:68 | 1 |
| Inter-29 | PD1 | PP1plus | L1 | 203 | Yes | Yes | AGGGCATTTTACTTGTGAGGT | HR-INTER-29-FW | TGCTTATTGTGTGTTAGCCA | HR-INTER-29-REV | K00114:354:HC225BBXX:7:1109:10916:247 | 1 |
| Inter-30 | PD10 | PP10plus | L1 | 211 | No | / | ACTTTCTACCAGTTTCTGCAC | HR-INTER-30-FW | TGGGAAATTTCAGGTCAAAGT | HR-INTER-30-REV | K00114:354:HC225BBXX:8:1102:28940:474 | 1 |
| Inter-31 | PD2 | FP2plus | L1 | 194 | Yes | Yes | TGAATTGAGGTTTTGCTACGT | HR-INTER-31-FW | TTTCTCCTGCGTCCTTTTCT | HR-INTER-31-REV | K00114:354:HC225BBXX:3:1109:14245:936 | 1 |
| Inter-32 | PD2 | KP2 | L1 | 159 | Yes | Yes | AGCAGTCTTGAAAATGAGCAT | HR-INTER-32-FW | AACTGAGTGGATGGATTGCA | HR-INTER-32-REV | K00114:354:HC225BBXX:1:2203:5081:4103 | 1 |
| Inter-33 | AD1 | TA1minus | ALU | 275 | Yes | Yes | TTCACCCACTGCGCTGAG | HR-INTER-33-FW | CCCATTTTACGTGAATCTGAAGC | HR-INTER-33-REV | K00114:354:HC225BBXX:3:1203:27113:831 | 1 |
| Inter-34 | AD1 | PA1minus | ALU | 225 | Yes | Yes | CCCCTAGCCAAATGACTGC | HR-INTER-34-FW | GGACAATAAGCACACCCTGG | HR-INTER-34-REV | K00114:354:HC225BBXX:5:1106:8065:4581 | 1 |
| Inter-35 | AD10 | FA10minus | ALU | 225 | Yes | Yes | CTGCAGGTCTCCAGGCTG | HR-INTER-35-FW | TTATCGGGTTAGCAGACACC | HR-INTER-35-REV | K00114:354:HC225BBXX:8:2116:30908:237 | 1 |
| Inter-36 | AD10 | KA10 | ALU | 156 | Yes | Yes | CCTTCCCCTACCCACATACC | HR-INTER-36-FW | AGGGACATGGGTATCTAGAAGG | HR-INTER-36-REV | K00114:354:HC225BBXX:1:1206:22262:335 | 1 |
| Inter-37 | AD2 | TA2minus | ALU | 220 | Yes | Yes | AGCCTAACTAAATGTCAGCAA | HR-INTER-37-FW | AGACAGGTCAGAGTAATGGAT | HR-INTER-37-REV | K00114:354:HC225BBXX:3:1225:5751:1747 | 1 |
| Inter-38 | AD1 | PA1plus | L1 | 190 | Yes | Yes | CCTGCGGCCTTAACTCTGT | HR-INTER-38-FW | GCTTGACGATATTCTCTTTTGTC | HR-INTER-38-REV | K00114:354:HC225BBXX:3:1121:20090:364 | 1 |
| Inter-39 | AD2 | LA2 | L1 | 245 | Yes | Yes | AGATTTCAAGGGCAATAGGATTG | HR-INTER-39-FW | GCAAAAGGCGTGACTTGATTG | HR-INTER-39-REV | K00114:354:HC225BBXX:1:1113:18010:466 | 1 |
| Inter-40 | AD3 | KA3 | L1 | 222 | Yes | Yes | AGGAAGACGTAGATTAGAATGGA | HR-INTER-40-FW | GGTGTGTTTCATAAAATGTGTGT | HR-INTER-40-REV | K00114:354:HC225BBXX:2:1119:5822:4253 | 1 |
| Inter-41 | AD3 | TA3plus | L1 | 234 | No | / | CAGTCTTTCAGGGTGTGGGA | HR-INTER-41-FW | TGGTGTCATGGAAGGCTTCT | HR-INTER-41-REV | K00114:354:HC225BBXX:7:2119:30868:149 | 1 |
| Inter-42 | AD5 | PA5plus | L1 | 236 | Yes | Yes | CTGCAGATACCTCCGGAGAC | HR-INTER-42-FW | ACCATGTGCTTCCCAGTCTG | HR-INTER-42-REV | K00114:354:HC225BBXX:3:2114:26890:655 | 1 |
| Inter-43 | AD6 | KA6 | L1 | 239 | Yes | Yes | GAATTACATTTGCAACCTCTACA | HR-HS1-FW-1 | CGGTATGGACTTTTCAGACA | HR-HS1-REV-1 | K00114:354:HC225BBXX:1:2111:31253:141 | 1 |
| Inter-44 | Control9 | KC9 | L1 | 159 | Yes | Yes | GGATTAAAAATCACTAAGCAGGG | HR-HS1-FW-2 | GGGGACTTTAATACATATGGCA | HR-HS1-REV-2 | K00114:354:HC225BBXX:1:2113:16051:201 | 2 |
| Inter-45 | Control5 | LC5 | L1 | 175 | No | / | GAATTACATTTGCAACCTCTACA | HR-HS1-FW-1 | TTCTGCTGCTGTAAAACTTACC | HR-HS1-REV-3 | K00114:354:HC225BBXX:1:2116:26768:328 | 1 |
| Inter-46 | AD9 | LA9 | L1 | 182 | Yes | Yes | GAATTACATTTGCAACCTCTATA | HR-HS1-FW-4 | TGGAACCTTACTTTGGATTGAGC | HR-HS1-REV-4 | K00114:354:HC225BBXX:2:1121:20730:509 | 1 |
| Inter-47 | Control7 | LC7 | L1 | 207 | Yes | Yes | GGATTAAAAATCACTAAGCAGGG | HR-HS1-FW-2 | TGCAGCCTACCTACATCTTGAT | HR-HS1-REV-5 | K00114:354:HC225BBXX:2:2102:14123:329 | 2 |
| Inter-48 | Control9 | TP9minus | L1 | 215 | Yes | Yes | GGATTAAAAATCACTAAGCAGGG | HR-HS1-FW-2 | CACTCTGTGGGCAAGGACT | HR-HS1-REV-6 | K00114:354:HC225BBXX:8:1125:20344:415 | 1 |
| Inter-49 | Control10 | KP10 | L1 | 163 | Yes | Yes | CCAAATTCTCTGAATTTGGAAAAG | HR-HS2-FW-1 | ATCCGTGATTCCCCTTCAGA | HR-HS2-REV-1 | K00114:354:HC225BBXX:2:1102:12469:161 | 1 |
| Inter-50 | AD3 | PA3plus | L1 | 175 | Yes | Yes | GTGCATGCCAAATTCTCTGA | HR-HS2-FW-2 | ACTGGTTGCCACACTTGATG | HR-HS2-REV-2 | K00114:354:HC225BBXX:3:1209:10947:130 | 1 |
| Inter-51 | PD7 | FP7minus | L1 | 98 | Yes | Yes | GGATTAAAAATCACTAAGCAGGG | HR-HS1-FW-2 | GTGACCAGAGAAACGGGAGT | HR-HS2-REV-3 | K00114:354:HC225BBXX:6:2215:20273:160 | 1 |
| Inter-52 | Control2 | FC2plus | L1 | 166 | Yes | Yes | GGATTAAAAATCACTAAGCAGGG | HR-HS1-FW-2 | ATCCTTCCACCCATCCCTAT | HR-HS2-REV-4 | K00114:354:HC225BBXX:7:2125:10866:498 | 1 |
| Inter-53 | Control6 | PC6plus | L1 | 165 | Yes | Yes | GGATTAAAAATCACTAAGCAGGG | HR-HS1-FW-2 | TGAATTTTCCAGCAATAGCTCA | HR-HS2-REV-5 | K00114:354:HC225BBXX:6:2227:21765:271 | 1 |
| Inter-54 | Control7 | KC7 | L1 | 243 | Yes | Yes | GCATTTGCATGTTCACAAACT | HR-HS3-FW-1 | TCCATCTTACCAAATCCAAAGA | HR-HS3-REV-1 | K00114:354:HC225BBXX:1:1215:12083:345 | 1 |
| Inter-55 | AD3 | KA3 | L1 | 135 | Yes | Yes | TGATGAAATGAAAGATAAGAAAAGC | HR-HS3-FW-2 | CTGCTAGGTACCTCCAGG | HR-HS3-REV-2 | K00114:354:HC225BBXX:2:1107:26362:200 | 2 |
| Inter-56 | PD2 | LP2 | L1 | 138 | Yes | Yes | TGATGAAATGAAAGATAAGAAAAGC | HR-HS3-FW-2 | TGAGTATCTTAGTCTCACTTTATC | HR-HS3-REV-3 | K00114:354:HC225BBXX:2:2121:24759:184 | 1 |
| Inter-57 | PD4 | FP4plus | L1 | 141 | Yes | Yes | TGATGAAATGAAAGATAAGAAAAGC | HR-HS3-FW-2 | TAGACGAGTAAGCTGGTCATTTTT | HR-HS3-REV-4 | K00114:354:HC225BBXX:3:2217:4259:1548 | 2 |
| Inter-58 | PD4 | FP4minus | L1 | 201 | Yes | Yes | TGATGAAATGAAAGATAAGAAAAGC | HR-HS3-FW-2 | CGTCTTTTACTCCTGTCATTAAGC | HR-HS3-REV-5 | K00114:354:HC225BBXX:5:1124:17817:473 | 2 |
| Inter-59 | Control2 | LC2 | L1 | 213 | Yes | Yes | CTGTATCTCAGATACATTTGAGG | HR-HS4-FW-1 | TCATTAATGATGCAAATGTAATGTT | HR-HS4-REV-1 | K00114:354:HC225BBXX:1:2104:29264:388 | 1 |
| Inter-60 | PD3 | FP3plus | L1 | 160 | Yes | Yes | CTGTATCTCAGATACATTTGAGG | HR-HS4-FW-1 | TAAATATTCATGGGATGCATGTG | HR-HS4-REV-2 | K00114:354:HC225BBXX:3:1128:13250:279 | 2 |
| Inter-61 | AD8 | TA8minus | L1 | 228 | Yes | Yes | CTGTATCTCAGATACATTTGAGG | HR-HS4-FW-1 | CCTCACATTTTCTTATCATTTCTT | HR-HS4-REV-3 | K00114:354:HC225BBXX:4:1208:30188:249 | 1 |
| Inter-62 | PD3 | PP3plus | L1 | 215 | Yes | Yes | CTGTATCTCAGATACATTTGAGG | HR-HS4-FW-1 | ACTCTTTTGTAAGTACATTGCATA | HR-HS4-REV-4 | K00114:354:HC225BBXX:7:2225:12286:236 | 2 |
| Inter-63 | PD1 | LP1 | L1 | 204 | Yes | Yes | TCTAGCCTTTCTAGAAATAAGG | HR-HS5-FW-1 | AACTAATTCATGCAGGGCTTTT | HR-HS5-REV-1 | K00114:354:HC225BBXX:2:1205:16812:349 | 1 |
| Inter-64 | PD8 | FP8minus | L1 | 197 | No | / | TCTAGCCTTTCTAGAAATAAGG | HR-HS5-FW-1 | TGCATGTAACTCAAAACCAGATT | HR-HS5-REV-2 | K00114:354:HC225BBXX:6:1106:3539:1040 | 4 |
| Inter-65 | Control10 | PC10minus | L1 | 242 | Yes | Yes | TAGCCTTTCTAGAAATAAGGCTA | HR-HS5-FW-3 | AATTTGTTCTTCCCATGCTCA | HR-HS5-REV-3 | K00114:354:HC225BBXX:8:1123:15625:443 | 1 |
| Inter-66 | PD10 | PP10plus | L1 | 224 | No | / | TCTAGCCTTTCTAGAAATAAGG | HR-HS5-FW-1 | GTGCAGACCTAACTCAGCAAA | HR-HS5-REV-4 | K00114:354:HC225BBXX:8:1211:5842:4189 | 1 |
| Inter-67 | Control7 | KC7 | ALU | 194 | Yes | Yes | AGGTGTACAATCACTGCCAGA | HR-HS6-FW-1 | GGCCTGGAAGGAGATGTTTA | HR-HS6-REV-1 | K00114:354:HC225BBXX:1:1123:20375:325 | 1 |
| Inter-68 | AD1 | KA1 | ALU | 228 | Yes | No | AGTGTCGGGAGGCTGAGG | HR-HS6-FW-2 | GAGGATGAGTCACTTAGCTATC | HR-HS6-REV-2 | K00114:354:HC225BBXX:2:1216:25063:232 | 1 |
| Inter-69 | AD5 | PA5plus | ALU | 188 | Yes | Yes | AGGTGTACAATCACTGCCAGA | HR-HS6-FW-1 | CCTCTTTGGAAAGTTTGCACT | HR-HS6-REV-3 | K00114:354:HC225BBXX:3:1113:2930:1793 | 1 |
| Inter-70 | PD6 | LP6 | ALU | 176 | Yes | Yes | TTGTTTAAGAGTGAAGAAGAACTT | HR-HS7-FW-1 | TATTTGTACATTTGTAAAATCTGC | HR-HS7-REV-1 | K00114:354:HC225BBXX:1:2101:12297:121 | 1 |
| Inter-71 | PD8 | FP8minus | ALU | 245 | Yes | Yes | GGACTCAGGTCTTTGGTGTCA | HR-HS7-FW-2 | GAGTCATTAAACATATCGAAGAAA | HR-HS7-REV-2 | K00114:354:HC225BBXX:6:1128:28564:144 | 2 |
| Inter-72 | AD9 | FA9plus | ALU | 207 | Yes | Yes | TTGTTTAAGAGTGAAGAAGAACTT | HR-HS7-FW-1 | TGCCGGCCATACGGTTTTTC | HR-HS7-REV-3 | K00114:354:HC225BBXX:6:2204:15148:112 | 1 |
| Inter-73 | Control7 | PC7minus | ALU | 212 | Yes | Yes | TGGTTGTTGTTTAAGAGTGAAGAA | HR-HS7-FW-4 | TCAGAATTCAGAAACCTTTTGG | HR-HS7-REV-4 | K00114:354:HC225BBXX:8:2108:5558:3307 | 1 |
| Inter-74 | Control7 | TC7plus | ALU | 202 | Yes | No | CCCTTAAGAAAAGACCGTGG | HR-HS8-FW-1 | AGGTTTTCACAGTAAGGAAGCAC | HR-HS8-REV-1 | K00114:354:HC225BBXX:4:2207:4817:4599 | 1 |
| Inter-75 | Control9 | PC9plus | ALU | 199 | Yes | Yes | CCCTTAAGAAAAGACCGTGG | HR-HS8-FW-1 | TTTTGAATACAGAAAAAGCCTTT | HR-HS8-REV-2 | K00114:354:HC225BBXX:6:2110:18974:613 | 1 |
| Inter-76 | AD9 | FA9minus | ALU | 262 | Yes | Yes | AGGGAGACATCTTGCCCTTA | HR-HS8-FW-3 | ACCTCGCCCAGCTCCTAT | HR-HS8-REV-3 | K00114:354:HC225BBXX:8:2128:17107:385 | 1 |
| Inter-77 | PD9 | LP9 | ALU | 178 | No | / | CAATGTCCCCACACTGAATAC | HR-HS9-FW-1 | TATGGGCCAGGCCCTTCTTT | HR-HS9-REV-1 | K00114:354:HC225BBXX:1:2122:5893:1110 | 1 |
| Inter-78 | Control5 | KC5 | ALU | 190 | Yes | No | CAATGTCCCCACACTGAATAC | HR-HS9-FW-1 | ATTCAACCCCGTCCAGAGAT | HR-HS9-REV-2 | K00114:354:HC225BBXX:2:2101:14884:436 | 1 |
| Inter-79 | Control10 | FC10plus | ALU | 173 | Yes | Yes | CCCTGAATACATGTTATAAAAATTAGC | HR-HS9-FW-3 | CCCAGCCTTCATTTCCTTCA | HR-HS9-REV-3 | K00114:354:HC225BBXX:8:2107:25905:303 | 1 |
| Intra-1 | AD3 | PA3minus | ALU | 183 | Yes | Yes | GCAACAGATAGCTTTCGTTT | HR-INTRA-1-FW | TTGACCTTGCTTTGTGAAAA | HR-INTRA-1-REV | K00114:354:HC225BBXX:5:2203:8471:3569 | 1 |
| Intra-2 | AD6 | PA6minus | ALU | 239 | Yes | Yes | CAAACTTGGGTCCACTAGTAGT | HR-INTRA-2-FW | AGTCTTCTTGTCCACCAGCT | HR-INTRA-2-REV | K00114:354:HC225BBXX:6:1123:24271:302 | 1 |
| Intra-3 | AD8 | LA8 | ALU | 229 | Yes | Yes | CCATGGTGCTGCTCAGAGTA | HR-INTRA-3-FW | TGTGAAGGTCCTGATGCCAT | HR-INTRA-3-REV | K00114:354:HC225BBXX:2:2219:13910:469 | 1 |
| Intra-4 | Control1 | PC1minus | ALU | 221 | Yes | Yes | AGTTTTATGTAATTTCCTGGAGG | HR-INTRA-4-FW | TGATCTCACATGGGCATAGT | HR-INTRA-4-REV | K00114:354:HC225BBXX:7:1108:9942:4246 | 1 |
| Intra-5 | Control2 | LC2 | L1 | 151 | Yes | Yes | GCTCTTGAAATAAAATGGTATCTCT | HR-INTRA-5-FW | CATTTTGCACAACTCCCTGC | HR-INTRA-5-REV | K00114:354:HC225BBXX:1:1113:13991:368 | 1 |
| Intra-6 | Control5 | LC5 | L1 | 242 | Yes | Yes | GGCTTCCATGTCTTGAGTTAA | HR-INTRA-6-FW | CACCAAAACCCACCACCAG | HR-INTRA-6-REV | K00114:354:HC225BBXX:1:1101:7436:1874 | 1 |
| Intra-7 | PD1 | LP1 | L1 | 251 | Yes | Yes | ACAAATTGTATGCTCCAGGTCA | HR-INTRA-7-FW | TTTCTGCTTGGGTCCTCTGT | HR-INTRA-7-REV | K00114:354:HC225BBXX:2:2223:12560:293 | 1 |
| Intra-8 | Control7 | FC7minus | ALU | 228 | Yes | Yes | CCAGTCAGGCCATGTTCT | HR-INTRA-8-FW | AACAATGCCTCCAGATCTAA | HR-INTRA-8-REV | K00114:354:HC225BBXX:4:1225:8085:4061 | 1 |
| Intra-9 | Control1 | PC1plus | ALU | 174 | Yes | No | ACCACATGACAATTACAGTATCC | HR-INTRA-9-FW | AGCCTGATTAAAAGAGAGCCCT | HR-INTRA-9-REV | K00114:354:HC225BBXX:5:2125:14448:378 | 1 |
| Intra-10 | Control1 | FC10plus | ALU | 204 | Yes | No | ACACTAATCAACTCAGTAACAATG | HR-INTRA-10-FW | TGGTTAGGGTCAATCAGTGGT | HR-INTRA-10-REV | K00114:354:HC225BBXX:8:2109:4503:2160 | 1 |
| Intra-11 | Control2 | PC2minus | ALU | 207 | No | / | TGTTCTACTGACATCTGGACTT | HR-INTRA-11-FW | AGGGAGATAACTGTGACCGG | HR-INTRA-11-REV | K00114:354:HC225BBXX:7:1222:11992:687 | 2 |
| Intra-12 | Control6 | LC6 | ALU | 242 | Yes | Yes | AATTTGGGCGACACACAAGG | HR-INTRA-12-FW | GGACCACAGCCTAAGAACCA | HR-INTRA-12-REV | K00114:354:HC225BBXX:2:2210:29701:343 | 1 |
| Intra-13 | Control7 | FC7minus | ALU | 244 | Yes | Yes | CCAGTCAGGCCATGTTCT | HR-INTRA-13-FW | GTGGTTAATAACTTATAACAATGCC | HR-INTRA-13-REV | K00114:354:HC225BBXX:4:1225:8085:4061 | 1 |
| Intra-14 | Control7 | PC7minus | ALU | 232 | Yes | Yes | AGACTAGTAAACAGAAGGTGGCT | HR-INTRA-14-FW | CAGGCATACTCAAGCTTTCT | HR-INTRA-14-REV | K00114:354:HC225BBXX:8:2202:20222:314 | 1 |
| Intra-15 | Control7 | TC7plus | ALU | 159 | Yes | Yes | AACAGGCTGAGAAAAGTGGC | HR-INTRA-15-FW | TCACAAACCTGGGCTAAAG | HR-INTRA-15-REV | K00114:354:HC225BBXX:4:1128:17005:413 | 1 |
| Intra-16 | Control1 | TC1plus | L1 | 185 | Yes | Yes | GGTAACAGTAATCGGATTTGCC | HR-INTRA-16-FW | AGTGGGAAATCGATAACATGGA | HR-INTRA-16-REV | K00114:354:HC225BBXX:3:2220:1925:2603 | 1 |
| Intra-17 | Control4 | PC4plus | L1 | 240 | Yes | Yes | TTAGAACTTCAGGCTGCTGA | HR-INTRA-17-FW | GGTAGGAGAAACTAATATGGTGAC | HR-INTRA-17-REV | K00114:354:HC225BBXX:5:1107:6837:1896 | 1 |
| Intra-18 | Control5 | KC5 | L1 | 207 | Yes | Yes | TGAGCAATATAGTTGACCCTT | HR-INTRA-18-FW | TCCAGGTTGACACAAAGATG | HR-INTRA-18-REV | K00114:354:HC225BBXX:2:2218:20638:318 | 1 |
| Intra-19 | Control6 | LC6 | L1 | 172 | Yes | Yes | AGGAGGACATGAGATAACATC | HR-INTRA-19-FW | TCAGCTTCATACCCAGATGT | HR-INTRA-19-REV | K00114:354:HC225BBXX:2:2114:14894:32 | 1 |
| Intra-20 | PD2 | LP2 | ALU | 216 | Yes | Yes | ACAACCAGACAACAAGTAGCTG | HR-INTRA-20-FW | CCTCCACTGTCCCTTGCAT | HR-INTRA-20-REV | K00114:354:HC225BBXX:2:2128:7689:3862 | 1 |
| Intra-21 | PD3 | PP3minus | ALU | 251 | Yes | Yes | TGTCCTGCACACAATCCTAGT | HR-INTRA-21-FW | TGTGCCTAGGAGAGAAGCTG | HR-INTRA-21-REV | K00114:354:HC225BBXX:3:1123:18365:35 | 1 |
| Intra-22 | PD4 | PP4plus | ALU | 202 | Yes | Yes | TTCTATTGCTTAGCCCCAGG | HR-INTRA-22-FW | GTTGAGAGAAGGGCCATGGT | HR-INTRA-22-REV | K00114:354:HC225BBXX:7:1204:8948:3718 | 1 |
| Intra-23 | PD6 | TP6minus | ALU | 214 | Yes | Yes | AGATGTGATAGCTTGCCAGGA | HR-INTRA-23-FW | TCTCAAGTGGTCGCCTCTG | HR-INTRA-23-REV | K00114:354:HC225BBXX:8:1117:21531:127 | 1 |
| Intra-24 | PD10 | KP10 | L1 | 189 | Yes | Yes | CCCTCCAGAGAAGGTTTGCT | HR-INTRA-24-FW | GCACTTTCCACAAGATACATGTC | HR-INTRA-24-REV | K00114:354:HC225BBXX:2:1105:20872:268 | 1 |
| Intra-25 | PD3 | KP3 | L1 | 228 | Yes | Yes | AAAGTCATGGATGTCTGGTAC | HR-INTRA-25-FW | CACTTTGTGCCAGACCCAC | HR-INTRA-25-REV | K00114:354:HC225BBXX:1:2114:8511:2661 | 1 |
| Intra-26 | PD8 | PP8minus | L1 | 236 | Yes | Yes | AAGTACCAAGAAAGGAGCCA | HR-INTRA-26-FW | GTGTATCTGATTCATCCCATTTC | HR-INTRA-26-REV | K00114:354:HC225BBXX:4:2117:30360:384 | 1 |
| Intra-27 | AD1 | LA1 | L1 | 236 | Yes | No | AGACGGTGCATTCTCCATTG | HR-INTRA-27-FW | GAAGCGTTAGAACACCACCA | HR-INTRA-27-REV | K00114:354:HC225BBXX:1:1223:20659:419 | 1 |
| Intra-28 | AD5 | TA5minus | L1 | 164 | Yes | Yes | CTTGATGACTTTAATGTAACAGCA | HR-INTRA-28-FW | TTCCTCTCTGCATTTGGGC | HR-INTRA-28-REV | K00114:354:HC225BBXX:3:2124:5923:3222 | 1 |
| Intra-29 | AD6 | PA6plus | L1 | 225 | Yes | Yes | AGCATCTTACACAATCATTAGCA | HR-INTRA-29-FW | TGTAAGTGTAGAAGTCCAAGAGG | HR-INTRA-29-REV | K00114:354:HC225BBXX:4:2217:7862:1983 | 1 |
| Intra-30 | AD8 | PA8minus | ALU | 204 | Yes | Yes | GCTCCCTGATCACCTCTTCA | HR-INTRA-30-FW | GCGAGACGTGCCTATGTG | HR-INTRA-30-REV | K00114:354:HC225BBXX:6:1101:13890:280 | 1 |
| Intra-31 | AD8 | TA8minus | ALU | 209 | Yes | No | AACTGTTGGGGTTGAAATGG | HR-HS10-FW-1 | CCTGCATTTGAATGCTTCTG | HR-HS10-REV-1 | K00114:354:HC225BBXX:4:2101:22972:466 | 1 |
| Intra-32 | AD4 | TA4plus | ALU | 200 | Yes | No | AACTGTTGGGGTTGAAATGG | HR-HS10-FW-1 | GAATGCTTCTGAACACATCTTC | HR-HS10-REV-2 | K00114:354:HC225BBXX:7:2107:15300:361 | 1 |
| Intra-33 | AD6 | TA6plus | ALU | 185 | Yes | No | ATGGGGGTGGGTGTCTAAAG | HR-HS10-FW-3 | CCTGCATTTGAATGCTTCTG | HR-HS10-REV-1 | K00114:354:HC225BBXX:8:1126:6370:1175 | 1 |

**PCR validation of putative somatic NAHR events in capture-seq libraries - PCR from GENOMIC DNA**

| Target Name | Donor | Library | Repeat | Exp. Amplicon size | Exp. Amplicon size N | Nested PCR Band of exp | Validated by Sanger | FW primer sequence | REV primer sequence | Nested FW primer sequence | Nested REV primer sequence | Contig ID | Supporting Contigs |
|---|---|---|---|---|---|---|---|---|---|---|---|---|---|
| Inter-1 | AD1 | FA1plus | ALU | 262 | 191 | | | CATTAAAGTGGGCCCAGTG | TTCTGACAGTTGCTTCCCA | CATCAACAAAGAAGGGCCA | TCCCAGCATGGATGTAAAGA | K00114:354:HC225BBXX:5:2106:26 | 1 |
| Inter-2 | AD1 | PA1minus | ALU | 228 | 204 | Yes | | CCAAATGACTGCTCTCCCTG | CGCAGTAGGTGGACAATAAGC | GCTCTCCCTGTCTCTCATG | GCACACCCTGGTTAAAGAAG | K00114:354:HC225BBXX:5:1106:80 | 1 |
| Inter-3 | Control1 | FC1minus | ALU | 245 | 183 | Yes | Yes | TGAAACCAGATATTTGAAACTTACC | CGACCCTCCCATCTCAGC | TGAAACTTACCTTCTTCTTAACGTCT | CCACCACACTCAGACTTTC | K00114:354:HC225BBXX:3:2214:16 | 1 |
| Inter-4 | Control1 | FC1plus | ALU | 221 | 197 | | | AAACTATGAGGAGCAGGCTT | GGTGGTCAGCAGCTCTAAAC | GAGGAGCAGGCTTGGATAA | ACCTAAGACCCCAACTGTG | K00114:354:HC225BBXX:7:1222:15 | 1 |
| Inter-5 | Control10 | LC10 | ALU | 149 | / | | | GCAATACTAAGAGAAAGGATGGA | GGCAAGAATCCTGACCCA | AAGAGAAAGGATGGAGGCC | CCACATCTGTATGCTAAGCC | K00114:354:HC225BBXX:2:2224:17 | 1 |
| Inter-6 | Control10 | TC10plus | ALU | 235 | / | | | AGATAAGCATAGCATAGCATGAT | ACAAATGAACAAAAGACGCACT | GAAGTCAGATAGAACTGAGCTCC | CACTTTAATAATTCTCCTGCCCG | K00114:354:HC225BBXX:4:1110:31 | 1 |
| Inter-7 | Control2 | LC2 | ALU | 229 | 164 | | | TGATTTACTGATGAAACCTACCA | AGGAGATGTGGTGTAGAGTTGA | GAGGATGGCAAATATTTTCCTACTAC | TTTCATCCCCACCTCTTGAT | K00114:354:HC225BBXX:1:2219:21 | 1 |
| Inter-8 | Control2 | TC2plus | ALU | 194 | 160 | | | ACCCCTGATTTAGTGATCTCA | AATATGCCCTATTAAGATTCGCT | GATCTCACTTAGAAACACCTCTCT | TCGCTGCAGAATATATCCCC | K00114:354:HC225BBXX:3:1214:21 | 1 |
| Inter-9 | Control3 | PC3minus | ALU | 263 | / | | | GCCTGTAAAGCAACCAGTG | GGAAAGTTACACAGGGACATT | GCAACCAGTGAGAATTTGCTTAGA | CCTTTAATGATAGATGCGGTGACTT | K00114:354:HC225BBXX:7:1209:15 | 1 |
| Inter-10 | Control3 | KC3 | ALU | 224 | 167 | | | CACTACGAATTGGTATGTGGGT | TCCCTTAACAGCACTGCAGA | TTTGAAAGTTAAGCCAGGCATG | TGCAGATAACTCCAAGTGGG | K00114:354:HC225BBXX:2:2204:28 | 1 |
| Inter-11 | Control3 | TC3plus | ALU | 236 | 216 | Yes | | ACCCACACATCCATCTATTGC | AGAGAAACCAGTGGAATAGAAGT | CCATCTATTGCTCCCAGGT | TGGAATAGAAGTAGCCGTTTCG | K00114:354:HC225BBXX:3:2105:10 | 1 |
| Inter-12 | Control4 | LC4 | ALU | 253 | 210 | | | AAGCAGCCTAAATGTCTAACTT | CGAAGGCAGTATTATGGACAAG | AGGAACCAGCTAAACAAACCT | CAAGGGATATTGGTAAGGTGTCT | K00114:354:HC225BBXX:1:2222:22 | 1 |
| Inter-13 | Control4 | TC4minus | ALU | 197 | 183 | Yes | | ACATCTGGGCTGTGACTGTC | TCAGAGAATGGGCCCTACAG | GCTGTGACTGTCTTCCAAAAG | AATGGGCCCTACAGAGAAG | K00114:354:HC225BBXX:5:2212:83 | 1 |
| Inter-14 | Control5 | FC5minus | ALU | 231 | 177 | | | TCCCATGGTAGAGCTTAGCA | TGAATAACAAGGCAATGGTGGG | CTAATGAAACAGACTGGGCG | AGGAATGGAGTGGGGTTAC | K00114:354:HC225BBXX:3:1225:24 | 1 |
| Inter-15 | Control5 | PC5plus | ALU | 251 | 192 | Yes | | AAAACGCAGGGGAAGGCA | AGCAGGATTTCACTCGATTTATT | CAAAGGCCTCTAAAAACGCA | AGCAGGATTTCACTCGATTTATTTTG | K00114:354:HC225BBXX:5:2209:1 | 1 |
| Inter-16 | Control6 | FC6minus | ALU | 262 | 244 | | | CCTTCTCCTGTTAGTTGCCC | ACCCTTTCCCGTGTCCTT | AGTTGCCCTAATTTCTTATGACAGT | CCCGTGTCCTTCAAAGTCT | K00114:354:HC225BBXX:4:2202:10 | 1 |
| Inter-17 | Control6 | KC6 | ALU | 248 | / | | | CCCTCAACTGAAATAGCCTTC | CTCAAGTAATCCACCTGCCT | CATTCAGAAAGAGGAGAAAAGTTGC | CAAAGTGCTGGGATTACAGG | K00114:354:HC225BBXX:1:1121:94 | 1 |
| Inter-18 | Control10 | TC10plus | L1 | 216 | 205 | Yes | | CATCTTGACGAGCTGTTTGGA | AGATTTCAGGCTGCATTAATTTG | TGACGAGCTGTTTGGAATTTC | CAGGCTGCATTAATTTGATCTGT | K00114:354:HC225BBXX:4:1118:20 | 1 |
| Inter-19 | Control2 | TC2plus | L1 | 253 | 187 | | | AGTGACCTTAGTTTTCAGTGTT | TGGGAATGCTGAAGGATAATAAA | AGTCAGAGGAAGTGTGTTGA | CTGAAGGATAATAAAAGACTGGTTTGTG | K00114:354:HC225BBXX:3:2110:21 | 1 |
| Inter-20 | Control4 | TC4minus | L1 | 191 | 183 | | | GGTTATTTACTACCCCTCCACT | TTGTGAAAGCCAACCAGACA | TACCCCTCCACTATGTCCA | CCAACCAGACATTTCATAGATATACCT | K00114:354:HC225BBXX:5:1204:22 | 1 |
| Inter-21 | Control6 | TC6minus | L1 | 242 | 148 | | | ACTGGCCAGAGTGAGATTC | AGGACCCAGAAACAGAAGCT | GATGTGTCTTGCCTTCTGG | GCAACTCCATCATTTTGACTCA | K00114:354:HC225BBXX:6:1110:23 | 1 |
| Inter-22 | Control7 | KC7 | L1 | 237 | 190 | | | TCAGGTCAACAAATTTACTCTGT | ACCCATAACCTGCTCCAG | GATCTCATTTATATTGGCTTTGGCG | TGCTCCAGCCAGTACTAC | K00114:354:HC225BBXX:1:1114:26 | 2 |
| Inter-23 | PD10 | FP10minus | ALU | 154 | 138 | | | GGGGTAGGCAGATGGGTG | CAGTTTTGAATGTGGCATTCTCA | AGATGGGTGAGCTAGTGGT | TGTGGCATTCTCATTAGGGA | K00114:354:HC225BBXX:6:1210:13 | 1 |
| Inter-24 | PD10 | LP10 | ALU | 130 | / | | | TTAGGAGTCCATTGCAAGATG | CTTAGGGAACAAGCACAGTCA | CCATTGCAAGATGGCTTGG | CTTAGGGAACAAGCACAGTCA | K00114:354:HC225BBXX:1:2101:77 | 1 |
| Inter-25 | PD10 | TP10plus | ALU | 218 | 153 | | | CAAAGCATGAAAGTGATGTCA | ACAGATGCCTCTTGAGAAGTG | GAGTGTATTCATGGCACACC | TAGAAATGGCTTCAATTGTGGCT | K00114:354:HC225BBXX:6:2206:23 | 1 |
| Inter-26 | PD2 | LP2 | ALU | 220 | 186 | | | AGCATCACTCTCATTCCTGGT | AGAGATGGGAAGGGAAAACA | CATTCCTGGTCTCCCTCC | AGAATGATGGGGAGGCAG | K00114:354:HC225BBXX:2:2217:12 | 1 |
| Inter-27 | PD3 | FP3plus | ALU | 237 | 216 | | | TCTCTAATGCCTCCTTCCCT | TGTGCATCTTAACTCTCTTCCA | CCTCCTTCCCTTCACAGTA | TCTCTTCCATACAAAGCATTCCT | K00114:354:HC225BBXX:3:1205:61 | 1 |
| Inter-28 | PD1 | FP1plus | L1 | 192 | 152 | | | AGCTGGAGTCCGTCATTAGA | ACTCATGTTCAAATATGGCAGTC | CCACTCACCCCATTGTTCT | CAGTCTCACAGTGTGGTTTTTC | K00114:354:HC225BBXX:3:2223:3 | 1 |
| Inter-29 | PD1 | PP1plus | L1 | 203 | 147 | | | AGGGCATTTTACTTGTGAGGT | TGCTTATTGTGTGTTAGCCA | GCATTTTACTTGTGAGGTTGGT | GTGTTAGCCAATGGGAGTTAC | K00114:354:HC225BBXX:7:1109:10 | 1 |
| Inter-30 | PD10 | PP10plus | L1 | 211 | 193 | | | ACTTTCTACCAGTTTCTGCAC | TGGGAAATTTCAGGTCAAAGT | AGTTTCTGCACTGAGGGG | TTCAGGTCAAAGTGGCATTG | K00114:354:HC225BBXX:8:1102:28 | 1 |
| Inter-31 | PD2 | FP2plus | L1 | 194 | / | | | TGAATTGAGGTTTTGCTACGT | TTTCTCCTGCGTCCTTTTCT | CATTTAGGTATCTGGGTGCAG | CCTTTTCTAACTTCATGCACTCAC | K00114:354:HC225BBXX:3:1109:14 | 1 |
| Inter-32 | PD2 | KP2 | L1 | 159 | 137 | | | AGCAGTCTTGAAAATGAGCAT | AACTGAGTGGATGGATTGCA | GAAAATGAGCATTATAGCACACCA | ATTGCACATAGTTCAAAATGATGACTG | K00114:354:HC225BBXX:1:2203:50 | 1 |
| Inter-33 | AD1 | TA1minus | ALU | 275 | 218 | | | TTCACCCACTGCGCTGAG | CCCATTTTACGTGAATCTGAAGC | AGTGAGGAAGATGCAGTCC | AAAAAAGCTGGCTGGGC | K00114:354:HC225BBXX:3:1203:27 | 1 |
| Inter-34 | AD1 | PA1minus | ALU | 225 | 210 | | | CCCCTAGCCAAATGACTGC | GGACAATAAGCACACCCTGG | CAAATGACTGCTCTCCCTG | AGCACACCCTGGTTAAAGA | K00114:354:HC225BBXX:5:1106:80 | 1 |
| Inter-35 | AD10 | FA10minus | ALU | 225 | 201 | | | CTGCAGGTCTCCAGGCTG | TTATCGGGTTAGCAGACACC | CTGACTCAGATTTCAAATGGGG | ACACCCTGTTAGTTGTTCTATCC | K00114:354:HC225BBXX:8:2116:30 | 1 |
| Inter-36 | AD10 | KA10 | ALU | 156 | 122 | | | CCTTCCCCTACCCACATACC | AGGGACATGGGTATCTAGAAGG | CCTACCTATCATTGGTCCTCTTG | GAAGGATTAAAAATGTTCTCCCCTC | K00114:354:HC225BBXX:1:1206:22 | 1 |
| Inter-37 | AD2 | TA2minus | ALU | 220 | 178 | | | AGCCTAACTAAATGTCAGCAA | AGACAGGTCAGAGTAATGGAT | TTAAGATCTAAGTATATTGGCTGGGTG | TATATTACCTCCCACATGCGC | K00114:354:HC225BBXX:3:1225:57 | 1 |
| Inter-38 | AD1 | PA1plus | L1 | 190 | 142 | | | CCTGCGGCCTTAACTCTGT | GCTTGACGATATTCTCTTTTGTC | CTGTTCCCATGGGTGCAG | GCTTGACGATATTCTCTTTTGTCTT | K00114:354:HC225BBXX:3:1121:20 | 1 |
| Inter-39 | AD2 | LA2 | L1 | 245 | 213 | | | AGATTTCAAGGGCAATAGGATTG | GCAAAAGGCGTGACTTGATTG | GGGCAATAGGATTGTGTTGTTAA | CCACACCATCATTACGTCCT | K00114:354:HC225BBXX:1:1113:18 | 1 |
| Inter-40 | AD3 | KA3 | L1 | 222 | 215 | Yes | | AGGAAGACGTAGATTAGAATGGA | GGTGTGTTTCATAAAATGTGTGT | AGGAAGACGTAGATTAGAATGGAATT | TCATAAAATGTGTGTCAAGGGTAGG | K00114:354:HC225BBXX:2:1119:58 | 1 |
| Inter-41 | AD3 | TA3plus | L1 | 234 | 212 | | | CAGTCTTTCAGGGTGTGGGA | TGGTGTCATGGAAGGCTTCT | GGGTGTGGGAAGGGTTT | GGCTTCTTGGAGGAAAGC | K00114:354:HC225BBXX:7:2119:30 | 1 |
| Inter-42 | AD5 | PA5plus | L1 | 236 | 183 | Yes | | CTGCAGATACCTCCGGAGAC | ACCATGTGCTTCCCAGTCTG | GCCTCCGTTTGAGAACTG | AGTCTGGGAGATCAAACTATGATATTC | K00114:354:HC225BBXX:3:2114:26 | 1 |
| Intra-1 | AD3 | PA3minus | ALU | 183 | 163 | | | GCAACAGATAGCTTTCGTTT | TTGACCTTGCTTTGTGAAAA | ATAGCTTTCGTTTCACATAGCTGT | GTGAAAAAGTGCAAAAATAAGACATGA | K00114:354:HC225BBXX:5:2203:84 | 1 |
| Intra-2 | AD6 | PA6minus | ALU | 239 | 225 | | | CAAACTTGGGTCCACTAGTAGT | AGTCTTCTTGTCCACCAGCT | TAGTAGTGCCCATTTGGTCAC | AGTCTTCTTGTCCACCAGCT | K00114:354:HC225BBXX:6:1123:24 | 1 |
| Intra-3 | AD8 | LA8 | ALU | 229 | 192 | | | CCATGGTGCTGCTCAGAGTA | TGTGAAGGTCCTGATGCCAT | TGCTCAGAGTACCAAAAGGC | CTGGTATCTTCCGATGGTCA | K00114:354:HC225BBXX:2:2219:13 | 1 |
| Intra-4 | Control1 | PC1minus | ALU | 221 | 164 | | | AGTTTTATGTAATTTCCTGGAGG | TGATCTCACATGGGCATAGT | AAAGGGTTCCAAAGTGGGG | GCATAGTGTAAGAGTTAAGAAGGTTG | K00114:354:HC225BBXX:7:1108:99 | 1 |
| Intra-5 | Control2 | LC2 | L1 | 151 | 140 | | | GCTCTTGAAATAAAATGGTATCTCT | CATTTTGCACAACTCCCTGC | AAATGGTATCTCTCCACCATGG | CATTTTGCACAACTCCCTGC | K00114:354:HC225BBXX:1:1113:13 | 1 |
| Intra-6 | Control5 | LC5 | L1 | 242 | 194 | | | GGCTTCCATGTCTTGAGTTAA | CACCAAAACCCACCACCAG | GCTATCGTGCTAGAGATACTGT | AGGGGCTGGAAAATACTACAG | K00114:354:HC225BBXX:1:1101:74 | 1 |
| Intra-7 | PD1 | LP1 | L1 | 251 | 213 | Yes | | ACAAATTGTATGCTCCAGGTCA | TTTCTGCTTGGGTCCTCTGT | GGTCAAGCTGTTAGGGTATTC | TGCCCAGCTACTCCCTTTA | K00114:354:HC225BBXX:2:2223:12 | 1 |
| Intra-8 | Control7 | FC7minus | ALU | 244 | 217 | Yes | Yes | CCAGTCAGGCCATGTTCT | GTGGTTAATAACTTATAACAATGCC | GGGCCCTAAAGAGAGAGA | ACTTATAACAATGCCTCCAGATCT | K00114:354:HC225BBXX:4:1225:80 | 1 |
| Intra-9 | Control1 | PC1plus | ALU | 174 | / | | | ACCACATGACAATTACAGTATCC | AGCCTGATTAAAAGAGAGCCCT | n/a | n/a | K00114:354:HC225BBXX:5:2125:14 | 1 |
| Intra-10 | Control1 | FC10plus | ALU | 204 | 177 | | | ACACTAATCAACTCAGTAACAATG | TGGTTAGGGTCAATCAGTGGT | TCAACTCAGTAACAATGAATACCC | TTCTCAAACCCCAGTCCAC | K00114:354:HC225BBXX:8:2109:45 | 1 |
| Intra-11 | Control2 | PC2minus | ALU | 207 | 193 | | | TGTTCTACTGACATCTGGACTT | AGGGAGATAACTGTGACCGG | TAAAACAGAATGCTGGCTGG | ACTGTGACCGGACATATGAC | K00114:354:HC225BBXX:7:1222:11 | 2 |
| Intra-12 | Control6 | LC6 | ALU | 242 | 184 | | | AATTTGGGCGACACACAAGG | GGACCACAGCCTAAGAACCA | TGAGGTGGAGTGGATGGA | GGCAGGTAAGAAAACTCTTGC | K00114:354:HC225BBXX:2:2210:29 | 1 |
| Intra-13 | Control7 | FC7minus | ALU | 228 | 212 | Yes | Yes | CCAGTCAGGCCATGTTCT | AACAATGCCTCCAGATCTAA | GCCCTAAAGAGAGAGATAAATCTA | AACAATGCCTCCAGATCTAA | K00114:354:HC225BBXX:4:1225:80 | 1 |
| Intra-14 | Control7 | PC7minus | ALU | 232 | 208 | Yes | | AGACTAGTAAACAGAAGGTGGCT | CAGGCATACTCAAGCTTTCT | AAGGTGGCTGGCTGATAA | AGCTTTCTTGTCCACAGATGA | K00114:354:HC225BBXX:8:2202:20 | 1 |
| Intra-15 | Control7 | TC7plus | ALU | 159 | 143 | Yes | Yes | AACAGGCTGAGAAAAGTGGC | TCACAAACCTGGGCTAAAG | AAAGTGGCTCACATCTGTTGG | CACAAACCTGGGCTAAAGAAA | K00114:354:HC225BBXX:4:1128:17 | 1 |
| Intra-16 | Control1 | TC1plus | L1 | 185 | 165 | | | GGTAACAGTAATCGGATTTGCC | AGTGGGAAATCGATAACATGGA | CGGATTTGCCCCAATGTTC | CGATAACATGGAATTTGAAGATCACT | K00114:354:HC225BBXX:3:2220:19 | 1 |
| Intra-17 | Control4 | PC4plus | L1 | 240 | 226 | Yes | | TTAGAACTTCAGGCTGCTGA | GGTAGGAGAAACTAATATGGTGAC | GGCTGCTGAAGGCATTTTTAC | GGAGAAACTAATATGGTGACAATTACC | K00114:354:HC225BBXX:5:1107:68 | 1 |
| Intra-18 | Control5 | KC5 | L1 | 207 | 193 | | | TGAGCAATATAGTTGACCCTT | TCCAGGTTGACACAAAGATG | ACCCTTGAGCAACACAGG | TCCAGGTTGACACAAAGATGTAAG | K00114:354:HC225BBXX:2:2218:20 | 1 |
| Intra-19 | Control6 | LC6 | L1 | 172 | 160 | | | AGGAGGACATGAGATAACATC | TCAGCTTCATACCCAGATGT | CTCAGACAATGGGAACCAG | CAGATGTAAAAATACCCAGCACAC | K00114:354:HC225BBXX:2:2114:1 | 1 |
| Intra-20 | PD2 | LP2 | ALU | 216 | 205 | Yes | | ACAACCAGACAACAAGTAGCTG | CCTCCACTGTCCCTTGCAT | AGACAACAAGTAGCTGAGTTACA | CTGTCCCTTGCATGGTGT | K00114:354:HC225BBXX:2:2128:76 | 1 |
| Intra-21 | PD3 | PP3minus | ALU | 251 | 207 | | | TGTCCTGCACACAATCCTAGT | TGTGCCTAGGAGAGAAGCTG | GGTTTCCATTTCTTTTCTGAGCA | CTGAAGCAAAGACCAGATGTG | K00114:354:HC225BBXX:3:1123:1 | 1 |
| Intra-22 | PD4 | PP4plus | ALU | 202 | 160 | | | TTCTATTGCTTAGCCCCAGG | GTTGAGAGAAGGGCCATGGT | CCCTCACAATGTGAGGTTTC | AACTGAATCCCAGAGGTCTG | K00114:354:HC225BBXX:7:1204:89 | 1 |
| Intra-23 | PD6 | TP6minus | ALU | 214 | 201 | Yes | Yes | AGATGTGATAGCTTGCCAGGA | TCTCAAGTGGTCGCCTCTG | TGATAGCTTGCCAGGATAAGTT | GTCGCCTCTGTTTGTCAAA | K00114:354:HC225BBXX:8:1117:21 | 1 |
| Intra-24 | PD10 | KP10 | L1 | 189 | / | | | CCCTCCAGAGAAGGTTTGCT | GCACTTTCCACAAGATACATGTC | GGTTTGCTTACTGTTCAGTATAATTTGG | AAACAACCTACAAACCAGCATG | K00114:354:HC225BBXX:2:1105:20 | 1 |
| Intra-25 | PD3 | KP3 | L1 | 228 | 187 | | | AAAGTCATGGATGTCTGGTAC | CACTTTGTGCCAGACCCAC | TGGATGTCTGGTACACATAATTTGAT | AGAAAGAGGCCTAGACACATG | K00114:354:HC225BBXX:1:2114:85 | 1 |
| Intra-26 | PD8 | PP8minus | L1 | 236 | 220 | | | AAGTACCAAGAAAGGAGCCA | GTGTATCTGATTCATCCCATTTC | GGAGCCAGAAGAGTCAGAG | TCTGATTCATCCCATTTCTAATTCC | K00114:354:HC225BBXX:4:2117:30 | 1 |
| Intra-27 | AD1 | LA1 | L1 | 236 | 205 | | | AGACGGTGCATTCTCCATTG | GAAGCGTTAGAACACCACCA | GTTATCTGCTTAGGGGTGC | CCACCAAATCACCCCAAA | K00114:354:HC225BBXX:1:1223:20 | 1 |
| Intra-28 | AD5 | TA5minus | L1 | 164 | 149 | | | CTTGATGACTTTAATGTAACAGCA | TTCCTCTCTGCATTTGGGC | CTTGATGACTTTAATGTAACAGCATTCATTT | GGCTAGCCTTCTATAGAAGCA | K00114:354:HC225BBXX:3:2124:59 | 1 |
| Intra-29 | AD6 | PA6plus | L1 | 225 | 198 | Yes | Yes | AGCATCTTACACAATCATTAGCA | TGTAAGTGTAGAAGTCCAAGAGG | TCATTAGCAGAAATTCGACTCCG | CCAAGAGGCTACTGGAATTAGA | K00114:354:HC225BBXX:4:2217:78 | 1 |
| Intra-30 | AD8 | PA8minus | ALU | 204 | / | | | GCTCCCTGATCACCTCTTCA | GCGAGACGTGCCTATGTG | ATCACCTCTTCAGAAGGTTTCA | AGTATAAATAATGTATATTGGCCGGGT | K00114:354:HC225BBXX:6:1101:13 | 1 |

**PCR validation of putative polymorphic NAHR events found in capture-seq libraries and ONT PromethION WGS libraries**

| Target ID | RE-1 Repeat Masker Annotation | | | | | RE-2 Repeat Masker Annotation | | | | | PromethION libraries count | PromethION libraries | Capture-seq libraries count | Capture-seq libraries | PromethION library chosen for primers design and PCR | FASTQ ID of read u | PCR amplicon of expected size | Validated by Sanger | FW | REV |
|---|---|---|---|---|---|---|---|---|---|---|---|---|---|---|---|---|---|---|---|---|
| Poly-Inter-1 | chr3 | 75414206 | 75414466 | + | AluYc | chr7 | 6922900 | 6923206 | + | AluY | 5 | TA1plus \| TC6plus \| TC8plus \| KC6 | 6 | KA1 \| LC5 \| PA10plus \| PP10plus | TC6plus | 97b33cf6-452e-400 | Yes | Yes | TCCAGAAGGTTCTG | TGGGTCTGTTATTGTGTGG |
| Poly-Inter-2 | chr22 | 23510368 | 23510489 | - | AluSx4 | chr9 | 129424466 | 129424600 | + | AluSx3 | 16 | TA10plus \| TA1plus \| TA3plus \| TA | 4 | FA10minus \| FC2plus \| FC8plus \| | TC5plus | 8fbc319f-c6cd-4cd | No | / | TAAAGACACTCCAA | AGATCCGACAGAATCACCA |
| Poly-Intra-1 | chr19 | 52816682 | 52816834 | + | AluSx | chr19 | 52853406 | 52853696 | + | AluSx | 10 | TA1plus \| TA4plus \| TA8plus \| TC | 9 | FA1minus \| LC3 \| LC4 \| LP9 \| PA | TC6plus | 432bfb80-57c8-45b | Yes | Yes | CTTATGGCCAAGGG | GAGAGGCTTGTGAAACAGG |
| Poly-Intra-2 | chr19 | 52888468 | 52888643 | + | AluSc8 | chr19 | 52915999 | 52916321 | + | AluJb | 14 | TA7plus \| TA8plus \| TC10plus \| TC | 7 | FA4plus \| FC4plus \| LC6 \| PA1plu | TC10plus | a2e1256c-08ca-47 | Yes | No | AGTGACATTCAGTA | GGGACAAATGACAACCCTTT |
| Poly-Intra-3 | chr15 | 20320009 | 20320301 | - | AluSx | chr15 | 20337957 | 20338260 | - | AluSx1 | 14 | TA1plus \| TA10plus \| TA4plus \| TA | 51 | FA10minus \| FA7plus \| FC10min | TA9plus | 40ba957e-f989-49 | Yes | Yes | TCTCACATTGCAATT | CAAACCCCAAATTCTAGAGGA |
| Poly-Intra-4 | chr12 | 79760846 | 79761157 | - | AluSq2 | chr12 | 79768800 | 79769037 | - | AluSc | 1 | TC7plus | 9 | KC7 \| LA1 \| PC3minus \| PC3plus | TC7plus | a7d0a60e-4a1b-47 | Yes | Yes | GAGCAAGTCTAGCA | GAATATGCTAATATTGGCTGAGCA |
| Poly-Intra-5 | chr5 | 181095728 | 181095915 | + | AluSx | chr5 | 181103475 | 181103783 | - | AluYk2 | 1 | TA9plus | 6 | FA9minus \| FA9minus \| KA9 \| LF | TA9plus | f8813092-18ad-4cc | Yes | Yes | GCAACAGAACAGA | CCGCAAGAACACGATAATCC |
| Poly-Intra-6 | chr3 | 132270115 | 132270397 | + | AluSz6 | chr3 | 132276184 | 132276347 | + | AluSc | 10 | TA10plus \| TA3plus \| TA4plus \| TA | 45 | FA10minus \| FA1minus \| FA2mi | TC9plus | 3a39e1df-3942-47 | No | / | ATTCTGGTGGAAG | GATAATTTCTCTATGGACGGGG |
| Poly-Intra-7 | chr14 | 58013757 | 58014047 | + | AluSx1 | chr14 | 58019592 | 58019808 | + | AluSx | 1 | TA5plus | 3 | FA5plus \| TA5minus \| TA5plus | TA5plus | 43f742a2-745b-43 | Yes | Yes | ATACTGATGATGGC | CATTGGTAGGATGCATGGAA |
| Poly-Intra-8 | chr8 | 40916868 | 40917180 | - | AluSx1 | chr8 | 40922039 | 40922334 | - | AluSc8 | 14 | TA1plus \| TA3plus \| TA4plus4 \| TA | 65 | FA1plus \| FA3plus \| FA5plus \| FA | TA6plus | cf483882-ef9d-4e7 | Yes | Yes | CTTGGTGGAGCATG | CCTTGGGGTCCTAAATTCC |
| Poly-Intra-9 | chr6 | 17376503 | 17376817 | + | AluY | chr6 | 17381033 | 17381342 | + | AluSx | 8 | TA6plus \| TA8plus \| TC6plus \| TC | 6 | FA1plus \| FA9minus \| KA1 \| KP1 | TA6plus | b86c03ff-5742-452 | Yes | Yes | AGACACACAAACCC | ATGGTCAGCAAACCAAGAG |
| Poly-Intra-10 | chr6 | 125861755 | 125862067 | + | AluSp | chr6 | 125866047 | 125866348 | + | AluSp | 11 | TA1plus \| TA10plus \| TA4plus \| TA | 14 | FA10plus \| FA4minus \| FA4plus | TC6plus | 8dd40f47-5cd7-471 | Yes | Yes | GTTAGGCCTGTGCT | GACCACTCTCAAATCAATGC |
| Poly-Intra-11 | chr2 | 172041178 | 172041476 | + | AluSx4 | chr2 | 172045379 | 172045683 | + | AluSp | 5 | TA7plus \| TC5plus \| TC8plus \| KC8 | 18 | FC8minus \| FP9minus \| KC5 \| KC | TC5plus | 63efc126-b1f0-436 | No | / | TACAAGAACTGAAG | GTAGGGTCAGGCTTATCGTA |
| Poly-Intra-12 | chr1 | 174003106 | 174003382 | + | AluSq | chr1 | 174007107 | 174007403 | + | AluY | 2 | TA7plus \| TC5plus | 14 | FA7minus \| FA7plus \| FC5plus \| | TC5plus | 8fa2551a-c651-47f | Yes | Yes | AGTCATTCAATCCTT | TAACTGGAGTGCTTTGTAACAG |
| Poly-Intra-13 | chr8 | 42332719 | 42333022 | + | AluSp | chr8 | 42336679 | 42336973 | + | AluSq2 | 20 | TA1plus \| TA10plus \| TA2plus \| TA | 128 | FA2minus \| FA2plus \| FA3minus | TC7plus | 96a47262-4166-45 | Yes | Yes | AAATGCCAACCAAA | GTAGGGAATTGTGGGATCA |
| Poly-Intra-14 | chr10 | 27181658 | 27181792 | + | AluSx4 | chr10 | 27185354 | 27185638 | + | AluSx | 1 | TA9plus | 2 | FA9minus \| TA9plus | TA9plus | 28c4c979-913d-47 | Yes | Yes | GCTGGAATGAAAG | AGGTGCCTGGTTTGTATTTG |
| Poly-Intra-15 | chr16 | 85117121 | 85117446 | - | AluY | chr16 | 85120814 | 85121125 | - | AluY | 1 | TC9plus | 10 | FC4plus \| FC9minus \| KC9 \| LC4 | TC9plus | 9d3edeb3-13ff-479 | Yes | Yes | GTTGTTTTGGGCAA | CACCCATCAAATCCAGAGG |
| Poly-Intra-16 | chr6 | 157278754 | 157279041 | - | AluSq | chr6 | 157282118 | 157282420 | - | AluSx | 5 | TA7plus \| TC10plus \| TC7plus \| KC | 20 | FA7minus \| FC10plus \| FP5plus \| | TC7plus | 6edd0985-a1d0-4a | Yes | Yes | CCACCCTGAAAACA | AGGCAAGACGGAGAGAAG |
| Poly-Intra-17 | chr6 | 22050609 | 22050900 | - | AluSc8 | chr6 | 22053965 | 22054311 | - | AluSx | 19 | TA1plus \| TA10plus \| TA2plus \| TA | 28 | FA1minus \| FA9plus \| FC2plus \| | TC7plus | a7e65086-3ad6-40 | Yes | Yes | TCAAGCATTTCTCCC | GGTTAAAGCCTGGATACGG |
| Poly-Intra-18 | chr21 | 29348812 | 29349109 | + | AluY | chr21 | 29352052 | 29352353 | + | AluSz | 1 | TC10plus | 2 | FP9minus \| TP2plus | TC10plus | e05222f0-a245-43 | Yes | Yes | TACAAATCCAAGCC | CTGAGCCAACATCCCTGTA |

### Table S6 - Calculation for thresholds of Recombination Index

#### CONTROL

| | Subfamily | Total number of elements found reocmbined at least once in aggregated capture-seq of post-mortem samples | Total number of recombination events in aggregated capture-seq of post-mortem samples | Total elements annotated in RepeatMasker | Recombination per element expected by random genomic distribution (random threshold) | Rounded | Number of elements exceeding the random threshold (hot elements) | % of elements exceeding the random threshold (% of hot elements per subfamily) |
|---|---|---|---|---|---|---|---|---|
| ALU | Alu Y | 128662 | 1191417 | 139234 | 8.556940115 | 8.6 | 57408 | 41.23130844 |
| | Alu S | 613460 | 2689306 | 678131 | 3.965761778 | 4 | 257928 | 38.03512891 |
| | Alu J | 174805 | 350730 | 309536 | 1.133083066 | 1.1 | 46103 | 14.89422878 |

| | Subfamily | Total number of elements found reocmbined at least once in aggregated capture-seq of post-mortem samples | Total number of recombination events in aggregated capture-seq of post-mortem samples | Total elements annotated in RepeatMasker | Recombination per element expected by random genomic distribution (random threshold) | Rounded | Number of elements exceeding the random threshold (hot elements) | % of elements exceeding the random threshold (% of hot elements per subfamily) |
|---|---|---|---|---|---|---|---|---|
| L1 | L1PA7 | 10612 | 69469 | 12978 | 5.352827863 | 5.4 | 4814 | 45.36373916 |
| | L1PA4 | 10137 | 94993 | 11834 | 8.027125232 | 8 | 4299 | 42.40899674 |
| | L1PA5 | 9689 | 98486 | 11228 | 8.771464197 | 8.8 | 4845 | 50.00516049 |
| | L1PA3 | 8766 | 72042 | 10757 | 6.697220415 | 6.7 | 3869 | 44.13643623 |
| | L1PA8 | 7866 | 31080 | 8047 | 3.862308935 | 3.9 | 3132 | 39.81693364 |
| | L1PA6 | 4946 | 44488 | 5887 | 7.556989978 | 7.6 | 2432 | 49.17104731 |
| | L1PA10 | 4741 | 14292 | 7007 | 2.039674611 | 2 | 1713 | 36.1316178 |
| | L1PA13 | 4140 | 5074 | 8829 | 0.574697021 | 0.6 | 566 | 13.67149758 |
| | L1PA2 | 4007 | 28271 | 4940 | 5.722874494 | 5.7 | 1702 | 42.47566758 |
| | L1PA11 | 2801 | 6862 | 4086 | 1.679393049 | 1.7 | 1149 | 41.02106391 |
| | L1PA16 | 2518 | 1262 | 13883 | 0.090902543 | 0.1 | 114 | 4.527402701 |
| | L1PA15 | 2296 | 1539 | 8220 | 0.187226277 | 0.2 | 257 | 11.19337979 |
| | L1PA8A | 1500 | 5847 | 2438 | 2.398277276 | 2.4 | 731 | 48.73333333 |
| | L1HS | 1000 | 4767 | 1620 | 2.942592593 | 2.9 | 471 | 47.1 |
| | L1PA14 | 782 | 522 | 3037 | 0.171880145 | 0.2 | 81 | 10.35805627 |
| | L1PA12 | 720 | 1148 | 1756 | 0.653758542 | 0.7 | 305 | 42.36111111 |
| | L1PA17 | 577 | 296 | 4723 | 0.06267203 | 0.1 | 27 | 4.679376083 |

## PD

| | Subfamily | Total number of elements found reocmbined at least once in aggregated capture-seq of post-mortem samples | Total number of recombination events in aggregated capture-seq of post-mortem samples | Total elements annotated in RepeatMasker | Recombination per element expected by random genomic distribution (random threshold) | Rounded | Number of elements exceeding the random threshold (hot elements) | % of elements exceeding the random threshold (% of hot elements per subfamily) |
|---|---|---|---|---|---|---|---|---|
| ALU | Alu Y | 127947 | 1036141 | 139234 | 7.441724004 | 7.4 | 54463 | 39.11616416 |
| | Alu S | 603439 | 2374191 | 678131 | 3.501080175 | 3.5 | 208936 | 30.8105661 |
| | Alu J | 167854 | 320364 | 309536 | 1.034981392 | 1 | 39275 | 12.68834643 |

| | Subfamily | Total number of elements found reocmbined at least once in aggregated capture-seq of post-mortem samples | Total number of recombination events in aggregated capture-seq of post-mortem samples | Total elements annotated in RepeatMasker | Recombination per element expected by random genomic distribution (random threshold) | Rounded | Number of elements exceeding the random threshold (hot elements) | % of elements exceeding the random threshold (% of hot elements per subfamily) |
|---|---|---|---|---|---|---|---|---|
| L1 | L1PA7 | 10612 | 48061 | 12978 | 3.703267067 | 3.7 | 4695 | 44.24236713 |
| | L1PA4 | 10137 | 64472 | 11834 | 5.448031097 | 5.4 | 4256 | 41.98480813 |
| | L1PA5 | 9689 | 67509 | 11228 | 6.012557891 | 6 | 3929 | 40.55114047 |
| | L1PA3 | 8766 | 50454 | 10757 | 4.690341173 | 4.7 | 3673 | 41.90052475 |
| | L1PA8 | 7866 | 22042 | 8047 | 2.73915745 | 2.7 | 2738 | 34.80803458 |
| | L1PA6 | 4946 | 30556 | 5887 | 5.190419569 | 5.2 | 2113 | 42.72139102 |
| | L1PA10 | 4741 | 10232 | 7007 | 1.460254032 | 1.5 | 1671 | 35.24572875 |
| | L1PA13 | 4140 | 4104 | 8829 | 0.464831804 | 0.5 | 1007 | 24.3236715 |
| | L1PA2 | 4007 | 20694 | 4940 | 4.189068826 | 4.2 | 1471 | 36.71075618 |
| | L1PA11 | 2801 | 5344 | 4086 | 1.307880568 | 1.3 | 823 | 29.38236344 |
| | L1PA16 | 2518 | 1311 | 13883 | 0.094432039 | 0.1 | 147 | 5.83796664 |
| | L1PA15 | 2296 | 1303 | 8220 | 0.158515815 | 0.2 | 209 | 9.102787456 |
| | L1PA8A | 1500 | 4230 | 2438 | 1.735028712 | 1.7 | 713 | 47.53333333 |
| | L1HS | 1000 | 3544 | 1620 | 2.187654321 | 2.2 | 368 | 36.8 |
| | L1PA14 | 782 | 413 | 3037 | 0.135989463 | 0.1 | 64 | 8.184143223 |
| | L1PA12 | 720 | 945 | 1756 | 0.538154897 | 0.5 | 262 | 36.38888889 |
| | L1PA17 | 577 | 291 | 4723 | 0.061613381 | 0.1 | 27 | 4.679376083 |

## AD

| | Subfamily | Total number of elements found reocmbined at least once in aggregated capture-seq of post-mortem samples | Total number of recombination events in aggregated capture-seq of post-mortem samples | Total elements annotated in RepeatMasker | Recombination per element expected by random genomic distribution (random threshold) | Rounded | Number of elements exceeding the random threshold (hot elements) | % of elements exceeding the random threshold (% of hot elements per subfamily) |
|---|---|---|---|---|---|---|---|---|
| ALU | Alu Y | 126892 | 871923 | 139234 | 6.262285074 | 6.3 | 49941 | 35.86839421 |
| | Alu S | 582485 | 1926184 | 678131 | 2.840430536 | 2.8 | 226208 | 33.35756661 |
| | Alu J | 142588 | 237802 | 309536 | 0.768253127 | 0.8 | 22922 | 7.405277577 |

| | Subfamily | Total number of elements found reocmbined at least once in aggregated capture-seq of post-mortem samples | Total number of recombination events in aggregated capture-seq of post-mortem samples | Total elements annotated in RepeatMasker | Recombination per element expected by random genomic distribution (random threshold) | Rounded | Number of elements exceeding the random threshold (hot elements) | % of elements exceeding the random threshold (% of hot elements per subfamily) |
|---|---|---|---|---|---|---|---|---|
| L1 | L1PA7 | 10612 | 44577 | 12978 | 3.43481276 | 3.4 | 4346 | 40.95363739 |
| | L1PA4 | 10137 | 62578 | 11834 | 5.287983776 | 5.3 | 4113 | 40.57413436 |
| | L1PA5 | 9689 | 65107 | 11228 | 5.798628429 | 5.8 | 4503 | 46.47538446 |
| | L1PA3 | 8766 | 48444 | 10757 | 4.503486102 | 4.5 | 3516 | 40.10951403 |
| | L1PA8 | 7866 | 20258 | 8047 | 2.517459923 | 2.5 | 2471 | 31.41367913 |
| | L1PA6 | 4946 | 29243 | 5887 | 4.967385765 | 5 | 2007 | 40.57824505 |
| | L1PA10 | 4741 | 8952 | 7007 | 1.277579563 | 1.3 | 1368 | 28.85467201 |
| | L1PA13 | 4140 | 3233 | 8829 | 0.366179635 | 0.4 | 751 | 18.14009662 |
| | L1PA2 | 4007 | 19390 | 4940 | 3.925101215 | 3.9 | 1677 | 41.85175942 |
| | L1PA11 | 2801 | 4526 | 4086 | 1.107684777 | 1.1 | 684 | 24.41985005 |
| | L1PA16 | 2518 | 842 | 13883 | 0.060649715 | 0.1 | 64 | 2.541699762 |
| | L1PA15 | 2296 | 878 | 8220 | 0.106812652 | 0.1 | 106 | 4.616724739 |
| | L1PA8A | 1500 | 3707 | 2438 | 1.520508614 | 1.5 | 649 | 43.26666667 |
| | L1HS | 1000 | 3427 | 1620 | 2.115432099 | 2.1 | 375 | 37.5 |
| | L1PA14 | 782 | 312 | 3037 | 0.10273296 | 0.1 | 32 | 4.092071611 |
| | L1PA12 | 720 | 759 | 1756 | 0.432232346 | 0.4 | 204 | 28.33333333 |
| | L1PA17 | 577 | 161 | 4723 | 0.034088503 | 0 | 8 | 1.386481802 |
