## Supplementary Document 1 for "Recombination of repeat elements generates somatic complexity in human genomes"

**Supplementary Document-1: Technical protocol for Capture-seq and reagents**

**Adapted from Gnirke et al, 2009 and from Fisher at al, 2011.**

**Adapters annealing**

1. Prepare the following mix by using equimolar amounts of forward and reverse single-stranded adapters:

- Adapter 1-Forward*P: n moles
- Adapter 1-Reverse: n moles
- NaCl 5M: 20µl
- $H_2O$: to 100µl

- Adapter 2-Forward*P: n moles
- Adapter 2-Reverse: n moles
- NaCl 5M: 20µl
- $H_2O$: to 100µl

2. Heat to 94°C for 5' and cool to 25C over 45 minutes.

3. Add 0.1x volume NaAc 3M, 2 volumes of EtOH and precipitate overnight at -20°C. Centrifuge at 13000 g for 30', wash pellet once with EtOH 75% and resuspend in 40µl of nuclease-free water. After measuring concentration, check the quality of the annealing on acrylamide gel, aliquot the adapters for single-use reactions and store at -80°C.

**DNA shearing.**

Note: Shearing is performed on a Covaris S220 machine using proprietary glass microtubes.

1. Dilute 1-5µg intact genomic DNA in 130µl $H_20$ or TE and load in Covaris SnapCap tube (try to avoid making bubbles in the tube)

2. Shear the DNA using a program to produce a 200bp fragments distribution (Duty cycle 10% - Intensity 5% - Cycler 200, 60s x 3 times).

3. After shearing, clean the DNA with AMPure XP (1.8x volume). Elute the DNA in 40µl $H_2O$.

**Preparation of DNA for capture and sequencing**

End repair / dA-tailing / Paired End Adaptors ligation / Library size selection (200bp) and Pre-Capture LM-PCR are performed using NEBNext DNA Library Prep Master Mix Set for Illumina (E6040) according to manufacturer's specifications. For Adapters ligation, the suggested molar ratio Adapters:DNA is 10:1. Small aliquots of DNA before/after the ligation/Before LM-PCR/After LM-PCR can be saved for Bioanalyzer quality control.

**Preparation of biotinylated riboprobes for DNA/RNA hybridization and capture.**

Note: the sequences of the ssDNA oligonucleotides matching the 5'- and 3'- regions of L1HS and AluYa5/b8/b9 elements are in Supplementary Document-2.
First and second PCR reactions are performed on 96-wells plates using one single oligonucleotide as template per well.

1. Dilute the oligonucleotides at 50ng/ul and perform the first PCR with Takara Ex Taq Hot Start in 25ul reaction volume (1ul ssDNA template oligonucleotide, 0.125ul Takara ExTaq HS, 2.5ul 10X buffer, 2ul dNTPs mix, Primer A-Fw 1uM, Primer B-Rev 1uM, $H_2O$ to 25ul).

2. Perform first PCR:

| | | |
|---|---|---|
| 1 cycle: | 5 min | 94°C |
| 8 cycles: | 20 sec | 94°C |
| | 30 sec | 55°C |
| | 30 sec | 72°C |

3. The second PCR is performed again with Takara ExTaq Hot Start version in 25ul reaction. Use 1ul from the first PCR reactions as template. The primers used for the second PCR are Primer T7-A-Fw and Primer B-Rev.

| | | |
|---|---|---|
| 1 cycle: | 5 min | 94°C |
| 10 cycles: | 20 sec | 94°C |
| | 30 sec | 55°C |
| | 30 sec | 72°C |

Pool in a single tube 2ul of the second PCR products, keep L1HS and AluY PCR products separated in two different tubes. Load 100ul from each tube on a preparative agarose gel and perform purification and cleaning of the DNA products with QIAquick Gel Extraction Kit. Elute in 40 ul $H_2O$ and measure the concentration with Nanodrop.

**In vitro transcription of biotinylated riboprobes.**

**1.** The reaction is performed using the T7 MAXIscript Transcription Kit in 100ul volume following manufacturer's instructions. Biotin is introduced using a Biotin RNA labeling mix. Two different reactions are performed for L1HS and AluY templates. Prepare the following mix:

| | |
|---|---|
| Template: | 500ng purified dsDNA oligos |
| Biotin RNA labeling mix: | 10µl |
| 10x buffer: | 10µl |
| T7 Enzyme: | 10µl |
| RNase Inhibitors: | 1µl |
| $H_2O$: | to 100ul |

2. Incubate the samples at 37C for 2 hours and stop the reaction with EDTA 0.5M (see manufacturer's protocol for details)

**3.** Add 0.1x volume of NaAc 3M and 2.5x volumes EtOH, mix well and incubate overnight at -80C. Precipitate the probes by centrifuging at 13000g for 30 minutes at 4C, wash the pellets once with 75% EtOH, air-dry them briefly and resuspend in RNAse free $H_2O$.

**Preparation of M-280 streptaviding beads.**

For each single DNA/RNA hybridization reaction aliquot 50ul of M-280 streptavidin beads per reaction and perform washes according to manufacturer's instructions. After the final wash resuspend the beads in 165ul 1x wash buffer per reaction.

**DNA/RNA hybridization**

**Note:** we have noticed an increase of L1HS enrichment efficiency when the hybridization is performed with separate reactions for L1HS and AluY.

1. Prepare hybridization buffer and heat it at 65C until clear with occasional shaking. Keep the buffer at 65C until use.

| | |
|---|---|
| 20x SSPE: | 500µl |
| H2O: | 260µl |
| 50x Denhardt's: | 200µl |
| 10% SDS: | 20µl |
| 0,5M EDTA: | 20µl |

2. Prepare the DNA samples:

| | |
|---|---|
| Adapters-ligated, LM-PCR amplified DNA: | 300ng for L1HS / 100ng for AluY |
| Salmon Sperm DNA: | 2.5ul |
| Blocking oligo-1 30uM: | 1.5ul |
| Blocking oligo-2 30uM: | 1.5ul |
| $H_2O$: | to 13ul |

3. Prepare RNA probes:

| | |
|---|---|
| Biotinylated RNA probes: | 300ng L1HS / 300ng AluY |
| RNAsin RNAse Inhibitor: | 1ul |
| $H_2O$: | to 7ul |

4. Set a thermocycler with the following program:

| | |
|---|---|
| 5min | 95°C |
| 5min | 65°C |
| 5min | 65°C |

Put the DNA sample in the thermocycler at 95°C for 5min; let the thermocycler go down to 65°C and keep for 5 minutes; in the meanwhile, put the RNA sample in a thermomixer or a thermoplate at 65°C and heat it for 2,5min (synchronize the heating of RNA sample so that when the 2,5min have passed, the DNA sample in the thermocycler will have completed the 5min at 65°C). After the 2,5min have passed, add the RNA sample to the DNA sample in the thermocycler, and add 13µl of hybridization sample to the DNA/RNA sample. Mix the sample by pipetting 10 times, close the lid and keep at 65°C for 5 minutes. Perform pipetting steps gently but quickly to avoid reducing the volume of reaction.

**DNA/RNA hybrids capture.**

1. Prepare the low-stringency and high-stringency wash buffers. Incubate the high-stringency buffer at 68C.

Low-stringency wash buffer:

| | |
|---|---|
| Nuclease-free water: | 23.5ml |
| 20× SSC: | 1.25ml |
| 10% SDS: | 250µl |

High-stringency wash buffer:

| | |
|---|---|
| Nuclease-free water: | 24.7ml |
| 20× SSC: | 125µl |
| 10% SDS: | 250µl |

2. After the hybridization is completed remove the samples from thermocycler or thermoblock and transfer each reaction to a labeled 1.5ml microcentrifuge tube

3. Add to each tube 165ul of washed and well-resuspended M-280 beads, resuspend again by pipetting and let the capture proceed for 5 minutes at RT. Tickle the tubes gently a few times during the 5 minutes to prevent the beads from depositing on the bottom of the tubes.

4. Add 165ul of low-stringency buffer to each sample, pipette to mix and incubate at RT for 15 minutes. Place the tubes on a magnet separator stand and allow the beads to separate for 2 minutes. Discard the supernatant.

5. Add 165ul of pre-warmed high-stringency buffer, vortex briefly to resuspend the beads and incubate in a heating block set at 68C for 10 minutes. Place the tubes on a magnet separator stand and allow the beads to separate for 2 minutes. Discard the supernatant. Repeat for a total of 3 washes.

6. After having discarded the supernatant from the third wash denature DNA/RNA hybrids by adding 50ul of 0.1N NaOH in each tube. Resuspend the beads by pipetting up and down gently for 15 times and incubate at RT for 10 minutes, occasionally tickling the tubes to avoid beads precipitation. Transfer the tubes to a magnetic stand separator for 2 minutes and transfer the supernatant containing the captured ssDNA to clean labeled tubes.

7. Add 50ul 1M Tris HCl pH 8 to neutralize the NaOH.

8. Clean the samples with 1.8x volumes AMPure XP and after the standard washes elute the ssDNA in 22ul nuclease-free $H_2O$.

**Enrichment of captured DNA by LM-PCR**

1. Prepare the following mix for each sample:

| | |
|---|---|
| 2X Phusion Hot Start Flex 2X master mix: | 25µl |
| Universal PE Forward primer 10uM: | 2.5µl |
| INDEX PE Reverse primer 10uM: | 2.5µl |
| Template captured ssDNA: | 20ul |

Run the following PCR program:

| | | |
|---|---|---|
| 1 cycle: | 30sec | 98°C |
| 12 cycles: | 10 sec | 98°C |
| | 1min | 72°C |
| 1 cycle: | 2min | 72°C |

2. Clean the reactions with 1.8x volumes of AMPure XP beads. Elute the libraries in 30ul nuclease-free $H_2O$.

**Quality control and sequencing.**

1. Run a qPCR with reagents of choice to calculate the enrichment efficiency for L1HS and AluY for all capture libraries. The primer sequences can be found in the "Primers list" section. For each library run in triplicate wells 100pg/reaction of pre-capture library, 100pg/reaction of L1HS post-capture library and 100pg/reaction of AluY post-capture library. The enrichment folds can be calculated as $2^{\Delta Ct(Pre\text{-}post)}$ (Supplementary Fig.3)

2. Calculate the concentration of L1HS and AluY libraries with Nanodrop and perform quality control on Bioanalyzer High Sensitivity DNA Kit. Calculate the molarity of L1HS and AluY libraries for the same DNA sample and make an equimolar mixture. The libraries are then loaded again on Bioanalyzer for a second quality control of the profile and molarity calculation.

3. Make an equimolar pool of the complete libraries with different indexes according to the initial experimental design and perform paired-end sequencing on Illumina platform using the primers indicated in the "Primers list" section.

**List of reagents and materials.**
Covaris S220 (Covaris)
microTUBE Snap-Cap AFA Fiber (Covaris)
Agencourt AMPureXP beads (Beckman Coulter)
NEBNext DNA Library Prep Master Mix Set for Illumina (New England Biolabs)
Ex Taq DNA Polymerase, Hot-Start Version (Takara)
QIAquick Gel Extraction kit (Qiagen)
T7 MAXIscript Transcription Kit (Thermo Fisher Scientific)
Biotin RNA labeling mix (Roche Life Science)
RNasin Ribonuclease Inhibitors (Promega)
Dynabeads M-280 Streptavidin (Thermo Fisher Scientific)
20X SSPE (Sigma-Aldrich)

Denhardt's solution 50x (Sigma-aldrich)
Salmon Sperm DNA, sheared (10 mg/mL) (Thermo Fisher Scientific)
20x SSC (Sigma-Aldrich)
Phusion Hot Start Flex 2X Master Mix (New England Biolabs)
SYBR Premix Ex Taq II (Tli RNaseH Plus) (Takara)

**List of PCR primers and other DNA sequences.**

**Primers for amplification of ssDNA oligonucleotides (probes)**
FIRST PCR
Primer A (FW): CTCACTATAGGGATCGCACCAGCGTGT
Primer B (REV): CGTGGATGAGGAGCCGCAGTG

SECOND PCR
Primer T7-A (FW): GGATTCTAATACGACTCACTATAGGGATCGCACCAGCGTGT
Primer B (REV: CGTGGATGAGGAGCCGCAGTG

**Single-strand Adapters for Paired-End sequencing**
Adapter1-FW: 5'-PHOSPH/GATCGGAAGAGCGTCGTGTAGGGAAAGAGTGT
Adapter1-REV: ACACTCTTTCCCTACACGACGCTCTTCCGATCT

Adapter2-FW: 5'-PHOSPH/GATCGGAAGAGCGGTTCAGCAGGAATGCCGAG
Adapter2-REV: CTCGGCATTCCTGCTGAACCGCTCTTCCGATCT

**Indexing primers for capture libraries construction and post-capture amplification (indexes are in bold)**

Universal-FW:
AATGATACGGCGACCACCGAGATCTACACTCTTTCCCTACACGACGC
REV-1:
CAAGCAGAAGACGGCATACGAGAT**CGTGAT**CTCGGCATTCCTGCTGAACC
REV-2:
CAAGCAGAAGACGGCATACGAGAT**GCCTAA**CTCGGCATTCCTGCTGAACC
REV-3:
CAAGCAGAAGACGGCATACGAGAT**TGGTCA**CTCGGCATTCCTGCTGAACC
REV-4:
CAAGCAGAAGACGGCATACGAGAT**CACTGT**CTCGGCATTCCTGCTGAACC
REV-5:
CAAGCAGAAGACGGCATACGAGAT**ATTGGC**CTCGGCATTCCTGCTGAACC
REV-6:
CAAGCAGAAGACGGCATACGAGAT**GATCTG**CTCGGCATTCCTGCTGAACC
REV-7:
CAAGCAGAAGACGGCATACGAGAT**TCAAGT**CTCGGCATTCCTGCTGAACC
REV-8:
CAAGCAGAAGACGGCATACGAGAT**CTGATC**CTCGGCATTCCTGCTGAACC
REV-9:
CAAGCAGAAGACGGCATACGAGAT**AAGCTA**CTCGGCATTCCTGCTGAACC

REV-10:
CAAGCAGAAGACGGCATACGAGAT**GTAGCC**CTCGGCATTCCTGCTGAACC
REV-11:
CAAGCAGAAGACGGCATACGAGAT**TACAAG**CTCGGCATTCCTGCTGAACC
REV-12:
CAAGCAGAAGACGGCATACGAGAT**TTGACT**CTCGGCATTCCTGCTGAACC
REV-13:
CAAGCAGAAGACGGCATACGAGAT**GGAACT**CTCGGCATTCCTGCTGAACC
REV-14:
CAAGCAGAAGACGGCATACGAGAT**TGACAT**CTCGGCATTCCTGCTGAACC
REV-15:
CAAGCAGAAGACGGCATACGAGAT**GGACGG**CTCGGCATTCCTGCTGAACC
REV-16:
CAAGCAGAAGACGGCATACGAGAT**CTCTAC**CTCGGCATTCCTGCTGAACC
REV-17:
CAAGCAGAAGACGGCATACGAGAT**GCGGAC**CTCGGCATTCCTGCTGAACC
REV-18:
CAAGCAGAAGACGGCATACGAGAT**TTTCAC**CTCGGCATTCCTGCTGAACC
REV-19:
CAAGCAGAAGACGGCATACGAGAT**GGCCAC**CTCGGCATTCCTGCTGAACC
REV-20:
CAAGCAGAAGACGGCATACGAGAT**CGAAAC**CTCGGCATTCCTGCTGAACC
REV-21:
CAAGCAGAAGACGGCATACGAGAT**CGTACG**CTCGGCATTCCTGCTGAACC
REV-22:
CAAGCAGAAGACGGCATACGAGAT**CCACTC**CTCGGCATTCCTGCTGAACC
REV-23:
CAAGCAGAAGACGGCATACGAGAT**GCTACC**CTCGGCATTCCTGCTGAACC
REV-24:
CAAGCAGAAGACGGCATACGAGAT**ATCAGT**CTCGGCATTCCTGCTGAACC
REV-25:
CAAGCAGAAGACGGCATACGAGAT**GCTCAT**CTCGGCATTCCTGCTGAACC
REV-26:
CAAGCAGAAGACGGCATACGAGAT**AGGAAT**CTCGGCATTCCTGCTGAACC
REV-27:
CAAGCAGAAGACGGCATACGAGAT**CTTTTG**CTCGGCATTCCTGCTGAACC
REV-28:
CAAGCAGAAGACGGCATACGAGAT**TAGTTG**CTCGGCATTCCTGCTGAACC
REV-29:
CAAGCAGAAGACGGCATACGAGAT**CCGGTG**CTCGGCATTCCTGCTGAACC
REV-30:
CAAGCAGAAGACGGCATACGAGAT**ATCGTG**CTCGGCATTCCTGCTGAACC

**Blocking oligos for inhibition of formation of daisy chain PCR templates during DNA/RNA hybridization**

Blocking oligo-1: 5’-
AATGATACGGCGACCACCGAGATCTACACTCTTTCCCTACACGACGCTCTTCCGA
TC/3InvdT/-3’

Blocking oligo-2: 5’-CTCGGCATTCCTGCTGAACCGCTCTTCCGATCT/3InvdT/-3’

**Sequencing primers.**
Read-1-Seq-primer: ACACTCTTTCCCTACACGACGCTCTTCCGATCT
Read-2-Seq-primer: CTCGGCATTCCTGCTGAACCGCTCTTCCGATCT
Index-Read-Seq-primer: AGATCGGAAGAGCGGTTCAGCAGGAATGCCGAG

**Primers for evaluation of enrichment efficiency of L1 and Alu capture libraries.**

Alu
FW: TCCTGCCTCAGCCTCCCAAG
REV: GTCAGGAGATCGAGACCATCCC

L1 5’-UTR
FW: CGGTGATTTCTGCATTTCCATC
REV: TTCCCAGGTGAGGCAATGCCT
