## Supplementary Document 2 for "Recombination of repeat elements generates somatic complexity in human genomes"

### Supplementary Document-2: Oligonucleotide sequences for synthesis of capture probes

| Probe_Name | Probe_sequence |
|---|---|
| L1HS-1 | ATCGCACCAGCGTGTAGGAGCCAAGATGGCCGAATAGGAACAGCTCCGGTCTACAGCTCCCAGCGTGAGCGACGCCACTGCGGCTCCTCA |
| L1HS-2 | ATCGCACCAGCGTGTGAGCCAAGATGGCCGAATAGGAACAGCTCCGGTCTACAGCTCCCAGCGTGAGCGACGCAGCACTGCGGCTCCTCA |
| L1HS-3 | ATCGCACCAGCGTGTAGCCAAGATGGCCGAATAGGAACAGCTCCGGTCTACAGCTCCCAGCGTGAGCGACGCAGACACTGCGGCTCCTCA |
| L1HS-4 | ATCGCACCAGCGTGTCCAAGATGGCCGAATAGGAACAGCTCCGGTCTACAGCTCCCAGCGTGAGCGACGCAGAAGCACTGCGGCTCCTCA |
| L1HS-5 | ATCGCACCAGCGTGTAAGATGGCCGAATAGGAACAGCTCCGGTCTACAGCTCCCAGCGTGAGCGACGCAGAAGACCACTGCGGCTCCTCA |
| L1HS-6 | ATCGCACCAGCGTGTAGATGGCCGAATAGGAACAGCTCCGGTCTACAGCTCCCAGCGTGAGCGACGCAGAAGACGCACTGCGGCTCCTCA |
| L1HS-7 | ATCGCACCAGCGTGTGATGGCCGAATAGGAACAGCTCCGGTCTACAGCTCCCAGCGTGAGCGACGCAGAAGACGGCACTGCGGCTCCTCA |
| L1HS-8 | ATCGCACCAGCGTGTTGGCCGAATAGGAACAGCTCCGGTCTACAGCTCCCAGCGTGAGCGACGCAGAAGACGGTGCACTGCGGCTCCTCA |
| L1HS-9 | ATCGCACCAGCGTGTCGAATAGGAACAGCTCCGGTCTACAGCTCCCAGCGTGAGCGACGCAGAAGACGGTGATTTCACTGCGGCTCCTCA |
| L1HS-10 | ATCGCACCAGCGTGTAATAGGAACAGCTCCGGTCTACAGCTCCCAGCGTGAGCGACGCAGAAGACGGTGATTTCTCACTGCGGCTCCTCA |
| L1HS-11 | ATCGCACCAGCGTGTTAGGAACAGCTCCGGTCTACAGCTCCCAGCGTGAGCGACGCAGAAGACGGTGATTTCTGCCACTGCGGCTCCTCA |
| L1HS-12 | ATCGCACCAGCGTGTGGAACAGCTCCGGTCTACAGCTCCCAGCGTGAGCGACGCAGAAGACGGTGATTTCTGCATCACTGCGGCTCCTCA |
| L1HS-13 | ATCGCACCAGCGTGTAACAGCTCCGGTCTACAGCTCCCAGCGTGAGCGACGCAGAAGACGGTGATTTCTGCATTTCACTGCGGCTCCTCA |
| L1HS-14 | ATCGCACCAGCGTGTCAGCTCCGGTCTACAGCTCCCAGCGTGAGCGACGCAGAAGACGGTGATTTCTGCATTTCCCACTGCGGCTCCTCA |
| L1HS-15 | ATCGCACCAGCGTGTGCTCCGGTCTACAGCTCCCAGCGTGAGCGACGCAGAAGACGGTGATTTCTGCATTTCCATCACTGCGGCTCCTCA |
| L1HS-16 | ATCGCACCAGCGTGTTCCGGTCTACAGCTCCCAGCGTGAGCGACGCAGAAGACGGTGATTTCTGCATTTCCATCTCACTGCGGCTCCTCA |
| L1HS-17 | ATCGCACCAGCGTGTCGGTCTACAGCTCCCAGCGTGAGCGACGCAGAAGACGGTGATTTCTGCATTTCCATCTGACACTGCGGCTCCTCA |
| L1HS-18 | ATCGCACCAGCGTGTGTCTACAGCTCCCAGCGTGAGCGACGCAGAAGACGGTGATTTCTGCATTTCCATCTGAGGCACTGCGGCTCCTCA |
| L1HS-19 | ATCGCACCAGCGTGTCTACAGCTCCCAGCGTGAGCGACGCAGAAGACGGTGATTTCTGCATTTCCATCTGAGGTACACTGCGGCTCCTCA |
| L1HS-20 | ATCGCACCAGCGTGTACAGCTCCCAGCGTGAGCGACGCAGAAGACGGTGATTTCTGCATTTCCATCTGAGGTACCCACTGCGGCTCCTCA |
| L1HS-21 | ATCGCACCAGCGTGTAGCTCCCAGCGTGAGCGACGCAGAAGACGGTGATTTCTGCATTTCCATCTGAGGTACCGGCACTGCGGCTCCTCA |
| L1HS-22 | ATCGCACCAGCGTGTCTCCCAGCGTGAGCGACGCAGAAGACGGTGATTTCTGCATTTCCATCTGAGGTACCGGGTCACTGCGGCTCCTCA |
| L1HS-23 | ATCGCACCAGCGTGTCCCAGCGTGAGCGACGCAGAAGACGGTGATTTCTGCATTTCCATCTGAGGTACCGGGTTCCACTGCGGCTCCTCA |
| L1HS-24 | ATCGCACCAGCGTGTCAGCGTGAGCGACGCAGAAGACGGTGATTTCTGCATTTCCATCTGAGGTACCGGGTTCATCACTGCGGCTCCTCA |
| L1HS-25 | ATCGCACCAGCGTGTGCGTGAGCGACGCAGAAGACGGTGATTTCTGCATTTCCATCTGAGGTACCGGGTTCATCTCACTGCGGCTCCTCA |
| L1HS-26 | ATCGCACCAGCGTGTGTGAGCGACGCAGAAGACGGTGATTTCTGCATTTCCATCTGAGGTACCGGGTTCATCTCACACTGCGGCTCCTCA |
| L1HS-27 | ATCGCACCAGCGTGTGAGCGACGCAGAAGACGGTGATTTCTGCATTTCCATCTGAGGTACCGGGTTCATCTCACTCACTGCGGCTCCTCA |
| L1HS-28 | ATCGCACCAGCGTGTGCGACGCAGAAGACGGTGATTTCTGCATTTCCATCTGAGGTACCGGGTTCATCTCACTAGCACTGCGGCTCCTCA |
| L1HS-29 | ATCGCACCAGCGTGTGACGCAGAAGACGGTGATTTCTGCATTTCCATCTGAGGTACCGGGTTCATCTCACTAGGGCACTGCGGCTCCTCA |
| L1HS-30 | ATCGCACCAGCGTGTCGCAGAAGACGGTGATTTCTGCATTTCCATCTGAGGTACCGGGTTCATCTCACTAGGGAGCACTGCGGCTCCTCA |
| L1HS-31 | ATCGCACCAGCGTGTCAGAAGACGGTGATTTCTGCATTTCCATCTGAGGTACCGGGTTCATCTCACTAGGGAGTGCACTGCGGCTCCTCA |
| L1HS-32 | ATCGCACCAGCGTGTGAAGACGGTGATTTCTGCATTTCCATCTGAGGTACCGGGTTCATCTCACTAGGGAGTGCCCACTGCGGCTCCTCA |
| L1HS-33 | ATCGCACCAGCGTGTAGACGGTGATTTCTGCATTTCCATCTGAGGTACCGGGTTCATCTCACTAGGGAGTGCCAGCACTGCGGCTCCTCA |
| L1HS-34 | ATCGCACCAGCGTGTACGGTGATTTCTGCATTTCCATCTGAGGTACCGGGTTCATCTCACTAGGGAGTGCCAGACCACTGCGGCTCCTCA |
| L1HS-35 | ATCGCACCAGCGTGTGGTGATTTCTGCATTTCCATCTGAGGTACCGGGTTCATCTCACTAGGGAGTGCCAGACAGCACTGCGGCTCCTCA |
| L1HS-36 | ATCGCACCAGCGTGTTGATTTCTGCATTTCCATCTGAGGTACCGGGTTCATCTCACTAGGGAGTGCCAGACAGTGCACTGCGGCTCCTCA |
| L1HS-37 | ATCGCACCAGCGTGTATTTCTGCATTTCCATCTGAGGTACCGGGTTCATCTCACTAGGGAGTGCCAGACAGTGGCACTGCGGCTCCTCA |
| L1HS-38 | ATCGCACCAGCGTGTTTCTGCATTTCCATCTGAGGTACCGGGTTCATCTCACTAGGGAGTGCCAGACAGTGGGCGCACTGCGGCTCCTCA |
| L1HS-39 | ATCGCACCAGCGTGTCTGCATTTCCATCTGAGGTACCGGGTTCATCTCACTAGGGAGTGCCAGACAGTGGGCGCACACTGCGGCTCCTCA |
| L1HS-40 | ATCGCACCAGCGTGTGCATTTCCATCTGAGGTACCGGGTTCATCTCACTAGGGAGTGCCAGACAGTGGGCGCAGGCACTGCGGCTCCTCA |
| L1HS-41 | ATCGCACCAGCGTGTATTTCCATCTGAGGTACCGGGTTCATCTCACTAGGGAGTGCCAGACAGTGGGCGCAGGCCCACTGCGGCTCCTCA |
| L1HS-42 | ATCGCACCAGCGTGTTTCCATCTGAGGTACCGGGTTCATCTCACTAGGGAGTGCCAGACAGTGGGCGCAGGCCAGCACTGCGGCTCCTCA |
| L1HS-43 | ATCGCACCAGCGTGTCGAAGCAGGGCGAGGCATTGCCTCACCTGGGAAGCGCAAGGGGTCAGGGAGTTCCCTTTCCACTGCGGCTCCTCA |
| L1HS-44 | ATCGCACCAGCGTGTAAGGGGAATATCACACTCTGGGGACTGTGGTGGGGTCGGGGGAGGGGGGAGGGATAGCATCACTGCGGCTCCTCA |
| L1HS-45 | ATCGCACCAGCGTGTGGGGAATATCACACTCTGGGGACTGTGGTGGGGTCGGGGGAGGGGGGAGGGATAGCATTGCACTGCGGCTCCTCA |
| L1HS-46 | ATCGCACCAGCGTGTGGAATATCACACTCTGGGGACTGTGGTGGGGTCGGGGGAGGGGGGAGGGATAGCATTGGGCACTGCGGCTCCTCA |
| L1HS-47 | ATCGCACCAGCGTGTAATATCACACTCTGGGGACTGTGGTGGGGTCGGGGGAGGGGGGAGGGATAGCATTGGGAGCACTGCGGCTCCTCA |
| L1HS-48 | ATCGCACCAGCGTGTTATCACACTCTGGGGACTGTGGTGGGGTCGGGGGAGGGGGGAGGGATAGCATTGGGAGATCACTGCGGCTCCTCA |
| L1HS-49 | ATCGCACCAGCGTGTTCACACTCTGGGGACTGTGGTGGGGTCGGGGGAGGGGGGAGGGATAGCATTGGGAGATATCACTGCGGCTCCTCA |
| L1HS-50 | ATCGCACCAGCGTGTACACTCTGGGGACTGTGGTGGGGTCGGGGGAGGGGGGAGGGATAGCATTGGGAGATATACCACTGCGGCTCCTCA |
| L1HS-51 | ATCGCACCAGCGTGTACTCTGGGGACTGTGGTGGGGTCGGGGGAGGGGGGAGGGATAGCATTGGGAGATATACCTCACTGCGGCTCCTCA |
| L1HS-52 | ATCGCACCAGCGTGTTGGGGACTGTGGTGGGGTCGGGGGAGGGGGGAGGGATAGCATTGGGAGATATACCTAATGCACTGCGGCTCCTCA |
| L1HS-53 | ATCGCACCAGCGTGTGGGACTGTGGTGGGGTCGGGGGAGGGGGGAGGGATAGCATTGGGAGATATACCTAATGCTCACTGCGGCTCCTCA |
| L1HS-54 | ATCGCACCAGCGTGTGACTGTGGTGGGGTCGGGGGAGGGGGGAGGGATAGCATTGGGAGATATACCTAATGCTAGCACTGCGGCTCCTCA |
| L1HS-55 | ATCGCACCAGCGTGTCTGTGGTGGGGTCGGGGGAGGGGGGAGGGATAGCATTGGGAGATATACCTAATGCTAGATCACTGCGGCTCCTCA |
| L1HS-56 | ATCGCACCAGCGTGTGTGGTGGGGTCGGGGGAGGGGGGAGGGATAGCATTGGGAGATATACCTAATGCTAGATGACACTGCGGCTCCTCA |
| L1HS-57 | ATCGCACCAGCGTGTGGTGGGGTCGGGGGAGGGGGGAGGGATAGCATTGGGAGATATACCTAATGCTAGATGACACACTGCGGCTCCTCA |
| L1HS-58 | ATCGCACCAGCGTGTTGGGGTCGGGGGAGGGGGGAGGGATAGCATTGGGAGATATACCTAATGCTAGATGACACACACTGCGGCTCCTCA |
| L1HS-59 | ATCGCACCAGCGTGTGGGTCGGGGGAGGGGGGAGGGATAGCATTGGGAGATATACCTAATGCTAGATGACACATTCACTGCGGCTCCTCA |
| L1HS-60 | ATCGCACCAGCGTGTGTCGGGGGAGGGGGGAGGGATAGCATTGGGAGATATACCTAATGCTAGATGACACATTAGCACTGCGGCTCCTCA |
| L1HS-61 | ATCGCACCAGCGTGTCGGGGGAGGGGGGAGGGATAGCATTGGGAGATATACCTAATGCTAGATGACACATTAGTGCACTGCGGCTCCTCA |
| L1HS-62 | ATCGCACCAGCGTGTGGGGAGGGGGGAGGGATAGCATTGGGAGATATACCTAATGCTAGATGACACATTAGTGGGCACTGCGGCTCCTCA |
| L1HS-63 | ATCGCACCAGCGTGTGGAGGGGGGAGGGATAGCATTGGGAGATATACCTAATGCTAGATGACACATTAGTGGGTGCACTGCGGCTCCTCA |
| L1HS-64 | ATCGCACCAGCGTGTAGGGGGGAGGGATAGCATTGGGAGATATACCTAATGCTAGATGACACATTAGTGGGTGCACACTGCGGCTCCTCA |
| L1HS-65 | ATCGCACCAGCGTGTGGGGGAGGGATAGCATTGGGAGATATACCTAATGCTAGATGACACATTAGTGGGTGCAGCCACTGCGGCTCCTCA |
| L1HS-66 | ATCGCACCAGCGTGTGGGAGGGATAGCATTGGGAGATATACCTAATGCTAGATGACACATTAGTGGGTGCAGCGCCACTGCGGCTCCTCA |
| L1HS-67 | ATCGCACCAGCGTGTGAGGGATAGCATTGGGAGATATACCTAATGCTAGATGACACATTAGTGGGTGCAGCGCACCACTGCGGCTCCTCA |
| L1HS-68 | ATCGCACCAGCGTGTGGGATAGCATTGGGAGATATACCTAATGCTAGATGACACATTAGTGGGTGCAGCGCACCACACTGCGGCTCCTCA |

L1HS-69 ATCGCACCAGCGTGTGATAGCATTGGGAGATATACCTAATGCTAGATGACACATTAGTGGGTGCAGCGCACCAGCCACTGCGGCTCCTCA
L1HS-70 ATCGCACCAGCGTGTTAGCATTGGGAGATATACCTAATGCTAGATGACACATTAGTGGGTGCAGCGCACCAGCATCACTGCGGCTCCTCA
L1HS-71 ATCGCACCAGCGTGTGCATTGGGAGATATACCTAATGCTAGATGACACATTAGTGGGTGCAGCGCACCAGCATGGCACTGCGGCTCCTCA
L1HS-72 ATCGCACCAGCGTGTATTGGGAGATATACCTAATGCTAGATGACACATTAGTGGGTGCAGCGCACCAGCATGGCACACTGCGGCTCCTCA
L1HS-73 ATCGCACCAGCGTGTTGGGAGATATACCTAATGCTAGATGACACATTAGTGGGTGCAGCGCACCAGCATGGCACACACTGCGGCTCCTCA
L1HS-74 ATCGCACCAGCGTGTGGAGATATACCTAATGCTAGATGACACATTAGTGGGTGCAGCGCACCAGCATGGCACATGCACTGCGGCTCCTCA
L1HS-75 ATCGCACCAGCGTGTAGATATACCTAATGCTAGATGACACATTAGTGGGTGCAGCGCACCAGCATGGCACATGTACACTGCGGCTCCTCA
L1HS-76 ATCGCACCAGCGTGTATATACCTAATGCTAGATGACACATTAGTGGGTGCAGCGCACCAGCATGGCACATGTATACACTGCGGCTCCTCA
L1HS-77 ATCGCACCAGCGTGTATACCTAATGCTAGATGACACATTAGTGGGTGCAGCGCACCAGCATGGCACATGTATACACACTGCGGCTCCTCA
L1HS-78 ATCGCACCAGCGTGTACCTAATGCTAGATGACACATTAGTGGGTGCAGCGCACCAGCATGGCACATGTATACATACACTGCGGCTCCTCA
L1HS-79 ATCGCACCAGCGTGTCTAATGCTAGATGACACATTAGTGGGTGCAGCGCACCAGCATGGCACATGTATACATATGCACTGCGGCTCCTCA
L1HS-80 ATCGCACCAGCGTGTAATGCTAGATGACACATTAGTGGGTGCAGCGCACCAGCATGGCACATGTATACATATGTACACTGCGGCTCCTCA
L1HS-81 ATCGCACCAGCGTGTTGCTAGATGACACATTAGTGGGTGCAGCGCACCAGCATGGCACATGTATACATATGTAACCACTGCGGCTCCTCA
L1HS-82 ATCGCACCAGCGTGTCTAGATGACACATTAGTGGGTGCAGCGCACCAGCATGGCACATGTATACATATGTAACTACACTGCGGCTCCTCA
L1HS-83 ATCGCACCAGCGTGTAGATGACACATTAGTGGGTGCAGCGCACCAGCATGGCACATGTATACATATGTAACTAACCACTGCGGCTCCTCA
L1HS-84 ATCGCACCAGCGTGTATGACACATTAGTGGGTGCAGCGCACCAGCATGGCACATGTATACATATGTAACTAACCTCACTGCGGCTCCTCA
L1HS-85 ATCGCACCAGCGTGTGACACATTAGTGGGTGCAGCGCACCAGCATGGCACATGTATACATATGTAACTAACCTGCCACTGCGGCTCCTCA
L1HS-86 ATCGCACCAGCGTGTCACATTAGTGGGTGCAGCGCACCAGCATGGCACATGTATACATATGTAACTAACCTGCACCACTGCGGCTCCTCA
L1HS-87 ATCGCACCAGCGTGTCATTAGTGGGTGCAGCGCACCAGCATGGCACATGTATACATATGTAACTAACCTGCACAACACTGCGGCTCCTCA
L1HS-88 ATCGCACCAGCGTGTTTAGTGGGTGCAGCGCACCAGCATGGCACATGTATACATATGTAACTAACCTGCACAATGCACTGCGGCTCCTCA
L1HS-89 ATCGCACCAGCGTGTAGTGGGTGCAGCGCACCAGCATGGCACATGTATACATATGTAACTAACCTGCACAATGTGCACTGCGGCTCCTCA
L1HS-90 ATCGCACCAGCGTGTTGGGTGCAGCGCACCAGCATGGCACATGTATACATATGTAACTAACCTGCACAATGTGCACACTGCGGCTCCTCA
L1HS-91 ATCGCACCAGCGTGTGGTGCAGCGCACCAGCATGGCACATGTATACATATGTAACTAACCTGCACAATGTGCACACACTGCGGCTCCTCA
L1HS-92 ATCGCACCAGCGTGTTGCAGCGCACCAGCATGGCACATGTATACATATGTAACTAACCTGCACAATGTGCACATGCACTGCGGCTCCTCA
L1HS-93 ATCGCACCAGCGTGTCAGCGCACCAGCATGGCACATGTATACATATGTAACTAACCTGCACAATGTGCACATGTACACTGCGGCTCCTCA
L1HS-94 ATCGCACCAGCGTGTGCGCACCAGCATGGCACATGTATACATATGTAACTAACCTGCACAATGTGCACATGTACCCACTGCGGCTCCTCA
L1HS-95 ATCGCACCAGCGTGTGCACCAGCATGGCACATGTATACATATGTAACTAACCTGCACAATGTGCACATGTACCCTCACTGCGGCTCCTCA
L1HS-96 ATCGCACCAGCGTGTACCAGCATGGCACATGTATACATATGTAACTAACCTGCACAATGTGCACATGTACCCTAACACTGCGGCTCCTCA

AluYa5-1 ATCGCACCAGCGTGTGGAAGAAGGGGGCCGGGCGCGGTGGCTCACGCCTGTAATCCCAGCACTTTGGGAGGCCGACACTGCGGCTCCTCA
AluYa5-2 ATCGCACCAGCGTGTAAGAAGGGGGCCGGGCGCGGTGGCTCACGCCTGTAATCCCAGCACTTTGGGAGGCCGAGGCACTGCGGCTCCTCA
AluYa5-3 ATCGCACCAGCGTGTAAGGGGGCCGGGCGCGGTGGCTCACGCCTGTAATCCCAGCACTTTGGGAGGCCGAGGCGGCACTGCGGCTCCTCA
AluYa5-4 ATCGCACCAGCGTGTAGGGGGCCGGGCGCGGTGGCTCACGCCTGTAATCCCAGCACTTTGGGAGGCCGAGGCGGGCACTGCGGCTCCTCA
AluYa5-5 ATCGCACCAGCGTGTGGGGCCGGGCGCGGTGGCTCACGCCTGTAATCCCAGCACTTTGGGAGGCCGAGGCGGGCGCACTGCGGCTCCTCA
AluYa5-6 ATCGCACCAGCGTGTGGCCGGGCGCGGTGGCTCACGCCTGTAATCCCAGCACTTTGGGAGGCCGAGGCGGGCGGACACTGCGGCTCCTCA
AluYa5-7 ATCGCACCAGCGTGTCCGGGCGCGGTGGCTCACGCCTGTAATCCCAGCACTTTGGGAGGCCGAGGCGGGCGGATCCACTGCGGCTCCTCA
AluYa5-8 ATCGCACCAGCGTGTGGGCGCGGTGGCTCACGCCTGTAATCCCAGCACTTTGGGAGGCCGAGGCGGGCGGATCACCACTGCGGCTCCTCA
AluYa5-9 ATCGCACCAGCGTGTGCGCGGTGGCTCACGCCTGTAATCCCAGCACTTTGGGAGGCCGAGGCGGGCGGATCACGACACTGCGGCTCCTCA
AluYa5-10 ATCGCACCAGCGTGTGCGGTGGCTCACGCCTGTAATCCCAGCACTTTGGGAGGCCGAGGCGGGCGGATCACGAGGCACTGCGGCTCCTCA
AluYa5-11 ATCGCACCAGCGTGTGGTGGCTCACGCCTGTAATCCCAGCACTTTGGGAGGCCGAGGCGGGCGGATCACGAGGTCCACTGCGGCTCCTCA
AluYa5-12 ATCGCACCAGCGTGTTGGCTCACGCCTGTAATCCCAGCACTTTGGGAGGCCGAGGCGGGCGGATCACGAGGTCAGCACTGCGGCTCCTCA
AluYa5-13 ATCGCACCAGCGTGTGCTCACGCCTGTAATCCCAGCACTTTGGGAGGCCGAGGCGGGCGGATCACGAGGTCAGGACACTGCGGCTCCTCA
AluYa5-14 ATCGCACCAGCGTGTTCACGCCTGTAATCCCAGCACTTTGGGAGGCCGAGGCGGGCGGATCACGAGGTCAGGAGACACTGCGGCTCCTCA
AluYa5-15 ATCGCACCAGCGTGTACGCCTGTAATCCCAGCACTTTGGGAGGCCGAGGCGGGCGGATCACGAGGTCAGGAGATCCACTGCGGCTCCTCA
AluYa5-16 ATCGCACCAGCGTGTGCCTGTAATCCCAGCACTTTGGGAGGCCGAGGCGGGCGGATCACGAGGTCAGGAGATCGACACTGCGGCTCCTCA
AluYa5-17 ATCGCACCAGCGTGTCTGTAATCCCAGCACTTTGGGAGGCCGAGGCGGGCGGATCACGAGGTCAGGAGATCGAGACACTGCGGCTCCTCA
AluYa5-18 ATCGCACCAGCGTGTGTAATCCCAGCACTTTGGGAGGCCGAGGCGGGCGGATCACGAGGTCAGGAGATCGAGACCCACTGCGGCTCCTCA
AluYa5-19 ATCGCACCAGCGTGTAATCCCAGCACTTTGGGAGGCCGAGGCGGGCGGATCACGAGGTCAGGAGATCGAGACCATCACTGCGGCTCCTCA
AluYa5-20 ATCGCACCAGCGTGTTCCCAGCACTTTGGGAGGCCGAGGCGGGCGGATCACGAGGTCAGGAGATCGAGACCATCCCACTGCGGCTCCTCA
AluYa5-21 ATCGCACCAGCGTGTAGTCCCAGCTACTTGGGAGGCTGAGGCAGGAGAATGGCGTGAACCCGGGAGGCGGAGCTTCACTGCGGCTCCTCA
AluYa5-22 ATCGCACCAGCGTGTTCCCAGCTACTTGGGAGGCTGAGGCAGGAGAATGGCGTGAACCCGGGAGGCGGAGCTTGCCACTGCGGCTCCTCA
AluYa5-23 ATCGCACCAGCGTGTCCAGCTACTTGGGAGGCTGAGGCAGGAGAATGGCGTGAACCCGGGAGGCGGAGCTTGCAGCACTGCGGCTCCTCA
AluYa5-24 ATCGCACCAGCGTGTAGCTACTTGGGAGGCTGAGGCAGGAGAATGGCGTGAACCCGGGAGGCGGAGCTTGCAGTGCACTGCGGCTCCTCA
AluYa5-25 ATCGCACCAGCGTGTCTACTTGGGAGGCTGAGGCAGGAGAATGGCGTGAACCCGGGAGGCGGAGCTTGCAGTGAGCACTGCGGCTCCTCA
AluYa5-26 ATCGCACCAGCGTGTACTTGGGAGGCTGAGGCAGGAGAATGGCGTGAACCCGGGAGGCGGAGCTTGCAGTGAGCCCACTGCGGCTCCTCA
AluYa5-27 ATCGCACCAGCGTGTTTGGGAGGCTGAGGCAGGAGAATGGCGTGAACCCGGGAGGCGGAGCTTGCAGTGAGCCGACACTGCGGCTCCTCA
AluYa5-28 ATCGCACCAGCGTGTGGGAGGCTGAGGCAGGAGAATGGCGTGAACCCGGGAGGCGGAGCTTGCAGTGAGCCGAGACACTGCGGCTCCTCA
AluYa5-29 ATCGCACCAGCGTGTGAGGCTGAGGCAGGAGAATGGCGTGAACCCGGGAGGCGGAGCTTGCAGTGAGCCGAGATCCACTGCGGCTCCTCA
AluYa5-30 ATCGCACCAGCGTGTGGCTGAGGCAGGAGAATGGCGTGAACCCGGGAGGCGGAGCTTGCAGTGAGCCGAGATCCCACTGCGGCTCCTCA
AluYa5-31 ATCGCACCAGCGTGTCTGAGGCAGGAGAATGGCGTGAACCCGGGAGGCGGAGCTTGCAGTGAGCCGAGATCCCGCCACTGCGGCTCCTCA
AluYa5-32 ATCGCACCAGCGTGTGAGGCAGGAGAATGGCGTGAACCCGGGAGGCGGAGCTTGCAGTGAGCCGAGATCCCGCCACACTGCGGCTCCTCA
AluYa5-33 ATCGCACCAGCGTGTGGCAGGAGAATGGCGTGAACCCGGGAGGCGGAGCTTGCAGTGAGCCGAGATCCCGCCACTCACTGCGGCTCCTCA
AluYa5-34 ATCGCACCAGCGTGTCAGGAGAATGGCGTGAACCCGGGAGGCGGAGCTTGCAGTGAGCCGAGATCCCGCCACTGCCACTGCGGCTCCTCA
AluYa5-35 ATCGCACCAGCGTGTGGAGAATGGCGTGAACCCGGGAGGCGGAGCTTGCAGTGAGCCGAGATCCCGCCACTGCACCACTGCGGCTCCTCA
AluYa5-36 ATCGCACCAGCGTGTAGAATGGCGTGAACCCGGGAGGCGGAGCTTGCAGTGAGCCGAGATCCCGCCACTGCACTCCACTGCGGCTCCTCA
AluYa5-37 ATCGCACCAGCGTGTAATGGCGTGAACCCGGGAGGCGGAGCTTGCAGTGAGCCGAGATCCCGCCACTGCACTCCACACTGCGGCTCCTCA
AluYa5-38 ATCGCACCAGCGTGTTGGCGTGAACCCGGGAGGCGGAGCTTGCAGTGAGCCGAGATCCCGCCACTGCACTCCAGCCACTGCGGCTCCTCA
AluYa5-39 ATCGCACCAGCGTGTGCGTGAACCCGGGAGGCGGAGCTTGCAGTGAGCCGAGATCCCGCCACTGCACTCCAGCCTCACTGCGGCTCCTCA
AluYa5-40 ATCGCACCAGCGTGTGTGAACCCGGGAGGCGGAGCTTGCAGTGAGCCGAGATCCCGCCACTGCACTCCAGCCTGGCACTGCGGCTCCTCA
AluYb9-1 ATCGCACCAGCGTGTGTAGGCAATGGGCCGGGCGCGGTGGCTCACGCCTGTAATCCCAGCACTTTGGGAGGCCGACACTGCGGCTCCTCA
AluYb9-2 ATCGCACCAGCGTGTAGGCAATGGGCCGGGCGCGGTGGCTCACGCCTGTAATCCCAGCACTTTGGGAGGCCGAGGCACTGCGGCTCCTCA

AluYb9-3 ATCGCACCAGCGTGTGCAATGGGCCGGGCGCGGTGGCTCACGCCTGTAATCCCAGCACTTTGGGAGGCCGAGGCGCACTGCGGCTCCTCA
AluYb9-4 ATCGCACCAGCGTGTAATGGGCCGGGCGCGGTGGCTCACGCCTGTAATCCCAGCACTTTGGGAGGCCGAGGCGGGCACTGCGGCTCCTCA
AluYb9-5 ATCGCACCAGCGTGTTGGGCCGGGCGCGGTGGCTCACGCCTGTAATCCCAGCACTTTGGGAGGCCGAGGCGGGTGCACTGCGGCTCCTCA
AluYb9-6 ATCGCACCAGCGTGTGGCCGGGCGCGGTGGCTCACGCCTGTAATCCCAGCACTTTGGGAGGCCGAGGCGGGTGGACACTGCGGCTCCTCA
AluYb9-7 ATCGCACCAGCGTGTCCGGGCGCGGTGGCTCACGCCTGTAATCCCAGCACTTTGGGAGGCCGAGGCGGGTGGATCCACTGCGGCTCCTCA
AluYb9-8 ATCGCACCAGCGTGTGGGCGCGGTGGCTCACGCCTGTAATCCCAGCACTTTGGGAGGCCGAGGCGGGTGGATCATCACTGCGGCTCCTCA
AluYb9-9 ATCGCACCAGCGTGTGCGCGGTGGCTCACGCCTGTAATCCCAGCACTTTGGGAGGCCGAGGCGGGTGGATCATGACACTGCGGCTCCTCA
AluYb9-10 ATCGCACCAGCGTGTGCGGTGGCTCACGCCTGTAATCCCAGCACTTTGGGAGGCCGAGGCGGGTGGATCATGAGGCACTGCGGCTCCTCA
AluYb9-11 ATCGCACCAGCGTGTGGTGGCTCACGCCTGTAATCCCAGCACTTTGGGAGGCCGAGGCGGGTGGATCATGAGGTCCACTGCGGCTCCTCA
AluYb9-12 ATCGCACCAGCGTGTTGGCTCACGCCTGTAATCCCAGCACTTTGGGAGGCCGAGGCGGGTGGATCATGAGGTCAGCACTGCGGCTCCTCA
AluYb9-13 ATCGCACCAGCGTGTGCTCACGCCTGTAATCCCAGCACTTTGGGAGGCCGAGGCGGGTGGATCATGAGGTCAGGACACTGCGGCTCCTCA
AluYb9-14 ATCGCACCAGCGTGTTCACGCCTGTAATCCCAGCACTTTGGGAGGCCGAGGCGGGTGGATCATGAGGTCAGGAGACACTGCGGCTCCTCA
AluYb9-15 ATCGCACCAGCGTGTACGCCTGTAATCCCAGCACTTTGGGAGGCCGAGGCGGGTGGATCATGAGGTCAGGAGATCCACTGCGGCTCCTCA
AluYb9-16 ATCGCACCAGCGTGTGCCTGTAATCCCAGCACTTTGGGAGGCCGAGGCGGGTGGATCATGAGGTCAGGAGATCGACACTGCGGCTCCTCA
AluYb9-17 ATCGCACCAGCGTGTCTGTAATCCCAGCACTTTGGGAGGCCGAGGCGGGTGGATCATGAGGTCAGGAGATCGAGACACTGCGGCTCCTCA
AluYb9-18 ATCGCACCAGCGTGTGTAATCCCAGCACTTTGGGAGGCCGAGGCGGGTGGATCATGAGGTCAGGAGATCGAGACCCACTGCGGCTCCTCA
AluYb9-19 ATCGCACCAGCGTGTAATCCCAGCACTTTGGGAGGCCGAGGCGGGTGGATCATGAGGTCAGGAGATCGAGACCATCACTGCGGCTCCTCA
AluYb9-20 ATCGCACCAGCGTGTTCCCAGCACTTTGGGAGGCCGAGGCGGGTGGATCATGAGGTCAGGAGATCGAGACCATCCCACTGCGGCTCCTCA
AluYb9-21 ATCGCACCAGCGTGTGCTACTGGGGAGGCTGAGGCAGGAGAATGGCGTGAACCCGGGAAGCGGAGCTTGCAGTGACACTGCGGCTCCTCA
AluYb9-22 ATCGCACCAGCGTGTTACTGGGGAGGCTGAGGCAGGAGAATGGCGTGAACCCGGGAAGCGGAGCTTGCAGTGAGCCACTGCGGCTCCTCA
AluYb9-23 ATCGCACCAGCGTGTCTGGGGAGGCTGAGGCAGGAGAATGGCGTGAACCCGGGAAGCGGAGCTTGCAGTGAGCCGCACTGCGGCTCCTCA
AluYb9-24 ATCGCACCAGCGTGTGGGGAGGCTGAGGCAGGAGAATGGCGTGAACCCGGGAAGCGGAGCTTGCAGTGAGCCGAGCACTGCGGCTCCTCA
AluYb9-25 ATCGCACCAGCGTGTGGAGGCTGAGGCAGGAGAATGGCGTGAACCCGGGAAGCGGAGCTTGCAGTGAGCCGAGATCACTGCGGCTCCTCA
AluYb9-26 ATCGCACCAGCGTGTAGGCTGAGGCAGGAGAATGGCGTGAACCCGGGAAGCGGAGCTTGCAGTGAGCCGAGATTGCACTGCGGCTCCTCA
AluYb9-27 ATCGCACCAGCGTGTGCTGAGGCAGGAGAATGGCGTGAACCCGGGAAGCGGAGCTTGCAGTGAGCCGAGATTGCGCACTGCGGCTCCTCA
AluYb9-28 ATCGCACCAGCGTGTTGAGGCAGGAGAATGGCGTGAACCCGGGAAGCGGAGCTTGCAGTGAGCCGAGATTGCGCCCACTGCGGCTCCTCA
AluYb9-29 ATCGCACCAGCGTGTAGGCAGGAGAATGGCGTGAACCCGGGAAGCGGAGCTTGCAGTGAGCCGAGATTGCGCCACCACTGCGGCTCCTCA
AluYb9-30 ATCGCACCAGCGTGTGCAGGAGAATGGCGTGAACCCGGGAAGCGGAGCTTGCAGTGAGCCGAGATTGCGCCACTGCACTGCGGCTCCTCA
AluYb9-31 ATCGCACCAGCGTGTAGGAGAATGGCGTGAACCCGGGAAGCGGAGCTTGCAGTGAGCCGAGATTGCGCCACTGCACACTGCGGCTCCTCA
AluYb9-32 ATCGCACCAGCGTGTGAGAATGGCGTGAACCCGGGAAGCGGAGCTTGCAGTGAGCCGAGATTGCGCCACTGCAGTCACTGCGGCTCCTCA
AluYb9-33 ATCGCACCAGCGTGTGAATGGCGTGAACCCGGGAAGCGGAGCTTGCAGTGAGCCGAGATTGCGCCACTGCAGTCCCACTGCGGCTCCTCA
AluYb9-34 ATCGCACCAGCGTGTATGGCGTGAACCCGGGAAGCGGAGCTTGCAGTGAGCCGAGATTGCGCCACTGCAGTCCGCCACTGCGGCTCCTCA
AluYb9-35 ATCGCACCAGCGTGTGGCGTGAACCCGGGAAGCGGAGCTTGCAGTGAGCCGAGATTGCGCCACTGCAGTCCGCAGCACTGCGGCTCCTCA
AluYb9-36 ATCGCACCAGCGTGTCGTGAACCCGGGAAGCGGAGCTTGCAGTGAGCCGAGATTGCGCCACTGCAGTCCGCAGTCCACTGCGGCTCCTCA
AluYb9-37 ATCGCACCAGCGTGTTGAACCCGGGAAGCGGAGCTTGCAGTGAGCCGAGATTGCGCCACTGCAGTCCGCAGTCCGCACTGCGGCTCCTCA
AluYb9-38 ATCGCACCAGCGTGTAACCCGGGAAGCGGAGCTTGCAGTGAGCCGAGATTGCGCCACTGCAGTCCGCAGTCCGGCCACTGCGGCTCCTCA
AluYb9-39 ATCGCACCAGCGTGTCCCGGGAAGCGGAGCTTGCAGTGAGCCGAGATTGCGCCACTGCAGTCCGCAGTCCGGCCTCACTGCGGCTCCTCA
AluYb9-40 ATCGCACCAGCGTGTCGGGAAGCGGAGCTTGCAGTGAGCCGAGATTGCGCCACTGCAGTCCGCAGTCCGGCCTGGCACTGCGGCTCCTCA
