## Supplementary Document 3 for "Recombination of repeat elements generates somatic complexity in human genomes"

**Supplementary Document-3: Preparation of nuclear suspensions for FANS (adapted from Matevossian et al, 2008).**

**Tissue processing**

1. Put ~1200mg of frozen tissue or pulverized tissue in 20ml NEB lysis buffer in a 40ml douncer
2. Leave on ice until tissue is completely thawed
3. Dounce the tissue with 55 strokes, keep the douncer in ice at all times
4. Transfer the 20ml lysate in 2 clear 40ml ultracentrifuge tubes (10ml+10ml), keep tubes on ice
5. Add 18ml of Sucrose cushion per tube. Pipette the sucrose cushion in the bottom of the tube to create a gradient with lysate on top
6. Weigh tubes and equilibrate them with NEB buffer if different
7. Ultracentrifuge in a SW28 rotor at 107163 RCF (24400rpm) for 2.5hrs at 4C
8. Remove supernatant and debris with a vacuum, be careful not to disturb the pellet
9. Add 900ul of PBS1x on the pellet in each tube, pipette to re-suspend the pellet and let sit on ice for 20 minutes to help the pellet dissolve easier
10. After the 20 minutes on ice, pipette up and down to dissolve thoroughly the nuclei clumps. Leave the sample on ice all time.

**Immunostaining**

1. For each different sample combine 300ul PBS 1x with 100ul blocking solution
2. For the "Unstained" control and "Secondary Antibody" control add 150ul PBS 1x with 30ul blocking solution in each tube
3. Add 2.9ul of NeuN Antibody and 2ul of secondary antibody in the tubes for FACS samples and mix thoroughly by pipetting up and down.
4. For the "Unstained" control no antibodies are added. For the "Secondary Antibody" control dilute 1:10 Secondary antibody (1ul Antibody + 9ul PBS 1x) and add 3ul of the dilution to the "Secondary Antibody" sample
5. Incubate for 5 minutes at RT in the dark
6. Meanwhile, re-suspend again the nuclei pellets and combine the content of the 2 pellets in a single tube
7. When incubation is complete, add 800ul of the re-suspended nuclei to the staining mixtures. Prepare also unstained and negative control by adding 50ul of re-suspended nuclei to each control tube.
8. Mix briefly by inverting the tubes, wrap each tube in aluminum foil and incubate samples in cold room on rotating wheel at low speed overnight.

**Nuclei sorting**

1. Filter samples with a 40um filter
2. After setting the filters for elimination of cellular debris, nuclear fragments and nuclei doublets according to manufacturer's suggestions, set proper gates and filters for FITCH detection and check the "Unstained" and the "Secondary Antibody" controls for background noise (Supplementary Fig. 1)
3. Load the nuclei samples and proceed to sorting the $NeuN^-$ and $NeuN^+$ fractions in separate collection tubes containing 500ul 1x PBS. Collect at least 1.5M nuclei per tube.

4. When sorting is finished, re-run each sample to assess the purity of the sorted fractions (Supplementary Fig. 2)

**After sorting**

1. Adjust the volume of sorted samples to 10ml with 1X PBS and add:
   - 2ml Sucrose 1.8M
   - 50ul 1M $CaCl_2$
   - 30ul 1M $Mg(Ac)_2$

   Keep samples on ice at all times.
2. Invert samples for several times and place on ice for 15 minutes
3. Centrifuge samples in a swing bucket rotor at 1786 RCF (1500rpm) at 4C for 15 minutes
4. Discard supernatant with a vacuum without disturbing the pellet
5. Freeze the pellet at -80C or dissolve them in lysis buffer in 900ul of lysis buffer per tube, pipette up and down to redissolve and store at -80C until further processing.

**List of Antibodies.**

Primary Antibody:
Anti-NeuN Antibody, clone A60, Millipore (MAB377)

Secondary Antibody:
Alexa Fluor 488 Goat Anti-Mouse IgG (H+L) Antibody (A28175), Thermo-Fisher

**Buffers composition.**

NEB (Nuclei Extraction Buffer):

0.32M Sucrose (Sigma), 5mM $CaCl_2$, 3mM $Mg(Ac)_2$, 0.1mM EDTA, 10mM Tris-HCl pH8, 1x, Protease inhibitors cocktail (Roche), 0.1mM PMSF (Sigma), 0.1% Triton X-100 (Sigma).

Sucrose Cushion for NEB:
1.8M Sucrose (Sigma), 3mM $Mg(Ac)_2$, 10mM Tris-HCl pH 8.

Blocking solution:
0.5% BSA (Sigma), 10% Normal Goat Serum (Life Technologies), 1X PBS.

The FACS used was a FACSAria II (BD). Initial set-up, filtering of cell doublets and triplets signals and gating were all performed according to manufacturer's instructions.
