## Supplementary Document 4 for "Recombination of repeat elements generates somatic complexity in human genomes"

**Supplementary Document-4: Sanger sequencing results**

LEGEND
< > = TE sequence start/end
[S>] [<S] = Sanger sequence validated start/end
... = split-split junction (breakpoint)

NOTE:
- For each target:
-> the first sequence is the sequenced contig
-> the second sequence is the sanger fasta

INTERCHROMOSOMAL RECOMBINATION

Inter-1
CTTGGGGACCATTAAAGTGGGCCCAGTGCTGTGCTACACTGTAGGCAGATTAGTTCCAAA[S>]CTGA
GGACATCAACAAAGAAGGGCCACTAATAAAGTAAAATCATGGCTGGAAGA<GTTGTTCACACC...GG
TAATCCCAGCACTTTGGGAGGCCAAGGCAGGCGGATCATTTGAGGTCGGGAGTTTGATACCAGCCTGA
CTAACATGGTAATATTTGAGTCTGATGAATTTTGTTGAAGTTCTTTACATCCATGCTGGGAAGCAACT
GTCAGAA[<S]ATGTTAA

NNNNNNNGNNANTAGTNNNNCTGAGGACATCAACAAAGAAGGGCCACTAATAAAGTAAAATCATGGC
TGGAAGAGTTGT
TCACACCGGTAATCCCAGCACTTTGGGAGGCCAAGGCAGGCGGATCATTTGAGGTCGGGAGTTTGATA
CCAGCCTGACTA
ACATGGTAATATTTGAGTCTGATGAATTTTGTTGAAGTTCTTTACATCCATGCTGGGAAGNNCTGTCA
GAAA

Inter-2
TGGAGCTGCATCAGGCATGATTGCCCCTAGCCAAATGACTGCTCTCCCTGTCTCTCATGATTTCTTCT
T[S>]CTCTTACTACTCCCC<TTTTTTTTCT...TGAGACGGAGTCTTGCTCTGTCGCCCAGCCTGGA
GGGCAGTGGTGCGATCTTAGCTCACTGCAACCTCCACCACCTGGGTTCAAGCAATTCTCCTGCCTCAG
CCTCCCAAGTAGTTG>ATCCAGTAGCTTCTTTAACCAGGGTGTGCTTATTGTCCACCTACTGCG[<S]
TGCCGGGCCCCGCCTGTC

NNNNNNCTCTCTTACTACTCCCCTTTTTTTTCTTGAGACGGAGTCTTGCTCTGTCGCCCAGGCTGGAG
GGCAGTGGTGCG
ATCTTAGCTCACTGCAACCTCCACCACCTGGGTTCAAGCAATTCTCCTGCCTCAGCCTCCCAAGTAGT
TGATCCAGTAGC
TTCTTTAACCAGGGTGTGCTTATTGTCCACCTACTGCGA

Inter-3
CTGAAACCAGATATTTGAAACTTACCTTCTTCTTAACGTCTAGAATAAAA[S>]TGTTAAAAGTGAAG
AAAAAG<GGCCAGGCGC...GGTGGCTCACACCTGTAATCCCAGCACTTTGGGAGGCCAAGGCAGGCG
GATCACCTGAGGTCAGGAGTTCGAGCC>GGAGTATGAGCTTTCATATTAAAAGAAAGTCTGAGTGTGG
TG[<S]GCTCATCCCTATAAACTCAGCACTTTGGGAGGCTGAGATGGGAGGGTCGCTTGAG

NTNNNNCNNANANATGTTAAAAGTGAAGAAAAAGGGCCAGGCGCGGTGGCTCACACCTGTAATCCCAG
CACTTTGGGAGG
CCAAGGCAGGCGGATCACCTGAGGTCAGGAGTTCGAGCCGGAGTATGAGCTTTCATATTAAAAGAAAN
CNGAGTGTGGTGA

Inter-4
CATCAATGAGTAGATAAACTATGAGGAGCAGGCTTGGATAAAAAAAATTACCATC<[S>]GACCGGGC
TC...GGTGGCTCACGCCTGTAATCCCAGCACTTTGGGAGGCCGAGGTGGGAGGACTGCTTGATGTAG
GAGTTCAAGACCAGCCTGGGCAACATAGGGAGGCCTCGTCTCTACAAAAAATAATAATAAAAA>AAAA
GTTATCACAGTTGGGGTCTTAGGTTTAGAGCTGCTGACCAC[<S]CAAATCTCACAGTGATTCAAATG
CCTTATCTTCCTTTATTA

NNNNNNNNGACCGGGCTCGGTGGCTCACGCCTGTAATCCCAGCACTTTGGGAGGCCGAGGTGGGAGGA
CTGCTTGATGTA
GGAGTTCAAGACCAGCCTGGGCAACATAGGGAGGCCTCGTCTCTACAAAAAATAATAATAAAAAAAAA
GTTATCACAGTTGGGGTCTTAGGTTTAGAGCTGCTGACCACNNNN

Inter-5
TATATGAGGGAAGAGAAAAAGGTAAGAGGAAAATAGCAATACTAAGAGAAAGGATGGA<GGCCGGGCA
A...GGT[S>]GGCTCACGCCTGTAATCCCAGCACTTTGGGAGGCCCAGGAGGGCGGATCACAAGGTC
AGAAAA>CATTGGCATCAACAGTGGCTTAGCATACAGATGTGGGTCAGGATTCTTGCC[<S]AATCAT
AAGGAAACAGT

NNNNNNNGGCTNNNCCTGTATCCCNGCACTTTGGGAGGCCCAGGAGGGCGGATCACAAGGTCAGAAAA
CATTGGCATCAACAGTGGCTTAGCATACAGATGTGGGTCAGGATTCTTGCCCN

Inter-6
AAATGTTTTGGCTTTTTAGATAAGCATAGCATAGCATGATAATGGAATTTGAAGTCAG[S>]ATAGAA
CTGAGCTCCT<GGTTAGGCGCAGTGGCTCA...CGCCTGTAATCCCAGCACTTTGGGAGGCCGAGGCA
GGCGGATGACGAGGTCAGGAGCTCGAGATCATCCTGGCTAACACGGTGAAACCCTGTCCTACTAAAAA
AACAAAAAAATTAGCCGGGC>AGGAGAATTATTAAAGTGCGTCTTTTGTTCATTTGT[<S]TTTAGTA
AAGATTACATGGATAATTAG

NNNNNNANTCNATAGACTGAGCTCCTGGTTAGGCGCAGTGGCTCACGCCTGTAATCCCAGCACTTTGG
GAGGCCGAGGCAGGCGGATGACGAGGTCAGGAGCTCGAGATCATCCTGGCTAACACGGTGAAACCCTGTCCTAC
TAAAAATACAAAAAAATTAGCCGGGCAGGAGAATTATTAAAGTGCGTCTTTTGTTCATTTGTA

Inter-7
CACTCCTCATATCATCTGTAGGTGATTTACTGATGAAACCTACCAGAGGATGGCAAATATTTTCC[S>
]TACTA<CTTGGGTGATCCACCTGCCTCAGCCTCCCAAAGTGCTGGGATTACAGG...TGTAAGTCAC
CGCACTCGGCC>TCTCCACATCTTCTCATCAGATATTTTTGAGTGGGTATTCTTCTCAGAATCAAGAG
GTGGGGATGAAAAACTGTCAGGGATATTCTTACTTCTCAACTCTACACCACATCTCCT[<S]CAATTC
TCAC

NNNNNNNNTTNNTACTACTTGGGTGATCCACCTGCCTCAGCCTCCCAAAGTGCTGGGATTACAGGTGT
AAGTCACCGCACTCGGCCTCTCCACATCTTCTCATCAGATATTTTTGAGTGGGTATTCTTCTCAGAATCAAGAG
GTGGGGATGAAAAACTGTCAGGGATATTCTTACTTCTCAACTCTACCNACATCTCCTNNN

Inter-8
TACTACTTACCTAAACCCCTGATTTAGTGATCTCACTTAGAAACACCTCTCTA<TTTT[S>]TTTTTA
AACGTTGTTTCACTCTGTCTCCCAGATTGGAGTGCAGTGGTGCGACCTCTGCTCACTGC...AAGCTC
CGCCTCCTGGGTTCACGCTATTCTCC>AGGCCACCTCTTTTTAAGGGGATATATTCTGCAGCGAATCT
TAATAGGGCATAT[<S]TTTCTGCTGCTTTCATATAAATCAT

NNNCNCNNNTTTTTTNAACGTTGTTTCACTCTGTCTCCCAGATTGGAGTGCAGTGGTGCGACCTCTGC
TCACTGCAAGCTCCGCCTCCTGGGTTCACGCTATTCTCCAGGCCACCTCTTTTTAAGGGGATATATTCTGCAGC
GAATCTTAATAGGGCATATNNAN

Inter-10
CCCTCACTACGAATTGGTATGTGGGTAATAAACTTTGATTACATAGAATTTGAAAG[S>]TTAA<GCC
AGGCATGGTGGCTCACGCCTGTAATCCTAGCACTCTGGGAGGCAGGGGCGGGCGGACTGTCTGAGCTC
AGGAGTTCGAGA...CCAGCCTGGGCAACATAGCAAGACCCCGTCTCTACAAAAATAAAAA>TAAATC
CCCACTTGGAGTTATCTGCAGTGCTGTTAAGGGA[<S]CTGCAGATGCTAGAGGAGTT

NNNANNNTNGATTNNNNNTTAAGCCAGGCATGGTGGCTCACGCCTGTAATCCTAGCACTCTGGGAGGC
AGGGGCGGGCGGACTGTCTGAGCTCAGGAGTTCGAGACCAGCCTGGGCAACATAGCAAGACCCCGTCTCTACAA
AAATAAAAATAAATCCCCACTTGGAGTTATCTGCAGTGCTGTTAAGGGAA

Inter-11
TGTACCCACACATCCATCTATTGCTCCCAGGTGCCAGGCCCTCCCGGAGAGCCA[S>]GGTCTGGCAT

CCCAGGCCCCTAGACG<TTGTTTGTTTCGTTTTTTTGAC...ACGGAGTCTCGCTCTGTCGCCCAGGC
TGGAGTGCAGTGGCGCGATCTCGGCTCACTGCAAACTCCGCCTCCCGGGTTCACGCCATTCTCCTGCC
TCAGCCTCCCG>CGAAACGGCTACTTCTATTCCACTGGTTTCTC[<S]TCTTTTTCTTGGGAGCTTGT
GCTCATCAATCTACAATG

NNNNNNGNCNNNNGANAGNNGGTCTGGCATCCCAGGCCCCTAGACGTTGTTTGTTTCGTTTTTTTGACACGGAG
TCTCGCTCTGTCGCCCAGGCTGGAGTGCAGTGGCGCGATCTCGGCTCACTGCAAACTCCGCCTCCCGGGTTCAC
GCCATTCTCCTGCCTCAGCCTCCCGCGAAACGGCTACTTCTATTCCCNGGTTTCTCNNN

Inter-12
CAATGTAAGCAGCCTAAATGTCTAACTTAAAAGGAACCAGCTAAACAAACCTCA[S>]GTATAACCTC
AGCATGGAATACCAAAGATC<TTTTTTTTTGAAACTGAGTTTCA...CTCTGTCGCCCAGGCTGGAGT
GCAGTGGCGTGATCTCGGCTCACTGCAAGCTCCACCTCCCGGGTTCACGCCATTATCCTGCCTCAGCC
TCCCGA>AAAACAACTTTCTTAATAGACACCTTACCAATATCCCTTGTCCATAATACTGCCTTCG[<S
]

NNNNNNNNNNNNCCTNNGTATAACCTCAGCATGGAATACCAAAGATCTTTTTTTTTGAAACTGAGTTT
CACTCTGTCGCCCAGGCTGGAGTGCAGTGGCGTGATCTCGGCTCACTGCAAGCTCCACCTCCCGGGTTCACGCC
ATTATCCTGCCTCAGCCTCCCGAAAAACAACTTTCTTAATAGACACCTTACCAATATCCCTTGTCCATAATACT
GCCTTCGA

Inter-13
TATAATTAAGGTTACAATAATTATCACATCTGGGCTGTGACTGTCTTCCAAAAGAAATGTT[S>]ATG
GG<GGCCGGGCGCGGTGGCTCA...CGCCTGTAATCCCAGCACTTTGGGAGGCCAAGGCGGGTGGATC
GCTTGAGCTCAGGAGTTCGAGAGCAGCCTGGGCAACATGGCAA>ACATGTTGTTTTCCATTCCCATTG
ACCCTTCTCTGTAGGGCCCATTCTCTGA[<S]TTTTCTCAACATGAAAATCAGTGGCCAGTGAGGCT

NNNNNANGGGGGCCGGGCGCGGTGGCTCACGCCTGTAATCCCAGCACTTTGGGAGGCCAAGGCGGGTG
GATCGCTTGAGCTCAGGAGTTCGAGAGCAGCCTGGGCAACATGGCAAACATGTTGTTTTCCATTCCCATTGACC
CTTCTCTGTAGGGCCCATTCTCTGAA

Inter-14
TTACAATTATCCCATGGTAGAGCTTAGCATCTGTTAGCTAATGAAACAG[S>]ACTGGGCGTCGCCAG
AGAAAAGCACCCA<CTGGTCTCAA...TCTCCTGACCTCGTGATCCGCCCGCCTCAGCCTCCCAAAGT
CTGGATTACAGGCATAAGCCACCGCGCACAGCC>CCATATCTTATTCTTAATCATGATTAAGGATTAA
GTAACCCCACTCCATTCCTCCCTCCCCACCATTGCCTTGTTATTCA[<S]GTAACCTTCATTTTTACC
GAGTGTGGACTAGA

NNNNNANNACTGGGCGTCGCCAGAGANAGCACCCACTGGTCTCAATCTCCTGACCTCGTGATCCGCCC
GCCTCAGCCTCCCAAAGTCTGGATTACAGGCATAAGCCACCGCGCACAGCCCCATATCTTATTCTTAATCATGA
TTAAGGATTAAGTAACCCCACTCCATTCCTCCCTCCCCACCATTGNNTGTTATTCAA

Inter-15
AGAGAACAAGCGGCCCGGAGACCACAGCCTCAGACAAGGTGCTGCAGCGTACAGCTCGGGCCAAAGGC
CTCTAAAAACGCAGGGGAAGGCAGGTA<GGGCGCAGTGGC...GCACGCC[S>]TGTGATCCCAGCAC
TGTGGGAGGCCGAGGCGGGAGGATTGCTTGGGGCCAGCAGTTCAAGACCAGCCTGGGCAACAAAGTGA
GACCCTTGTCTCTACCAACAACAAAAAAA>AGTGAACTTTTCTTACAAAATAAATCGAGTGAAATCCT
G[<S]CTGCGAGGCCCACATTGTAGAGT

NNNNNNNNNNNNTGTGANCNNGCACTGTGGGAGGCCGAGGCGGGAGGATTGCTTGGGGCCAGCAGTTCAAGACC
AGCCTGGGCAACAAAGTGAGACCCTTGTCTCTACCAACAACAAAAAAAAGTGAACTTTTCTTACAAAATAAATC
GAGGNAATCCTGNNN

Inter-16
TCCTTCTCCTGTTAGTTGCCCTAATTTCTTATGACAGTAAGAATAACTTAA[S>]AATATTCTTTTA<
AATCCCAGCTACTTGGGAG...GCTGAGACAGGAGGGTCACTTGAACCTGGGAGGCGGAGGTTGCAGT
GAGCCGAGATTATGGCACTGCACTCCAGACTGGATGACGAAGTGAAATTCAATTAAAAAAAAAAAAAA
AAA>GATAAGGTTATGGGCTGTAGCTTGCTAATTTCTGGTATAAGACTTTGAAGGACACGGGAAAGGG
T[<S]

NANNNNNANNCTTNAATATTCTTTTAAATCCCAGCTACTTGGGAGGCTGAGACAGGAGGGTCACTTGA
ACCTGGGAGGCGGAGGTTGCAGTGAGCCGAGATTATGGCACTGCACTCCAGCCTGGGTGACGGAGTGAGATTCT
GTTTCAAGAAAAAAAAAAAAAGATAAGGTTATGGGCTGTAGCTTGCTAATTTCTGGTATAAAACTTTGAAGGACC
GNGAAAGGGTAAN

Inter-18
AATTTCCCATCTTGACGAGCTGTTTGGAATTTCCCAT[S>]ACTGACTGGAAGTGTTTGT<TTAATGG
GTGCA...GCACACTAACATGGCACATGTATACATATGTAACAAATCTGTATGTTGTGCACATGTACC
CTAGAGCTTAAGGTATATAAAAAAA>GAGCAATTTGTCCCAATATGTAAAACATATAATCATTCCTGT
ACAGATCAAATTAATGCAGCCTGAAATC[<S]TTATT

NNNNACTGNANTGTTTGTTTATGGGTGCAGCACACTAACATGGCACATGTATACATATGTAACAAATC
TGTATGTTGTGCACATGTACCCTAGAGCTTAAGGTATATAAAAAAAGAGCAATTTGTCCCAATATGTAAAACAT
ATAATCATTCCTGTACAGATCAAATTAATGCAGCCTGAAATCNAGNN

Inter-19
TGGAGTGACCTTAGTTTTCAGTGTTAGAAATTAAGTCAGAAAGCAG[S>]TTTCTAAAATTTCTCACA
GAAAAAGTCAGAGGAAGTGTGTTGAAATAT<CACCA...GCATGGCACATGTATACATGTGTAACTAA
CCTGCACAATGTGCACATGTACCCTAAAACTTAAAGTATAATAAAAAAAAAAGAA>TACGTTAATATG
TTTAAATAGAAATATCTTAAATAATTCACAAACCAGTCTTTTATTATCCTTCAGCATTCCCAAA[<S]
TTATTTTAGTGAAAAAAAAAAAAAAACTG

NNNNNNNNNNNTTTCTAAATTTCTCACAGAAAAAGTCAGAGGAAGTGTGTTGAAATATCACCAGCAT
GGCACATGTATACATGTGTAACTAACCTGCACAATGTGCACATGTACCCTAGAACTTAAAGTATAATAAAAAAA
AAAGAATACGTTCATATGTTTAAATAGAAATATCTTAAATAATTCACAAACCAGTCTTTTATTATCCTTCAGCA
TTCCCAANN

Inter-21
CATGTGCCCACTGGCCAGAGTGAGATTCCTCAACAGGTAGCCATGGCTTATTTTCAGA[S>]TGTGTC
TTGCCTTCTGGGGCCATAAGGATATCG<GCGGCAAACCACCATGGCACATGTATACCTATGTAACAAA
CCTGCACGTTCTGCACATTTATCCCAGAAC...TTAAAAGTATAATAAAAAAAGA>TGAGTCAAAATG
ATGGAGTTGCCACATCTACCTTATGTGCACATCTCACAGCTTCTGTTTCTGGGTCC[<S]TATTCTTC
CTGGNNNNNNNGNNNNNNNNNATGTGTCTTGCCTTCTGGGGCCATAAGGATATCGGCGGCAAACCACCATGG
CACATGTATACCTATGTAACAAACCTGCACGTTCTGCACATTTATCCCAGAACTTAAAAGTATAATAAAAAAAG
ATGAGTCAAAATGATGGAGTTGCCACATCTACCTTATGTGCACATCTCACAGCTTCTGNNCTGGGTCCNN

Inter-22
GGATTTAATTCAGGTCAACAAATTTACTCTGTAGTCACACTTAGTAAGAT[S>]CTCATTTATATTGG
CTTT<GGCGGGGGTGGGAGGGATAGCATTAGGAGATATACCTAATGTAAATGACAAGTTAATGGGTGC
AGCACACCAACATGGCACATG...TATACATATGTAACAAACCTGCACGTTGTGTACATGTACCCTAG
AACTTAAAGTATTAAAAAAAAAAA>GTAGTACTGGCTGGAGCAGGTTATGGGT[<S]CTCCAAGACATT
ATCTGGAAAAATAATTT

NNNNNNNNANCTCTTTATATTGGCTTTGGCGGGGGTGGGAGGGATAGCATTAGGAGATATACCTAATG
TAAATGACAAGTTAATGGGTGCAGCACACCAACATGGCACATGTATACATATGTAACAAACCTGCACGTTGTGT
ACATGTACCCTAGAACTTAAAGTATTAAAAAAAAAAGTAGTACTGGCTGGAGCAGGTTATGGGTA

Inter-23
AAGTCAGACCTGGGGGTAGGCAGATGGGTGAGCTAGTGGTAACTGCTGTTAAAACCTTCATT<TTCCA
G...CACTTTGGGAGGCCGAGGCGGGCAGATCACCTGCGGTCAGG>GATCAAGGTACGCAGCGTTTTC
CAACTTTCCCTAATGAGAATGCCACATTCAAAACTG[<S]TAGAAAGGTTTATACTTTCTCCATTTAT
TAACTAGAGGAAAT

NNNNNNNNTNNNNNNTTCTTTTCNNCACTTTGGGAGGCCGAGGCGGGCAGATCACCTGCGGTCAGGGA
TCAAGGTACGCAGCGTTTTCCAACTTTCCCTAATGAGAATGCCACATTCAAAACTGA

Inter-24
ATTGGATAAGAATGAACCCTTGTATGGTAAGATACATTATTTAGGAGTCCATTGCAAGATGGCTTGGT

ACCTGT<AGGCTGAGACAGGAGAAT[S>]GGCG...TGAACCCGGGAGGCGGAGCTTGCAGTGAGC>A
ACTTCTTTTTTTTTGTTTGTTTTTGACTGTGCTTGTTCCCTAAG[<S]GAACAAAG
NNNNNNNNNNNNANGGCGNNNCCCGGNAGCGGANCTTGCAGTGAGCAACTTCTTTTTTTTTGTTTGTT
TTTGACTGTGCTTGTTCCCTAAGA

Inter-25
AACAAAGGAAAGTTTACTCAACAACATGTTTTCATTTGGACTTCATTTTGTCAAAGCATGAAAGTGAT
GTCAGAAATAGAAATGAGTGTATTCATG<GCACA[S>]CCACCACACCCAGCTAATTTTTGTACTTTT
TGCAGAGAACTCCTGACCTCAAATGATCCACCTGCCTC...AGCCTCCCAAAGTGTTGGGATTACAGG
CGTGAGCCACCATACCCAGCC>ACAATTGAAGCCATTTCTACCCCCATCCTGGCCACTTCTCAAGAGG
CATCTGT[<S]

NNNNGANTGNTTCTGGNNNCCACCACACCCAGCTAATTTTTGTACTTTTTGCAGAGAACTCCTGACCT
CAAATGATCCACCTGCCTCAGCCTCCCAAAGTGTTGGGATTACAGGCGTGAGCCACCATACCCAGCCACAGTTG
AAGCCATTTCTACCCCCATCCTGGCCACTTCTCAAGAGGCATCTGTA

Inter-26
CAGAAAAGCATCACTCTCATTCCTGGTCTCCCTCCCTCCACTGCTCTCCTCTACAA[S>]TCCC<ACC
TCAGGTGATCCTCCCGCCTCG...GCCTCCCAAAGTGCTGGGATTACAGGTGTGAGCTGCCGTGTCTG
GTCTGCCTCTCCGTCTTTCTCTCTCTCTGTCTTCCT>CCATCTCTCTTCGCATCGCTTTCTGCCTCCC
CATCATTCTCCATGTTTTCCCTTCCCATCT[<S]CTCC

NNNNNNTCTCTCTACATCCCNCCTCAGGTGATCCTCCCGCCTCGGCCTCCCAAAGTGCTGGGATTAC
AGGTGTGAGCTGCCGTGTCTGGTCTGCCTCTCCGTCTTTCTCTCTCTCTGTCTTCCTCCATCTCTCTTCGCATC
GCTTTCTGCCTCCCCATCATTCTCCATGTTTTCCCTTCCCATCTNNNNN

Inter-27
AAATTACAGAAGAATGCAAGTTTTAAACCATTCTCTAATGCCTCCTTCCCTTCACAGTATAAATAAAT
AAATA[S>]TATATTTA<TTTCACCGTGTTAGCCAGGATGGTCTCGATCTCCTGATCTCC...TGATC
CACCTGCCTCGGCCTCCCAAAGTGCTGGGATTACAGGCATGAGACACCGCACCCGGCC>AATAATTTT
TAAATATTTATATGAGAGGCAGAGAAATTTTGTAGGAAAGGAATGCTTTGTATGGAAGAGAGTTAAGA
TGCACA[<S]AAGACAGTAAGTGGTTCAAG

NNNNNNNNNNTATATTTATTTCCCGTGTTAGCCAGGATGGTCTCGATCTCCTGATCTCCTGATCCACC
TGCCTCGGCCTCCCAAAGTGCTGGGATTACAGGCATGAGACACCGCACCCGGCCAATAATTTTTAAATATTTAT
ATGAGAGGCAGAGAAATTTTGTAGGAAAGGAATGCTTTGTATGGAAGAGAGTNAGATGCACAA

Inter-28
TAGGATAGTCAAAGCCCCCAAGCTGGAGTCCGTCATTAGAGCGCCACTCACCCCATTGTTCTTTTAAA
TAGACCAGTCCCCCGGTGCTTTTAAA<ATGGCACACGTATAC...TTATGTAACAAACCTGCACGTTC
TGCACATGTACCCCAGAACTTAAAGTATAATAATAAAAAAAGAA>AAACCACACTGTGAGACTGC[<S
]CATATTTGAACATGAGTTTATACGTCCTACATTCAAAA

NNNNTNNNNTTTTAATAGACCNGTCCCCCGGTGCTTTTAAAATGGCACACGTATACTTATGTAACTAA
CCTGCACGTTCT
GCACATGTACCCCAAAACTTAAAGTATAATAATAAAAAAAAAACCACCCTGNTGNNNTGCTATAATNG
ANCATCATGANTA

Inter-29
AAAAAGAAAGAGGTGCTTTTGAATCATTTAATTTGTTAGGGCATTTTACTTGTGAGGTTGGTATTAT<
TAATGCTAGATT...A[S>]CGAGTTAGTGGGTGCAGCGCACCAGCATGGCACATGTATACATATGTA
ACTAACCTGCACATTGTGCACATATACCCTAAAACTTAAAGTATAATTAAAAAAAAAAA>GAAGCTAC
ATCATTTCTATTTTTGTAACTCCCATTGGCTAACACACAATAAGCA[<S]ATCAATCAATAA

NNNNTNCNNANNCGAGTTAGTGGGTGCAGCGCACCAGCATGGCACATGTATACATATGTAACTAACCT
GCACATTGTGCACATATACCCTAAAACTTAAAGTATAATTAAAAAAAAAAAGAAGCTACATCATTTCTATTTTT
GTAACTCCCATTGGCTAACACACAATAAGCAA

Inter-31

CTCTTGAATTGAGGTTTTGCTACGTTTTATTTTTGAGGTTTAAGAAT[S>]TATTCATTTAGGTATC<
TGGGTGCAGCGCACCAGCGTGGCACATGTATACATATGTAACTAATCTGCACAATG...TGCACATGT
ACCCCTGAACTTATAAGTTGGAAATAAAAAAA>AAAGTGAGTGCATGAAGTTAGAAAAGGACGCAGGA
GAAA[<S]ATTTTTCATGTGGTGATCTTTAGAACCTAAGCTTTT

NNNNNNANTATTCTTTAGGTATCTGGGTGCAGCGCACCAGCGTGGCACATGTATACATATGTAACTA
ATCTGCACAATGTGCACATGTACCCCTGAACTTATAAGTTGGAAATAAAAAAAAAAGTGAGTGCATGAAGTTAG
AAAAGGACGCAGGAGAAAAA

Inter-32
AACAATCGAAATGAATGAATTTAAAGGTATGATGTAATCACCTAGAAAAATGACATTAAAAGCAGTCT
TGAAAATGAGCATTAT<AGCACACCAGCATGGCACA[S>]TG...TATACCTATGTAACAAACCTGCA
TGTTTAGCACATGTATCCCAGAACTTAAAGTAAGATAATAAA>TCCACATTTCAGTCATCATTTTGAA
CTATGTGCAATCCATCCACTCAGTT[<S]CACTCATTTATC

NNNNNNNNNNNTGTATACCTATGTAACAAACCTGCATGTTTAGCACATGTATCCCAGAACTTAAAGTA
AGATAATAAATCCACATTTCAGTCATCATTTTGAACTATGTGCAATCCATCCACTCAGTTN

Inter-33
CTTGTTCACCCACTGCGCTGAGCACTGGGTCTGCCCAGTGAGGAAGATG[S>]CAGTCCTGG<TTTTT
TTATTTTTTATTTTTGAGACGGAGTCTTGCTCTGTCCCCAGGCTGGAGTGCAGTGGCGTGATCTCAGC
TCACTGCAAGCTCCACCTCCCAGGTTCACACCATTCTCTCGCCTCAGCCTCCCGAGTAGCTGGGACTA
CAGGCGCCCG...CCACCATGCCCAGC>CTGGGCACCCAGCCTGCCCAGCCAGCTTTTTTGAGCTTCA
GATT[<S]CACGTAAAATGGGAATGAGA

NNNNNNNNANGNNNNCAGTCCTGGTTTTTTTATTTTTTATTTTTGAGACGGAGTCTTGCTCTGTCCCC
AGGCTGGAGTGCAGTGGCGTGATCTCAGCTCACTGCAAGCTCCACCTCCCAGGTTCACACCATTCTCTCGCCTC
AGCCTCCCGAGTAGCTGGGACTACAGGCGCCCGCCACCATGCCCAGCCTGGGCACCCAGCCTGCCCAGCCAGCT
TTTTTGAGCTTCAGATTN

Inter-34
TGGAGCTGCATCAGGCATGATTGCCCCTAGCCAAATGACTGCTCTCCCTGTCTCTCATGATTTCT[S>
]TCTTCTCTTACTACTCCCC<TTTTTTTTCT...TGAGACGGAGTCTTGCTCTGTCGCCCAGCCTGGA
GGGCAGTGGTGCGATCTTAGCTCACTGCAACCTCCACCACCTGGGTTCAAGCAATTCTCCTGCCTCAG
CCTCCCAAGTAGTTG>ATCCAGTAGCTTCTTTAACCAGGGTGTGCTTATTGTCCA[<S]CCTACTGCG
TGCCGGGCCCCGCCTGTC

NNNNNNNANNTNNTCTTCTCTTACTACTCCCCTTTTTTTTCTTGAGACGGAGTCTTGCTCTGTCGCCC
AGGCTGGAGGGCAGTGGTGCGATCTTAGCTCACTGCAACCTCCACCACCTGGGTTCAAGCAATTCTCCTGCCTC
AGCCTCCCAAGTAGTTGATCCAGTAGCTTCTTTAACCAGGGTGTGCTTATTGTCCCAN

Inter-35
TGAGTTTGCATCGCTGGTGGGTGCCTGCAGGTCTCCAGGCTGTGCTGACTCAGATTTCAAAT[S>]GG
GGCTG<GGGCACAGTGG...CTCATGCCTGTAATCCCAGCACTTTGGGAGGCCAAGGTGGGTGGATCA
TTTGGGGTCAGGAGTTCAAGACCAACCAGCCCGGCCAACATGGCAAAACCCTGTCTCTA>TGGGTACC
TCCTAGGGTCTTTGTAAGGATAGAACAACTAACAGGGTGTCTGCTAACCCGATAA[<S]T

NNNNNNNGGGGCTGGGGCCAGTGGCTCATGCCTGTAATCCCAGCACTTTGGGAGGCCAAGGTGGGTGG
ATCATTTGGGGTCAGGAGTTCAAGACCAACCAGCCCGGCCAACATGGCAAAACCCTGTCTCTATGGGTACCTCC
TAGGGTCTTTGTAAGGATAGAACAACTAACAGGGTGTCTGTNACCCGATAAANA

Inter-36
ACATCCTTCCCCTACCCACATACCTACCTATCATTGGTC[S>]CTCTTGTTTGTG<TTTTTTTGTTGT
TG...TTTTTTTTTGAGAGGGAGTCTCGCTCTGTCGCCCAGGCTGGAGTGCAGTGGCATGA>GGGGAG
AACATTTTTAATCCTTCTAGATACCCATGTCCC[<S]TATTATTTTTCAACTTTTGTTTCTCAAGAGT
CTT

NNNNNCTCTTGTTTNNGTTTTTTTGTTGTTGTTTTTTTTTGAGAGGGAGTCTCGCTCTGTCGCCCAGG
CTGGAGTGCAGTGGCATGAGGGGAGAACATTTTTAATCCTTCTAGATACCCATGTCCCNCNN

Inter-37
AGATTTGTACAAGAAATCTAGACCCTTAGAAATAGCCTAACTAAATGTCAGCAATTAAGATCTAAGTA
TAT[S>]T<GGCTGGGTGCAGTGGCTCACGCTAGCACTTTGGGAGGCTGAGGTGGGGCAGATCACCTG
AGGTCAGGAGTTCAAGACCAGCCTAGTCAACATGGTGAAACCCC...GTCTCTACTAAAAATACAAAA
TTAGTCAGGCATGGTAGCGC>ATGTGGGAGGTAATATATCCATTACTCTGACCTGTCT[<S]TTTAAA
AACAGAGTGGTAGACCAGGTA

NNNNTNTGGCTGGGTGCAGTGGCTCACGCTAGCACTTTGGGAGGCTGAGGTGGGGCAGATCACCTGAG
GTCAGGAGTTCAAGACCAGCCTAGTCAACATGGTGAAACCCCGTCTCTACTAAAAATACAAAATTAGTCAGGCA
TGGTAGCGCATGTGGGAGGTAATATATCCATTACTCTGACCTGTCTANN

Inter-38
CCCTACATTTCTATTCAGAGTACTGTTTTTAGATGTGAAATATAAGCCTGCGGCCTTAACTCTGTATT
AAAAAAAATGTTTTTGTTTAAAAAAAAC[S>]TGTTCCC<AT...GGGTGCAGCAAACCAACATGGCA
CATGTATACATATGTAACAAACCTGCACATTGTGCACATGTACCCTAGAACTTAAAGTATAATTTAAA
AAATAAAAAA>AATAAAAAGACAAAAGAGAATATCGTCAAGC[<S]AACTTGACTTCTTTTACTTCTT
G

NNNTNNTTNNNNNNTGTTCCCATGGGTGCAGCAAACCAACATGGCACATGTATACATATGTAACAAAC
CTGCACATTGTGCACATGTACCCTAGAACTTAAAGTATAATTTAAAAAATAAAAAAAATAAAAAGACAAAAGAG
AATATCGTCAAGCN

Inter-39
TGAAGAAGATTTCAAGGGCAATAGGATTGTGTTGTTAAAAAAGTGTAACAT[S>]AGACAAATTTTGC
ATAATACAGAGTACTAGAGTCTA<TTTTTTTTTATTATACTTTAAGTTCTAGGGTACATGTGCACAAT
...GTGCAGGTTAGTTACATATGTATACATGTGCCATGTTGGTGTGCTGCACCCATTAACTCGTCATT
TAACATT>TTAGGAGGACGTAATGATGGTGTGGAAACAATCAAGTCACGCCTTTTGC[<S]CTTGGCT
G

NNNNNNNNNNGTACNAGACAATTTTGCATAATACAGAGTACTAGAGTCTATTTTTTTTTATTATACTT
TAAGTTCTAGGGTACATGTGCACAATGTGCAGGTTAGTTACATATGTATACATGTGCCATGTTGGTGTGCTGCA
CCCATTAACTCGTCATTTAACATTTTAGGAGGACGTAATGATGGTGTGGAAACAATCAAGTCNNCCTTTTGCA

Inter-40
TTACTCTAAGGAAGACGTAGATTAGAATGGAATTATTTTTATTCAAAAACT[S>]GTTT<TTATTATA
CC...TTAAGTTCTGGGGTACCTGTGCACAACGTGCAGGTTTATTACATAGGAATACATGTGCCATGT
TGGTTTGCTGCACCCATCAACCCATCATTTACGTTAGGTATTTCTCCTAATGCAATCCCTCCCCCAAA
CCCCTACCC>TTGACACACATTTTATGAAACACAC[<S]CTTTATTTCTTAAAGTTTCTTCTGGATCT
CATTTAGTAA

NNNNNNNNANNGTTTTTATTATACCTTAAGTTCTGGGGTACCTGTGCACAACGTGCAGGTTTATTACA
TAGGAATACATGTGCCATGTTGGTTTGCTGCACCCATCAACCCATCATTTACGTTAGGTATTTCTCCTAATGCA
ATCCCTCCCCCAAACCCCTACCCTTGACACACATTTTATGAAACACAC

Inter-42
AATCACATGGACGCTAGAGGCTGCAGATACCTCCGGAGACGAGGCTTGGCCTGGCCTGGGGCCTCCG[
S>]TTTG<AGAACTGGGAGATACACCTAATGCTAGATGACAAGTTAGTGGGTGCAGCGCACCAGCATG
GCACATGTATACATATGTAACTAACCTGCACATTGTGCACATGTAC...CCTAAAAGTTAAAGTGTAA
TAAAAAAAAA>GAGTAGGATTGAATATCATAGTTTGATCTCCCAGACTGGGAAGCACATGG[<S]TGT
TGCTTTAAACTCGGACGCCAGAGCTGAAT

NNNNCNNNNNNGGCNNNTTTGAGACTGGGAGATACACCTAATGCTAGATGACAAGTTAGTGGGTGCAG
CGCACCAGCATGGCACATGTATACATATGTAACTAACCTGCACATTGTGCACATGTACCCTAAAAGTTAAAGTG
TAATAAAAAAAAAAGAGTAGGATTGAATATCATAGTTTGATCTCCCAGACTGNAANGCACATGGNNN

Inter-43
GCTGAGTAAGAGTTTTTGTAGAATTACATTTGCAACCTCTACAGGATTAAAAATCACTAAGCA<G[S>

]GGTGATGCACACCAGCATGGCACATGTATACGTATGTCACTAACCTGCACATTGTGCACATGTACCC
TAAAACTTAAAG...CATAATAATAATAAAAAAAAAGAATAACCTTGCTGTTACAAAGTGCCTGGTAA
TCATTATCCTAACTTTACTTGATTAG>GCAAAGAAAGCAGTATTTTGTCTGAAAAGTCCATACCG[<S
]GGG

NNNNNNNNNNNNNNNNNGGTGATGCACACCAGCATGGCACATGTATACGTATGTCACTAACCTGCACA
TTGTGCACATGTACCCTAAAACTTAAAGCATAATAATAATAAAAAAAAAGAATAACCTTGCTGTTACAAAGTGC
CTGGTAATCATTATCCTAACTTTACTTGATTAGGCAAAGAAAGCAGTATTTTGTCTGAAAAGTCCATACCGAN

Inter-44
GGATTAAAAATCACTAAGCA<GGGTGATGCACACCAGCATGGCACATGTATAC[S>]GTATGT...AA
CTAACCTGCACGTTGTGCACATGTACCCTAAAACTTAAAAGTATAATAAAAAAAAGA>TCAAAAATGT
AAATTTGGCTGCCATATGTATTAAAGTCC[<S]CCTCAGTTTAA

GNNNNNNNNNNNGNNNNGTATACGTATGTAACTAACCTGCACGTTGTGCACATGTACCCTAAAACTTA
AAAGTATAATAAAAAAAAGATCAAAAATGTAAATTTGGCTGCCATATGTATTAAAGTCCNNNNN

Inter-46
TTGGAAAAACGCTTATTGTTCTGATAAAATTGCTGAGTAAGAGTTTTTGTAGAATTACATTTGCAACC
TCTATAGGATTAAAAATCACTAAGCA<[S>]GGGTG...CAGCACACCAGCATGGCACATGTATACAT
ATGTAACTAACATGCACATTGTGCACAAGTACCCTAAAACGTAAAGTAAATTAAAAATAAA>TACAAA
AGAATGCTTACAAATTGCTCAATCCAAAGTAAGGTTCCA[<S]

NNNNNNNNNNNNNNNNNGGTGCAGCACACCAGCATGGCACATGTATACATATGTAACTAACATGCACAT
TGTGCACAAGTACCCTAAAACGTAAAGTAAATTAAAAATAAATACAAAAGAATGCTTACAAATTGCTCAATCCA
AAGTAAGGTTCCAAA

Inter-47
ATTACATTTGCAACCTCTACAGGATTAAAAATCACTAAGCA<GGGTGATGCACACCAGCATGGCACA[
S>]TGTATAC...ACATGTAACAAACCTGCAGGTTGTGCACATGTACCCTAAAACTTAAAAGTATAAT
AAACAAAACAAAA>CAAAAAAACAACAAATCTTATAGCTCTGTAGATCAGAAGTCTGATATGGTTCTC
ACTGAGTTAAATCAAGATGTAGGTAGGCTGCA[<S]TTTTTTT

NNNNNNNNNNNTGNNNTGTATACACATGTAACAAACCTGCAGGTTGTGCACATGTACCCTAAAACTTA
AAAGTATAATAAACAAAACAAAACAAAAAAACAACAAATCTTATAGCTCTGTAGATCAGAAGTCTGATATGGTT
CTCACTGAGTTAAATCAAGATGTAGNTAGGCTGCAAA

Inter-48
ATTTGCAACCTCTACAGGATTAAAAATCACTAAGCA<GGGTGATGCACACCAGCATGGCACA[S>]TG
TATACGTATGTCACTAACCTGCACATTGTGCACATGTACCCTAA...AACTTAAAGTATAATAAA>TT
ACTGTGCAGAAGCTCTTTAGTTTAATTAGATCCCGTTTGTCAATTTTGGCTTCTGTTGCCATTGCTTT
TGGTGTTTTAGACATGAAGTCCTTGCCCACAGAGTGA[<S]ACAGGCAACCTACAGAATGGGA

NNNNNNNNNNNGNNNTGTATACGTATGTCACTAACCTGCACATTGTGCACATGTACCCTAAAACTTAA
AGTATAATAAATTACTGTGCAGAAGCTCTTTAGTTTAATTAGATCCCGTTTGTCAATTTTGGCTTCTGTTGCCA
TTGCTTTTGGTGTTTTAGACATGAAGTCCTTGNCCACAGAGTGA

Inter-49
CCAAATTCTCTGAATTTGGAAA<AGAGGGTGCAGCGCACCAGCATGGCACA[S>]TGTATACATATGT
AACTAACCTGCACATTGTGCA...TGTGTACCCTAAAACTTAAAGTATAATAATAAAAAAA>GTAAAA
AAAAAAGAGTCTTCACTCTGAAGGGGAATCACGGATAGTG[<S]ACCCTTTAAAATTTACCTTCTTGA
CTT

NNNNNNNNNNNGNNNTGTATACNTATGTAACTAACCTGCACATTGTGCATNTGTACCCTAAAACTTAA
AGTATAATAATAANAAANNNAAAAAAAAAAGAGTCTTCACTCTGAAGGGGAATCACGGATAGTGNNNNN

Inter-50
TGAGTGCATGCCAAATTCTCTGAATTTGGAAA<AGAGGGTGCAGCGCACCAGC[S>]ATG...TCATA
TGTATACATATGTAACTAACCTGCACATTGTGCACATGTACCCTAAAACTTAAAGTAGAATAATAATA

A>TAATAAAAAGAGGCACTTTAGTCCTCTACATCAAGTGTGGCAACCA[<S]GTATGAAATACTGCCT
AC

NNNNNNNNNNNNNGNNNNNNATGTCATATGTATACATATGTAACTAACCTGCACATTGTGCACATGTA
CCCTAAAACTTAAAGTAGAATAATAATAATAATAAAAAGAGGCACTTTAGTCCTCTACATCAAGTGTGGCAACC
ANNN

Inter-51
AGGCTCTGCAATCAGAAAGCAGAGTTGTTGTCCCCAAGGAATGTCATCTCTAGCTAGAGATGGAAAAT
CAGCACCAGAAAACAGTTCAGGATTGAGTGCATGCCAAATTCTCTGAATTTGGAAA<AGAGGGTGCAG
CGCACCAGCAT[S>]GGCACATGTATACATATGTAACTAACCTGCACATTGTGCACATGTACCCTAAA
ACTTAAAGTATAATAA...AATAAATAAA>TAAATAAAATACTCCCGTTTCTCTGGTCA[<S]CTTTT
AAAAACTGTAACTAGTATCTTTTCTTGCTA

NNNNNNNNNNNNNNNNNNNNNATGGCACNTGTATACATATGTAACTAACCTGCACATTGTGCACATGT
ACCCTAAAACTTAAAGTATAATAAAATAAATAAATAAATAAAATACTCCCGTTTCTCTGGTCANN

Inter-52
GTGCATGCCAAATTCTCTGAATTTGGAAA<AGAGGGTGCAGCGCACCAGC[S>]ATGGCACATG...C
ATACGTATGTAACTAACCTGCACAATGTGCACATGTACCCTAAAACTTAAAGTATAAAAAAAAAAAGA
AAA>AGAATTTGCATCTTTATAGGGATGGGTGGAAG[<S]GATGGTA

NNNNNNNNNNNNCNNNNNNATGGNACNTGCATACGTATGTAACTAACCTGCACAATGTGCACATGTAC
CCTAAAACTTAAAGTATAAAAAAAAAAAGAAAAAGAATTTGCATCTTTATAGGGATGGGTGGAAGNNNNN

Inter-53
TCAGAAAGCAGAGTTGTTGTCCCCAAGGAATGTCATCTCTAGCTAGAGATGGAAAATCAGCACCAGAA
AACAGTTCAGGATTGAGTGCATGCCAAATTCTCTGAATTTGGAAA<AGAGGGTGCAGCGCACCAGCAT
GGC[S>]ACATGTATACATATGTAACTAACCTGCACATTGTGCACATGTACCCTAAAACTTA...AAG
TATAATAAAAAAGAAAAAA>GAAGTCATAATGTTGAGCTATTGCTGGAAAATTCA[<S]TAGAAATAT
ATT

NNNNNNNNNNNNNNNNNNNNNNNNNACATGTATACATATGTAACTAACCTGCACATTGTGCACATGTA
CCCTAAAACTTAAAGTATAATAAAAAAGAAAAAAGAAGTCATAATGTTGAGCTATNGNTGGAAAATTCAN

Inter-54
GCATTTGCATGTTCACAAACTAATTCAATGTTTACAATTGAAACAG[S>]GTGTTATTATAGTGATTA
GAAAACTGATGAAATGAAAGATAAGAAAAGCTAGTTA<ACCAGCATGACACATGTATACATATGTAAC
AAACCTGCACGTTGTGCACATGTACCCTAGAACTTAAAGTG...TAATAATAATAATAATAATAATAA
TAATAATAATAATAAAATA>GACATTTCTTTGGATTTGGTAAGATGGA[<S]GTTATCAGTGACTCGT
AAATAACAAATTGAGATGGGAGCCAGATTG

NNNNNNNNGNNCNGTGTTATTATAGTGATTAGAAAACTGATGAAATGAAAGATAAGAAAAGCTAGTTA
ACCAGCATGACACATGTATACATATGTAACAAACCTGCACGTTGTGCACATGTACCCTAGAACTTAAAGTGTAA
TAATAATAATAATAATAATAATAATAATAATAATAAAATAGACATTTCTTTGGATTTGGTAAGATGGAN

Inter-55
CACAAACTAATTCAATGTTTACAATTGAAACAGGTGTTATTATAGTGATTAGAAAACTGATGAAATGA
AAGATAAGAAAAGCTAGTTA<ACCAGCATGA...CAC[S>]ATGTATACATATGTAACAAACCTGCAC
GTTATGCACATGTACCCTAAAACTTAAAGTATAATTTAAAAAAAAA>CCTGGAGGTACCTAGCAGAA[
<S]TAAATACATAGAACTACTAAAAAAAAAAAAAAAAGAA

NNNNNNNNANNTGTATACNTATGTAACAAACCTGCACGTTATGCACATGTACCCTAAAACTTAAAGT
ATAATTTAAAAAAAAACCTGGAGGTACCTAGCAGAA

Inter-56
TGTTCACAAACTAATTCAATGTTTACAATTGAAACAGGTGTTATTATAGTGATTAGAAAACTGATGAA
ATGAAAGATAAGAAAAGCTAGTTA<ACCAGCATGACACATGTATACATATGTAACAAACCTGCAC...
ATTCTGCACATGTATCCCCCAGAATTTAAAGTAAAATTGAAAA>GATAAAGTGAGACTAAGATAC[<S

]TCAGT

NNNNNNNNNNNANNTGTATACNTATGTAACAAACCTGCACATTCTGCACATGTATCCCCCAGAATTTA
AAGTAAAATTGAAAAGATAAAGTGAGACTAAGATACNNNNN

Inter-57
GTTATTATAGTGATTAGAAAACTGATGAAATGAAAGATAAGAAAAGCTAGTTA<ACCAGCATGACACA
TG[S>]TATACATATGTAACAAA...CCTGCACATTGTGCACATGTACCCTAGAAGTTAAAGTATAAT
AAAAAA>TTTTTAAAAATGACCAGCTTACTCGTCTA[<S]TAAAAAAAAAAAATACTTTTGAATAATG
AAGAATTGATAAAGACT

NNNNNNNNNNNNNNNNNTATACNTATGTAACAAACCTGCACATTGTGCNCNNNNACCCTAGAAGTTA
AAGTATAATAAAAAATTTTTAAAAATGACCAGCTTACTCGTCTAAN

Inter-58
TGATGAAATGAAAGATAAGAAAAGCTAGTTA<ACCAGCATGACACAT[S>]GTATACATATGTAACAA
ACCTGCAC...GTTCTGCACGTGTATCCCAGAACTTAAAGTAAAATAAAAA>CTTTTTAAAAAGAACT
ATGTCTATAACATATTAGAGTGACTAGATAAACAAGATTCCTTTAGTATTGCTTAATGACAGGAGTAA
AAGACGA[<S]AAATTGTTAACA

NNNNNNNNTGANCTGTATACNTATGTAACAAACCTGCACGTTCTGCACGTGTATCCCAGAACTTAAAG
TAAAATAAAAACTTTTTAAAAAGAACTATGTCTATAACATATTAGAGTGACTAGATAAACAAGATTCCTTTAGT
ATTGCTTAATGACAGGAGTAAAAGACGA

Inter-59
CTGTATCTCAGATACATT<TGAGGTGGGGGGAGGGGGGAGGGATAG[S>]CATTGGGGGATATACCTA
ATGCTAGATGACGAGTTAGTGGGTGCAGCACACCAGCATGGCAAATGTATACATATGAAACTAACCTG
CACATTGTGCA...CATGTACCCTAAAACTTAAAGTATAATAATAATAAAAAA>CAAAACAAAACATT
ACATTTGCATCATTAATGA[<S]

NNNNNNNNNNNNNNANNNCATTGGGGGANATACCTAATGCTAGATGACGAGTTAGTGGGTGCAGCACA
CCAGCATGGCAAATGTATACATATGAAACTAACCTGCACATTGTGCACATGTACCCTAAAACTTAAAGTATAAT
AATAATAAAAAACAAAACAAAACATTACATTTGCATCATTAATGAA

Inter-60
CAAAACCACAATGAGATACGATGGTATCTGTATCTCAGATACATT<TGAGGTGGGGGGAGGGGGGAGG
GATAG[S>]CATTGGGGGATATACCTAATGCTAGATGACGAGTTAGTGGGTGCAGCAAACCACCATGG
CACACGTATACCTACGTAGCAAACCTGCACATTCTGCACATGCATCCC>ATGAATATTTA[<S]

NNNNNNNNNNNNNNANNNNCATTGGGGGNNATACCTAATGCTAGATGACGAGTTAGTGGGTGCAGCAN
ACCACCATGGCACACGTATACCTACGTAGCAAACCTGCACATTCTGCACATGCATCCCATGAATATTTAA

Inter-61
CCACAATGAGATACGATGGTATCTGTATCTCAGATACATT<TGAGGTGGGGGGAGGGGGGAGGGATA[
S>]GCATT...AGGAGATATACCTAATGTAAATGACGAGTTAATGGGTGCAGCACACCAACATGGCAC
ATGTATACATATGTAACAAACCTGCACGTTGTGCACATGTACCCTAAAACTTAAAGTGTATATAAAAA
AAAAGAA>CAAAAGTATTAAAAAAAAAGAAGAAGAAATGATAAGAAAATGTGAGG[<S]

NNNNNNNNNNNNNGNNGCATTAGGAGATATACCTAATGTAAATGACGAGTTAATGGGTGCAGCACACC
AACATGGCACATGTATACATATGTAACAAACCTGCACGTTGTGCACATGTACCCTAAAACTTAAAGTGTATATA
AAAAAAAAGAACAAAGGTATTAAAAAAAAAGAAGAAGAAATGATAAGAAAATGTGAGGA

Inter-62
GTATCTGTATCTCAGATACATT<TGAGGTGGGGGGAGGGGGGAGGGATAG[S>]CATTGGGGGATATA
CCTAATGCTAGATGACGAGTTAGTGGGTGCAGCACACCAGCATGGCAAATGTATACATATGAAACTAA
CCTGCACATTGTGCACATGTACCCTAAAACTTAAAGTATAATAA...AAAA>GCAATAAATGAATTAA
TATATGCAATGTACTTACAAAAGAG[<S]TTTAA

NNNNNNNNNNNNNNNNNNNCATTGGGGGANATACCTAATGCTAGATGACGAGTTAGTGGGTGCAGCAC

ACCAGCATGGCAAATGTATACATATGAAACTAACCTGCACATTGTGCACATGTACCCTAAAACTTAAAGTATAA
TAAAAAAGCAATAAATGAATTAATATATGCAATGTACTTACAAAAGAGNNN

Inter-63
TCTAGCCTTTCTAGAAATAA<GGCTACTGGGAGGGGGGAGGGATAG[S>]CATTAGGAGATATACCTA
ATGTTAAATGACGAGTTAATGGGTGCAGCACACCAACATGGCACATGTATACATAT...GTAACAAAC
CTGCACGTTCTGCACATGTATCCCAGAACTTTAAGTATAATAATAAAAAAAA>GTAAAAGCCCTGCAT
GAATTAGT[<S]TAACC

NNNNNNANNNGNAANNNCATTAGGAGANATACCTAATGTTAAATGACGAGTTAATGGGTGCAGCACAC
CAACATGGCACATGTATACATATGTAACAAACCTGCACGTTCTGCACATGTATCCCAGAACTTTAAGTATAATA
ATAAAAAAAAGTAAAAGCCCTGCATGAATTAGTNNNN

Inter-65
TAGCCTTTCTAGAAATAA<GGCTACTGGGAGGGGGGAGGGATAGC[S>]ATTAGGAGATATACCTAAT
GTTAAATGACGAGTTAATGGGTGCAGCACACCAACATGGCACATGTATACATATGTAAC...TAACCT
GCACAATGTGCACATGTACCCTAAAACTTAAAGTATAATAAAAAAAAAACATTAAAAAAAATAAATAA
ATAAAATTAAAAAAAAAA>GTAGAATTGAGCATGGGAAGAACAAAT[<S]TATGTACATTTTTGCTTT
TCCTACAATGCTTCTGTGCACC

NNNNNNNNNANANNATTAGGAGANATACCTAATGTTAAATGACGAGTTAATGGGTGCAGCACACCAAC
ATGGCACATGTATACATATGTAACTAACCTGCACAATGTGCACATGTACCCTAAAACTTAAAGTATAATAAAAA
AAAAACATTAAAAAAAATAAATAAATAAAATTAAAAAAAAAAGTAGAATTGAGCATGGGAAGAACAAATAAN

Inter-67
GAAGAATGCCACATAGTACTGATTATTTGTACAGAACTTTAAACTTCTTAAATCAGGTGTACAATCAC
TGCCAGAGTTTTCTAAAAACTGGCAA{S>]GTG<TCGG...GAGGCTGAGGCAGGAGAATCGCTTGAA
CCTGGGAGGCGGAGGTTGCAGTCAGCCGAGATTGCGCCACTGCACTCCAGCCTGGGTGAGAGTGAGAC
TCCATCTCAAACAAACAAACAAACAATAGAA>TTAAACATCTCCTT[<S]CCAGGCCAGT

GGNNNNNNNGGNNGTGTCGGGAGGCTGAGGCAGGAGAATCGCTTGAACCTGGGAGGCGGAGGTTGCAG
TCAGCCGAGATTGCGCCACTGCACTCCAGCCTGGGTGAGAGTGAGACTCCATCATAGAATTAAACAACAAACAA
ACAATAGAATTAAACATCNCCTTCCAGGCCNN

Inter-69
AACTTCTTAAATCAGGTGTACAATCACTGCCAGAGTTTTCTAAAAACTGGCAA[S>]GTG<TCGGGAG
GCTGAGGCAGGAGAATGGCGTGAACCCCGGAAGCGGAGCTTGCAGTGAGCCGAGATTGCGCCACTGCA
GTCCACAGTCCC...GCCTGGGCGACAGA>AAATTTAACGTTCAGGGTTCTAAAGTGCAAACTTTCCA
AAGAGG[<S]TTCTC

NNNNNNNNNGNNGTGTCGGGAGGCTGAGGCAGGAGAATGGCGTGAACCCCGGAAGCGGAGCTTGCAGT
GAGCCGAGATTGCGCCACTGCAGTCCACAGTCCCGCCTGGGCGACAGAAAATTTAACGTTCAGGGTTCTAAAGT
GCAAACTTTCCAAAGAGG

Inter-70
TTGTTTAAGAGTGAAGAAGAA<CTTGGGAGGCTGAGGCAGGAGA[S>]ATGGCATGAACCCGGGAGGC
GAAGGTTGCA...GTGAGCCAAGATCGCACCACTGCACTCCAGCCTGGGTGACAAAGCAAGACTCCGC
CTCAAAAATAAAAAATAAAAAAA>TGCAGATTTTACAAATGTACAAATA[<S]TACAAAAAATACTTT
GTACATT

GGGNNNGNTNNNNNNNATGGCATGAACCCGGGNNGCGAAGGTTGCAGTGAGCCAAGATCGCACCACTG
CACTCCAGCCTGGGTGACAAAGCAAGACTCCGCCTCAAAAATAAAAAATAAAAAAATGCAGATTTTACAAATGT
ACAAATAAA

Inter-71
CATGAACTGAAGCACATCTTAAAGGACTCAGGTCTTTGGTGTCACTCCACTCTGTTGTCTG[S>]CTT
CTTATGGTTGTTGTTTAAGAGTGAAGAAGAA<CTTGGGAGGCTGAGGCAGGAGAATGGCATGAACCCG
GGAGGCGAAGGTTGCAGTGAGCCAAGATCATGCCACTGCACTCCAGCCTGGGCGACAGAGCGAGACTC
CGTCTCAAAA...CAAACAAACAAAAAAAAAAAAAAAAAAA>AACACATTTTCTTCGATATGTTTAAT

GACT[<S]CTATTTA

NNNNNNGNCNGCTTCTTATGGTTGTTGTTTAAGAGTGAAGAAGAACTTGGGAGGCTGAGGCAGGAGAA
TGGCATGAACCCGGGAGGCGAAGGTTGCAGTGAGCCAAGATCATGCCACTGCACTCCAGCCTGGGCGACAGAGC
GAGACTCCGTCTCAAAACAAACAAACAAACAAACAAACCAAAAAAAACACATTTTCTTCGATATGTTAATGACT
NNNNN

Inter-72
GGACTCAGGTCTTTGGTGTCACTCCACTCTGTTGTCTGCTTCTTATGGTTGTTGTTTAAGAGTGAAGA
AGAA<CTTGGGAGGCTGAGGCAGGAGAATGGCATGAACCCGGGAGGC...AGAGGTTGCAGTGAGCAG
AGATCGCGCCACTGCACTCCAGCTTGGGCGACAGAGCAAAACTCTGTCTCAAAAAAGAAAAA>CCGTA
TGGCCGGCA[<S]CGGTGGCTCATGCCTGTAATCCCAGCACTTTGGGAGGCCGAGGCGGGCAAATCAA
GACCATCCTGGCTAATACGGTG

NNNNNNNNNNNNNNNNNANGGCATGAACCCGGGAGGCANAAGGTTGCAGTGAGCCNAGATCANTCGCC
ACTGCACTCCAGCNTGGGCGACAGAGCAANACTCTGTCTCAAAAAAGAAAAACCGTATGGCCGGCAA

Inter-73
TTATGGTTGTTGTTTAAGAGTGAAGAAGAA<CTTGGGAGGCTGAGGCAGGAGA[S>]ATGGCATGAAC
CCGGGAGGCGAAGGTTGCAGTGAGCC...GAGATTGCACCATTGCACTCCAGCCTGGGCAACAAGAGT
GAAACTCCATCTCAAAATAAATACATACATACATACATACATACATACATACTGAAAGAAGTAAA>CC
AAAAGGTTTCTGAATTCTGAA[<S]GTG

NNNNNNNNNAGGNTGANGNNGANATGGCATGAACCCGGGAGGCGAAGGTTGCAGTGAGCCGAGATTGC
ACCATTGCACTCCAGCCTGGGCAACAAGAGTGAAACTCCATCTCAAAATAAATACATACATACATACATA
CATACATACTGAAAGAAGTAAACCAAAAGGTTTCTGAATTCTGAA

Inter-75
CTGCAGATTGCAAAAAGGGAGACATCTTGCCCTTAAGAAAAGACCGT<GGAGTGGTGGCAGGCGCCTG
TA[S>]GTCCCAGCTACTCAGGAGGCTGAGGCAGGAGAATGGCGTGAACCCGGGAGGCGGAGC...TT
GCAGTGAACCGAGATCGCGCCACTGCACTCCAGCCTAGGCGACAGAGGGAGACTTAGAAAAAAAAAAA
A>AAGAAAAAAAAAGGCTTTTTCTGTATTCAAAA[<S]

NNNNNNGNNCNGNNGTCCNGCTACTCAGGAGGCTGAGGCAGGAGAATGGCGTGAACCCGGGAGGCGGA
GCTTGCAGTGAACCGAGATCGCGCCACTGCACTCCAGCCTAGGCGACAGAGGGAGACTTCGTCTCAAAAAAAAA
AGAAAAAAAAAGGCTTTTTCTGTATTCAAAAAA

Inter-76
AAGGGAGACATCTTGCCCTTAAGAAAAGACCGT<GGA...GT[S>]GGCGGCGGGCGCTTGTAGTCCC
AGCTACTCGGGAGGCTGAGGCAGGAGAATGGCGTGAACCCGGGAGGCGGAGCTTGCAGTGAGCGAAGA
TCGCACCACTGCACTCCAGCCTGGGGGACAGAGCGAGACTCCGTCTCAAAAAACAAAACAAAACAAAA
ACAAAACAAAAACAAA>GGGTTCATACAAAGAATTATCCTAAGAAGTGCATAGGA[<S]GCTGGGCGA
GGTGGCTCACGCCTGTAATCCCA

NNNNNNNNNNGNCGGCGGGCGCTTGTAGTCCCAGCTACTCGGGAGGCTGAGGCAGGAGAATGGCGTGA
ACCCGGGAGGCGGAGCTTGCAGTGAGCGAAGATCGCACCACTGCACTCCAGCCTGGGGGACAGAGCGAGACTCC
GTCTCAAAAAACAAAACAAAACAAAAACAAAACAAAAACAAAGGGTTCATACAAAGAATTATCCTAAGAAGTGC
ATAGGANNNGGGCGAGGNNN

Inter-79
ACCCTGAATACATGTTATAAAAATTAGCATA<GCAAGAGAATGGCGTGAAC[S>]CCGGGAGGCGGAG
CTTGCAGTGAGCCGAGATTGCGCCACTGCACTCCCGCCTGGGCCACAGAGCGAG...GCTCCCTCTAA
AAGAAAAACAAAAAAAGAAAAGGAAA>TGAAGGAAATGAAGGCTGGG[<S]CATGGTAGCTCATGCCT
GTAA

NNNNNNNNNNGGGCGTGANCCGGGNGGCGGAGCTTGCAGTGAGCCGAGATTGCGCCACTGCACTCCCG
CCTGGGCCACAGAGCGAGGCTCCCTCTAAAAGAAAAACAAAAAAAGAAAAGGAAATGAAGGAAATGAAGGCTGG
GA

INTRACHROMOSOMAL RECOMBINATION
Intra-1
GAACAAGACAGAATTTCAGTTTCTTTTGAAACAAGTCTCAGTCCTTATGGAATCTGAGACAGCAACAG
ATAGCTTTCGTTT<CACATAGCTGTGTCTGGAGCTCCCCACC[S>]GCAGGTGACCCGCCCGCCTCGG
CCTCCCAAAGTGCTAGGATGACAGGCGTGAGGCACC...ACGCCTGGCC>CAAAAATTCTTATTTTGA
CTATTCATGTCTTATTTTTGCACTTTTTCACAAAGCAAGGTCA[<S]AAAAGCAAAGGAAA

NNNNNNNNNNNNNNNNGCAGGTGANCCGCCCGCCTCGGCCTCCCAAAGTGCTAGGATGACAGGCGTGA
GGCACCACGCCTGGCCCAAAAATTCTTATTTTGACTATTCATGTCTTATTTTTGCACTTTTTCACAAAGCAAGG
TCAAA

Intra-2
TGGTTTCCAAACTTGGGTCCACTAGTAGTGCCCATTTGGTCACATAAAAA[S>]TCCCCTGTG<GGAG
GGCGCCTGTGGTCCCAGCTACTCTGGAGGCTGAGGCAGGAGAATGGCGTGAACCCGGGAGGCGGAGCT
TGCAGGGAGCCGAGATCGCACCACTGCACTCCAGCCTGGGGGACAGAGC...CAGACCCTGTCTCAAA
AACAAACAAACAAATAAAA>TTTATGTTTTATAGCTGGTGGACAAGAAGACT[<S]GGTAATATCAAA
GACCAAAAAT

NNNNNNNNNNNNTCCCCTGTGGGAGGGCGCCTGTGGTCCCAGCTACTCTGGAGGCTGAGGCAGGAGAA
TGGCGTGAACCCGGGAGGCGGAGCTTGCAGGGAGCCGAGATCGCACCACTGCACTCCAGCCTGGGGGACAGAGC
CAGACCCCGTCTCAACAACAAACAAACAAATAAAATTTATGTTTTATAGCTGGTGNCCAAGAAGACTN

Intra-3
CTTTTTGCCACCTTTCATAAGGCACTTGTTCTTAACAACCGCCATGGTGCTGCTCAGAGTACCAAAAG
GCTAAA<TTTTGTA[S>]TTTTTGGTAGAGACAGGGCTTCACCATGTTGGCCAGGCTGGTCTTCAACT
CCTGAC...CTCGTGATCTGCCTGCCTCAGCCTCCCAAAGAGCTGTGACCCACCGCGCTTAGC>GCTG
CATGTTCTCTTTTTACAATAAATTCATGACCATCGGAAGATACCAGTTTGACATACATGGCATCAGGA
CCTTCAC[<S]AGCCACCACAG

NNNNNNNNTTTTTGGTAGAGACAGGGCTTCACCATGTTGGCCAGGCTGGTCTTCAACTCCTGACCTCG
TGATCTGCCTGCCTCAGCCTCCCAAAGTGCTGTGACCCACCGCGCTTAGCGCTGCATGTTCTCTTTTTACAATA
AATTCATGACCATCGGAAGATACCAGTTTGACATACATGGCATNNNACCTTCACAA

Intra-4
AGTTTTATGTAATTTCCTGGAGGTGTATTATATTCACAAT[S>]AAAAAGGGTTCCAAAGTG<GGGTA
CAGTGGTGCATTCCTGTAGTCAGAACAACACAGGAGACTGAGACAGGAGGATCGCTTGAGCCCACCTC
GGCCTCCCAAAGTGCTGGATTACAGGTGTGAGCCACCGTGCC...TGGCC>AACCTTCTTAACTCTTA
CACTATGCCCATGTGAGATCA[<S]AATAAATCCCCTTAAAAGCTA

NNNNNNNNNNANAAGGGTTCNAAGTGGGGTACAGTGGTGCATTCCTGTAGTCAGAACAACACAGGAGA
CTGAGACAGGAGGATCGCTTGAGCCCACCTCGGCCTCCCAAAGTGCTGGATTACAGGTGTGAGCCACCGTGCCT
GGCCAACCTTCTTAACTCTTACACTATGCCCATGTGAGATCAA

Intra-5
ATGTTTTTTCTAAATCTCAGAAATAAAGGGCAGTTAAAATTTAGTAGTCTTAAGCTCTTGAAATAAAA
TGGTATCTCT<CCACCATGGCACA[S>]CGTTTACC...TATGTAACTAACCTGCACAATGTGCACAT
GTACCCTAAAACTTAAAGTATAATAAAAAAAAA>CCTCATATTAACCCTAATTTATTTGCAGGGAGTT
GTGCAAA[<S]ATGATCTTCTGACCCAAAAGTTTTACTTAATGATTTTTCTCTTTTATTTCCAGTTTT

NNNNNNCGTTNNCCTATGTAACTAACCTGCACAATGTGCACATGTACCCTAAAACTTAAAGTATAATA
AAAAAAAACCTCATATTAACCCTAATTTATTTGCAGGGAGTTGTGCAAAANGA

Intra-6
GGCTTCCATGTCTTGAGTTAAGAAAAAAATTCAAATTAGCTATCGTGCTAGAGATA[S>]CTGTATAG
AATAAAATTGAATATAGGGCAAAAATT<GCATGGCACATGTATACATATGTAACTAACCTGCACATT.
..CTGCACATGTATCCTGGAACTTAAAGTACAATTAGAAAAAA>TAATAATAATAACAAACAAACCAC
TGTGGGGATACTGTAGTATTTTCCAGCCCCTGGTGGTGGGTTTTGGTG[<S]

NNNNNNNNNCGTGCTAGANANCTGTATAGAATAAAATTGAATATAGGGCAAAAATTGCATGGCACATG
TATACATATGTAACTAACCTGCACATTCTGCACATGTATCCTGGAACTTAAAGTACAATTAGAAAAAATAATAA
TAATAACAAACAAACCACTGTGGGGATACTGTAGTATTTTCCAGCCCCTGGTGNGGGTTTTGGTGN

Intra-7
CAGTTTTATGAAGCTAACAAATTGTATGCTCCAGGTCAAGCTGTTAGGGTATTCAG[S>]CTCTTTTG
CAAACTGCTTTTAGAAAGAGAAC<CCGACATGGCACATGTATA...TCTATGTAACAAACCTGCACGT
TCTGCACATGTATCCCAGAACTTAAAGTAAAATAATAATAATAAAA>GGACCAGAGGTGGGATATCAA
AAAGAAGGGTGAGATGACCTTGTTAATGGGGGCTAAAGGGAGTAGCTGGGCAGACAGAGGACCCAAGC
AGAAA[<S]GGGCTG

NNNNNNNTNCNCTCTTTTGCAACTGCTTTTAGAAAGAGAACCCGACATGGCACATGTATATCTATGTA
ACAAACCTGCACGTTCTGCACATGTATCCCAGAACTTAAAGTAAAATAATAATAATAAAAGGACCAGAGGTGGG
ATATCAAAAAGAAGGGTGAGATGACCTTGTTAATGGGGGCTAAAGGGAGTAGCTGGGCAGACAGAGGANNAAGC
AGAAAA

Intra-8
TACCATGAAATTAATGCCTTCCAGTCAGGCCATGTTCTGGGCCCTAAAGAGAGAGAT<AAATCT[S>]
AGCCGGGC...GTGGTGGTGGGTGTCTGTAATCCCAGCTACTCAGGAGGCTGAGGTAGAGAACTGCTT
GAACCCAGGAGGTGGAGGTTGCAGTGAGGCGAGATCGCGCAAATGCACTCCAGGCTGGGGGCCAGAAT
GAGACTCCATATAAAAAAAAAAAAAAAAAAAAAA>TTAGATCTGGAGGCATTGTT[<S]ATAAGTTATT
AACCACAT

NNNNNNNNNAGCCGGGCGTGGTGGTGGGTGTCTGTAATCCCAGCTACTCAGGAGGCTGAGGTAGAGAA
CTGCTTGAACCCAGGAGGTGGAGGTTGCAGTGAGCCGAGATCGCGCCACTGCACTCCAGGCTGGGGGCCAGAAT
GAGACTCCATCTCAAAAAAAAAGAAAAAAAAAATTAGATCTGGAGGCATTGTTANN

Intra-12
ACGTGGTGAAAATTTGGGCGACACACAAGGTTATTAATTTTGAGGTGGAGTGGATGGA[S>]ATAGAG
AGGGGCGATATACTCGGGGT<TAGAGATGAGGTTTAATCATGTTAGCCAGGATGGTCTCAATCTCCTG
ACCTTGTTATCCGCCTGCCTCAGCCTCCCAAAGGGCTGGGATTAC...AGGCGTGACCAAACACACCT
GGCC>ATGGCAAGAGTTTTCTTACCTGCCATAAATTAGTGGTTCTTAGGCTGTGGTC[<S]CCCTGGA
GAGGACCATCAGCA

NNNNNNGNNNNTGNNGNATAGAGAGGGGCGATATAGTCGGGGTTAGAGATGAGGTTTCATCATGTTAG
CCAGGATGGTCTCAATCTCCTGACCTTGTGATCCGCCTGCCTCAGCCTCCCAAAGTGCTGGGATTACAGGCGTG
ACCCAACACACCTGGCCATGGCAAGAGTTTTCTTACCTGCCATAAATTAGTGGTTCTTAGGCTGTGGTCNNNN

Intra-13 (Duplicate of Intra-8, not counted in final results)
TACCATGAAATTAATGCCTTCCAGTCAGGCCATGTTCTGGGCCCTAAAGAGAGAGAT<AAAT[S>]CT
AGCCGGGC...GTGGTGGTGGGTGTCTGTAATCCCAGCTACTCAGGAGGCTGAGGTAGAGAACTGCTT
GAACCCAGGAGGTGGAGGTTGCAGTGAGGCGAGATCGCGCAAATGCACTCCAGGCTGGGGGCCAGAAT
GAGACTCCATATAAAAAAAAAAAAAAAAAAAAAA>TTAGATCTGGAGGCATTGTTATAAGTTATTAACC
[<S]ACAT

NNNNNNNNNNCTAGCCGGGCGTGGTGGTGGGTGTCTGTAATCCCAGCTACTCAGGAGGCTGAGGTAGA
GAACTGCTTGAACCCAGGAGGTGGAGGTTGCAGTGAGCCGAGATCGCGCCACTGCACTCCAGGCTGGGGGCCAG
AATGAGACTCCATCTCAAAAAAAAAGAAAAAAAAAATTAGATCTGGAGGCATTGTTATAATNATTAACCANN

Intra-14
GGAGATACTGTTATTTTACTGAAATAATTAACTTGAAATAGACTAGTAAACAGAAGGTGGCTGGCTGA
TAACCTCAC<TTTTTTTTTCTTTTTTATTTTAGACA[S>]GTCTCGCTTTGTGGCCCAGGCTGGAGTG
CAGTGGTGGGATCTCGGCTCACTGCAA...GCTCCGCCTCCCAGGTTCACGCCATTCTCCTGCCTCAG
CCTCCCGAGTAGCTGGGACTACAGGCCCCC>TCTGTCTAGCTATTCATCTGTGGACAAGAAAGCT[<S
]TGAGTATGCCTG

NNNNNNNCNNNNANTNANAAGTCTCGCTTTGTGGCCCAGGCTGGAGTGCAGTGGTGGGATCTCGGCTC

ACTGCAAGCTCCGCCTCCCAGGTTCACGCCATTCTCCTGCCTCAGCCTCCCGAGTAGCTGGGACTACAGGCCCC
CTCTGTCTAGCTATTCATCTGTGGACAAGAAAGCTAAA

Intra-15
AAAAAAGTACTATCTCTTTAAAAAGTACTGGCTCAAAAAAATAATAGAAAATAAGAAACAAAAAACAG
GCTGAGAAAAGTGGCTCACATC<TGTTGG...TCAGGCTGGCCTCAAACTCCTGACCTCAGGTGATCC
ACCCGCCTCAGCCTCCCAAAGTGCTGGGATTACAGGCTTGAGCCACCGTGCCCGGC>AAAAGAAATTT
CTTTAGCCCAGGTTTGTGA[<S]AAGAGCCTTCTCTAATTTTCCAGGATGGGGGTGATAAGATCAA

NNNNNNNNNCNNNNNTCCTGNCCTCAGGNNATCCACCCGCCTCAGCCTCCCAAAGTGCTGGGATTACA
GGCTTGAGCCACCGTGCCCGGCAAAAGAAATTTCTTTAGCCCAGGTTTGTGAA

Intra-16
AACTTTTCTTGTGTTACCTTGGTAACAGTAATCGGATTTGCCCCAATGTTCT<TTTATTTTTTATTAT
T[S>]ATATTTTAAGTTCTAAGGTACATGTGCACAACGGGCAGGNTTGTTACATATGCATACATGTGC
CATGTTGGTGTGCTGCACCCATTAACTCG...TCA>AAAGTGAAGTGATCTTCAAATTCCATGTTATC
GATTTCCC[<S]ACTATGATTATGTGTCATAGTAGAAAAATTTTAAAGTTGCTAAAGAGTAATCTAAA
C

NNNNNNNNTNNNNATNATNATATTTTAAGTTCTAAGGTACATGTGCACAACGGGCAGGTTTGTTACAT
ATGCATACATGTGCCATGTTGGTGTGCTGCACCCATTAACTCGTCAAAAGTGAAGTGATCTTCAAATTCCATGT
TATCGATTTCCCANNNNN

Intra-17
AAAGCTCTTAGAACTTCAGGCTGCTGAAGGCATTTTTACTTA<TTTATTTATTTTTATTAT[S>]ACC
TTAAGTTCTAGAGTACATGTGCACAACG...TGCAGGTTAGTTACATATGTATACATGTGCCATGCTG
ATGTGCTGCACCCACTAACTCATCATCTAGCATTAGGTATATCTCCCAATGCTATCCCTCCCCCCTCC
CCC>TGGGATTTCTTATTTTATGGTAATTGTCACCATATTAGTTTCTCCTACCCA[<S]AAGAGGAGG
TAGAGATGAAGTCCCATTAATGGT

NNNNTNNANNANACCTTAAGTTCTAGAGTACATGTGCACAACGTGCAGGTTAGTTACATATGTATACA
TGTGCCATGCTGATGTGCTGCACCCACTAACTCATCATCTAGCATTAGGTATATCTCCCAATGCTATCCCTCCC
CCCTCCCCCTGGGATTTCTTATTTTATGGTAATTGTCACCATATTATTTCTCCTACCANN

Intra-18
ATTAACATTTTTGAGCAATATAGTTGACCCTTGAGCAACACAGGTTTT[S>]TTTATTTATTTATTTA
<TTTTTTATTATTATTATACTTTAAGTTTTAGGGTACATGTGCACAATGTGCAGGTTAGTTACATATG
TATACATGTGCCATGC...TGGTGTGCTGCACCCAGTTGTTGGTTTTTTTTGTTTGTTTGTTT>TAAA
CTTACATCTTTGTGTCAACCTGGA[<S]CTTCCTGGACTCTGATCTGAGGCTAAAATGATAGTTTTAT

NNNNNNNNNNNNNATTTATTTATTTTTTATTATTATTATACTTTAAGTTTTAGGGTACATGTGCACAA
TGTGCAGGTTAGTTACATATGTATACATGTGCCATGCTGGTGTGCTGCACCCAGTTGTTGGTTTTTTTTGTTTG
TTTGTTTTAAACTTACATCTTTGTGTCAACCTGGAA

Intra-19
AATTAAGGAGGACATGAGATAACATCTTCTCCTCAGACAATGGGAACCAGTTTTTTAAAA<TTTTTTA
AATTGTACTTTAAGTTCTAGGGTACATGTGCACCATGTGCAGGTG...TGTTACATATGTACACATGT
GCCATGTTGGTGTGCTG>GGTATTTTTACATCTGGGTATGAAGCTGA[<S]TTCTATCTCTACCTTCC
T

NNNNTNGGANNN[S>]TTTTTTAAATTTTTTAAATTGTACTTTAAGTTCTAGGGTACATGTGCACCAT
GTGCAGGTGTGTTACATATGTATACATGTGCCATGTTGGTGTGCTGGGTATTTTTACATCTGGGTATGAAGCTG
AA

Intra-20
AACACAAAAAAGGACAACCAGACAACAAGTAGCTGAGTTACAAAG<CAGCTACT[S>]CGGGAGGCTG
AGACAGAAAAATCACTTGAGCCCAGGAGG...CGGAGGTTGCACCACTACACTCCAGCCTGGACGACA
GAGCAAGACTCCATCTCAAACAAACAAGCAAAACAAACAAAAGCAAAAGAAAAAAAAAA>AACGCCTA
CTCTGTGCAGGACACCATGCAAGGGA[<S]CAGTGGAGG

NNNNNNNNNNCGGGAGGCTGAGACAGAAAAATCACTTGAGCCCAGGAGGCGGAGGTTGCACCACTACA
CTCCAGCCTGGACGACAGAGCAAGACTCCATCTCAAACAAACAAGCAAAACAAACAACAGCAAAAGAAAAAAAA
AAAACGCCTACTCTGGGCAGGACACCATGCAAGNNNAAGGGGAGGG

Intra-21
TGTCCTGCACACAATCCTAGTGGTTTCCATTTCTTTTCTGAGCAG<T[S>]TTTTGTTTTTTTTTTTG
AGACAGAGTCTCACTCTGTTGCCCAGGCTGGAGTGCCGTGGCACAATCTCGGTTCACAGCAACCTCTG
CCTCCCGTGTTCAAGCCATTCTCCTGTCTCAGCCTCCCAAGTAGCTGGGATTACAGGCATGTA...CC
ACCACGCCTGGC>CACATCTGGTCTTTGCTTCAGCAGCAGCTTCTCTCCTAGGCAC[<S]AGTCTGCT
CCCTCC

GNNNTNNNNNNNANNNNTTTTGTTTTTTTTTTTGAAACAAAGTCTCACTCTGTTGCCCAGGCTGGAGT
GCCGTGGCACAATCTCGGTTCACAGCAACCTCTGCCTCCCGTGTTCAAGCCATTCTCCTGTCTCAGCCTCCCAA
GTAGCTGGGATTACAGGCATGTACCACCACGCCTGGCCACATCTGGTCTTTGCTTCAGCAGCAGCTTCTCNCCT
AGGCACAAN

Intra-22
TCCTAATGCTCAGCAGCAAATTTTTTCTATTGCTTAGCCCCAGGCCCCCTCACAATGT<GAGGT[S>]
TTCGTCACATTGGCCAGGCTGGTCTTGAACTCCTGACATCAAGTGATCCGCCCACCTTGGGCTCCCAA
AGTGCTAGGATTACAAGCATA...AGCCACCACGCCCAGCCTCTTTCTTCTTTTATT>GCAGACCTCT
GGGATTCAGTTTACCATGGCCCTTCTCTCAACA[<S]GCAGCCA

NNNNNGANGNTTCGTCNTTGGCCAGGCTGGTCTTGAACTCCTGACATCAAGTGATCCGCCCACCTTGG
GCTCCCAAAGTGCTAGGATTACAAGCATAAGCCACCACGCCCAGCCTCTTTCTTCTTTTATTGCAGACCTCTGG
GATTCAGTTTACCATGGCCCTTCTCTCAACA

Intra-23
TCCTTCCTAGATGTGATAGCTTGCCAGGATAAGTTGGCAAAC<TTTTTT...G[S>]GTTTTTTTTTT
TTGAGACGGAGTCTTGCTCTGTAACCCAGGCTGGAGTGCAGTGGTGCGATCTCGGCTCACTGCAAGCT
CCGCCTCCCAGGTTCACGCCATTCTCCTGCCTCAGCCTCCCAAGTAGCTGGGAC>CATCTGAGACTGT
TTTGACAAACAGAGGCGACCACTTGA[<S]GAGA

NNNNNTNNTNGTTTTTTTTTTTTTGAGACGGAGTCTTGCTCTGTAACCCAGGCTGGAGTGCAGTGGTGC
GATCTCGGCTCACTGCAAGCTCCGCCTCCCAGGTTCACGCCATTCTCCTGCCTCAGCCTCCCAAGTAGCTGGGA
CCATCTGAAACTGTTTTGACAAACAAAGGCGACCACTTGAAAAA

Intra-24
GTGCTTCCCTCCAGAGAAGGTTTGCTTACTGTTCAGTATAATTTGGATAGTGTTTCTTTTTTTTTA[S
>]G<TTTTATTATTATTCTACCGTAAGTTTTAGGGTACATGTGCACAATGTGCAG...GTTTGTTACA
TATATATACATGTGCCATGCTGGTTTG>TAGGTTGTTTTTTATTTTGACATGTATCTTGTGGAAAGTG
[<S]C

NNNNNNNTTGNANNNGTTTCTTTTTTTTTAGTTTTATTATTATTCTACCGTAAGTTTTAGGGTACATG
TGCACAATGTGCAGGTTTGTTACATATATATACATGTGCCATGCTGGTTTGTAGGTTGTTTTTTATTTTGACAT
GTATNNNTGGAAAGTGNN

Intra-25
AAAAAAGTCATGGATGTCTGGTACACATAATTTGATTAATATTA[S>]TTTCTTTGTTTATGAAACAT
ATTAATCTATTAACCATCTTATACCCATTTGTTATCTATTTAG<TTTTTTTT...TTATTATACTTTA
AGTTTTAGGGTACATGTGCACAATGTGCAGGTTAGTTACATATGTATACATGTGTC>TAGGCCTCTTT
[<S]CTTAGACATTTTGGGTTAGTGGGTCTGGCACAAAGTGCTGAAATCTTAATTTTTATCAAGATCC
ATAGTTTAGTCTACTGGGTA

NNNNNNNNNNNTTTCTTTGTTTATGAAACATATTAATCTATTAACCATCTTATACCCATTTGTTATCT
ATTTAGTTTTTTTTTTTATTATACTTTAAGTTTTAGGGTACATGTGCACAATGTGCAGGTTAGTTACATATGTAT
ACATGTGTCTAGGCCTCTTTNNNNNN

Intra-26

ATGGTAAATTAGGAGGACATGCCAGCACCTCCTTGGCTAAGTAGTAAGAAAGTACCAAGAAAGGAGCC
AGAAGAGTCAGAGAGATTGAAATGTTAAA[S>]GTGGATTTCTCAGTAAGAACAGAAAACCC<TGATA
...GGTACAGCACACCAACATGGAACATGTATATGTATGTAACAAACCTGCACATTGTGCACATGTAC
CCTAGAACTTCAAGTATAATTTAAAAAATGAAAAAAAA>AAACAAGAAGTTGAGATTTTAAAGGAATT
AGAAATGGGA[<S]TGAATCAGATACAC

NNNNNNNNTNATGNTAAGTGGATTTCTCAGTAAGAACAGAAAACCCTGATAGGTACAGCACACCAACA
TGGCACATGTATATGTATGTAACAAACCTGCACATTGTGCACATGTACCCTAGAACTTCAAGTATAATTTAAAA
AATGAAAAAAAAAAACAAGAAGTTGAAATTTTAAAGGAATTAAAAATGGGANNANCAAATACACAN

Intra-28
CTTCTTGTTCATTCATTCTTGATGACTTTAATGTAACAGCATTCA<TTTAT...TTTAAGTT[S>]TT
AGGGTACATGTGCACATTGTGCAGGTTAGTTACATATGTATACATGTGCCATGCTGGTGCGCTGCACC
CACTAACTCGTCA>GTATTGCTTCTATAGAAGGCTAGCCCAAATGCAGAGAGGAA[<S]AATG

NNNNNNNNNTTNNGGNNANTGTGCACATTGTGCAGGTTAGTTACATATGTATACATGTGCCATGCTGG
TGCGCTGCACCCACTAACTCGTCAGTATTGCTTCTATAGAAGGCTAGCCCAAATGCAGAGAGGAAA

Intra-29
TTTTCTGCTCACAGAGTTGGGAGTCTATGCAATAGCATCTTACACAATCATTAGCAGAAATTCG[S>]
ACTCCGTAAATTTT<TTTTTAAATTTTTTTATTACACTTTTAAGTTTTAGGGTACAGGTGCACAATGT
GCAGGTT...TGTTACATATGTATACATGTGCCATGTTAGTGTGATGCACCCATTAACTCGTCATTTA
CATTAAGGAAATACCCTA>TCATTGATTCTAATTCCAGTAGCCTCTTGGACTTCTACACTTACA[<S]
TTTTCTTATACCTCTG

NNNNACTCNNANTTTTTTTTAATTTTTTTATTACACTTTTAAGTTTTAGGGTACAGGTGCACAATGTG
CAGGTTTGTTACATATGTATACATGTGCCATGTTAGTGTGATGCACCCATTAACTCGTCATTTACATTAAGGAA
ATACCCTATCATTGATTCTAATTCCAGTAGCCTCTTGGACTTCTACACTTACAA

Intra-30
GGCTGCAATCTGTCACCCTGCAGCACACTGAACTAGTTTTTGCGGTGCTCAGGCTTAGAGCTCCCTGA
TCACCTCTTCAGAAGGTTTCATTTAA<TTTTTTTTATAGACTCAA[S>]GATCTCACTATGTTGCTCA
GGCTGGTCTTGAACTCCTGGGCTCACGTCACCCTCCTGCCTCGGCCTCCCAAAGTGCAAGGATTATAG
GCATGAGCCACCACACCCG...GCC>AATATACATTATTTATACTGCGTACCACATAGGCA[<S]CGT
CTCGCCCG

NNTNTNTNNNACTCNGATCTCACTATGTTGCTCAGGCTGGTCTTGAACTCCTGAGCTCAAGTGACCCT
CCTGCCTCGGCCTCCCAAAGTGCAAGGATTATAGGCATGAGCCACCACACCCGGCCAATATACATTATTTATAC
TGCGTACCACATAGGCA

PCR VALIDATION OF PUTATIVE SOMATIC NAHR EVENTS FROM GENOMIC DNA
SANGER SEQUENCES FOR 6 POSITIVE TARGETS

>INTER-3
NNNNNNNNNNNNNNNNNNNNANNNGGGCCAGGCGCGGTGGCTCACACCTGTAATCCCAGCACTTTGGG
AGGCCAAGGCAG
GCGGATCACCTGAGGTCAGGAGTTCGAGCCGGAGTATGAGCTTTCATATTAAAAGAAAGTCTGAGTGT
GGTGG

>INTRA-8
GGNNNNGNNNNNNTNNNNNNNGGTATCCAGCTACTCAGGAGGCTGAGGTAGGAGAACTGCTTGAACCC
AGGAGGTGGAGG
TTGCAGTGAGCCGAGATCGCGCCACTGCACTCCAGGCTGGGGGCCAGAATGAGACTCCATCTCAAAAA
AAAAGAAAAAAA
AAATTAAATCTGGAGGCATTGTTAAN

>INTRA-13
ANNNNNNNNTTTTTTTTNAGANGGAGTCTCATTCTGGCCCCCAGCCTGGAGTGCAGTGGCGCGATCTC

GGCTCACTGCAA
CCTCCACCTCCTGGGTTCAAGCAGTTCTCTACCTCAGCCTCCTGAATAACTGGGATTACAGACGCCCC
CCACCCCCCCCG
GGTAAATTTATTTCTTTTTTTAAGGCCCAAAAA

>INTRA-15
NNNCNNNNNNNNNNNGACCTCNGTGACCTATGTGCCTTGGCCTCCCGACGTGCTGGGATTACAGGCTT
GAGCCACCGTGC
CCGGCAAAAGAAATTTCTTTAGCCCAGGTTTGTGA

>INTRA-23
NNNNNNNNNNNNNTTTTTTTGAGAAGGGCTTCACTCTGTTGCCCAGGCTGGAGTGCAGTGGCGTGATC
TCTGCTCATTGC
AACCTCCGCCTCCTGGGTTCAAGCAATTCTCCTGCCTCAGCCTCCCAAGTAGCTGGGACCATCTGAGA
CTGTTTTGACAA
ACAGAGGCGACA

>INTRA-29
NNNNNNNNNNNNNTTTTNATTACNCTTTTAAGTTTTAGGGTACAGGTGCACAATGTGCAGGTTTGTTA
CATATGTATACA
TGTGCCATGTTAGTGTGATGCACCCATTAACTCGTCATTTACATTAAGGAAATACCCTATCATTGATT
CTAATTCCAGTAGCCTCTTGGA

PCR VALIDATION FROM GENOMIC DNA OF PUTATIVE POLYMORPHIC NAHR EVENTS
DETECTED IN CAPTURE-SEQ and PROMETHION WGS LIBRARIES
SANGER SEQUENCES FOR 16 POSITIVE TARGETS

Repeat elements sequences are in lower case
[xxx] indicates the junction between the recombined repeat elements

>POLY-INTER-1
CAGACTGCATAAACTCTAATAACTTACCTTTCCATGGAAAGAGAATATGT
GGGTATTCTCTTTGAAGACAAATGGTGGAGAGTAGACTCTCATCCAAGAC
CCTCTGGCAGTAATGAGTTGCATTCCAGAAGGTTCCAGAAGGTTCTGTGG
AAGGCTGCAGTTTGGGATTGGTTCTCAGGTGTCTGTCTAACTGCTTTCTA
AGGATGAAGACggccgggcacggtggctcacgcctataatcccagcactt
tgggaggccaagacgggcggatcacgaggtcaggagatcgagaccatcct
tgttaacactgtgaaacctc
[xxx]
gtctctactaaaaatacaaaaaattagccaggcgtggtggttggcgcctg
tagtcccagctcctccagaggctgaggcaggagaatggcatgaacccggg
aggcggggcttgcagtgagccgagatcgcaccactgcactccagcctggg
tgacagagtgagactccgtctcaaaaaaaaaaaggatgaaaaCTCCTGGT
TCGTGCTTTAGCTACATCCACAGGAGCAACAGACTAAGACATTCTTCCAC
ACAATAACAGACCCAGGCAGTGCAGAGCCATGTGGGTCTTGTCTGTCCTG
TTGTCCACAGTCTGGTTTTATCTTAGACCTAGCCTGGTGCAATCCATGTT
GTGTATTGGCACTCCACTTAAGTTTCCTGTTTGTAAAAAGCCCCTTTCAT

>POLY-INTRA-3
TTTTAAAGTCTCATAACGATGCAGAAATACACGTGTACGAGACTTGCTCT
GACAGCTAACACAGAACTGACAGGAGTGGCTCCCCAGTGAGGCTGGAAGG
AGGGTGGAGGATGTGACAGGGAAAGGAGATGTCTTATGGCCAAGGGTCTA
AGCTGCAGCTTTGCTTGGAGTTTTACCACAAAACAAATATATATATCATG
TGATGGTGAAATATAACCATTTATATTTAATAATAACTGGTGGggctggg
catgatggcttgcgcttgtaatcccaacacactggggggccgaggcgggc
agaccacctgaggttaggaggttgaggccagcctggccagcatgctgaaa
cgctgtttctactaaaaatacaaaaaaattagctgggcgtggtggtgcac
acctataaccccagctactaggtgggctgaggcaggagaatca

[xxx]
cttcaacctgggagacagaggttgtagtaagctgagatcatgccactgca
ctgcagcctgggcaacaaagcaagactccgtttcagaaaataataataat
aataataataataataataataataaAACGTGGAATGAGAATACAGCTTT
ATCCTCTTAGATAAAAAAGCAGTACTCTCCAATATTAGAGAAAAATTGCA
ACCACTTTTACCAAAAACACCTGTTTCACAAGCCTCTCCACAGTAACACC
ATAGTCTCCGTGAGCTCGATGTGTTAAGAATGGTTGAATGACTCTCCTGC
TATTATTTACATTGATTTCATGTTCTTAAAATAATGAAGGAATA

>POLY-INTRA-5
TTTAGTATAGCAGTCAAAGTATTAATTTCTCACATTGCAATTTCCTTCAA
AGACATGAATACAACCTTTCTAATGACTCCTTGTTTATCAAGATACCTCT
TCAAATTATTCTATTTATTTCATTCAGTATATTATCTGTGTATACCGATA
TGATATTACACttttttttttgagatggaatctcattctgttactgatgc
tggagtgaggaggcatgatctcggttcactgcaacctccacctcccaggt
tcaagcgattctcctgtctcagccccacgggtagctaggactacaggtgc
acaccaccatg
[xxx]
cctggctaattttttgtatttttagcagagacggggtttcaccatgttggc
caggctggtctcgaactcctgaccttaggagatccacctgcctcggcctc
ccaaagtgctgggattacaggcatgagccactgcgcctggccTCTCTTCT
TACATATTTCTAGAACTCCTCTAGAATTTGGGGTTTGTTTTTCTTAATTA
CAAGGAATCAAGTTGAATCATTAGTGCATATATAAATATACATTTTATTT
TTAGTACACATTATATACCTCAGGAATgtacaatgctcagtgcctgggtg
acgggatgattcataccccaaacctcagcaacgtacaatatcctcaggtc
acaaagctgcccgtggatcccctgaatct

>POLY-INTRA-6
CCTCTAGGTGATTTAGCTTAGTTTTCTAGTATAATTATCCTTAACAACTT
ATATAATTCTTTTATTAAAAAATATTTATAGTTATACAGTATTTGTGGGT
TTTATTAAGAAGAATTACAAATTCTTCTTGGGAAAAGGATATATAGATAA
GGTTTTTGAGGGAGCAAGTCTAGCATATGGTCAATAAATATCTAAATGAA
GTCCTGAGtttagctctcctgttgcctctctgactttctctcctattact
ctacttcttgctcactccaatataaccatattaaccttactgtttctcca
acatatgagaatatctgcacctcaaggcctttgcttttgttcctctgcat
ggaaatggtggtctcctagacatctgcataatttttgtccaacgtttctt
gcagttttggctcaggtactacattctcagtgaggctttctctaattagt
tctcccttcgctcctgaattcttttcctggcctagtttttgcacgatctc
agctcactgcaacctctgcctcccgggttcaagtgattctcctgcttcag
cctcttgagtagatgggactacaggcacgcaccaccatgcccagctaatt
tttgtatttttagtagagacggggtttcaccatgttggccagatggtctc
atctcttgacctcgtgatccacctgccttggcctcccaaagtgctgggat
tacagg
[xxx]
cgtgagccaccatgctcagccAATATTAGCATATTCATAGTTCATATTTT
AGGACTAATCTTGGAACTAAtcagtactatgtatttcaaatatcttctcc
cattatgtgacttgtttcattttgtgtatgtttttatgcagaaaattt
ttaaagttgaatataatatacta

>POLY-INTRA-7 (Sanger sequences not long enough to reach the
recombination breakpoint, but sequences confirmed otherwise)
aggaggctgaggcaggagaatctcttgaacccaggaggcagaggttgcag
tgagcagagatcatgccactgcactccagcctgggcgacagagccagact
ctatctcaaaaaaaaaagaaagaaaagaaaagaaaagaaatacctgaggc
tgggtaattgataagaaaagaggcttaattggctcatggctctgcaggct
gtgcaggaagcataccgccagcttctgcttctggcgaggcttcaggaagc
ttccaatcacggcagaaggcaaagagggagcaggcacatcacatggcgaa
ggtgggagcaagaaggcgggggggtgccacacactttttaaaaccttatca
ggggagaactcataatagggaggacagcaccaaggccgtggtgctaaacc

attcatgagaaatccatcctcatgatccaataacctcccaccaggcccca
cctccaatattgggaattacatttcaacgtaaattttgacgaggacataa
atacaaactatattaCTACCCTTAAACAATTGGAAATTGCTTTGTTTTTG
AGAAATAAAAGAATGCTGAATTCTGTTCTGTTGCTTTGGTTGTACACTTT
AGTGAAAATTGAAAGGTTATGTTAATTTCTGAAGAATTATAATGATAAAA
AACGGTGCAGACACAACCTGCACTTTGCAAGCATTGCAGACTTAAGTTTT
TTGGGGGAAACTATACATAATTAAATAGTTTCCCAGTGAGCTCAGCCAGC
CCATTCAGGGCAGTGCCCAGACATATGCTTGTGGCAAACTCTAGCCAAGC
AACTTACCTACAGGAGAAAAAAAGACTCACTGTTTCAAGAGGAGCTGTGG
TGCTTCTCTCCTTTCTCCTCTTTTTCTTGTATAAACCCTACATTCACCTG
TATTCAACAAAATCAGCCAACTAATTGGTTTCTGTGTGTGTATTCAAGAA
(...)
gagtagctgggactacaggcgcccgccaccgcgccctaatttttttgtatt
tttagtagagacggggtttcaccatgttggtcaggctggtctcgatctcc
tgacctcgtgatccgcccgcctcggcctcccaaagtgctgggattacagg
cgtgagcccccgcgcccggccGGGATTATCGTGttcttgcggctgagtgg
tgctctgctgcgcatccacctcggtgcctccgcccactcctgcaccgacg
gccgctgcctttgctcgcagatctgggctattgtgaacagcgctgcggta
aacgtggcggcacagacatctctcgggcaaaccgaGGATGCGATACGGAT
ACTTCCTCCGTCATTTCCACTCCCCTTTCTCCAGGCCCTGAAGCTCCGTC
TCCACGCCCGAACTCGTGGCCTCCCATTCCCCAGGCAGCTTCTCCCGAAT
GTCCCCTCTCAGAAACCGGTGCCAGCGCGTCCCCGCTGTGCCACCCCCAA

>POLY-INTRA-9
GCATAGATATACTTTAACCAAGAAGTCGGTGCCTCCATTTACAATCTGACTCTAGTAATAATATATCT
GAAAATACTGATGATGGCTAGCATGCACACAGCTCCACCCtttacaaagcatgccacatgcattgcat
cacttgaacagcaaaccaatgacaaacatgaaagtcaggtcagattatgctcagggaaaagatgagga
aactggggctcaagggaagctaagagacttgctcatggttaaccagagagtaaggtgtcattggggcc
gagcacggtggctcacacctgtaaccccagcacttggagaggcagaggcaggtgaagttcaagaccag
cctggccaacat
[xxx]
ggtgaaaccccgtctctactaaaaatacaaaaattagccgagtatggtgg
catgcacctgtagtcccagctactcaggagaccaaggcaggagaatcgct
tgaacccaggaggtggaggttgcagtgagccgagatcacgccactgcact
ccagcctgggtgacagagtgagactctgtctcaaaaaaataataataaaa
TATAATTTAAAAAAATAAAATAACATTTCCATGCATCCTACCAATGGATG
TGTTTTAATGGGCTGTGGCTGAAAAGTATCCCAAACTGTTTATTTAGAGG
AGTATGTTTAATTTTCATTTATTTCCAGTGACAAAGTTTTATGGACTATA
TTTATCCCAGATATATGCATAAATTCCCTTTTACTAAAGGGGTAGGGATG

>POLY-INTRA-10
AAGCTTCTCTTGTAGACCCGGGTGCATTGCTTGCCAGAGAGATCTCTGAA
AGGTGGTTCTGCTTTCCAGGGAACTATTTAGCCAATATGCATTACCCAGC
TCCTCAACCTTTTTATGATTCCTGCCTGATGCTGCCTTTTGGTGATTTCT
GGTAAAGCATCAGCACCTTATCATCTGCTTTAGTGACCAGCTTGGGAGAG
TAACGTTAAGTCACGACCAGAATATGTCTTGGTGGAGCATGAACAAAGCA
TAGATTTTTTGTGGGGGAAAAAAGGTGTtacttagtttcaaattgccatt
caatgatttgttagtagatttagtgacattgtatacatttctcaggctgt
tagactctctatttttttaatccataaaataagagtaataatagctacctt
acaaggtttttgtgaagattataagaaacacataaaatgtataATatata
taataatataacatgaaacattacgtatagtatgtatcataagttatctt
ttttattgtttgtttttgtttttattttttgagatagagtctcactctgt
ggcccaggctggagtgcagtggcacgatcttggctcactgcaacctccat
ctcccaggttcaagcgattctcctgcctcagcctcctgagtagctgggac
tgcaggtgcccgccaccgcacccagctaattttttgtatttatagtagaga
cggggtttcaccatgttggccaggctggtctcaag
[xxx]
ctcctgacctcgtgatccgccagccttggcctcccaaagtgctggaatta
caggcgtgagccaccgcgcccagccctgagaaaccattttaatgaata

aattgttttccattctttagttgtatcatcagttatttaaccaacctcct
atgtgggacatttgggtagcttctgatattttttccattacactaatggc
tgagatgaatattcttgaacataaatcttgtacataaatttgattattgc
ctaggataaagtcctaaaactaaaattactggctcaaagtgtatgaacac
ttcaaggctgttcatatataaattgccaaataaccaaccctccagaaagc
ttgcaccaatgtatatccccatcagcagcagtgggtgaggggtgagaaca
cccatttctccacatcctcggcagtaaatttccaccaatgagataagcaa
aaactttccatgtttataacttgcatttctttggtggctgtggggtccac
gtctttccttgtttgttgggcatcagcttcggctgtattgtgaatgtcct
atcatattcttcctgtttgtccatCACTATGATTGGGGATGCTGGAATTT
AGGACCCCAAGGAGGCTCTGCCTGACCTTTTCAGCACTTAGCGTTGTTCC
ACTTGTGGTGAGGATGAGGCAGAGTCCATTCTCCATGAAAGAAAGACAGT
TGCTCCCTCGATACTTAGCAAATGTcattccta

>POLY-INTRA-11
tgcttgaacctgggagttggagattgcagtgagctgagatcgcaccactg
cactccagcctgggtgacaaagcgagactccatctcaaaGACACACAAAC
CCTTAAGgcaaggtgcttcaacatataaaaattatgataatatactacat
taacagaatgaagggaaaaaagctacatggtcatctcaattgatgtagca
aaagaattctcgacaaattcagcaccctttcatgacaaaaatacactcaa
caaattagaagtagaagaaaatggccaagcgcgttg
[xxx]
gctcacgcctgtaatcccagcactttgggaggccgaggcgggtggatcac
ctgaggtcaggagttccagaccagcctggccaacatggcgaaaccccatc
tctactaaaaatacaaaaattagccaggtgtggtggctcactcttgtaat
cccagctacgtgggaggctgaggcaggagaatcacttgaacccgggaggc
ggaagttgcagtgagccgagattgcacctctgcactccagcttgggtgac
agagcaagactccgtctcaaaaaaagaagaagaagaaaaaagaaaGTTAT
CTTGGCCCCTGCTGCTCTTGGTTTGCTGACCATGTTTGAACATGATAAAT
TTGAACCAACGTGAGAAATGAAATTTGCTGTCTGCTTACTTCCAAAGCCA
AGAACTGAAGAGTGATTGAACTGCACTGACTGCCTGTCTTAAAATACACA
AAGCAAATGCTACCATGAGTCAGCCAAGATCAAGCCCCGCTCCAGAATGC
AGACTGTTCATCACTGCTCCAGAAACCAGGGAAAGAC

>POLY-INTRA-12
TGATCGTCCTGGTACTTTAGATCTTAAAACATCAAAGAATATAGGATTAA
AATGGATTTTCATGTAGAAAAAGGATGCCAAGAATCCAATTAATATTTTT
TTAATCAAGCAGAATGAGATTTGGGTATAAAATAAATGGAAATTATTTGG
ATGAAAATAGTTAGGCCTGTGCTTCTTGGGACCATCAAAGAAGTCAAATT
AGTGTTTTCGTGACATACCTGGgtggtgagaggacaaacctgaggtgtgc
ccccagaaagcatgcagtgtggctgaaggaatgaaagtgataagagaagc
tgggcgtggtggctcacgcctataatcccagcactttgggaggcggaggt
gagtggatcacctgagttcaggagttcgagaccagcctgaccaacatgga
gaaaccctgtctctactaaaaatacaaaaattagctgggcg
[xxx]
tggtggcacatgcctgtaatcccagctgttcaggaggctgaggcaggaga
atcgcttgaacccgtgaggtggagtttttgatgagccgagatcacgccat
tgcactccagcctggacaacaagagcgaaattccgtctcagaaaaaagaa
aaaaaaaaGATATATAACTTTATTTTTTGCTATTCAGTCTTGAATCTCTC
TGCATTGATTTGAGAGTGGTCCAAATTTTTCTCCAAATCTTAGCCATTAT
TCACCTAGCTTCATTTAATTATAATCCTTAAAGTCCCttgaac

>POLY-INTRA-14
TATTTTAACAAGACTAAACAGTTTTCCAGGTCATCTTTTGAAATCAGTCC
TCTTGCTGCAATAATGGTTCATTTTTATAAAGGAAGTTTACTCCTCTTGT
CCAGTTATTTTGAGACACACAAATCTGACTGGCAAATAATCATGTGGGAT
GCAGGGACAAATAATCACAATGGAAAGAAAATCTAACCTGTGCCTGATAA
ATTTATGTTTCAGGTTGTTAAATACTATCCTATAGAATCCTCAGGAATAT
GATGAAATTAAGACATTTTAGAGGGTGGCACTAATATTCTGTTTCTTCAA

CTTGTCCTTCATTATTAGTTCTTTCCCCACTGGCATGCTTAAATCTTTGT
TCAGGAGCTGAAAGAAATTTGTCTCCAACTTGTCTCATCTAAAAAAGTCA
TTCAATCCTTGTCCCTgtttccatccaccactattgtggaataattctta
acaagcacattaagaaccttatgaaattcaagagtcatatctcaattctt
attttacttctttctatatagcatttaataatattaaagttactcctaca
aaagctgccctttttctaggttctatacaacatactgtccaatgattggca
agccactccttgcccatatctattgagggcttcttcttctctattcatcc
ttggaatagtgttggttcttagaattccttccccagttctcttccctaga
aagtttccctgagtaatttcatctgtttttatgttttcaactaccaacta
gatcctgctgacttccaaatctctctctcaagttcagacttctcttctga
gtgccagggccatataaccagttgactactaagtcatccatgttcatctc
ctccccaccaaacctttttgttctctttgtcagtgactgagtgacatacca
attactgtgataaaagcatggggaactgccaggcgtggtggcttat
[xxx]
gcctgtaatcccagcactttgggaggccaaggcaggcagatcacgaggtc
aggaaattgagaccatcctggctaacacggggaaacaccatctctactaa
aaatacaaaaaaattagccgggcgtggtggcgggcgcctgtagtcccagc
tactcgggaggctgaggcaggagaatggcgtgaacccgggaggcggagct
tgcagtgagccgagatcactccactgcactccagcctgggcgacagagcg
agactccatctcaaaaaaaaaaaaaaatctatataacggaatcatcagtat
gtattcttttgtacttggcttctttcatgcaatgttttgttgttgttgtt
atttgagtcagggtcttgctttatcaccgaggctggagtgcagtggcatg
atcatggctcactgcagcctcaaacccctgggttcaaacgaccctcccac
ctcagcctcctgaggagctgggactatgagcatgggccaccatgccgggc
aactttttacaatttttgtagacatgggctatattgcccaggctggtctt
aaactcctgacctcaagtgatcctcctacctcagcctcccagatcagtag
gatgtcaggtgtgagccaccgcgtccggcccacacaacattatttttgtt
agattTTTTTTCCCAGCATTTACTACATTTTAAACTGTTACAAAGCACTC
CAGTTAAAATGATCACAGTTAGGAAGCTACCCTTTGTGAACTTATTTTTG
GAGCAATGATGACTTGTAGATTTTGAATGCAGCAGATTTCTCATGTTGAA

>POLY-INTRA-15
AAATAACACCCGAAAGGCAAGGACGGGCAGATTGGGGAGGGAAAGGATGT
TGGGCTAAGGGCTGTGAGCTTATGTTACAGGCAACTGAGCCACTGAAGAA
TTTTGACGAAGAAAATGCCAACCAAAGCAGTCATTTTAAAAGTTTATGGC
TGTTCAGTTACAGGACAAGTTGTGaaaagaaagaaaaaaaaatggaaaaaa
aaaaaGTTTATGGCTGAAACAGTGTAATTGATTAAAAAGTGAAAATCCAg
gccgggcgcggtggctcacgcctgtaattccaacactttgggaggctgag
gcaggcagatcacctgaggtcgggagttcgagaccagcctgaccaatatg
gagaaaccccgtctctactaaaaatacaaaagccaggcgtggtgggacat
gcctgtaatcccagctactcgggaggctgaggcaggagaatcg
[xxx]
cttgaacccgggaggcagaggttgcagtgagccgagatcgtgccattgca
ctctggcctgcgggacaagagtgaaactctgtctcaaaacaaaaaaaTTT
GTAAAACTGATCCCACAATTCCCTACATTTGTAACTTTTAAAATTCAGAG
GTAAATACATTTTTATTTAAGGAGGAATAATTCTTCAGAGCAAAATAATC
TGAATATCACCTGGAATAGAGATTGGACAGAAAAAAGTGTGAACAGGGAA

>POLY-INTRA-16
ctgagatcacgcttttgcactccagcctgggtgacaagagtgaaactcca
tctcaaaaaacaaaaaaaaGTAGTCATTTATCTCTGGATACTATTATTTT
GAGCAGTAATAAACAAGTTTATTTGTATATGTATAATTTTAGATATCCCT
TGGCCAGAAGGTGAAGAAAAGTTATCTGATAATGCTCAAAGTGCAGTAGA
AATACTTTTAACCATTGATGATACAAAGAGAGCTGGAATGAAAGGTATGG
TTTTGTGTTAATACATTGTTTTACCATTGATTTTTTGCAGATGGTGAATA
TTTATAAAAATAGCAAGTTCTggctgggcgtggtggctcacgcctg
[xxx]
taatcccagcactttgggaggctgaggcgggtggatcccctggggtcggg
agttcgagaccagcctggccaacatggtgaaaccccatctctactaaaaa

tacaaaaattagccaggcttggtggtacacatctgtaatcccagctactc
aggaggatgaggtgggagaatcgcttgaacccgggaggcagagtttgcag
tgagccgagattgtgccattgcactccagcctaggtgacagacaaaaaaa
aaaaaaaaaaaaaattagccaggcatggtagcactcacctgtagtcccagc
tacttgggaggctgaggcacgagaaccactcaacctgggagatggaggtt
gcagtgagccaagattgcaccactacactccagattgggcactggagaga
gactccatttcaaaaaaaaaaaaaGATGTGAGTgccgggcatggtgactc
acacctgtaatcccagcactttggaaggccaaggtgggcagatcacctga
ggtcaggagttcgagaccaacctgcccaacatggcgaaaccccgtctcta
ctaaaaatacaaaattagcctggtgtggtggcacacacctgtaatcccag
ctactcgggaggctgaggtaggagaatcgcttgaacccgggaggcggcgg
ttgcagtgagcagagatcgcgccactgcacttcagcctgggcaacggagc
gagacctcgtctcaaaaaaaaaGCTGTGAGAAAGATAGGCTTCTAAGTTA
AGGCAAATCATTCATTCTGTCATTAAACAAATACAAACCAGGCACCTGTC
ATATGCCAAGTGATATTCAAAATGGCCCATGTAGACCTTTGTGAAGTATG
TGGCCTAACAGACATTAAACAAATGTCTGTGAAACTGACATAATAAAGTA
AGGTAAGTTATATGTGAGACATTCTCTTTTTATAATAATTCCTGTAAAGC
AGTACTTACTTAGGTAATGATATCATACTGTTTTGTTTTATATTTTTCCT

>POLY-INTRA-17
gttccattccccaatctacagacttcattcagattctacagttgtctcaa
taacaccgttttagcaaaCAGGCAAGCCATTGTCTCCTGGCCCAGGACAG
CCACCCGCTGCCTTTAGCAGGCGCGATACTGACGTTTTGGGGACTCTGAG
CCAGTTGTTTTGGGCAACATCCCTCGGTGCCCATTTTCTCGAGCACCGAT
TCAGGCTGAGGTCATGGCAGGATGGCTCAGAGGtgcccctgggccctgcc
tctgttttattttattgtatttatttattttattttattttattttttga
gatggagtctcgttctgttgcccaggctggagtgcagtggtgggatctcg
gctcactgcaagctctgcctcccgggttcacc
[xxx]
ccattctcctgcctcggcctcccgagtagctgggactacaggcgcccgcc
accacgcctggctaatttttttgtatttttagtagagtcggggtttcaccg
tgttagccaggatggtctccatctcctgacctcatgatccgcccaccttg
gcctcccaaagtgctgggattacaggcgtgagccactgtgcccggccaat
atttgttatttttctatctttttatttAAAAAGATTTCAAATACATAGAA
CAATGAAAAAAAGAAGAGCATGCAGTTTGTGGCCCACCCTCTGGATTTGA
TGGGTGTTGGCGTCTGCACAGTGATCAAGCTCGTTTTAGTGGCTGGGTGG
TATTGCACTAGGGCACACGTGTCTGGGCTCGGtcagttactgaatgaata

>POLY-INTRA-18
TTACTATAATCATTGTTATTTCTGATTCACCGTGCAAT
TCGGCCATTTATATTTATAATGCTATTTGAGAGATTGTGAAAAAGCAACA
CCACCCTGAAAACACACAGTCTTCCTCTAGGCTAGATAGGCTGCAACAGA
TTAGCGCTTTTCCCTCCTTCCttttttttttttttgagacagattcttgc
tcttgttgcccagactggaatgcagtggcacaatttcggttcattgtaac
ctccgcctcccaggttcaagcgattctcctgcctcagc
[xxx]
ctcctgagtagctgggactggcaggcacacgccaccccgcccagctaatg
tttttgtatttttagtagagatgaggtttcttcatgttggccaggctggt
ctcaaactcctgacctcaggtgattcgcccgccttggcctcccaaagttc
tgggattacaggtatgagccacaacacccagccTTTAACGTGCCTTTCTG
ACTCATCTCCTTCTCTCCGTCTTGCCTATCCCACCACCCACTGCACCGTG
GTCTCGGGCTCACTGTGCCACCAACAAGTCTTCCTCTTGGCGCCAGCTTC
TTCTGCTCCTCTGAGCTTCTCCTGCTCCTCTCTGCCTCATCTCCCAAAGC

>POLY-INTRA-19
TGATACCATGTGATATCTTTTAGGGTCAGGAATGTAGTTCTATTTCTTGA
ACTAATTATGGTGGCTAATTTGAATTTGCAGCGTGTAAACAGGCTATTTG
GAATTCAAGCATTTCTCCCTTTCCTTGTCATCTATTTAGATGGTGAATTC
TTCTTTTCCTATTATTTATTATCGAAACACTTTGACCACAAGGATACttt

tttttttttttttgagacggctcgttctgttgcccaggctggagtgcagtg
gtgcgatcttggctcactgcaagctctacctcctgggttcaagtgattt
[xxx]
tcctgcttcagcctcccaagtagctgggattacaggcatgcactaccatg
cgcggctgatttttatatttttagtagagatggggtttcatcatgttggc
taggctggtctccaactccagacctcagatgatccacctgccttggcctc
ctcccaaagtgctgggattacaggcgtgagccactgtgcctagcTTCAAA
aacttttatttgaaatatgggggtacatgtgcagatttgttgcatgggaa
tattgtgtgatgctgaggtttgaagtacagatcccatcacccaggtagtg
agcatagtagccaataggtagtctaacttgtcccATCCGTATCCAGGCTT
TAACCATACTGAGAGAGTGGGAAAAAAACTGAGCTGCCTACTATGACCCA
GTGGAATGGAAAGTTTTCTACCATTTTGGTTCTATGTCATTTTCATATTA
CTTTTTTAAGGAAATTCAATTTTTCTGCCATTCTACTAGACTGTATCTCT
TTGATCACTACCTATGTGTCCAAATCACTGCTTCTCCCAACTTTTTTTGT
GGGCCATCATCCATGAGCATGGTA

>POLY-INTRA-20
CACATAGCTGCTAAGATAAGGTAGGAGAATTTGGCTAGTAAAAGCACTTA
ATGGCCTGAACTTGTGTTTACTAGGGCCTGGGAGAGCAGACAGCTGCCAG
CATACTTAAAATAGGTTGAGTATGAAGTATGCACTTAAGGACTAAAAGAA
TTGCAGAAATTTAAGCAATGACTCAGTTACAAATCCAAGCCCCTTCAGAA
AGTACaggaactatggggaggaatagaaagaacataaacctgcagtgaga
cagacctagttttctttttttttttttttgagatggagtctcactttttt
gcccaggctggagtgcagtggcgcgatcagagctcacagcaacctctgcc
tcctgggttcaagccattcttctgcctcagcctcccaagtagct
[xxx]
gggactacaggcacctgccaccacgcccggctaatttttttgtatttttagtagagatggggtttcatc
gtgttagccaggatggtctcaatctcctgacctcgtgatccgcccacctcagcctcccaaagtgctgg
gattacaggcgtgagccaccctgccgggccGTAAAGATAGATTTAAGACGCCTCCTCCATCCCTAGGA
GGCAACTTTCTTAGGCCTAAACATGCATTCTGGCTCAGGCCTGAAAACGCCTGTATACAGGGATGTTG
GCTCAGAGCCCCTAGAGGGAGCCCTCACTCAACAGTCCAGGTACCAGAAACAGCTTGTTGGAAGGGGA
AATGTAAATTAAGGTGTCATGCAATGGTGGTCTTTCCCCCAGGAATGATCAGTGTATACGGCCAAAAT
TAAGCCCTTTTTGAGCGTCCTTCCAATAGAAATTAGGATCACGAG
